# Did dietary change drive natural selection? A paleo-empirical evaluation across 6,000 years in Britain

**DOI:** 10.1101/2025.06.26.661798

**Authors:** Zexuan Chen, Andrew Millard, Eva Fernandez

## Abstract

Multiple genetic variants associated with diet-related traits show strong signatures of natural selection^1–17^. To test whether these signals were indeed diet-driven, we conducted an empirical investigation. We compiled an isotopic dataset comprising 6,064 ancient human samples and 5,635 food resource samples from Britain. We developed a Bayesian mixing model to estimate individual dietary proportions based on isotopic data and subsequently constructed a temporal dietary model. A dairy-use time series was also constructed^18^. Using 1,038 ancient DNA samples, we reconstructed derived allele frequency trajectories for 14 strongly selected single nucleotide polymorphisms (SNPs) via bootstrap resampling^19^. A generalized additive model (GAM) was then applied to estimate mean and time-varying selection coefficients, while accounting for evolutionary forces beyond selection. Finally, we applied the convergent cross mapping (CCM)^20^ algorithm for causal discovery between the time-varying selection coefficients and their corresponding dietary variables. Our findings indicate that C_3_ plant consumption drove selection on rs12401678 and rs653178, and dairy consumption on rs4988235. Selection signals at rs174570 and rs174594 are likely linked to marine fish and terrestrial meat intake, whereas the remaining SNPs show more complex selection dynamics in which any diet-related signal cannot be clearly identified. Our results underscore the complexity of natural selection at the genetic level and highlight the need for more careful evaluation when identifying its potential drivers.

## Main

As one of the major environmental factors to which humans have had to adapt, diet has long been discussed for its central role in human evolution^21–31^. With the advent of the genomic era, its evolutionary impact has been increasingly investigated from a genomic perspective^1–17^. Such studies have generally followed two principal lines of research. The first approach examines variation in the frequencies of genetic variants, including single nucleotide polymorphisms (SNPs) and copy number variations (CNVs), among populations with distinct dietary practices or across different time periods, particularly following the emergence of agriculture^12,13,15–17^. For instance, populations with high-starch diets tend to carry more copies of the amylase genes (AMY1, AMY2A, AMY2B) than those with low-starch diets, suggesting possible adaptation to different diets^13,17^. Ancient genomic data further provide a temporal dimension, revealing that haplotypes with higher copy numbers of amylase genes increased rapidly in frequency over the past 12,000 years in West Eurasia^15^, particularly among European farmers during the last 4,000 years^16^. The second approach combines modern or ancient genomic data with various statistical methods to detect signals of natural selection at loci associated with diet-related phenotypes^1–11,14^, such as fatty acid metabolism, vitamin D levels, and lactase persistence. Collectively, these findings reinforce the view that human evolution has been driven by natural selection responding to dietary change^1,2,5^.

The studies mentioned above represent qualitative analyses from a genomic perspective. Although they provide some evidence supporting diet-driven selection, no direct empirical evidence has demonstrated a causal link between dietary variables and natural selection coefficients. After all, qualitative associations do not necessarily imply causation^32^. Evershed et al. (2022)^18^ combined ancient genomic data with isotopic evidence of dairy use from pottery sherds and applied a likelihood comparison framework, showing that a model of selection driven by dairy consumption did not fit the data better than a constant selection model. Although the likelihood comparison approach is not a causal discovery or inference method, the idea of this study is nevertheless highly informative. Building on this inspiration, we developed an interdisciplinary empirical framework to provide direct causal evidence addressing whether dietary change acted as a driver of natural selection in humans (Fig. 1). To this end, we integrated large-scale ancient isotopic and genomic datasets from Britain (Extended Data Fig. 1) with a Bayesian mixing model, bootstrap resampling^19^, generalized additive modeling (GAM), and convergent cross mapping (CCM)^20^ in a unified analytical framework. Five food resource groups were modeled—terrestrial herbivores, terrestrial omnivores, C_3_ plants, marine, and freshwater fish. Fourteen diet-related SNPs under strong selection^1,2,5,8^ were analyzed, each paired with its putative dietary driver (Extended Data Table 1; see Supplementary Information Section 1, SI 1, for matching criteria). The following sections present (i) the reconstructed dietary patterns in ancient Britain, (ii) the temporal dynamics of derived allele frequencies and selection coefficients, and (iii) the causal discovery results and their interpretation.

**Fig. 1.**
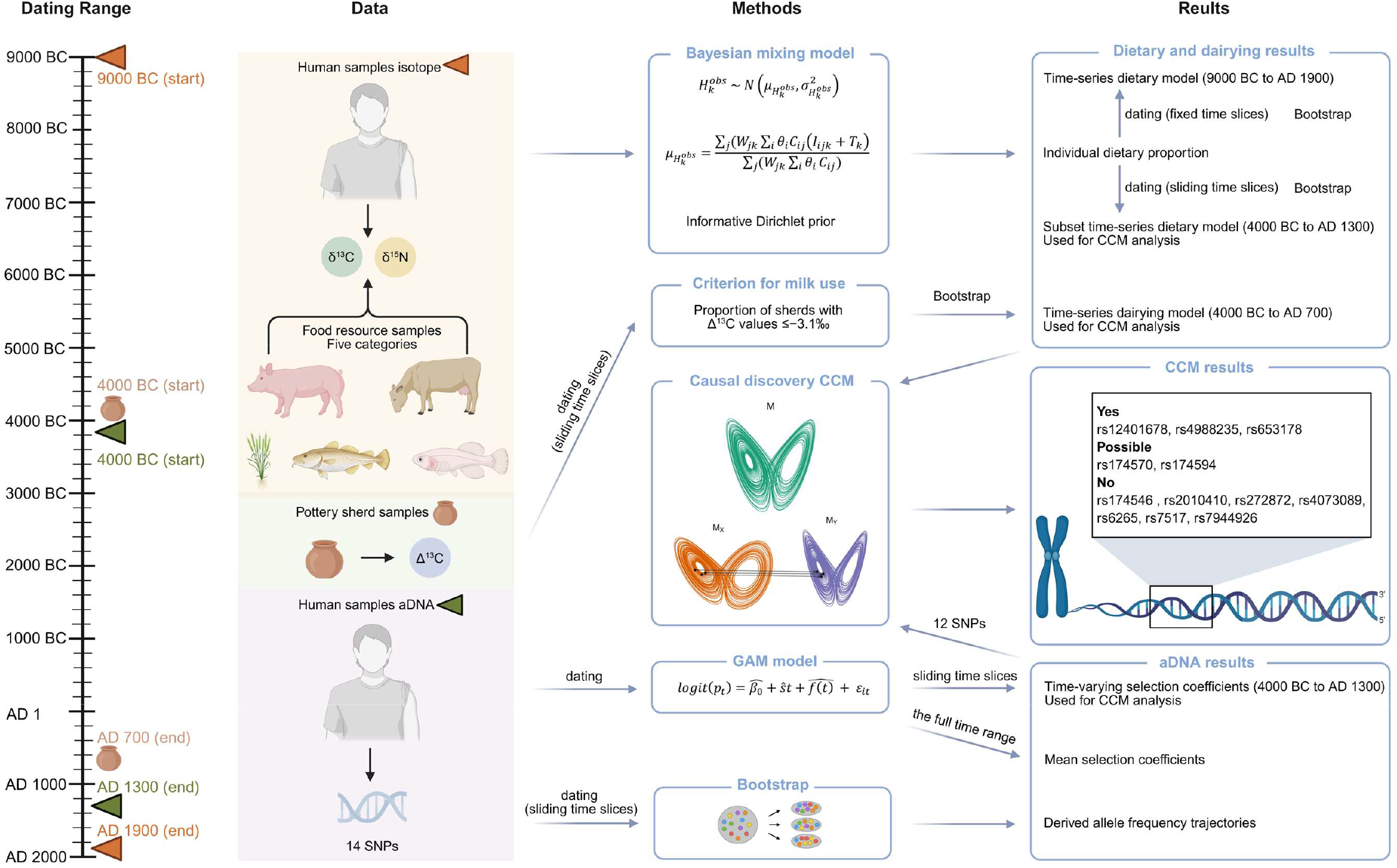
Overview of the study design.

### Ancient British diet

The time-series dietary model for ancient Britain is shown in Fig. 2. Due to the limited sample size and broad chronological distribution of the Paleolithic specimens, they were not included in the time-series modeling. Posterior estimates for each Paleolithic individual are provided in SI 2A.

**Fig. 2.**
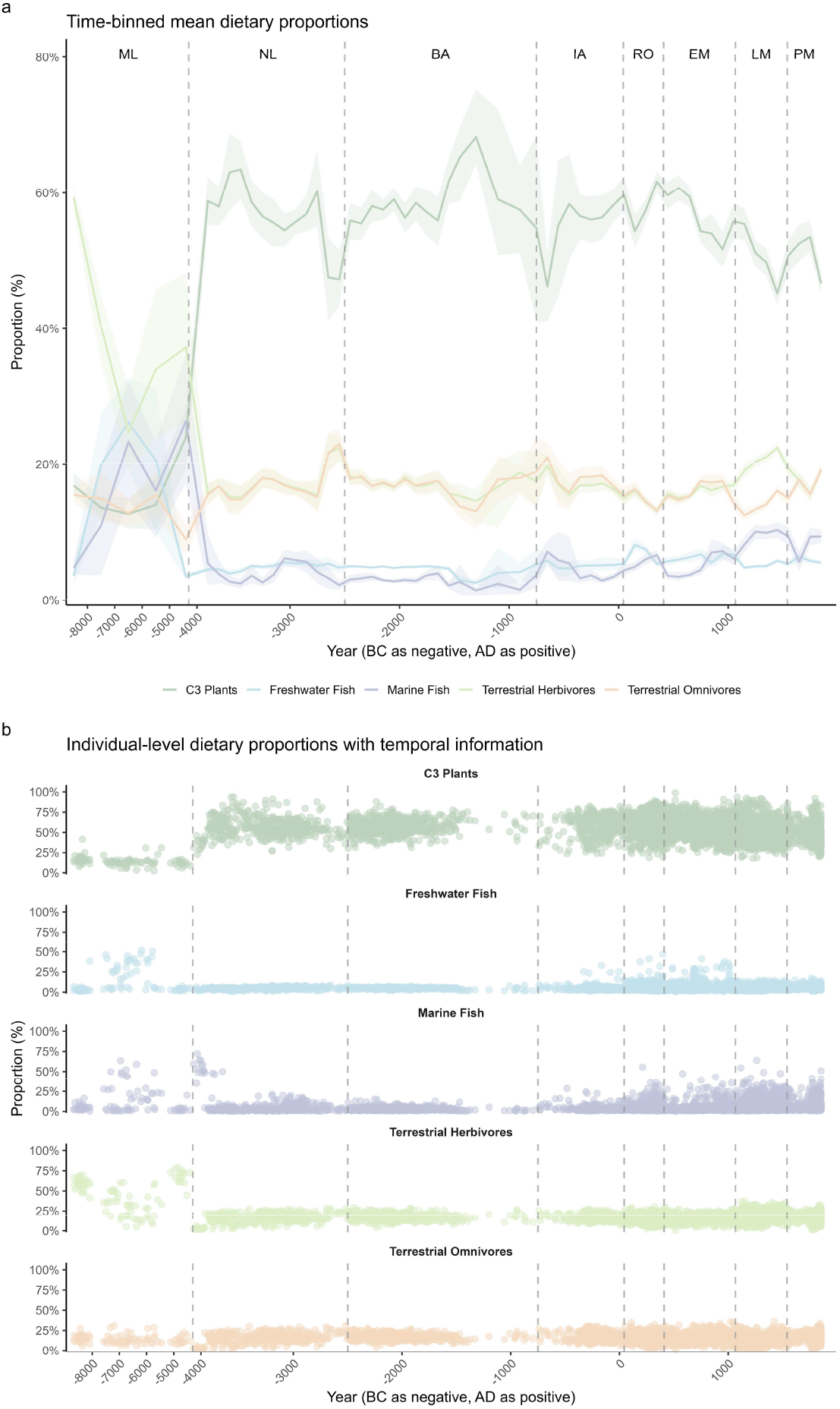
**a**, Mean proportions of each food group within each time slice. Lines connect mean proportion estimates plotted at the midpoint of each time slice. Shaded bands represent 95% confidence intervals. Both the mean estimates and confidence intervals were estimated from the posterior mean dietary proportions of individuals assigned to each slice using nonparametric bootstrap resampling (1,000 iterations). **b**, Posterior mean dietary proportions of all individuals. Abbreviations: ML – Mesolithic; NL – Neolithic; BA – Bronze Age; IA – Iron Age; RO – Roman period/Roman Iron Age; EM – Early Medieval period; LM – Later Medieval period; PM – Post-Medieval period.

The Mesolithic diet followed a hunter-gatherer subsistence strategy, with individuals relying primarily on locally available resources^33^. As a result, there was substantial dietary variation across Britain. As shown in Fig. 2b, individuals from this period exhibit marked heterogeneity in their diets. Inland populations primarily consumed terrestrial herbivores, whereas individuals from specific regions, such as Doggerland^34^ and southern Wales^35–37^, exhibited substantial intake of freshwater and marine fish, respectively (SI 2B).

With the onset of the Neolithic period, diet underwent a marked shift, characterized by a substantial increase in C_3_ plants and domesticated animals, reflecting the introduction of agriculture^38^. Concurrently, the consumption of marine and freshwater fish declined substantially. From around 3300 to 2500 BC, there is a slight decline in the proportion of plant-based foods in the diet, accompanied by a modest increase in terrestrial animal protein consumption (Extended Data Fig. 2). This trend broadly supports the view that agriculture declined in certain regions of Britain during this time, with some populations shifting towards a more pastoral lifestyle^39–41^. However, the modest magnitude of this shift is more consistent with localized agricultural decline, rather than a widespread phenomenon across Britain^42^ (SI 2C). Moreover, there is an increase in the consumption of plant resources following the arrival of the Beaker culture (2500 BC), which seems to contradict the previous view that agriculture did not recover until around 1500 BC^40^ (Extended Data Fig. 3).

From the beginning of the Bronze Age, dietary patterns appear to have remained relatively stable for nearly two millennia^43–49^. Diets continued to rely heavily on terrestrial resources, with minimal consumption of marine or freshwater foods (SI 2D). The onset of the Iron Age marked a slight departure from this trend (Fig. 2b), with increasing dietary heterogeneity and a modest rise in the use of aquatic resources (SI 2E). This shift became more pronounced during the Roman period, with a greater number of individuals consuming marine and freshwater fish (Fig. 2b)—an effect that is clearly reflected in the increase in average aquatic resource consumption across Britain (Extended Data Fig. 4). During this period, marine fish were particularly favored by the elite^50,51^, and dietary differences among individuals became more evident (SI 2F).

Marine resource consumption declined during the early phase of the Early Medieval period, suggesting that the infrastructure which had supported the supply of marine foods and the sea during the Roman era had largely collapsed^52^. However, in the subsequent Viking period (around AD 800), certain individuals with Viking affiliations^53–56^ exhibited substantial reliance on marine foods (SI 2G), which contributed to an increase in average marine resource consumption across Britain (Fig. 2a; Extended Data Fig. 5).

Marine resource consumption increased substantially during the Later Medieval period (Extended Data Fig. 5). The rise in fish consumption coincided with population growth, the expansion of the fish trade, and the widespread adoption of Christian fasting practices in Britain^52^. However, not all individuals had a high intake of marine fish (Fig. 2b). Our models reveal considerable dietary heterogeneity across different sites in the Later Medieval period. While some sites exhibit marked reliance on marine fish (e.g., Orkney^46,53,54^, Portmahomack^57,58^, and York^52^), a substantial number of others show little to no evidence of such consumption (SI 2H). Freshwater resource consumption remained stable before the Early Medieval period but peaked during the second and third centuries AD (Extended Data Fig. 4). During the Early Medieval period, it increased gradually, followed by a sharp decline in the Later Medieval period (Fig. 2; Extended Data Fig. 5). This pattern is consistent with the conclusion that freshwater fish played only a limited role in the diet during the Later Medieval period. Nevertheless, a notable proportion of freshwater resource intake is still observed among individuals from more remote regions^59^ (e.g., those from Wharram Percy^60^) (SI 2H).

In the Post-Medieval period, marine resource consumption appears to have declined during the 17th century (Extended Data Fig. 6). However, our modeling results indicate that, at least for samples dating to the 18th and 19th centuries, the proportion of marine resource consumption was comparable to that observed during the Later Medieval period (Extended Data Fig. 6). In fact, substantial marine intake persisted in major urban centers during the 18th and 19th centuries, such as in York^52^ and London^61–64^ (SI 2I). Thus, despite the waning influence of Christian fasting practices^52^, current data do not indicate a marked decline in marine resource consumption during the Post-Medieval period.

In addition to the dietary time-series model used to illustrate dietary trends, we constructed a separate model for causal discovery, which included dairy data^18^ and used the same temporal range and time bins as those applied to estimating time-varying selection coefficients. The results reveal gradual dietary shifts, with increasing freshwater and marine fish after 1000 BC, a slight decline in terrestrial resources, and a steady decrease in dairy (Extended Data Fig. 7; SI 2J).

### Allele frequency dynamics and the landscape of natural selection in Britain

The constructed trajectories of derived allele frequencies are shown in Fig. 3 (see also SI 3A). Although previous studies suggested strong selection on these SNPs over the past 10,000 years^1–8^, data from Britain between 4000 BC and AD 1300 indicate that such selection was not consistently strong for some loci. Several loci (rs12401678, rs2010410, rs3891176, rs4073089, rs7517, and rs7944926) exhibit only minor fluctuations without any clear directional change (Fig. 3). In a broader European context^5^ (Extended Data Fig. 8), all these SNPs, except rs3891176 and rs7944926, exhibit a comparable pattern of limited frequency change during this period, with major shifts occurring earlier (8000–3500 BC). For rs3891176, allele frequencies show a consistently strong upward trend in Europe, particularly after 500 BC, whereas the British data display an atypically stable trajectory (Fig. 3; Extended Data Fig. 8). For rs7944926, both British and European data show an initial rise followed by a decline. However, European frequencies peaked higher around 3500 BC (∼45% vs. ∼35% in Britain), producing a clearer downward trend (Extended Data Fig. 8). In contrast, the British trajectory rises and then falls back to approximately its initial level (Fig. 3).

**Fig. 3.**
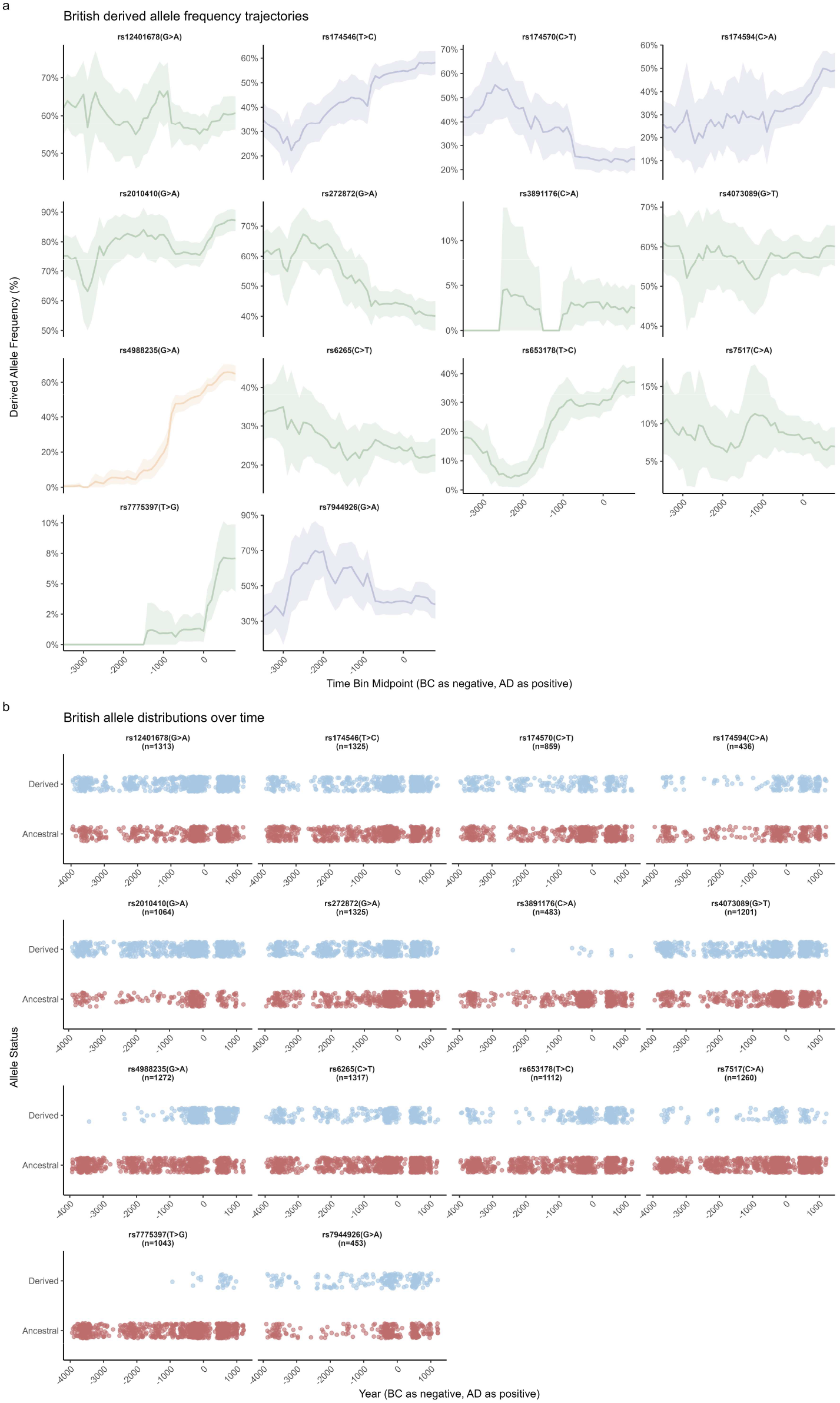
Derived allele frequency trajectories and allele distributions over time. **a**, Derived allele frequency trajectories constructed by estimating the mean and 95% confidence intervals of derived allele frequencies within each time bin using bootstrap resampling. SNPs associated with marine fish, C_3_ plants, and dairy are highlighted in blue, green, and yellow, respectively. rs174546, rs174570, and rs174594 are also associated with terrestrial animal consumption. **b**, Temporal distributions of ancestral and derived alleles.

SNPs rs174546, rs174594, rs4988235, rs653178, and rs7775397 exhibit clear upward trends in derived allele frequency, closely mirroring patterns observed across Europe^5^. Among them, rs174546, rs174594, and rs653178 display nearly identical trajectories compared with those observed in Europe (Fig. 3; Extended Data Fig. 8). rs4988235 and rs7775397 also show similar upward trends, but the frequency increase is notably stronger in Britain than in Europe. The derived allele of rs4988235 reaches ∼60% in the final time bin in Britain, compared with ∼30% in Europe (Fig. 3; Extended Data Fig. 8). Given that the European dataset includes British samples, the frequency elsewhere must be even lower, suggesting that the rapid rise of the lactase persistence allele was likely driven primarily by the British population. This interpretation agrees with previous estimates showing stronger selection for lactase persistence in Britain (selection coefficient: 1–6.3%) than in continental Eurasia (1.1–1.7%)^18^. The limited British data in that study yielded a broad estimated range, whereas our larger dataset provides stronger support for this pattern. A similar pattern is observed for rs7775397, whose derived allele frequency in Britain reaches ∼8% in the final time bin (Fig. 3), nearly twice that of Europe (Extended Data Fig. 8).

In contrast, SNPs rs174570, rs272872, and rs6265 show clear decreases in derived allele frequency over time. For rs174570 and rs272872, the magnitude and pattern of decline closely match those observed across Europe (Fig. 3; Extended Data Fig. 8). rs6265 differs slightly: in Britain, its derived allele frequency was higher around 3500 BC and declined gradually thereafter (Fig. 3), whereas in Europe it dropped sharply between 3500 and 3000 BC before stabilizing (Extended Data Fig. 8).

The estimated selection coefficients are shown in Fig. 4. Average estimates closely mirror the overall trends in frequency trajectories (Fig. 3; Fig. 4a). SNPs with minimal frequency changes have mean coefficients near zero, whereas those with clear upward or downward trajectories exhibit correspondingly positive or negative estimates. The strongest signal of positive selection remains at rs4988235, with a mean selection coefficient of 6.01%, higher than the corresponding estimate for Europe (4.21%)^5^. Although the GAM model incorporates a smooth term accounting for evolutionary forces beyond selection, the derived allele frequencies remain strongly correlated with the estimated selection coefficients, suggesting that natural selection is the dominant driver of allele frequency change (see SI 3B).

**Fig. 4.**
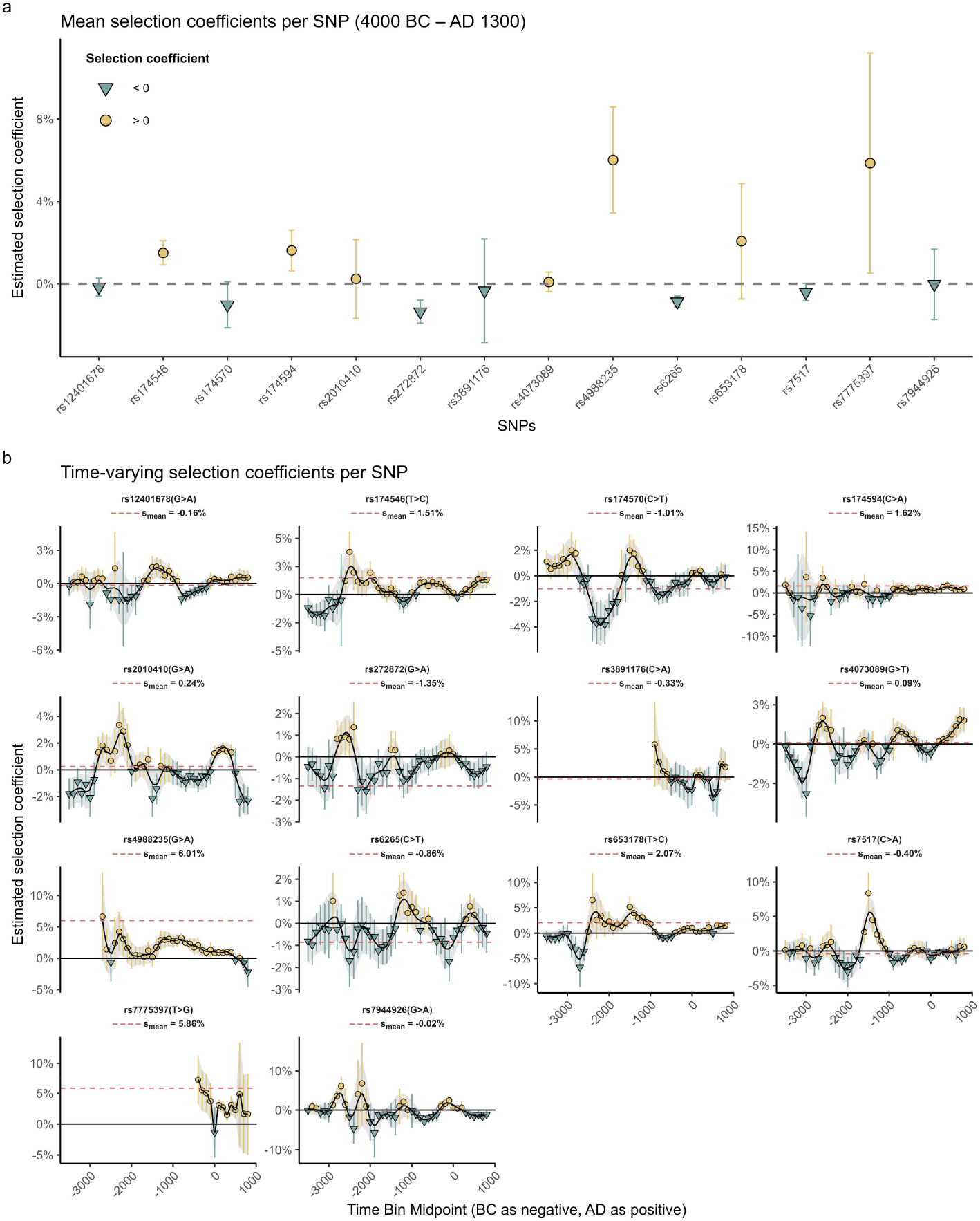
The landscape of natural selection. **a**, Mean selection coefficients estimated by fitting the GAM model across the full time span. **b**, Time-varying selection coefficients estimated by fitting the GAM model within each time bin. Red dashed lines indicate the point estimate of mean selection coefficient. Error bars represent the standard error. Black curves represent the temporal trend of the selection coefficients, with the shaded areas indicating the corresponding smoothed upper and lower standard-error bounds.

Time-varying coefficients (Fig. 4b) reveal that no SNP maintained consistently positive or negative selection across all time intervals. Thus, while some loci appear under overall positive, negative, or neutral selection across the 5,300-year span, their selection landscapes varied markedly through time. SNPs with average coefficients near zero (rs12401678, rs2010410, rs3891176, rs4073089, rs7517, rs7944926) oscillate between positive and negative selection, producing an overall signal of neutrality. Conversely, loci with clear long-term trends in frequency trajectories show more intervals with coefficients acting in the same direction (e.g., positive—rs174546, rs174594, rs4988235, rs653178, rs7775397; negative—rs174570, rs272872, rs6265). This suggests that the strength and direction of natural selection are temporally dynamic, highlighting that the landscape of selection may be far more complex than previously assumed. Even loci with strong mean selection signals^1–3^ may have undergone periods of weak or even reversed selection.

Model validation using fitted GAM simulations reproduced the observed trajectories well (SI 3B), confirming the reliability of our estimates, except for rs3891176 and rs7775397. Consequently, these two SNPs were excluded from subsequent causal analyses (see SI 3C).

### Causal discovery with convergent cross mapping

Fig. 5, Extended Data Fig. 9, and Extended Data Table 2 show the results of causal discovery using CCM^20^. SNPs rs12401678, rs4988235, and rs653178 exhibited strong indicators of causal relationships, meeting both criteria of high ρ_Lmax_ values (>0.25) and convergent trends in the coefficient (|Δρ_last 5L_| **<** 0.02) (Fig. 5a, h, j; Extended Data Table 2). SNPs rs174570 and rs174594 also show signs of potential causal relationships (Fig. 5c, d; Extended Data Fig. 9b, c; Extended Data Table 2). Although their ρ values do not fully converge, their ρ_Lmax_ values remain substantially above zero. In particular, rs174570 already shows a rather strong convergence trend (Fig. 5c; Extended Data Fig. 9b). Therefore, it is plausible that natural selection at these two SNPs was also influenced by diet. The remaining SNPs show no evidence of causality. Their ρ values either approach zero (rs174546 for meat consumption; rs2010410, rs272872, rs4073089, rs6265, rs7517) (Extended Data Fig. 9a; Fig. 5e, f, g, i, k; Extended Data Table 2) or fall below zero and continue to decline at the maximum library length (rs174546 for marine fish consumption; rs7944926) (Fig. 5b, l; Extended Data Table 2).

**Fig. 5.**
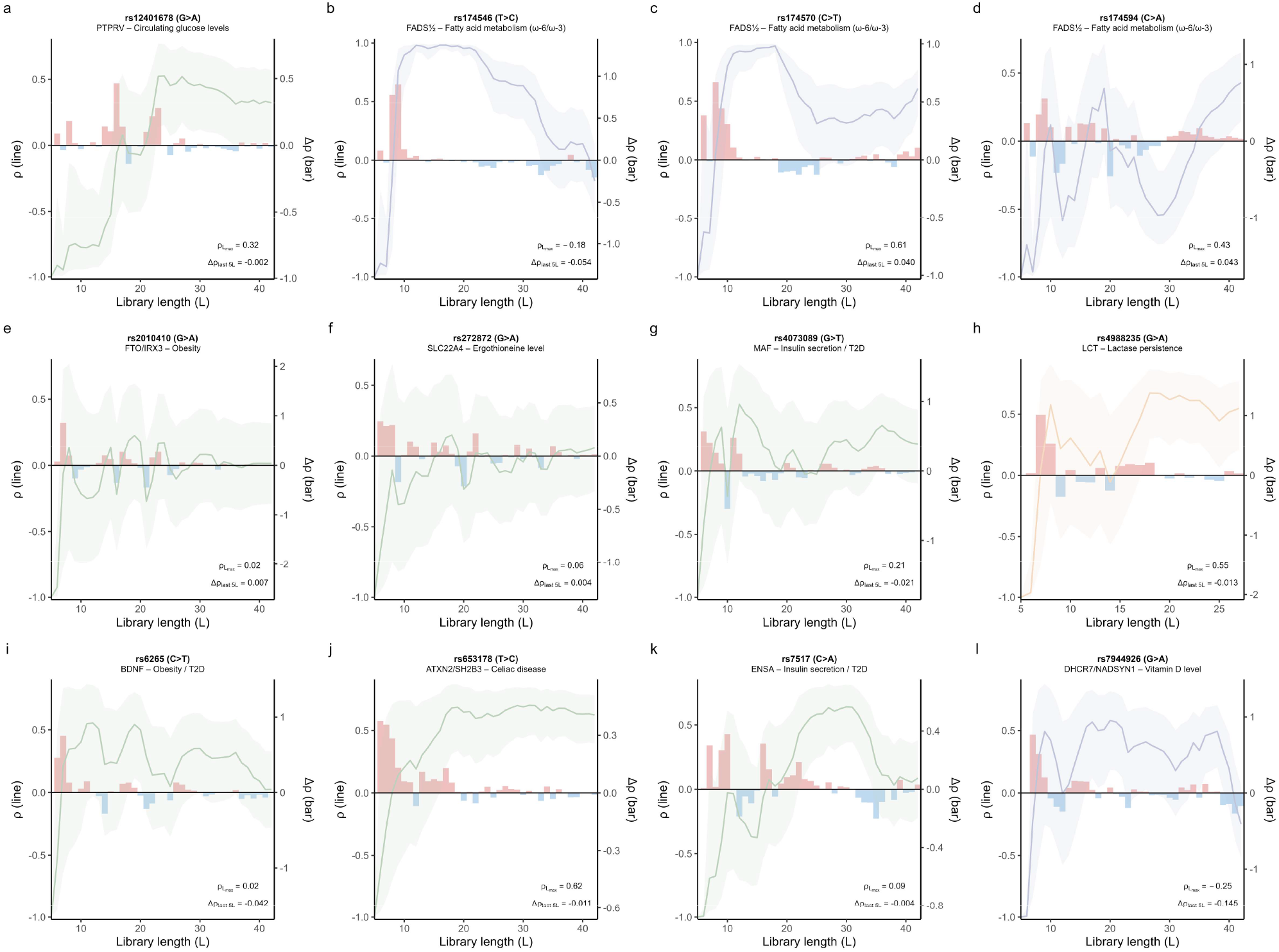
CCM results for 12 SNPs. **a-l**, CCM outputs for SNPs rs12401678 to rs7944926 (ordered left to right, top to bottom). As the library length increases, convergence of the coefficient (ρ) above zero indicates causality, with larger values denoting stronger effects. Stepwise changes are given by Δρ(L) = ρ(L) − ρ(L – 1). ρ_Lmax_ is the ρ at the final L, and Δρ_last 5L_ is the mean Δρ across the last five L steps. Evidence for causality was discovered when ρ_Lmax_ >0.25 and |Δρ_last 5L_| < 0.02. Line: ρ; Shade: 95% CI; Bars: Δρ. CCM results are based on the point estimates of dietary variables and selection coefficients. SNPs associated with marine-based diets, C_3_ plant consumption, and dairy are highlighted in blue, green, and yellow, respectively. rs174546, rs174570, and rs174594 are also associated with terrestrial animal consumption. The key parameters: the embedding dimension E=3, the time delay τ=1, and the number of nearest neighbors k=4 for all CCM tests.

Overall, rs12401678 (C_3_ plants), rs4988235 (dairy), and rs653178 (C_3_ plants) appear to be the strongest candidates for diet-driven selection. SNPs whose selection potentially exhibits causal relationships with diet include rs174570 (meat or marine fish) and rs174594 (meat or marine fish). Taken together, these patterns suggest that C_3_ plant consumption, dairy use, and the intake of meat and marine fish may all have contributed to natural selection at specific human genetic loci. Notably, although its robustness is somewhat lower (see SI 4A), our findings for rs4988235 differ from those of previous studies. Evershed et al. (2022)^18^, using a maximum likelihood comparison approach, concluded that a milk-driven selection model did not outperform a constant selection model. However, the likelihood comparison approach is not a traditional causal analysis method. Moreover, their model assumes a specific functional relationship between milk consumption and the selection coefficient, which is unlikely to reflect the true underlying dynamics. In addition, other important evolutionary forces were not incorporated into their modeling framework. In our study, the estimation of selection coefficients incorporated a smooth term to account for additional evolutionary forces. Its effectiveness has been validated through our reverse simulations (SI 3B). Furthermore, CCM does not impose any predefined functional relationship between variables but instead infers causality by reconstructing the underlying state space^20,65,66^, making it particularly well-suited for complex and nonlinear systems. However, due to the scarcity of ancient data and the inherent complexity of the research topic, we cannot claim that our conclusion is closer to the truth. Rather, our findings should be interpreted as the outcome from the currently available data and the specific analytical framework.

For those SNPs that did not exhibit causal signals (rs174546, rs2010410, rs272872, rs4073089, rs6265, rs7517, rs7944926), two possible explanations can be considered. First, diet may indeed not have been a driver of selection at these loci. Second, diet could have been a contributing factor, but its signal potentially obscured by other confounding factors. The second explanation is based on the fact that strong noise interference can distort the outputs of CCM^67^. Due to the scarcity of ancient data and complexity of our case, it is not feasible to control these confounders and estimate causal effects in the same way as in econometrics or epidemiology^68–79^ (see SI 4B). Therefore, for SNPs that did not exhibit causal signals, both of the aforementioned explanations are plausible. A more detailed discussion will be provided in the next section.

Since both the dietary data and selection coefficients are estimated values, they inherently carry uncertainty. Therefore, in addition to conducting CCM based on point estimates, we generated 100 simulated values for dietary proportions and selection coefficients of each time slice based on their probability distributions. This allowed us to construct 100 independent simulated time series for the selection coefficient of each SNP and its corresponding dietary proportions. We then performed 100 separate CCM tests per SNP using these simulations to test the robustness. The results remain highly consistent across simulations, indicating that our CCM findings are relatively robust (see SI 4A).

## Discussion

Of the C_3_ plant-associated SNPs, rs12401678 and rs653178 display clear signatures of diet-driven selection, whereas rs2010410, rs272872, rs4073089, rs6265, and rs7517 show no such signals. rs2010410 is primarily associated with obesity-related traits, particularly body mass index (BMI). Although high-carbohydrate diets may contribute to obesity, the trait is influenced by many other factors^80,81^, such as physical activity and vegetable intake. Consequently, any causal signal attributable to C_3_ plant consumption at this locus may have been overwhelmed by substantial background noise. rs272872 is associated with circulating ergothioneine levels. Ergothioneine occurs primarily in fungi^82^, whereas C_3_ plants contain only minimal amounts. As C_3_ plants are not a major dietary source of ergothioneine, the absence of a causal signal is unsurprising. Although rs6265 has been associated with type 2 diabetes (T2D)^81,83,84^, it is primarily linked to the brain-derived neurotrophic factor (BDNF) gene^2^. Given the multifactorial etiology of T2D^85–87^ and the indirect influence of rs6265 on it, any causal signal linked to C_3_ plant consumption was likely masked by noise.

The other T2D-associated genes (rs4073089 and rs7517) display stronger results than rs6265 (Fig. 5g, i, k), although their CCM outputs still do not meet our criteria for a causal signal. The stronger results are likely because they are related to insulin secretion, making their connection to glucose levels and T2D more direct. Nonetheless, insulin secretion is not directly equivalent to plasma glucose levels or T2D—the phenotypes on which natural selection acts directly. Consequently, the causal signals at these loci may also be partially attenuated. In contrast, traits such as circulating glucose levels (rs12401678) and celiac disease susceptibility (rs653178) represent phenotypes that are subject to direct selection pressures. These traits are shaped mainly by C_3_ plant consumption and are less affected by other dietary or environmental factors, making potential causal relationships easier to detect. While these traits are also subject to additional influences, the performance of CCM remains robust as long as the noise is not excessively strong.

This line of reasoning likewise applies to SNPs associated with the consumption of marine fish, meat, and dairy. The SNPs associated with meat and marine fish consumption that exhibited possible causal signals were those involved in fatty acid metabolism (rs174570, rs174594)^6,8^. The SNP lacking such signals is linked to vitamin D levels (rs7944926)^1^. Vitamin D levels are influenced by a broader range of factors^88,89^, including mushrooms and sunlight exposure. In contrast, the main sources of ω-3 and ω-6 fatty acids are relatively fixed, making any causal signals easier to detect. ω-3 fatty acids are obtained primarily from marine fish^90^. Although flaxseeds, algae, walnuts, and soybeans also contain ω-3 fatty acids^91^, these foods are generally consumed only as minor dietary components. Similarly, most ω-6 fatty acids are derived from terrestrial animal fat^92^. While plant oils and nuts contain substantial amounts of ω-6 fatty acids^92^, their consumption in ancient populations was relatively low^93^, making them a minor contributor to overall ω-6 intake. Because fatty acid metabolism involves multiple dietary sources (marine fish and terrestrial meat), analyzing any single dietary variable introduces noise. This likely explains why the CCM results for rs174570 and rs174594, although significantly above zero, did not converge and can only be considered possible causal signals (Fig. 5c, d; Extended Data Fig. 9b, c). SNP related to dairy consumption (rs4988235)^1,18^ requires little elaboration, as lactase persistence is a trait that is mainly determined by genetic factors and dairy consumption. A particularly noteworthy exception is rs174546^1^. Although this SNP is also involved in fatty acid metabolism, it showed no causal signal. Its selection profile and frequency trajectory closely resemble those of rs174594 (Fig. 3; Fig. 4), yet the CCM results differ markedly between these two SNPs, a discrepancy that requires additional data and approaches to validate.

The above analyses indicate that SNPs showing diet-driven selection signals share two features: their phenotypes are shaped mainly by a single dietary factor and are direct targets of natural selection. Natural selection acts directly on phenotypes^94^ and only indirectly on genotypes. When a SNP is strongly associated with a phenotype that is influenced primarily by a single dietary factor and is directly targeted by natural selection, it is reasonable to expect that individuals with different phenotypes will exhibit differential fitness under dietary selection pressures. This, in turn, can result in detectable signals of diet-driven selection at the genetic level. In contrast, if the phenotype of a SNP is not directly targeted by natural selection and is shaped by multiple environmental factors, the selective forces acting at the genetic level may be more difficult to attribute. Overall, our results show that C_3_ plant consumption drove the natural selection acting on rs12401678 and rs653178, and that dairy consumption drove the selection on rs4988235. The selection signals for rs174570 and rs174594 are possibly explained by the consumption of marine fish and terrestrial meat. The remaining SNPs were likely shaped by more complex selective processes, whose specific drivers cannot be resolved within the current empirical framework (see SI 5 for limitations of this study). Our results highlight the complexity of natural selection. For many diet-related SNPs, natural selection likely reflects the combined effects of multiple environmental factors rather than diet alone. It is insufficient to determine whether loci were truly diet-driven based solely on their selection signals and associated diet-related traits. The underlying selective mechanisms at these loci remain complex and warrant further investigation.

## Methods

### Data

#### Ancient isotopic data

We collected isotopic data (δ^13^C and δ^15^N) from human remains^34–37,43–45,47–54,56,57,59– 64,95–162^ and potential dietary sources^34,35,37,41,46–49,51,52,54,55,57– 59,61,95,96,99,102,103,105,107,109,112–115,117,119–124,135,137,140–143,147,148,152,154,156,163–192^ spanning the past 30,000 years in Britain (Supplementary Data 1). In total, the dataset includes 11,699 samples: 6,064 ancient British human samples, 3,810 ancient British faunal and plant remains, 1,565 modern British faunal and plant remains, and 260 contemporary British soil samples. The associated archaeological information for each sample (including, but not limited to, sample ID, site location, and dating information^193–202^) is also provided (see SI 6A and Supplementary Data 1). These samples collectively encompass nearly all available isotopic information relevant to ancient British diets. Pottery sherd data on dairy product use in Britain were also incorporated (2022)^18^.

#### Ancient DNA data

We collected ancient DNA (aDNA) data from 1,038 British samples based on previously published studies^203–216^. Relevant publications were identified through the Allen Ancient DNA Resource (AADR v62.0)^217,218^. All samples were aligned to the GRCh37/hg19 reference genome coordinates.

For samples with available BAM files (obtained from the European Nucleotide Archive or via direct communication with the corresponding authors), we used SAMtools^219^ (v1.12) to generate mpileup data at the positions of our target SNPs, using default quality thresholds. Diploid genotypes were called at sites with a read depth of at least two to maximize data usage; for sites covered by a single read, only one allele was called.

For a small subset of samples lacking BAM files^220^, we processed raw FASTQ data by aligning the reads to the GRCh37 reference genome (NCBI RefSeq assembly: GCF_000001405.13) using the Burrows-Wheeler Aligner^221^ (BWA aln, v0.7.17, r1188) with the following parameters: base quality ≥15, seed disabled, mismatch penalty (-n) set to 0.01, and gap open penalty (-o) set to 2, as recommended for aDNA analysis^222^. The resulting alignments were then sorted, filtered for mapping quality ≥20 (-F 4 -q 20), and deduplicated (-F 1024) using SAMtools. Genotypes were subsequently called from these BAM files following the same procedure described above (Supplementary Data 5 and Supplementary Data 6). All analyses were performed within the nf-core/eager^223^ (v2.5.1) workflow. Genotype calling was carried out in R.

### Bayesian estimation for ancient diet

#### Dietary reconstruction for individuals

We used the likelihood function provided by Food Reconstruction Using Isotopic Transferred Signals (FRUITS) ^224^, with modifications to the prior specifications of key parameters to ensure estimates that better reflect realistic dietary proportions.

The construction of the likelihood function, along with the specification of parameter priors, is detailed as follows. The observed isotopic values for each human individual were assumed to follow a normal distribution:

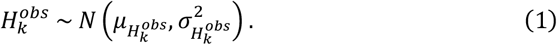

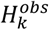 represents the observed isotopic value of the *k*-th dietary proxy (δ^13^C or δ^15^N in our case) for an individual. It is assumed to vary around a mean value 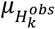 with a measurement error 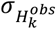 . The measurement error is empirically set to 0.5 for both δ^13^C and δ^15^N, and treated as a known parameter in the model.

The mean value 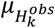 is modeled as a function of other parameters, as specified by the following equation:

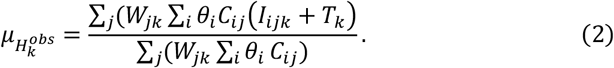

By combining Equations (1) and (2), we obtained the full likelihood function.

*W*_*jk*_ represents the weight contribution of the *j*-th food fraction — protein or energy (i.e., lipids and carbohydrates), to the *k*-th dietary isotope proxy. It is assumed to follow a normal distribution: 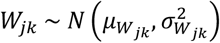 . In our model, the prior distributions for *W*_*jk*_ are specified as follows, based on evidence from animal feeding experiments^225–229^ and previous statistical analysis^230^: 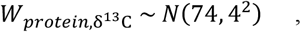 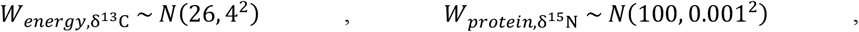 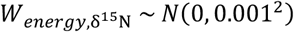.

*C*_*ij*_ denotes the concentration of the *j*-th food fraction in the *i*-th food group. In our case, the food groups include terrestrial herbivores (H), terrestrial omnivores (O), C3 plants (P), marine fish (M), and freshwater fish (F). Each *C*_*ij*_ is modeled as a normally distributed variable: 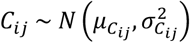 . Based on empirical evidence from previous studies^100,231^ and data from the Food and Agriculture Organization of the United Nations (FAO)^232^, the prior distributions of food fraction concentrations for each species were defined as follows: *C*_*H,protein*_ ∼ *N*(60, 5^2^) , *C*_*H,energy*_ ∼ *N*(40, 5^2^) , *C*_*O,protein*_ ∼ *N*(60, 5^2x^) , *C*_*O,energy*_ ∼ *N*(40, 5^2^) , *C*_*P,protein*_ ∼ *N*(10, 5^2^) , *C*_*P,energy*_ ∼ *N*(90, 5^2^) , *C*_*M,protein*_ ∼ *N*(75, 5^2^) , *C*_*M,energy*_ ∼ *N*(25, 5^2^) , *C*_*F,protein*_ ∼ *N*(65, 5^2^), *C*_*F,energy*_ ∼ *N*(35, 5^2^).

*θ*_*i*_ represents the dietary proportion of the *i*-th food group, which is the primary parameter of interest in our analysis. We assumed that these proportions follow a Dirichlet distribution. For each individual, we manually specified informative Dirichlet priors based on prior knowledge from previous studies and archaeological evidence. For example, if earlier research indicates that an individual from the post-agricultural period primarily consumed terrestrial resources, with moderate animal protein intake and little to no evidence of fish remains at the corresponding site, the informative Dirichlet prior would be set as (1.5, 1.5, 6, 0.5, 0.5), representing herbivores, omnivores, C3 plants, marine fish, and freshwater fish, respectively (see Supplementary Data 3 for the informative Dirichlet priors assigned to each individual). Although previous studies have suggested using logical constraints^224^ as priors for *θ*_i_, we found such priors to be overly restrictive and prone to causing sampling failures in the Markov Chain Monte Carlo (MCMC) process. In contrast, the informative Dirichlet prior provides a more flexible and reliable alternative.

*I*_*ijk*_ denotes the observed isotopic value of the *k*-th dietary proxy measured from the *i*-th food group and the *j*-th food fraction. It is modeled as a normally distributed variable: 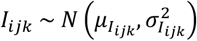 . The prior distribution of each *I*_*ijk*_ is derived from the collected isotopic data of food resources. For each individual, only their accessible food resources were considered—ideally those from the same geographic region and time period. For each food resource category, the mean and standard deviation of the isotopic values were calculated and used to define a normal distribution (Supplementary Data 2), which was then assigned as the prior distribution for the corresponding *I*_ijk_ (see SI 6B for details).

*T*_*k*_ represents the diet-to-tissue offset for the *k*-th dietary isotope proxy. It is modeled as a normally distributed parameter: 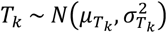 . The prior distributions were informed by empirical findings from previous studies^100,231^, specified as follows: 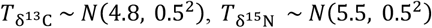 .

After defining the likelihood and specifying prior distributions for all parameters, we employed the MCMC method to estimate the posterior distributions of *θ*_*i*_ for each individual. Three MCMC chains were run in parallel. An adaptive phase of 5,000 iterations was used to allow the sampler to tune its proposal distributions, followed by a burn-in phase of 5,000 iterations. After burn-in, we collected 5,000 posterior samples per chain, resulting in a total of 15,000 posterior draws across all chains for posterior inference. All procedures were conducted using Just Another Gibbs Sampler^233^ (JAGS, v4.3.2) via R.

Existing tools such as FRUITS^224^ scale poorly, lack batch prior specification, and rely on uninformative Dirichlet priors whose limitations we have noted above. Our code enables individualized informative priors, batch processing, and greatly improved computational efficiency.

#### Time-series dietary models (9000 BC to AD 1900)

After reconstructing the diet of each individual, we divided the past 11,000 years into a series of time slices. For the period before 3800 BC, where the sample density is low, we used broader time slices of 1000 years (9000–8000 BC, 8000–7000 BC, 7000–6000 BC, 6000–5000 BC, 5000–3800 BC). From 3800 BC onwards, we applied finer slices of 100 years to better capture dietary shifts over time, with the exception of several Bronze Age periods (1400–1200 BC, 1200–1000 BC, and 1000–800 BC), which were grouped into 200-year slices due to particularly limited sample sizes.

Each individual was assigned to a specific time slice based on their dating. For each time slice, we used the posterior mean dietary proportions of the assigned individuals as input data, and applied nonparametric bootstrap resampling (1,000 iterations) to estimate the bootstrap mean of the mean dietary proportions and their 95% confidence intervals (Supplementary Data 4; SI 2J).

#### Subset time-series dietary model (4000 BC to AD 1300) and time-series dairying model (4000 BC to AD 700) for causal discovery

The original dietary model used fixed 100-year intervals since the Neolithic, which was incompatible with the temporal resolution of the ancient DNA data. For causal discovery, we reconstructed a subset dietary time series using only the data from 4000 BC to AD 1300 to match the temporal resolution of the ancient DNA data. This subset time series was generated using a sliding-window scheme (1,000-year windows with 100-year steps; Supplementary Data 9). In addition to the five dietary components, dairy consumption data^18^ (4000 BC to AD 700) were incorporated using the same sliding-window approach, with the proportion of pottery sherds exhibiting Δ^13^C values below −3.1‰ serving as an index of dairy use (Supplementary Data 9). For each sliding-window slice, we assigned individuals to their corresponding slices based on their dating (a single individual may be assigned to multiple slices). Within each slice, we then performed non-parametric bootstrap resampling of individuals (1,000 iterations) to estimate the bootstrap mean of the mean dietary proportions and the proportion of dairy use, together with their 95% confidence intervals (see also SI 2J).

### Allele frequency trajectories construction and time-varying selection coefficients estimation

#### Construction of Allele Frequency Trajectories

To reconstruct the allele frequency trajectories, we applied a sliding time window approach to 14 SNPs. Each window spans 1,000 years and slides forward in 100-year steps, starting from the window spanning 4000 BC to 3000 BC (centered at 3500 BC) and ending at the window spanning AD 300 to AD 1300 (centered at AD 800). Each individual was subsequently assigned to a specific time slice according to their chronological information.

For each time slice, we performed bootstrap resampling^19^ with 1000 iterations. In each iteration, individuals were sampled with replacement, and the derived allele frequency was calculated. The mean of 1000 bootstrap estimates (derived allele frequencies) was used as the point estimate for that time slice, and the standard deviation was used to quantify its uncertainty (Supplementary Data 7; SI 3A).

Furthermore, we obtained derived allele frequency data for all targeted SNPs from the Ancient Genome Selection (AGES) database^5^, covering the broader European region, and constructed their trajectories to facilitate comparative analysis with the British dataset.

#### The estimation of time-varying selection coefficients

We assumed an additive model (1, 1+s, 1+2s,) for our SNPs. Under standard theoretical models in evolutionary genetics, the change in allele frequency over time follows the deterministic equation^234^:

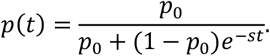

Then^235^,

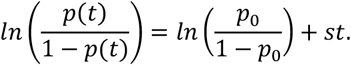

We employed this equation as the basis for constructing a Generalized Additive Model (GAM).

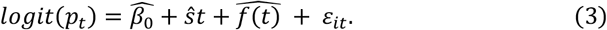

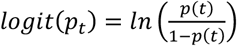 is the link function to transfer observed alleles (ancestral/derived) into its corresponding allele frequency p(t)/1-p(t) at time t. p(t) represents the derived allele frequency at time t. 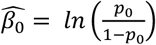 , where 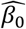 is the intercept to be inferred. Here, *p*_0_ represents derived allele frequency at time *t*_0_. ŝ is the estimated regression coefficient, interpreted as the selection coefficient. 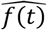 denotes the smooth function to be inferred. It serves as a black-box function of time that captures the aggregate effects of all other time-varying factors influencing allele frequency, excluding natural selection—such as genetic drift, gene flow, non-random mating, and recurrent mutation. *ε*_*it*_ is the error term, automatically incorporated by the GAM model. It captures residual variation in allele frequency not explained by either the linear selection component or the smooth non-linear term.

The smooth term 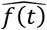 was specified with thin plate regression splines and a first-order penalty, allowing minimal curvature while avoiding overfitting^236^. We used the Restricted Maximum Likelihood approach (REML) for stable parameter estimation, and set select = TRUE to allow shrinkage of the smooth term toward zero if unsupported by the data. This penalized smoothing strategy helps to reduce potential concurvity between the linear and smooth components^237^, ensuring that the linear term reflects the effect of natural selection, while the smooth term accounts for other effects not explained by selection.

Our explanatory variable is time t, measured in units of twice the generation interval (i.e., 60 years). The response variable is the observed allele at time t, modeled as a Bernoulli-distributed random variable *y*_*i*_ ∼ *Bernoulli* (*p*_*t*_), where *y*_*i*_ = 0 represents the ancestral allele, *y*_*i*_ = 1 represents the derived allele. Here, i indexes each sample (i.e., allele) at time t. The parameter *p*_t_ denotes the derived allele frequency at the corresponding time. Each pair (*y*_*i*_, *t*) is incorporated into the model to estimate the coefficients of the GAM model. We first applied the model across the full temporal range to estimate the average selection coefficients (Supplementary Data 8). Then, the time-varying selection coefficients were estimated by fitting the model separately to each sliding time window (Supplementary Data 8). When estimating time-varying selection coefficients, the model specification remained unchanged; the only difference was that we used MLE instead of REML to estimate the model parameters. This choice was motivated by the typically smaller sample size within each time bin, which can lead REML to produce overly conservative estimates of smoothing parameters^236^. ML is more appropriate, as it tends to perform better with limited data, allowing the model to capture meaningful patterns without excessive penalization.

After estimating the selection coefficients, we used the fitted model to simulate allele frequency trajectories and compared the simulated trajectories with the original frequency trajectories derived from aDNA data. This comparison served to validate the reliability and accuracy of our model (SI 3B).

The number of time-varying selection coefficients is relatively small for rs3891176, rs4988235, and rs7775397 (especially rs3891176 and rs7775397). The reason is that the number of time intervals for which selection coefficients could be reliably estimated was limited for these SNPs (SI 3C).

### Causal discovery with convergent cross mapping

Convergent cross mapping (CCM)^20^ is a method specifically designed for causal discovery in time-series data. It does not assume predefined functional form between variables and identifies causal relationships directly from the data, making it well suited for complex nonlinear systems such as ours. After reviewing a wide range of methods for causal discovery^20,238–244^ and causal inference^68–79,245–254^ , we found CCM to be the most appropriate for our research context and data structure (see SI 4B).

The output of CCM results typically plots library length (L) on the x-axis against the correlation coefficient (ρ) on the y-axis. A steadily increasing ρ that converges to a value significantly above zero as L grows provides evidence for a causal effect. The magnitude of ρ reflects the strength of the causal relationship. Because CCM has no universally accepted threshold for determining convergence above zero, we adopted a conservative criterion: the correlation coefficient ρ must exceed 0.25 at the maximum library length (ρ_Lmax_ >0.25), and the mean absolute change in ρ over the last five library steps must be no greater than 0.02 (|Δρ_last 5L_| < 0.02).

In our case, time-series dietary data (including dairy) serve as the treatment variable X, while the corresponding time-series selection coefficients act as the outcome variable Y. We initially conducted CCM analyses using the point estimates of time-series selection coefficients for each SNP and their corresponding time-series dietary proportions (Dairy proportion: rs4988235; Marine fish proportion: rs7944926, rs174546, rs174594, rs174570; C_3_ plants proportion: rs653178, rs272872, rs7517, rs4073089, rs6265, rs12401678, rs2010410; terrestrial animal meat proportion: rs174546, rs174594, rs174570). However, as these values are inferred rather than directly observed, all variables carry inherent uncertainty at each time slice. To account for this, we assumed that diet proportions and selection coefficients of each time slice all follow normal distributions, with point estimates (means) and standard deviations as parameters. We then generated 100 simulated time series for each SNP’s selection coefficients and its corresponding dietary proportions. These simulations were used to perform 100 separate CCM analyses per SNP, allowing us to evaluate the robustness of causal discovery under uncertainty (Supplementary Data 10; SI 4A).

All CCM analyses were conducted using the same parameter settings: the embedding dimension E=3, the time delay τ=1, and the number of nearest neighbors k=4 (see SI 4C for the rationale behind these parameter choices). In our case, the total number of time points N in the time series is 29 for rs4988235 and the dairy data, and 44 for all other SNPs and dietary variables.

Given the large-scale nature of our data, we did not rely on existing R packages for CCM, but instead implemented the algorithm from scratch based on its mathematical foundations to facilitate high-throughput processing. The mathematical details of the CCM algorithm and the philosophy underlying how CCM identifies causality are provided in SI 4C.

This study benefited from the use of OpenAI’s ChatGPT, which assisted in editing the manuscript, annotating R scripts, and refining code structure and documentation. All intellectual decisions and substantive content remain the responsibility of the author.

## Data Availability

All supplementary data used in this study are available at GitHub: https://github.com/zexuan-chen/Chen_bioRxiv_2025.

## Code Availability

All code used in this study is available at GitHub: https://github.com/zexuan-chen/Chen_bioRxiv_2025. The code is licensed under the MIT License, which permits unrestricted use, distribution, and modification, provided that the original author is properly credited.

## Acknowledgements

We would like to sincerely thank Dr. Stephan Schiffels, Dr. Joscha Gretzinger, Prof. Martin Richards, Dr. Ceiridwen Edwards, and Dr. George Foody for generously sharing the BAM files used in their respective studies. I am also deeply grateful to Prof. Mike Parker Pearson and Dr. Alison Sheridan for their invaluable guidance in accessing the isotopic data of Beaker-associated individuals. We thank Dr. Iain Mathieson for his valuable comments and suggestions on earlier versions of this manuscript. We gratefully acknowledge the financial support provided by Durham University and the China Scholarship Council (CSC).

## Author contributions

Z.C. conceived the study, designed the analytical workflow, analyzed the data, and wrote the manuscript. A.M. and E.F.D. provided comments on the manuscript.

## Supplementary information

Supplementary Information is available as a single PDF file accompanying this manuscript.

---

**Extended Data Fig. 1.**
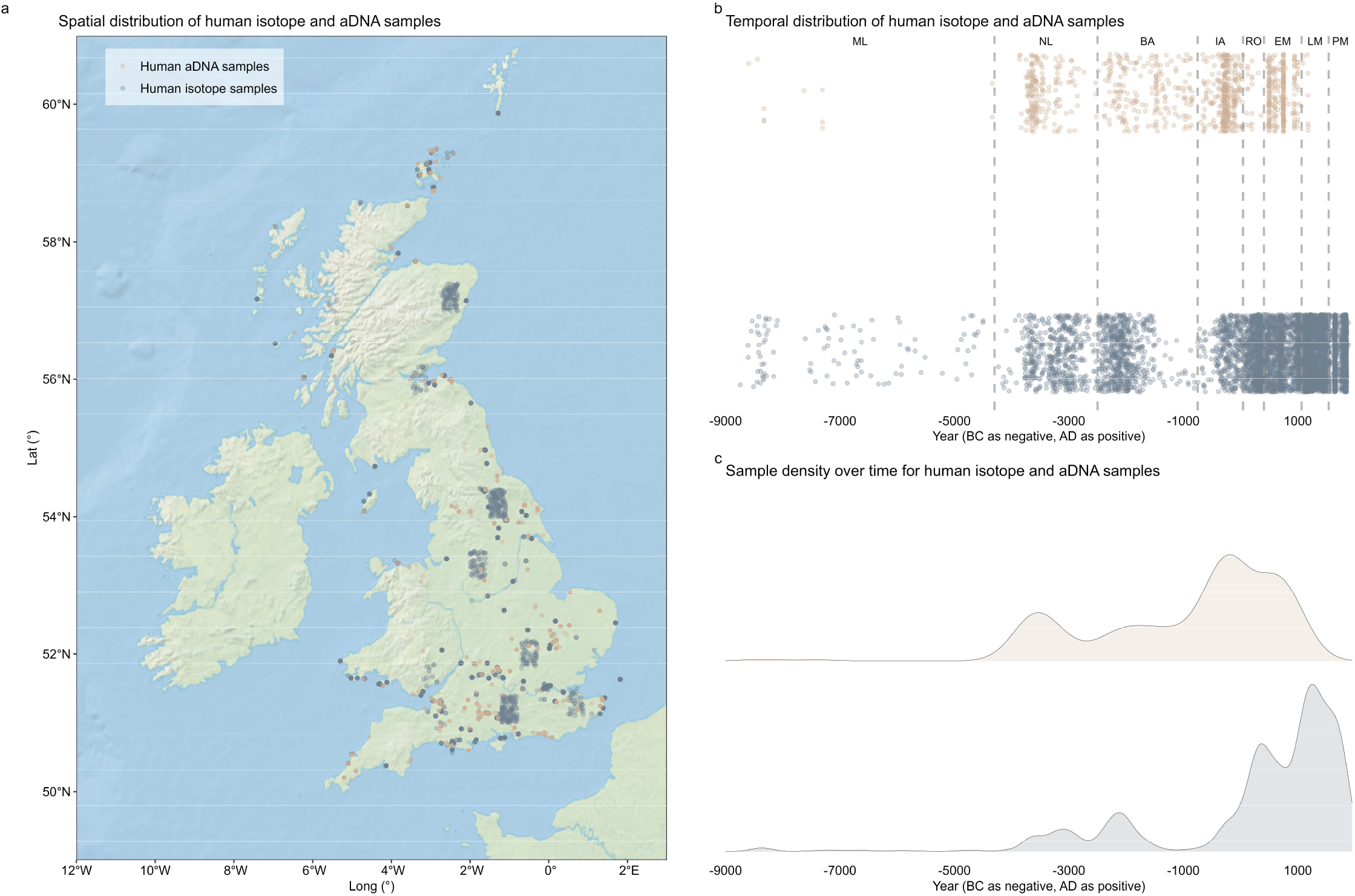
Distributions of human isotope (n = 6,046) and ancient DNA (n = 1,038) samples. **a**, Spatial distribution. **b**, Temporal distribution. **c**, Density over time. Abbreviations: ML – Mesolithic; NL – Neolithic; BA – Bronze Age; IA – Iron Age; RO – Roman period/Roman Iron Age; EM – Early Medieval period; LM – Later Medieval period; PM – Post-Medieval period. Although a few ancient DNA samples fall within the Mesolithic period, their number is too small for reliable analysis; therefore, only samples dated between 4000 BC and AD 1300 were used to construct derived allele frequency trajectories and to estimate selection coefficients. There are 6,064 human isotope samples in total. The Paleolithic samples (n=18) were excluded from the time-series modeling and are not shown in this figure.

**Extended Data Fig. 2.**
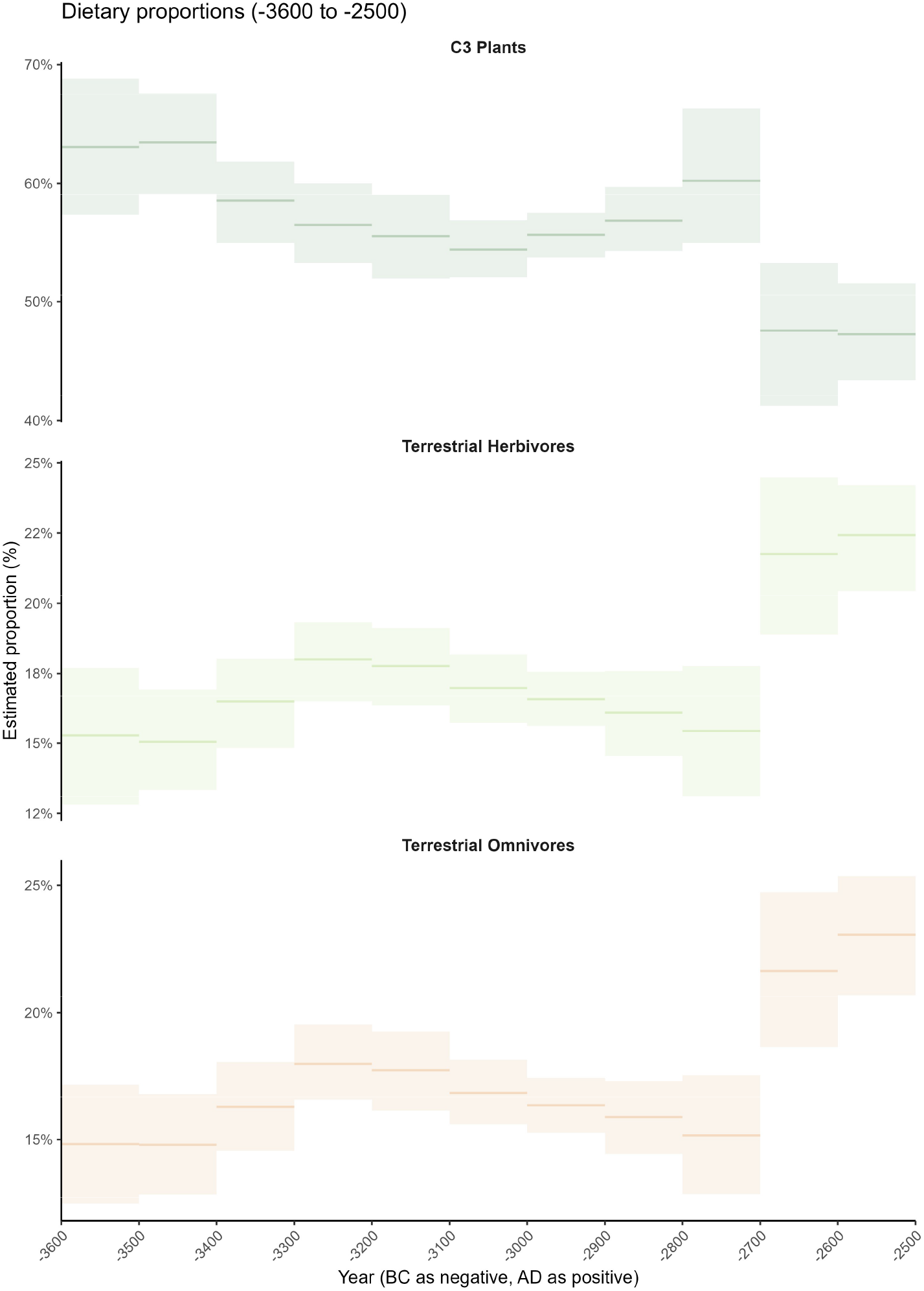
Mean proportions of C_3_ plants, terrestrial herbivores, terrestrial omnivores across time (3600-2500 BC). Lines and bars represent the mean and 95% confidence interval within each slice, calculated from the posterior mean dietary proportions of individuals assigned to that slice. Estimates were obtained using nonparametric bootstrap resampling (1,000 iterations).

**Extended Data Fig. 3.**
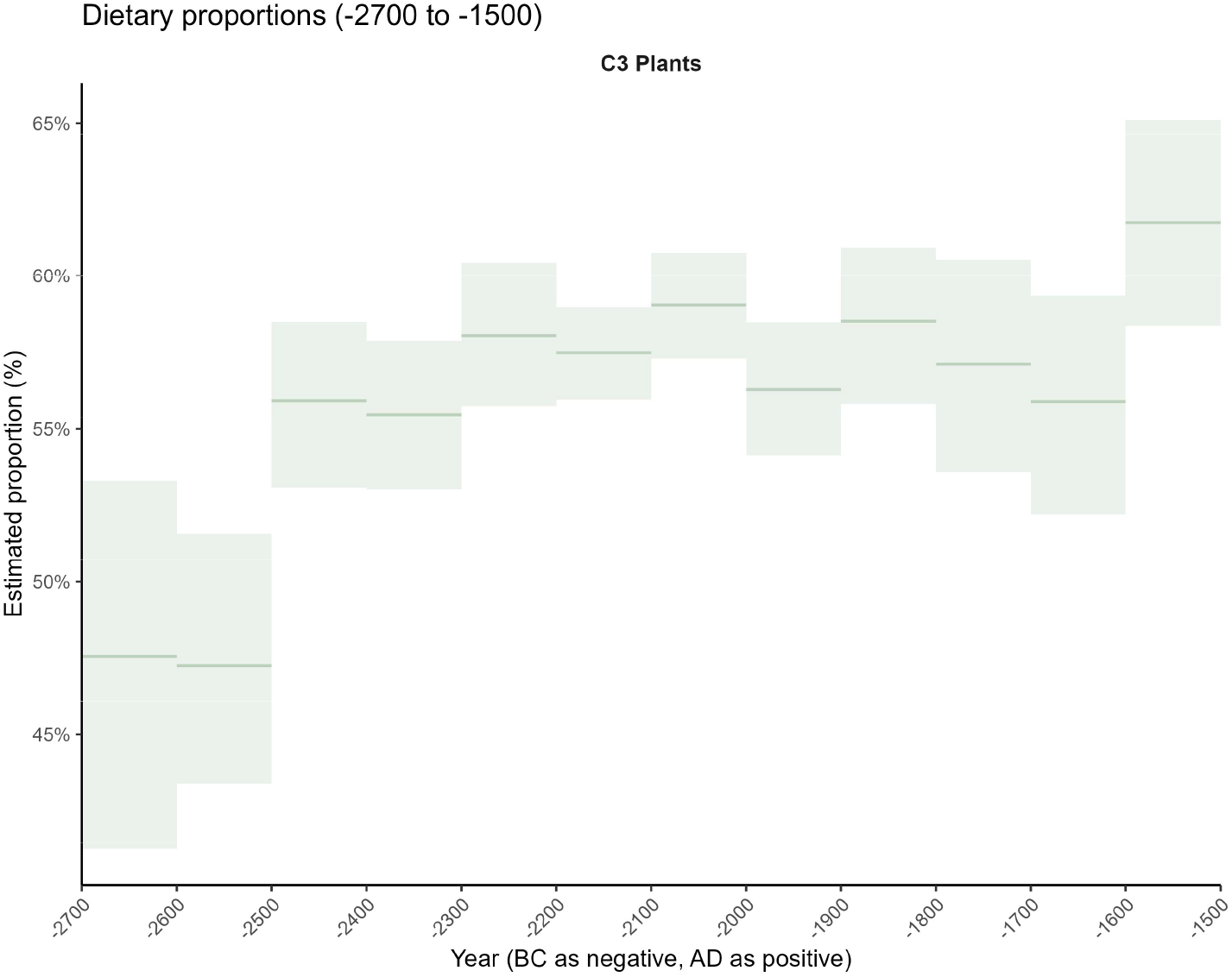
Mean proportions of C_3_ plants across time (2700-1500 BC). Lines and bars represent the mean and 95% confidence interval within each slice, calculated from the posterior mean dietary proportions of individuals assigned to that slice. Estimates were obtained using nonparametric bootstrap resampling (1,000 iterations).

**Extended Data Fig. 4.**
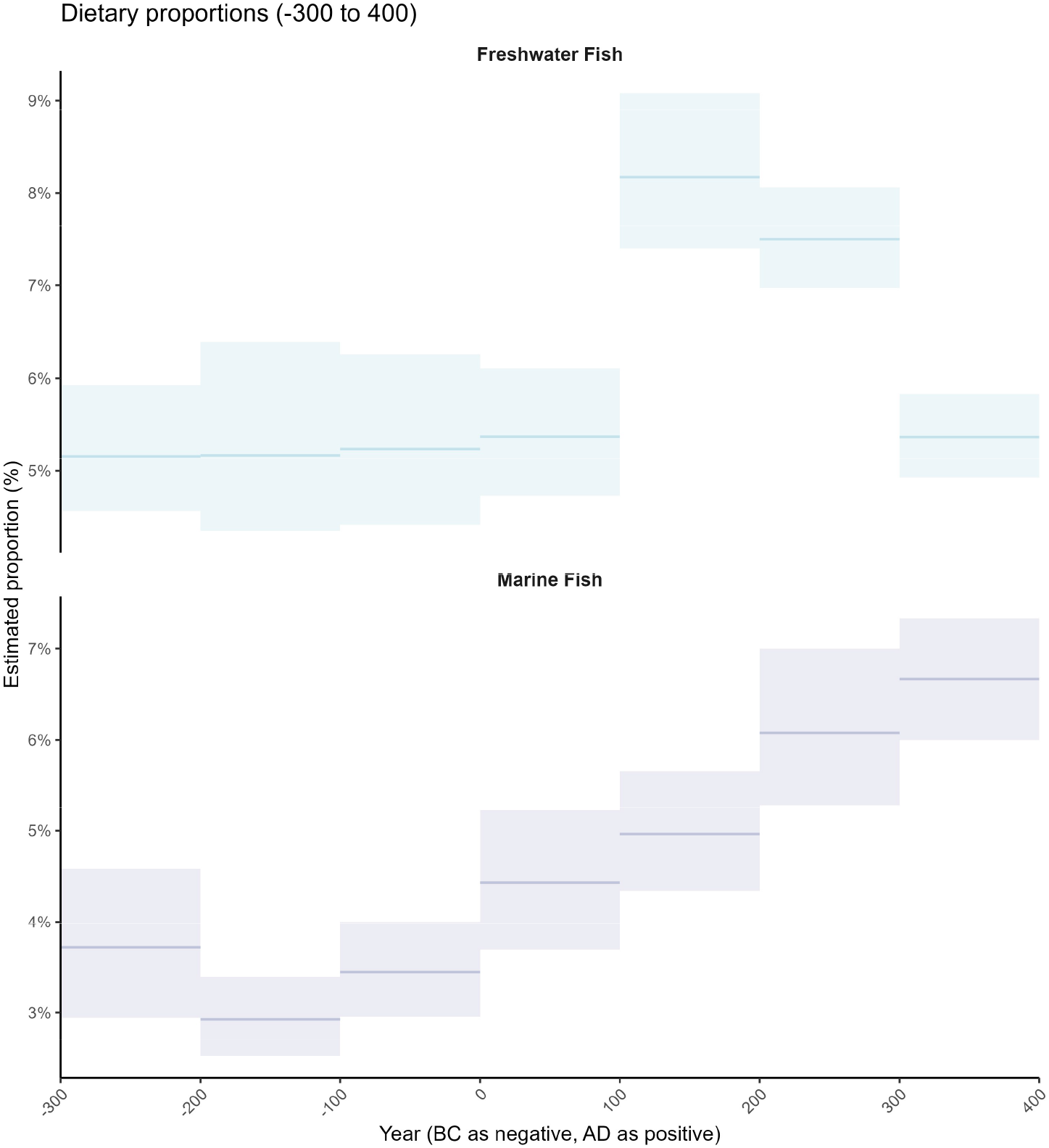
Mean proportions of marine and freshwater fish across time (300 BC-AD 400). Lines and bars represent the mean and 95% confidence interval within each slice, calculated from the posterior mean dietary proportions of individuals assigned to that slice. Estimates were obtained using nonparametric bootstrap resampling (1,000 iterations).

**Extended Data Fig. 5.**
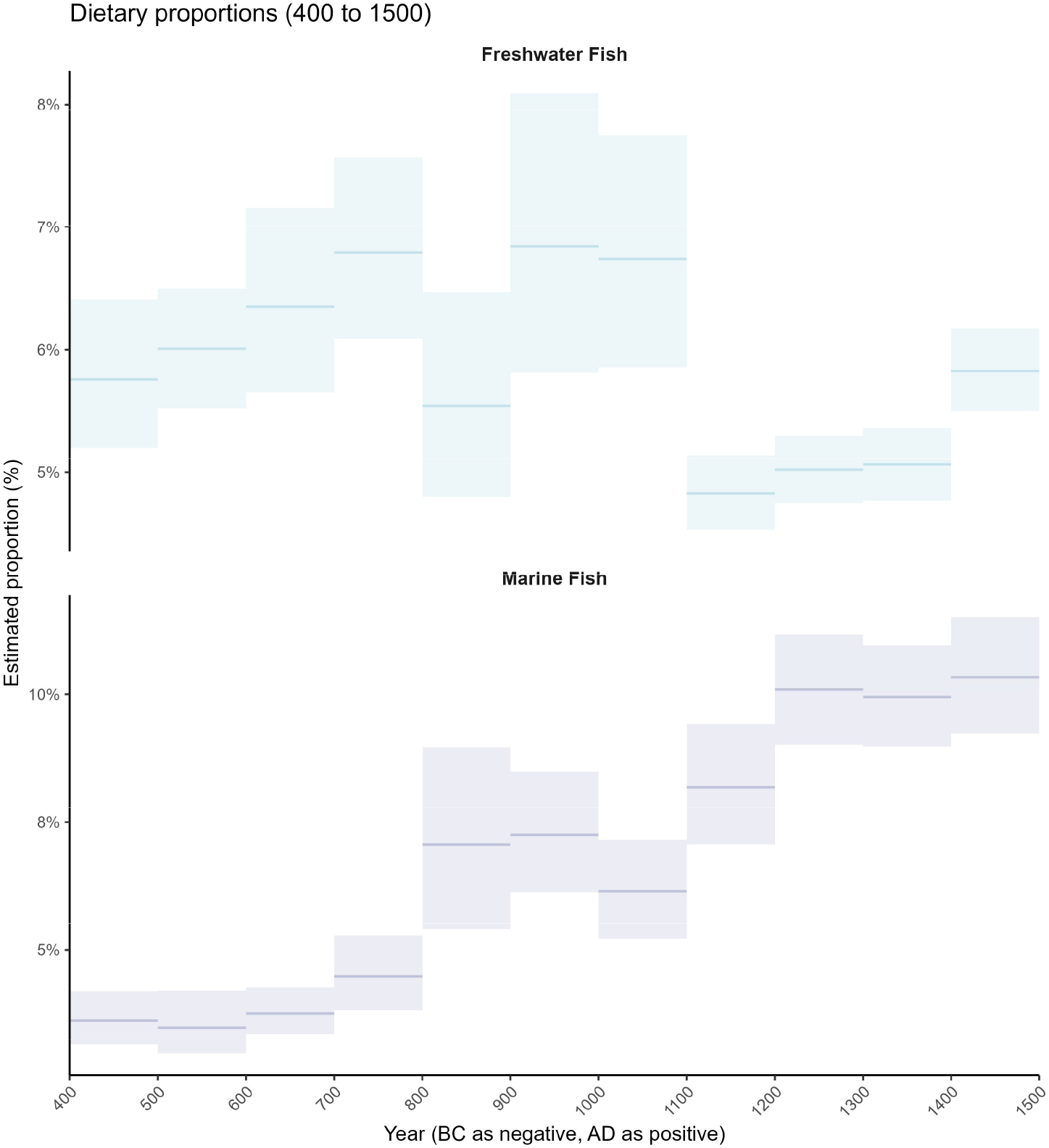
Mean proportions of marine and freshwater fish across time (AD 400-1500). Lines and bars represent the mean and 95% confidence interval within each slice, calculated from the posterior mean dietary proportions of individuals assigned to that slice. Estimates were obtained using nonparametric bootstrap resampling (1,000 iterations).

**Extended Data Fig. 6.**
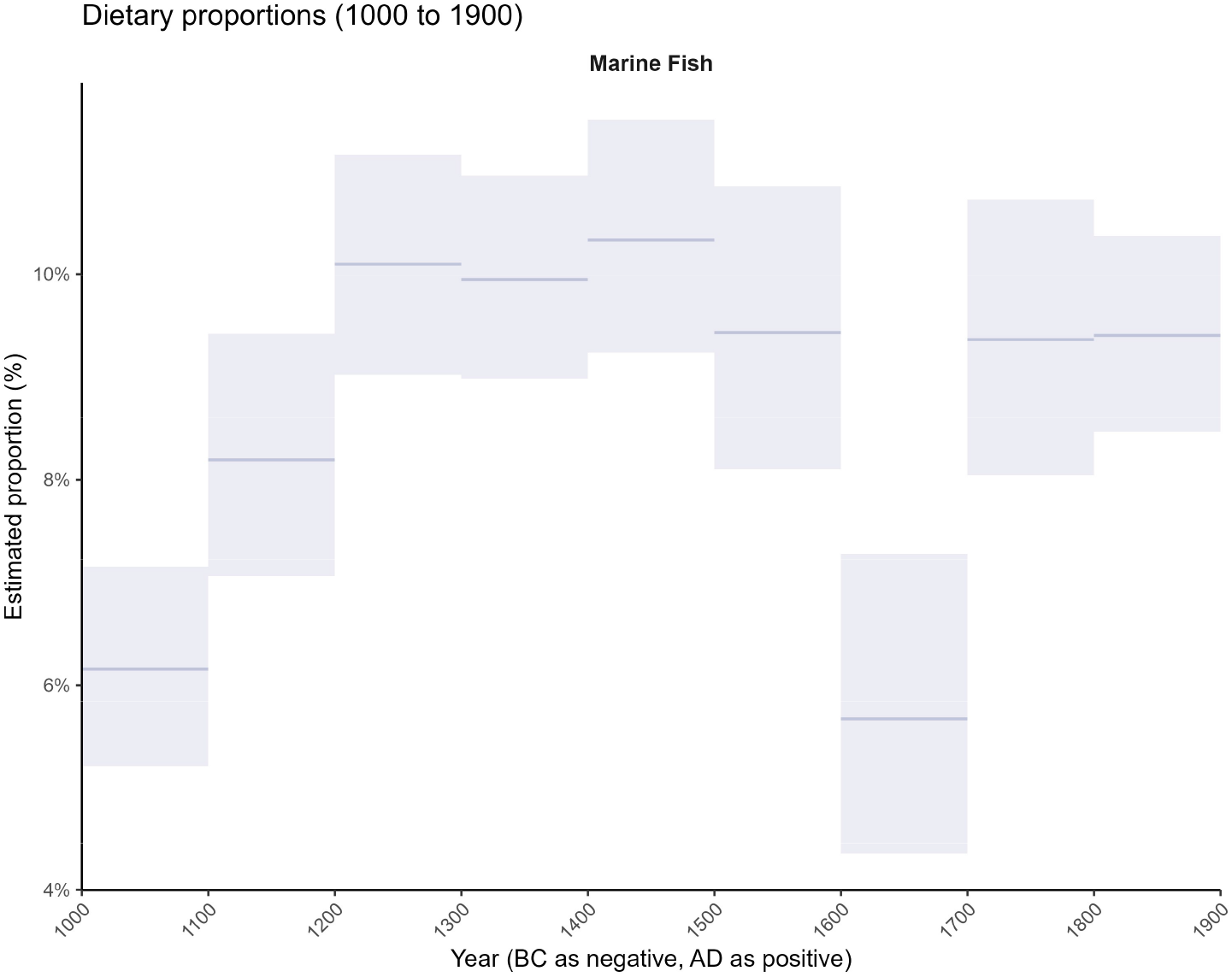
Mean proportions of marine fish across time (AD 1000-1900). Lines and bars represent the mean and 95% confidence interval within each slice, calculated from the posterior mean dietary proportions of individuals assigned to that slice. Estimates were obtained using nonparametric bootstrap resampling (1,000 iterations).

**Extended Data Fig. 7.**
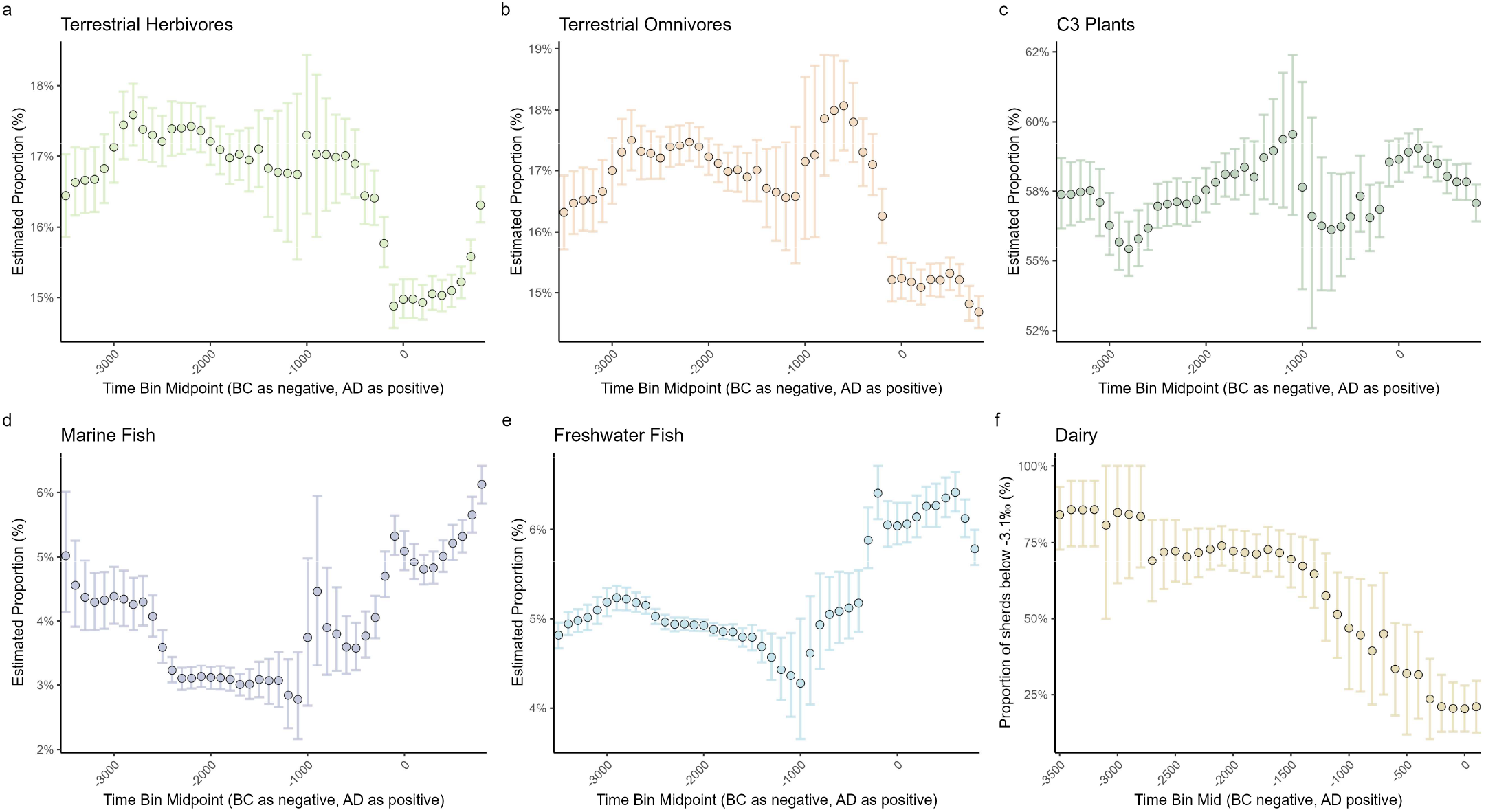
Time-series diet for causal discovery. The dietary model was constructed using sliding time windows to align with the corresponding time slices of selection coefficients. The error bars indicate the 95% confidence intervals.

**Extended Data Fig. 8.**
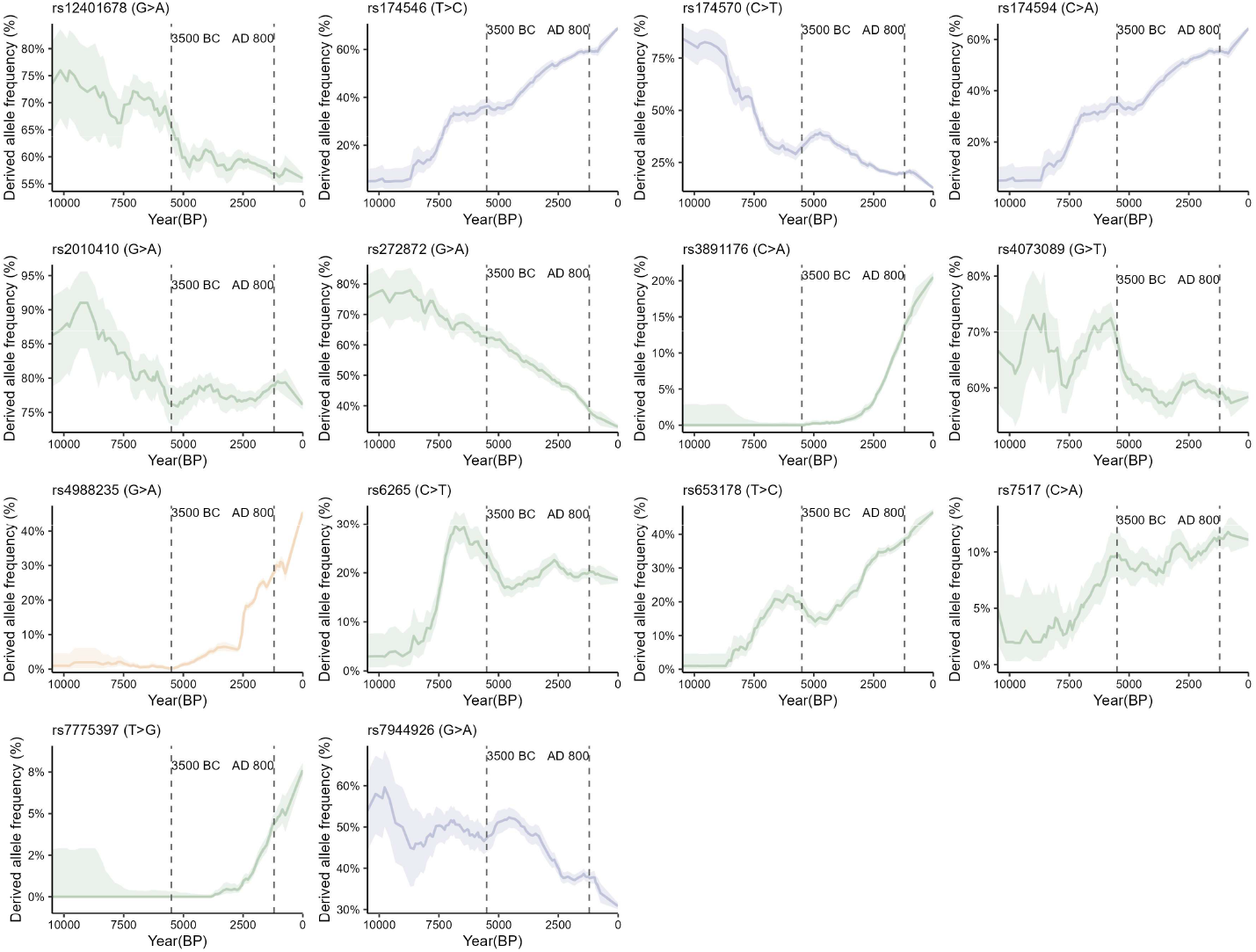
Allele frequency trajectories for 14 SNPs (based on European data). Data were obtained from the Ancient Genome Selection Database (AGES). The shaded area represents the 95% confidence interval.

**Extended Data Fig. 9.**
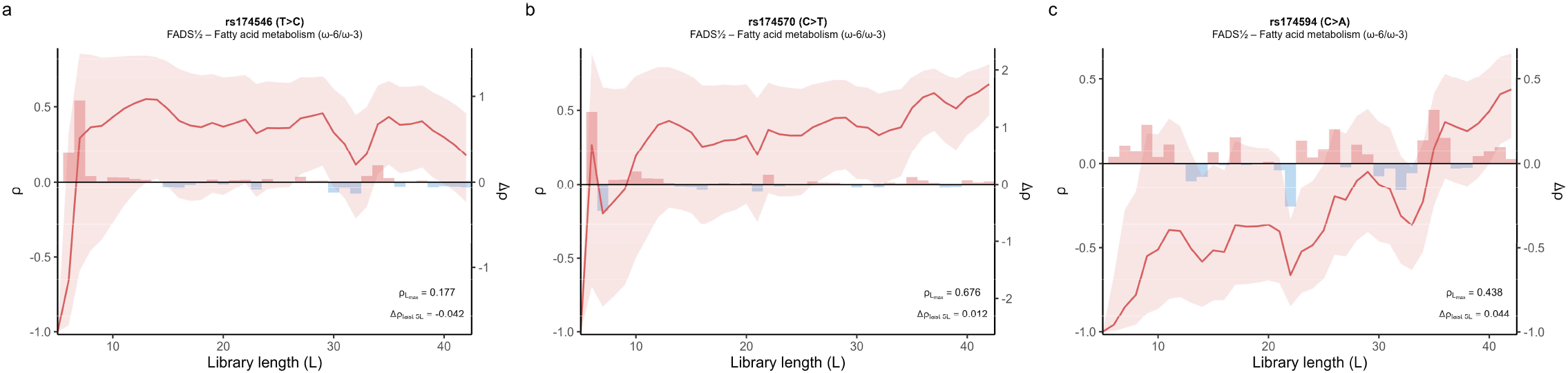
CCM results for three fatty acid metabolism SNPs (dietary driver: terrestrial animals = herbivores + omnivores). **a**, rs174546. **b**, rs174570. **c**, rs174594. As the library length increases, convergence of the coefficient (ρ) above zero indicates causality, with larger values denoting stronger effects. Stepwise changes are given by Δρ(L) = ρ(L) − ρ(L – 1). ρLmax is the ρ at the final L, and Δρlast 5L is the mean Δρ across the last five L steps. Line: ρ; Shade: 95% CI; Bars: Δρ. CCM results are based on the point estimates of dietary variables and selection coefficients. The key parameters: the embedding dimension E=3, the time delay τ=1, and the number of nearest neighbors k=4 for all CCM tests.

**Extended Data Table 1.** Detailed information on the 14 targeted SNPs, their corresponding phenotypes and potential dietary drivers.

| RSID | Allele | Food | Gene | Phenotype |
| --- | --- | --- | --- | --- |
| rs12401678 | G>A | C<sub>3</sub> plant | PTPRV | Circulating glucose levels |
| rs174546 | T>C | Marine fish<br>Terrestrial animals | FADS1/2 | Fatty acid metabolism (omega-6 and omega-3) |
| rs174570 | C>T | Marine fish<br>Terrestrial animals | FADS1/2 | Fatty acid metabolism (omega-6 and omega-3) |
| rs174594 | C>A | Marine fish<br>Terrestrial animals | FADS1/2 | Fatty acid metabolism (omega-6 and omega-3) |
| rs2010410 | G>A | C<sub>3</sub> plant | FTO/IRX3 | Obesity |
| rs272872 | G>A | C<sub>3</sub> plant | SLC22A4 | Ergothioneine level |
| rs3891176 | C>A | C<sub>3</sub> plant | HLA-DQB1 | Celiac disease susceptibility |
| rs4073089 | G>T | C<sub>3</sub> plant | MAF | Insulin secretion<br>Type 2 diabetes |
| rs4988235 | G>A | Dairy | LCT | Lactase persistence |
| rs6265 | C>T | C<sub>3</sub> plant | BDNF | Obesity<br>Type 2 diabetes |
| rs653178 | T>C | C<sub>3</sub> plant | ATXN2/SH2B3 | Celiac disease susceptibility |
| rs7517 | C>A | C<sub>3</sub> plant | ENSA | Insulin secretion<br>Type 2 diabetes |
| rs7775397 | T>G | C<sub>3</sub> plant | TSBP1 | Celiac disease susceptibility |
| rs7944926 | G>A | Marine fish | DHCR7/NADSYN1 | Vitamin D level |

**Extended Data Table 2.**
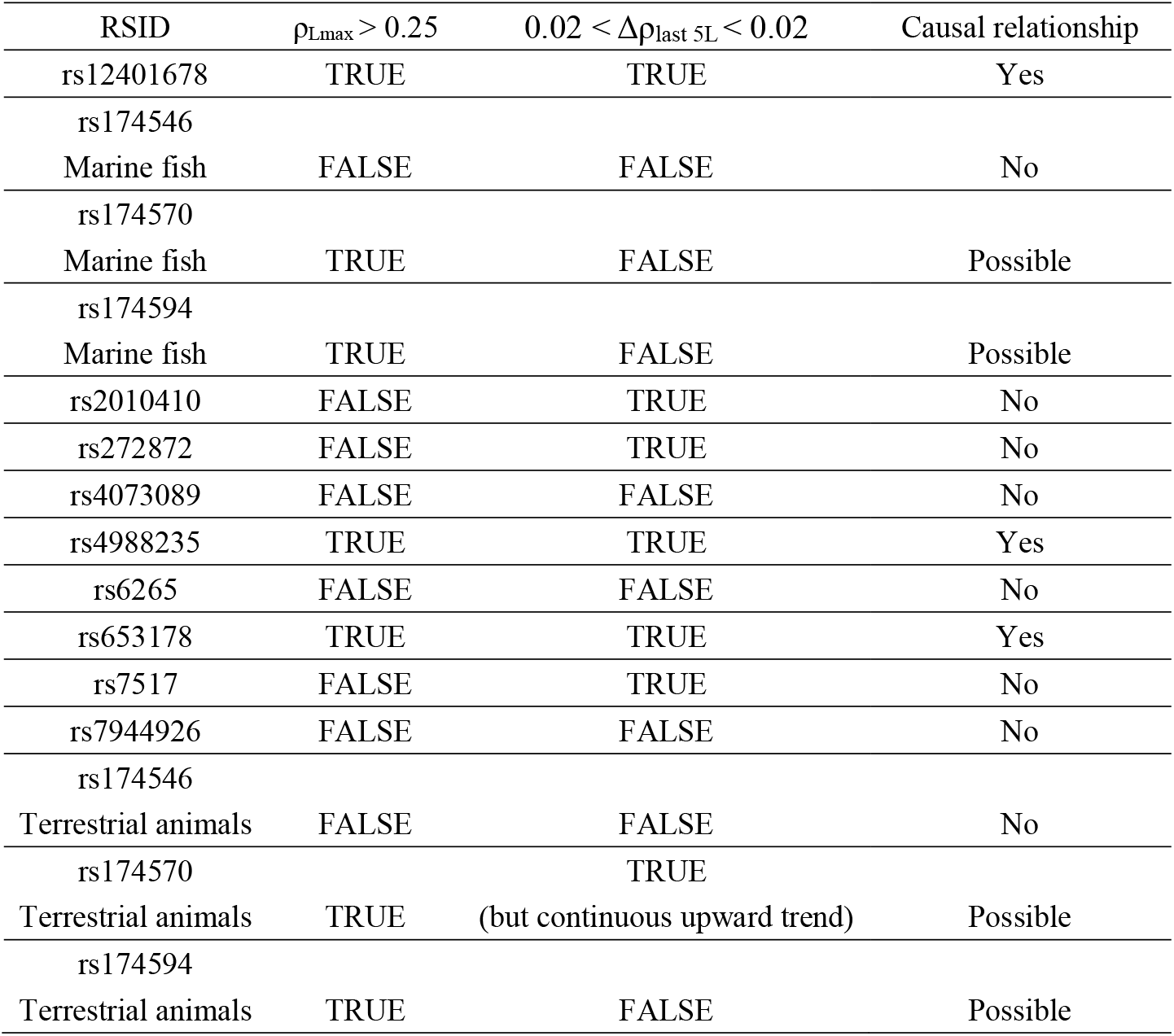
Summary of causal discovery results across 12 SNPs.

## References

1. Mathieson, I. et al. Genome-wide patterns of selection in 230 ancient Eurasians. Nature 528, 499–503 (2015).

2. Le, M. K. et al. 1,000 ancient genomes uncover 10,000 years of natural selection in Europe. bioRxiv https://doi.org/10.1101/2022.08.24.505188 (2022) doi:10.1101/2022.08.24.505188.

3. Kerner, G. et al. Genetic adaptation to pathogens and increased risk of inflammatory disorders in post-Neolithic Europe. Cell Genomics 3, 100248 (2023).

4. Irving-Pease, E. K. et al. The selection landscape and genetic legacy of ancient Eurasians. Nature 625, 312–320 (2024).

5. Akbari, A. et al. Pervasive findings of directional selection realize the promise of ancient DNA to elucidate human adaptation. bioRxiv https://doi.org/10.1101/2024.09.14.613021 (2024) doi:10.1101/2024.09.14.613021.

6. Fumagalli, M. et al. Greenlandic Inuit show genetic signatures of diet and climate adaptation. Science (1979). 349, 1343–1347 (2015).

7. Field, Y. et al. Detection of human adaptation during the past 2000 years. Science (1979). 354, 760–764 (2016).

8. Ye, K., Gao, F., Wang, D., Bar-Yosef, O. & Keinan, A. Dietary adaptation of FADS genes in Europe varied across time and geography. Nat. Ecol. Evol. 1, 0167 (2017).

9. Tishkoff, S. A. et al. Convergent adaptation of human lactase persistence in Africa and Europe. Nat. Genet. 39, 31–40 (2007).

10. Bersaglieri, T. et al. Genetic Signatures of Strong Recent Positive Selection at the Lactase Gene. Am. J. Hum. Genet. 74, 1111–1120 (2004).

11. Pontremoli, C. et al. Natural selection at the brush-border: Adaptations to carbohydrate diets in humans and other mammals. Genome Biol. Evol. 7, 2569–2584 (2015).

12. Hancock, A. M. et al. Human adaptations to diet, subsistence, and ecoregion are due to subtle shifts in allele frequency. Proc. Natl. Acad. Sci. U. S. A. 107, 8924–8930 (2010).

13. Scheer, K. et al. Rapid Adaptive Increase of Amylase Gene Copy Number in Indigenous Andeans. bioRxiv https://doi.org/10.1101/2025.03.25.644684 (2025) doi:10.1101/2025.03.25.644684.

14. Rees, J., Castellano, S. & Andrés, A. M. Global impact of micronutrients in modern human evolution. Am. J. Hum. Genet. 112, 2538–2561 (2025).

15. Bolognini, D. et al. Recurrent evolution and selection shape structural diversity at the amylase locus. Nature 634, 617–625 (2024).

16. Yilmaz, F. et al. Reconstruction of the human amylase locus reveals ancient duplications seeding modern-day variation. Science (1979). 386, eadn0609 (2024).

17. Perry, G. H. et al. Diet and the evolution of human amylase gene copy number variation. Nat. Genet. 39, 1256–1260 (2007).

18. Evershed, R. P. et al. Dairying, diseases and the evolution of lactase persistence in Europe. Nature 608, 336–345 (2022).

19. Efron, B. Bootstrap Methods: Another Look at the Jackknife. The Annals of Statistics 7, 1–26 (1979).

20. Sugihara, G. et al. Detecting causality in complex ecosystems. Science (1979). 338, 496–500 (2012).

21. Eaton, S. & Konner, M. Paleolithic Nutrition: A Consideration of Its Nature and Current Implications. New England Journal of Medicine 312, 283–289 (1985).

22. Cordain, L. et al. Origins and evolution of the Western diet: Health implications for the 21st century. American Journal of Clinical Nutrition 81, 341–354 (2005).

23. Konner, M. & Boyd Eaton, S. Paleolithic nutrition: Twenty-five years later. Nutrition in Clinical Practice 25, 594–602 (2010).

24. Eaton, S. B., Eaton, S. B. & Konner, M. J. Paleolithic nutrition revisited: A twelve-year retrospective on its nature and implications. Eur. J. Clin. Nutr. 51, 207–216 (1997).

25. Andrews, P. & Johnson, R. J. Evolutionary basis for the human diet: consequences for human health. J. Intern. Med. 287, 226–237 (2020).

26. Leonard, W. R. Food for thought: Dietary change was a driving force in human evolution. Sci. Am. 287, 106–115 (2002).

27. James, W. P. T. et al. Nutrition and its role in human evolution. J. Intern. Med. 285, 533–549 (2019).

28. Bragazzi, N. L., Del Rio, D., Mayer, E. A. & Mena, P. We Are What, When, And How We Eat: The Evolutionary Impact of Dietary Shifts on Physical and Cognitive Development, Health, and Disease. Advances in Nutrition 15, 100280 (2024).

29. Alt, K. W., Al-Ahmad, A. & Woelber, J. P. Nutrition and Health in Human Evolution–Past to Present. Nutrients 14, 3594 (2022).

30. Babbitt, C. C., Warner, L. R., Fedrigo, O., Wall, C. E. & Wray, G. A. Genomic signatures of diet-related shifts during human origins. Proceedings of the Royal Society B: Biological Sciences 278, 961–969 (2011).

31. Luca, F., Perry, G. H. & Di Rienzo, A. Evolutionary adaptations to dietary changes. Annu. Rev. Nutr. 30, 291–314 (2010).

32. Altman, N. & Krzywinski, M. Association, correlation and causation. Nat. Methods 12, 899–900 (2015).

33. Mithen, S. & Milner, N. Hunter-gatherers of the Mesolithic. in The Archaeology of Britain 36–59 (London, 1999).

34. van der Plicht, J., Amkreutz, L. W. S. W., Niekus, M. J. L. T., Peeters, J. H. M. & Smit, B. I. Surf’n Turf in Doggerland: Dating, stable isotopes and diet of Mesolithic human remains from the southern North Sea. J. Archaeol. Sci. Rep. 10, 110–118 (2016).

35. Schulting, R. J. & Richards, M. P. Finding the coastal Mesolithic in southwest Britain: AMS dates and stable isotope results on human remains from Caldey Island, South Wales. Antiquity 76, 1011–1025 (2002).

36. Meiklejohn, C., Chamberlain, A. T. & Schulting, R. J. Radiocarbon dating of Mesolithic human remains in Great Britain. Mesolithic Miscellany 21, 20–58 (2011).

37. Schulting, R. et al. Mesolithic and neolithic human remains from Foxhole Cave, Gower, South Wales. The Antiquaries Journal 93, 1–23 (2013).

38. Richards, M. P., Schulting, R. & Hedges, R. E. M. Sharp shift in diet at onset of Neolithic. Nature 423, 136–137 (2003).

39. Woodbridge, J. et al. The impact of the Neolithic agricultural transition in Britain: A comparison of pollen-based land-cover and archaeological 14C date-inferred population change. J. Archaeol. Sci. 51, 216–224 (2014).

40. Stevens, C. J. & Fuller, D. Q. Did neolithic farming fail? The case for a Bronze Age agricultural revolution in the British Isles. Antiquity 86, 707–722 (2012).

41. Treasure, E. R., Gröcke, D. R., Caseldine, A. E. & Church, M. J. Neolithic farming and wild plant exploitation in western Britain: Archaeobotanical and crop stable isotope evidence from Wales (c. 4000–2200 cal BC). Proceedings of the Prehistoric Society 85, 193–222 (2019).

42. Bishop, R. R. Did Late Neolithic farming fail or flourish? A Scottish perspective on the evidence for Late Neolithic arable cultivation in the British Isles. World Archaeol. 47, 834–855 (2015).

43. Jay, M. & Richards, M. P. Carbon and nitrogen isotopic analysis. in The Beaker People: Isotopes, Mobility and Diet in Prehistoric Britain (eds. Mike Parker Pearson et al.) 303–339 (Oxbow Books, Oxford, 2019).

44. Pearson, M. P. et al. Evidence for mummification in Bronze Age Britain. Antiquity 79, 529–546 (2005).

45. Parker Pearson, M. et al. Beaker people in Britain: migration, mobility and diet. Antiquity 90, 620–637 (2016).

46. Lawrence, D. M. Orkney’s first farmers. Reconstructing biographies from osteological analysis to gain insights into life and society in a Neolithic community on the edge of Atlantic Europe. (University of Bradford, 2012).

47. Armit, I., Schulting, R., Knusel, C. J. & Shepherd, I. A. G. Death, decapitation and display? The bronze and iron age human remains from the sculptor’s cave, covesea, North-east Scotland. Proceedings of the Prehistoric Society 77, 251–278 (2011).

48. Stevens, R. E., Lightfoot, E., Allen, T. & Hedges, R. E. M. Palaeodiet at Eton College Rowing Course, Buckinghamshire: Isotopic changes in human diet in the Neolithic, Bronze Age, Iron Age and Roman periods throughout the British Isles. Archaeol. Anthropol. Sci. 4, 167–184 (2012).

49. Lightfoot, E. et al. An investigation into diet at the site of yarnton, oxfordshire, using stable carbon and nitrogen isotopes. Oxford Journal of Archaeology 28, 301–322 (2009).

50. Richards, M. P., Hedges, R. E. M., Molleson, T. I. & Vogel, J. C. Stable Isotope Analysis Reveals Variations in Human Diet at the Poundbury Camp Cemetery Site. J. Archaeol. Sci. 25, 1247–1252 (1998).

51. Redfern, R., Gowland, R., Millard, A., Powell, L. & Gröcke, D. ‘From the mouths of babes’: A subadult dietary stable isotope perspective on Roman London (Londinium). J. Archaeol. Sci. Rep. 19, 1030–1040 (2018).

52. Müldner, G. & Richards, M. P. Stable isotope evidence for 1500 years of human diet at the city of York, UK. Am. J. Phys. Anthropol. 133, 682–697 (2007).

53. Barrett, J. H. & Richards, M. P. Identity, gender, religion and economy: New isotope and radiocarbon evidence for marine resource intensification in early historic Orkney, Scotland, UK. Eur. J. Archaeol. 7, 249–271 (2004).

54. Richards, M. P., Fuller, B. T. & Molleson, T. I. Stable isotope palaeodietary study of humans and fauna from the multi-period (Iron Age, Viking and Late Medieval) site of Newark Bay, Orkney. J. Archaeol. Sci. 33, 122–131 (2006).

55. Jarman, C. L., Biddle, M., Higham, T. & Bronk Ramsey, C. The Viking Great Army in England: new dates from the Repton charnel. Antiquity 92, 183–199 (2018).

56. Pollard, A. M. et al. ‘Sprouting like cockle amongst the wheat’: The St Brice’s Day Massacre and the isotopic analysis of human bones from St John’s College, Oxford. Oxford Journal of Archaeology 31, 83–102 (2012).

57. Curtis-Summers, S., Pearson, J. A. & Lamb, A. L. From Picts to Parish: Stable isotope evidence of dietary change at medieval Portmahomack, Scotland. J. Archaeol. Sci. Rep. 31, 102303 (2020).

58. Curtis-Summers, S., Montgomery, J. & Carver, M. Stable isotope evidence for dietary contrast between Pictish and medieval populations at Portmahomack, Scotland. Mediev. Archaeol. 58, 21–43 (2014).

59. Müldner, G. & Richards, M. P. Fast or feast: Reconstructing diet in later medieval England by stable isotope analysis. J. Archaeol. Sci. 32, 39–48 (2005).

60. Britton, K., Fuller, B. T., Tütken, T., Mays, S. & Richards, M. P. Oxygen isotope analysis of human bone phosphate evidences weaning age in archaeological populations. Am. J. Phys. Anthropol. 157, 226–241 (2015).

61. Bleasdale, M. et al. Multidisciplinary investigations of the diets of two post-medieval populations from London using stable isotopes and microdebris analysis. Archaeol. Anthropol. Sci. 11, 6161–6181 (2019).

62. Beaumont, J. et al. Victims and survivors: Stable isotopes used to identify migrants from the Great Irish Famine to 19th century London. Am. J. Phys. Anthropol. 150, 87–98 (2013).

63. Nitsch, E. K., Humphrey, L. T. & Hedges, R. E. M. The effect of parity status on δ15N: looking for the ‘ pregnancy effect’ in 18th and 19th century London. J. Archaeol. Sci. 37, 3191–3199 (2010).

64. Nitsch, E. K., Humphrey, L. T. & Hedges, R. E. M. Using stable isotope analysis to examine the effect of economic change on breastfeeding practices in Spitalfields, London, UK. Am. J. Phys. Anthropol. 146, 619–628 (2011).

65. Takens, F. Detecting strange attractors in turbulence. in Dynamical Systems and Turbulence, Warwick 1980 (eds. Rand, D. & Young, L.-S.) 366–381 (Springer-Verlag Berlin Heidelberg, 1981).

66. Kostelich, E. J. & Schreiber, T. Noise reduction in chaotic time-series data: A survey of common methods. PHYSICAL REVIEW E VOLUME 48, 1752–1763 (1993).

67. Mønster, D., Fusaroli, R., Tylén, K., Roepstorff, A. & Sherson, J. F. Causal inference from noisy time-series data — Testing the Convergent Cross-Mapping algorithm in the presence of noise and external influence. Future Generation Computer Systems 73, 52–62 (2017).

68. Angrist, J. D., Imbens, G. W. & Rubin, D. B. Identification of Causal Effects Using Instrumental Variables. J. Am. Stat. Assoc. 91, 444–455 (1996).

69. Imbens, G. W. & Angrist, J. D. Identification and Estimation of Local Average Treatment Effects. Econometrica 62, 467–475 (1994).

70. Angrist, J. D. & Pischke, J.-S. Mostly Harmless Econometrics: An Empiricist’s Companion. (Princeton University Press, Princeton, NJ, 2008).

71. Imbens, G. W. & Lemieux, T. Regression discontinuity designs: A guide to practice. J. Econom. 142, 615–635 (2006).

72. Card, D. & Krueger, A. B. Minimum Wages and Employment: A Case Study of the Fast-Food Industry in New Jersey and Pennsylvania. Am. Econ. Rev. 84, 772–793 (1994).

73. Robins, J. M., Ngel Hernán, M. A. & Brumback, B. Marginal Structural Models and Causal Inference in Epidemiology. Epidemiology 11, 550–560 (2000).

74. Robins, J. A NEW APPROACH TO CAUSAL INFERENCE IN MORTALITY STUDIES WITH A SUSTAINED EXPOSURE PERIOD-APPLICATION TO CONTROL OF THE HEALTHY WORKER SURVIVOR EFFECT. Mathematical Modelling 7, 1393–1512 (1986).

75. Greenland, S., Pearl, J. & Robins, J. M. Causal Diagrams for Epidemiologic Research. Epidemiology 10, 37–48 (1999).

76. Hernán, M. A., Hernández-Díaz, S. & Robins, J. M. A structural approach to selection bias. Epidemiology 15, 615–625 (2004).

77. Hernán, M. A. & Robins, J. M. Causal Inference: What If. (Chapman & Hall/CRC, 2024).

78. Abadie, A. & Gardeazabal, J. The Economic Costs of Confict: A Case Study of the Basque Country. Am. Econ. Rev. 93, 113–132 (2003).

79. Abadie, A., Diamond, A. & Hainmueller, A. J. Synthetic control methods for comparative case studies: Estimating the effect of California’s Tobacco control program. J. Am. Stat. Assoc. 105, 493–505 (2010).

80. Little, M., Humphries, S., Patel, K. & Dewey, C. Factors associated with BMI, underweight, overweight, and obesity among adults in a population of rural south India: A crosssectional study. BMC Obes. 3, 12 (2016).

81. Heald, A. et al. A study to investigate genetic factors associated with weight gain in people with diabetes: analysis of polymorphisms in four relevant genes. Adipocyte 12, 2236757 (2023).

82. Tian, X., Thorne, J. L. & Moore, J. B. Ergothioneine: An underrecognised dietary micronutrient required for healthy ageing? British Journal of Nutrition 129, 104–114 (2023).

83. Ranjan, S. & Sharma, P. K. Association of Brain-Derived Neurotrophic factor (BDNF) gene SNP G196A with Type 2 Diabetes and Obesity: A Meta-Analysis. Res. J. Pharm. Technol. 10, 4297–4305 (2017).

84. Robiou-Du-Pont, S. et al. Contribution of 24 obesity-associated genetic variants to insulin resistance, pancreatic beta-cell function and type 2 diabetes risk in the French population. Int. J. Obes. 37, 980–985 (2013).

85. Viñuela, A. et al. Genetic variant effects on gene expression in human pancreatic islets and their implications for T2D. Nat. Commun. 11, 4912 (2020).

86. Su, Y. et al. Periodontitis as a promoting factor of T2D: current evidence and mechanisms. Int. J. Oral Sci. 15, 25 (2023).

87. Alkhalidy, H., Wang, Y. & Liu, D. Dietary flavonoids in the prevention of T2D: An overview. Nutrients 10, 438 (2018).

88. Maurya, V. K. & Aggarwal, M. Factors influencing the absorption of vitamin D in GIT: an overview. J. Food Sci. Technol. 54, 3753–3765 (2017).

89. Zgaga, L. et al. Diet, environmental factors, and lifestyle underlie the high prevalence of vitamin D deficiency in healthy adults in Scotland, and supplementation reduces the proportion that are severely deficient. Journal of Nutrition 141, 1535–1542 (2011).

90. Durmuş, M. Fish oil for human health: Omega-3 fatty acid profiles of marine seafood species. Food Science and Technology (Brazil) 39, 454–461 (2019).

91. Meyer, B. J. et al. Dietary intakes and food sources of omega-6 and omega-3 polyunsaturated fatty acids. Lipids 38, 391–398 (2003).

92. Saini, R. K. & Keum, Y. S. Omega-3 and omega-6 polyunsaturated fatty acids: Dietary sources, metabolism, and significance — A review. Life Sci. 203, 255–267 (2018).

93. Roccisano, D., Kumaratilake, J., Saniotis, A. & Henneberg, M. Dietary Fats and Oils: Some Evolutionary and Historical Perspectives Concerning Edible Lipids for Human Consumption. Food Nutr. Sci. 07, 689–702 (2016).

94. Darwin, C. On the Origin of Species by Means of Natural Selection. (John Murray, London, 1859).

95. Craig-Atkins, E. et al. The dietary impact of the Norman Conquest: A multiproxy archaeological investigation of Oxford, UK. PLoS One 15, e0239640 (2020).

96. Bownes, J. Reassessing the Scottish Mesolithic-Neolithic Transition: Questions of diet and chronology. (University of Glasgow, Glasgow, 2018).

97. Walter, B. S., DeWitte, S. N., Dupras, T. & Beaumont, J. Assessment of nutritional stress in famine burials using stable isotope analysis. Am. J. Phys. Anthropol. 172, 214–226 (2020).

98. Schulting, R. J. et al. AVELINES’S HOLE: AN UNEXPECTED TWIST IN THE TALE. Proceedings of the University of Bristol Spelaeological Society 28, 9–63 (2019).

99. Schulting, R. J. & Richards, M. P. The Wet, the Wild and the Domesticated: the Mesolithic--Neolithic Transition On the West Coast of Scotland. Eur. J. Archaeol. 5, 147–189 (2002).

100. Pickard, C. & Bonsall, C. Post-glacial hunter-gatherer subsistence patterns in Britain: dietary reconstruction using FRUITS. Archaeol. Anthropol. Sci. 12, 142 (2020).

101. Schulting, R. ‘Tilbury man’: A mesolithic skeleton from the lower thames. Proceedings of the Prehistoric Society 79, 19–37 (2013).

102. Jacobi, R. M. & Higham, T. F. G. The early Lateglacial re-colonization of Britain: new radiocarbon evidence from Gough’s Cave, southwest England. Quat. Sci. Rev. 28, 1895–1913 (2009).

103. Hemer, K. A., Lamb, A. L., Chenery, C. A. & Evans, J. A. A multi-isotope investigation of diet and subsistence amongst island and mainland populations from early medieval western Britain. Am. J. Phys. Anthropol. 162, 423–440 (2017).

104. Eckardt, H., Müldner, G. & Speed, G. The Late Roman Field Army in Northern Britain Mobility, Material Culture and Multi-Isotope Analysis at Scorton (N Yorks.). Britannia 46, 191–223 (2015).

105. Moore, J. et al. A multi-isotope (C, N, O, Sr, Pb) study of Iron Age and Roman period skeletons from east Edinburgh, Scotland exploring the relationship between decapitation burials and geographical origins. J. Archaeol. Sci. Rep. 29, 102075 (2020).

106. Halldórsdóttir, H. H. et al. Continuity and individuality in Medieval Hereford, England: A stable isotope approach to bulk bone and incremental dentine. J. Archaeol. Sci. Rep. 23, 800–809 (2019).

107. Amkreutz, L. et al. What lies beneath … Late Glacial human occupation of the submerged North Sea landscape. Antiquity 92, 22–37 (2018).

108. Roberts, P. et al. The men of Nelson’s navy: A comparative stable isotope dietary study of late 18th century and early 19th century servicemen from Royal Naval Hospital burial grounds at Plymouth and Gosport, England. Am. J. Phys. Anthropol. 148, 1–10 (2012).

109. Charlton, S. et al. Finding Britain’s last hunter-gatherers: A new biomolecular approach to ‘unidentifiable’ bone fragments utilising bone collagen. J. Archaeol. Sci. 73, 55–61 (2016).

110. Bonsall, L. A. & Pickard, C. Stable isotope and dental pathology evidence for diet in late Roman Winchester, England. J. Archaeol. Sci. Rep. 2, 128–140 (2015).

111. Gigleux, C., Richards, M. P., Curtis, N., Hutchison, M. & Britton, K. Reconstructing diet at the Neolithic stalled cairn of the Knowe of Rowiegar, Rousay, Orkney, using stable isotope analysis. J. Archaeol. Sci. Rep. 13, 272–280 (2017).

112. Buckberry, J. et al. Finding Vikings in the Danelaw. Oxford Journal of Archaeology 33, 413–434 (2014).

113. Stevens, R. E., Lightfoot, E., Hamilton, J., Cunliffe, B. & Hedges, R. E. M. Investigating dietary variation with burial ritual in iron age hampshire: An isotopic comparison of suddern farm cemetery and danebury hillfort pit burials. Oxford Journal of Archaeology 32, 257–273 (2013).

114. Stevens, R. E., Jacobi, R. M. & Higham, T. F. G. Reassessing the diet of Upper Palaeolithic humans from Gough’s Cave and Sun Hole, Cheddar Gorge, Somerset, UK. J. Archaeol. Sci. 37, 52–61 (2010).

115. Bownes, J., Clarke, L. & Buckberry, J. The importance of animal baselines: Using isotope analysis to compare diet in a British medieval hospital and lay population. J. Archaeol. Sci. Rep. 17, 103–110 (2018).

116. Bell, L. S., Lee Thorp, J. A. & Elkerton, A. The sinking of the Mary Rose warship: a medieval mystery solved? J. Archaeol. Sci. 36, 166–173 (2009).

117. Redfern, R. C., Hamlin, C. & Athfield, N. B. Temporal changes in diet: A stable isotope analysis of late Iron Age and Roman Dorset, Britain. J. Archaeol. Sci. 37, 1149–1160 (2010).

118. Lamb, A. L., Evans, J. E., Buckley, R. & Appleby, J. Multi-isotope analysis demonstrates significant lifestyle changes in King Richard III. J. Archaeol. Sci. 50, 559–565 (2014).

119. Stevens, R. E., Lightfoot, E., Hamilton, J., Cunliffe, B. & Hedges, R. E. M. Stable Isotope Investigations Of The Danebury Hillfort Pit Burials. Oxford Journal of Archaeology 29, 407–428 (2010).

120. Richards, M. P., Hedges, R. E. M., Jacobi, R., Current, A. & Stringer, C. FOCUS: Gough’s Cave and Sun Hole Cave Human Stable Isotope Values Indicate a High Animal Protein Diet in the British Upper Palaeolithic. J. Archaeol. Sci. 27, 1–3 (2000).

121. Richards, M. P., Jacobi, R., Cook, J., Pettitt, P. B. & Stringer, C. B. Isotope evidence for the intensive use of marine foods by Late Upper Palaeolithic humans. J. Hum. Evol. 49, 390–394 (2005).

122. Privat, K. L., O’connell, T. C. & Richards, M. P. Stable isotope analysis of human and faunal remains from the Anglo-Saxon Cemetery and Berinsfield, Oxfordshire: Dietary and social implications. J. Archaeol. Sci. 29, 779–790 (2002).

123. Jay, M. Iron Age Diet at Glastonbury Lake Village: The isotopic evidence for negligible aquatic resource consumption. Oxford Journal of Archaeology 27, 201–216 (2008).

124. Hedges, R. E. M., Stevens, R. E. & Pearson, J. A. Carbon and Nitrogen Stable Isotope Compositions of Animal and Human Bone. in Building Memories: The Neolithic Cotswold Long Barrow at Ascott-under-Wychwood, Oxfordshire (eds. Benson, D. & Whittle, A.) 239–246 (Oxbow Books, Oxford, 2007).

125. Chenery, C. A., Evans, J. A., Score, D. & Boyle, A. A Boat Load of Vikings? Journal of the North Atlantic 7, 43–53 (2014).

126. Dhaliwal, K., Rando, C., Reade, H., Jourdan, A. L. & Stevens, R. E. Socioeconomic differences in diet: An isotopic examination of post-Medieval Chichester, West Sussex. Am. J. Phys. Anthropol. 171, 584–597 (2020).

127. Richards, M. Chapter 12 Human consumption of plant foods in the British Neolithic: Direct evidence from bone stable isotopes. in Plants in Neolithic Britain and Beyond (ed. Fairburn, A. S.) vol. 5 123–136 (Oxbow Books, Oxford, 2000).

128. Schulting, R. J., Chapman, M. & Chapman, E. J. AMS 14 C DATING AND STABLE ISOTOPE (CARBON, NITROGEN) ANALYSIS OF AN EARLIER NEOLITHIC HUMAN SKELETAL ASSEMBLAGE FROM HAY WOOD CAVE, MENDIP, SOMERSET. Proceedings of the University of Bristol Spelaeological Society 26, 9–26 (2013).

129. Schulting, R., Sheridan, A., Crozier, R. & Murphy, E. Revisiting Quanterness: new AMS dates and stable isotope data from an Orcadian chamber tomb. Proceedings of the Society of Antiquaries of Scotland 140, 1–50 (2010).

130. Schulting, R. J., Gardiner, P. J., Hawkes, C. J. & Murray, E. THE MESOLITHIC AND NEOLITHIC HUMAN BONE ASSEMBLAGE FROM TOTTY POT, CHEDDAR, SOMERSET. Proceedings of the University of Bristol Spelaeological Society 25, 75–95 (2010).

131. Richards, M. P. & Sheridan, J. A. New AMS dates on human bone from Mesolithic Oronsay. Antiquity 74, 313–315 (2000).

132. O’Connell, T. C. & Lawler, A. Chapter 5. Economic Resources VI. Stable isotope analysis of human and faunal remains. in The Anglo-Saxon Settlement and Cemetery at Bloodmoor Hill, Carlton Colville, Suffolk (eds. Lucy, S., Tipper, J. & Dickens, A.) vol. EAA 131 317–321 (East Anglian Archaeology, Barnsley, 2009).

133. Moore, F. E. Diet and subsistence in the Anglo-Saxon Trent Valley: a stable isotope investigation of Broughton Lodge Anglo-Saxon cemetery, Nottinghamshire. (University of Nottingham, Nottingham, 2017).

134. Millard, A. R. et al. Scottish soldiers from the Battle of Dunbar 1650: A prosopographical approach to a skeletal assemblage. PLoS One 15, e0243369 (2020).

135. Macpherson, P. M. Tracing Change: An Isotopic Investigation of Anglo-Saxon Childhood Diet. (University of Sheffield, Sheffield, 2005).

136. Lucy, S. et al. The Burial of A Princess? The Later Seventh-Century Cemetery At Westfield Farm, Ely. The Antiquaries Journal 89, 81–141 (2009).

137. Jay, M. & Richards, M. P. British iron age diet: Stable isotopes and other evidence. Proceedings of the Prehistoric Society 73, 169–190 (2007).

138. Redfern, R. C. et al. Going south of the river: A multidisciplinary analysis of ancestry, mobility and diet in a population from Roman Southwark, London. J. Archaeol. Sci. 74, 11–22 (2016).

139. Scorrer, J. et al. Diversity aboard a Tudor warship: Investigating the origins of the Mary Rose crew using multi-isotope analysis. R. Soc. Open Sci. 8, 202106 (2021).

140. Cummings, C. & Hedges, R. Chapter 5: Human remains: Carbon and nitrogen stable isotope analysis. in The late Roman cemetery at Lankhills, Winchester: Excavations 2000-2005 (eds. Booth, P. et al.) vol. 10 411–421 (Oxford Archaeology Monograph, Oxford, 2010).

141. Chenery, C., Müldner, G., Evans, J., Eckardt, H. & Lewis, M. Strontium and stable isotope evidence for diet and mobility in Roman Gloucester, UK. J. Archaeol. Sci. 37, 150–163 (2010).

142. Chenery, C., Eckardt, H. & Müldner, G. Cosmopolitan Catterick? Isotopic evidence for population mobility on Rome’s Northern frontier. J. Archaeol. Sci. 38, 1525–1536 (2011).

143. Montgomery, J. et al. Strategic and sporadic marine consumption at the onset of the Neolithic: Increasing temporal resolution in the isotope evidence. Antiquity 87, 1060–1072 (2013).

144. Redfern, R. C., Millard, A. R. & Hamlin, C. A regional investigation of subadult dietary patterns and health in late Iron Age and Roman Dorset, England. J. Archaeol. Sci. 39, 1249–1259 (2012).

145. Müldner, G., Chenery, C. & Eckardt, H. The ‘Headless Romans’: Multi-isotope investigations of an unusual burial ground from Roman Britain. J. Archaeol. Sci. 38, 280–290 (2011).

146. Hannah, E. L., McLaughlin, T. R., Keaveney, E. M. & Hakenbeck, S. E. Anglo-Saxon diet in the Conversion period: A comparative isotopic study using carbon and nitrogen. J. Archaeol. Sci. Rep. 19, 24–34 (2018).

147. Müldner, G. Stable isotopes and diet: Their contribution to Romano-British research. Antiquity 87, 137–149 (2013).

148. Müldner, G. et al. Isotopes and individuals: Diet and mobility among the medieval Bishops of Whithorn. Antiquity 83, 1119–1133 (2009).

149. Pollard, A. M. et al. ‘These boots were made for walking’: The isotopic analysis of a C4 Roman inhumation from Gravesend, Kent, UK. Am. J. Phys. Anthropol. 146, 446–456 (2011).

150. Müldner, G. & Richards, M. P. Diet and diversity at Later Medieval fishergate: The isotopic evidence. Am. J. Phys. Anthropol. 134, 162–174 (2007).

151. Lamb, A. L., Melikian, M., Ives, R. & Evans, J. Multi-isotope analysis of the population of the lost medieval village of Auldhame, East Lothian, Scotland. J. Anal. At. Spectrom. 27, 765–777 (2012).

152. Richards, M. P., Pettitt, P. B., Stiner, M. C. & Trinkaus, E. Stable isotope evidence for increasing dietary breadth in the European mid-Upper Paleolithic. Proc. Natl. Acad. Sci. U. S. A. 98, 6528–6532 (2001).

153. Mays, S. & Beavan, N. An investigation of diet in early Anglo-Saxon England using carbon and nitrogen stable isotope analysis of human bone collagen. J. Archaeol. Sci. 39, 867–874 (2012).

154. Jay, M. & Richards, M. P. Diet in the Iron Age cemetery population at Wetwang Slack, East Yorkshire, UK: Carbon and nitrogen stable isotope evidence. J. Archaeol. Sci. 33, 653–662 (2006).

155. Burt, N. M. Stable isotope ratio analysis of breastfeeding and weaning practices of children from medieval Fishergate House York, UK. Am. J. Phys. Anthropol. 152, 407–416 (2013).

156. Hedges, R., Saville, A. & O’Connell, T. Characterizing the diet of individuals at the Neolithic chambered tomb of Hazleton North, Gloucestershire, England, using stable isotopic analysis. Archaeometry 50, 114–128 (2008).

157. Haydock, H., Clarke, L., Craig-Atkins, E., Howcroft, R. & Buckberry, J. Weaning at Anglo-Saxon raunds: Implications for changing breastfeeding practice in britain over two millennia. Am. J. Phys. Anthropol. 151, 604–612 (2013).

158. Fuller, B. T., Molleson, T. I., Harris, D. A., Gilmour, L. T. & Hedges, R. E. M. Isotopic evidence for breastfeeding and possible adult dietary differences from Late/Sub-Roman Britain. Am. J. Phys. Anthropol. 129, 45–54 (2006).

159. Cheung, C., Schroeder, H. & Hedges, R. E. M. Diet, social differentiation and cultural change in Roman Britain: New isotopic evidence from Gloucestershire. Archaeol. Anthropol. Sci. 4, 61–73 (2012).

160. Armit, I., Shapland, F., Montgomery, J. & Beaumont, J. Difference in Death? A Lost Neolithic Inhumation Cemetery with Britain’s Earliest Case of Rickets, at Balevullin, Western Scotland. Proceedings of the Prehistoric Society 81, 199–214 (2015).

161. Britton, K. et al. Isotopes and new norms: Investigating the emergence of early modern U.K. breastfeeding practices at St. Nicholas Kirk, Aberdeen. Int. J. Osteoarchaeol. 28, 510–522 (2018).

162. Brunning, R. An Early Mesolithic Cemetery at Greylake, Somerset, UK. Archaeology in the Severn Estuary 22, 67–70 (2013).

163. Schulting, R. J. ‘… PURSUING A RABBIT IN BURRINGTON COMBE’: NEW RESEARCH ON THE EARLY MESOLITHIC BURIAL CAVE OF AVELINE’S HOLE. Proceedings of the University of Bristol Spelaeological Society 23, 171–265 (2005).

164. Jones, J. R. & Mulville, J. Isotopic and zooarchaeological approaches towards understanding aquatic resource use in human economies and animal management in the prehistoric Scottish North Atlantic Islands. J. Archaeol. Sci. Rep. 6, 665–677 (2016).

165. Jones, J. R., Mulville, J. A., McGill, R. A. R. & Evershed, R. P. Palaeoenvironmental modelling of δ13C and δ15N values in the North Atlantic Islands: Understanding past marine resource use. Rapid Communications in Mass Spectrometry 26, 2399–2406 (2012).

166. Jones, J. R., Mulville, J. & Evershed, Richard. P. Fruits of the sea: investigating marine resource use in the North Atlantic islands. in Ancient Maritime Communities and the Relationship between People and Environment along the European Atlantic Coasts (eds. Daire, M. Y. et al.) 501–511 (Archaeopress, Oxford, 2013).

167. Mulville, J. et al. Isotopic Analysis of Faunal Material from South Uist, Western Isles, Scotland. Journal of the North Atlantic 2, 51–59 (2009).

168. Schulting, R. J., Vaiglova, P., Crozier, R. & Reimer, P. J. Further isotopic evidence for seaweed-eating sheep from Neolithic Orkney. J. Archaeol. Sci. Rep. 11, 463–470 (2017).

169. Jones, G. Chapter 7 Evaluating the importance of cultivation and collecting in Neolithic Britain. in Plants in Neolithic Britain and beyond (ed. Fairburn, A. S.) vol. 5 79–84 (Oxbow Books, Oxford, 2000).

170. Russell, N. Marine Radiocarbon Reservoir Effects (MRE) in Archaeology: Temporal and Spatial Changes through the Holocene within the UK Coastal Environment. (University of Glasgow, Glasgow, 2011).

171. Hammond, C. & O’Connor, T. Pig diet in medieval York: Carbon and nitrogen stable isotopes. Archaeol. Anthropol. Sci. 5, 123–127 (2013).

172. Hutchinson, W. F. et al. The globalization of naval provisioning: Ancient DNA and stable isotope analyses of stored cod from the wreck of the Mary Rose, AD 1545. R. Soc. Open Sci. 2, 150199 (2015).

173. Jones, J. R. & Mulville, J. A. Norse Animal Husbandry in Liminal Environments: Stable Isotope Evidence from the Scottish North Atlantic Islands. Environmental Archaeology 23, 338–351 (2018).

174. Barrett, J. H. et al. Interpreting the expansion of sea fishing in medieval Europe using stable isotope analysis of archaeological cod bones. J. Archaeol. Sci. 38, 1516–1524 (2011).

175. Orton, D. C. et al. Stable isotope evidence for late medieval (14th-15th C) origins of the eastern Baltic cod (Gadus morhua) fishery. PLoS One 6, e27568 (2011).

176. Nehlich, O. et al. Application of sulphur isotope ratios to examine weaning patterns and freshwater fish consumption in Roman Oxfordshire, UK. Geochim. Cosmochim. Acta 75, 4963–4977 (2011).

177. Madgwick, R., Mulville, J. & Stevens, R. E. Diversity in foddering strategy and herd management in late Bronze Age Britain: An isotopic investigation of pigs and other fauna from two midden sites. Environmental Archaeology 17, 126–140 (2012).

178. Kancle, L., Montgomery, J., Gröcke, D. R. & Caffell, A. From field to fish: Tracking changes in diet on entry to two medieval friaries in northern England. J. Archaeol. Sci. Rep. 22, 264–284 (2018).

179. Hamilton, W. D., Sayle, K. L., Boyd, M. O. E., Haselgrove, C. C. & Cook, G. T. ‘Celtic cowboys’ reborn: Application of multi-isotopic analysis (δ 13 C, δ 15 N, and δ 34 S) to examine mobility and movement of animals within an Iron Age British society. J. Archaeol. Sci. 101, 189–198 (2019).

180. Schulting, R. J. et al. The ups & downs of Iron Age animal management on the Oxfordshire Ridgeway, south-central England: A multi-isotope approach. J. Archaeol. Sci. 101, 199–212 (2019).

181. Millard, A. R., Jimenez-Cano, N. G., Lebrasseur, O. & Sakai, Y. Isotopic Investigation of Animal Husbandry in the Welsh and English Periods at Dryslwyn Castle, Carmarthenshire, Wales. Int. J. Osteoarchaeol. 23, 640–650 (2013).

182. Stevens, R. E., Lightfoot, E., Hamilton, J., Cunliffe, B. W. & Hedges, R. E. M. One for the master and one for the dame: Stable isotope investigations of Iron Age animal husbandry in the Danebury Environs. Archaeol. Anthropol. Sci. 5, 95–109 (2013).

183. Sakai, Y. Transition from the Late Roman Period to the Early Anglo-Saxon Period in the Upper Thames Valley Based on Stable Isotopes. (University of Oxford, Oxford, 2017).

184. Knapp, Z. The Zooarchaeology of the Anglo-Saxon Christian Conversion: Lyminge, a case study. (University of Reading, Reading, 2018).

185. Blanz, M. et al. Identifying seaweed consumption by sheep using isotope analysis of their bones and teeth: Modern reference δ13C and δ15N values and their archaeological implications. J. Archaeol. Sci. 118, 105140 (2020).

186. Britton, K., Müldner, G. & Bell, M. Stable isotope evidence for salt-marsh grazing in the Bronze Age Severn Estuary, UK: implications for palaeodietary analysis at coastal sites. J. Archaeol. Sci. 35, 2111–2118 (2008).

187. Towers, J., Jay, M., Mainland, I., Nehlich, O. & Montgomery, J. A calf for all seasons? The potential of stable isotope analysis to investigate prehistoric husbandry practices. J. Archaeol. Sci. 38, 1858–1868 (2011).

188. Sykes, N. J. et al. New evidence for the establishment and management of the European fallow deer (Dama dama dama) in Roman Britain. J. Archaeol. Sci. 38, 156–165 (2011).

189. Lightfoot, E. & Stevens, R. E. Stable isotope investigations of charred barley (Hordeum vulgare) and wheat (Triticum spelta) grains from Danebury Hillfort: implications for palaeodietary reconstructions. J. Archaeol. Sci. 39, 656–662 (2012).

190. Lodwick, L., Campbell, G., Crosby, V. & Müldner, G. Isotopic Evidence for Changes in Cereal Production Strategies in Iron Age and Roman Britain. Environmental Archaeology 26, 13–28 (2021).

191. Richards, M. 7 Human remains and diet: Stable isotope values. in HAMBLEDON HILL, DORSET, ENGLAND. Excavation and survey of a Neolithic monument complex and its surrounding landscape (eds. Mercer, R. & Healy, F.) vol. 2 522–527 (Liverpool University Press, Historic England, Liverpool, 2008).

192. Hamerow, H. et al. An Integrated Bioarchaeological Approach to the Medieval ‘Agricultural Revolution’: A Case Study from Stafford, England, c. ad 800-1200. Eur. J. Archaeol. 23, 585–609 (2020).

193. Jay, M., Haselgrove, C., Hamilton, D., Hill, J. D. & Dent, J. Chariots and context: New radiocarbon dates from wetwang and the chronology of iron age burials and brooches in East Yorkshire. Oxford Journal of Archaeology 31, 161–189 (2012).

194. Bayliss, A. et al./person-group>. 7. Interpreting Chronology: The Radiocarbon Dating Programme. in Building Memories: The Neolithic Cotswold Long Barrow at Ascott-under-Wychwood, Oxfordshire (eds. Benson, D. & Whittle, A.) 221–236 (Oxbow Books, Oxford, 2007).

195. Barrett, J. H., Beukens, R. P. & Brothwell, D. R. Radiocarbon dating and marine reservoir correction of Viking Age Christian burials from Orkney. Antiquity 74, 537–543 (2000).

196. Hemer, K. A., Evans, J. A., Chenery, C. A. & Lamb, A. L. No Man is an island: Evidence of pre-Viking Age migration to the Isle of Man. J. Archaeol. Sci. 52, 242–249 (2014).

197. Groom, P. et al. Two early medieval cemeteries in Pembrokeshire: Brownslade Barrow and West Angle Bay. Archaeologia Cambrensis 160, 133–203 (2011).

198. Cunliffe, B. Danebury Ring Hillfort – The Main Phases. Digital Digging https://www.digitaldigging.net/danebury-ring-hillfort-main-phases/ (2013).

199. Hedges, R. E. M., Housley, R. A., Bronk, C. R. & Klinken, G. J. V. RADIOCARBON DATES FROM THE OXFORD AMS SYSTEM: ARCHAEOMETRY DATELIST 13. Archaeometry 33, 279–296 (1991).

200. Kirby, M. Excavations at Musselburgh Primary Health Care Centre: Iron Age and Roman discoveries to the north of Inveresk Roman Fort, East Lothian. Scottish Archaeological Internet Reports 89, 1–153 (2020).

201. Holbrook, N. & Thomas, A. An Early-medieval monastic cemetery at Llandough, Glamorgan: Excavations in 1994. Mediev. Archaeol. 49, 1–92 (2005).

202. Carver, M. Portmahomack: Monastery of the Picts. (Edinburgh University Press, Edinburgh, 2016).

203. Dulias, K. et al. Ancient DNA at the edge of the world: Continental immigration and the persistence of Neolithic male lineages in Bronze Age Orkney. Proc. Natl. Acad. Sci. U. S. A. 119, e2108001119 (2022).

204. Allentoft, M. E. et al. Population genomics of post-glacial western Eurasia. Nature 625, 301–311 (2024).

205. Gretzinger, J. et al. The Anglo-Saxon migration and the formation of the early English gene pool. Nature 610, 112–119 (2022).

206. Rohland, N. et al. Three assays for in-solution enrichment of ancient human DNA at more than a million SNPs. Genome Res. 32, 2068–2078 (2022).

207. Margaryan, A. et al. Population genomics of the Viking world. Nature 585, 390–396 (2020).

208. Olalde, I. et al. The Beaker phenomenon and the genomic transformation of northwest Europe. Nature 555, 190–196 (2018).

209. Patterson, N. et al. Large-scale migration into Britain during the Middle to Late Bronze Age. Nature 601, 588–594 (2022).

210. Scheib, C. L. et al. East Anglian early Neolithic monument burial linked to contemporary Megaliths. Ann. Hum. Biol. 46, 145–149 (2019).

211. Martiniano, R. et al. Genomic signals of migration and continuity in Britain before the Anglo-Saxons. Nat. Commun. 7, 10326 (2016).

212. Fowler, C. et al. A high-resolution picture of kinship practices in an Early Neolithic tomb. Nature 601, 584–587 (2022).

213. Brace, S. et al. Ancient genomes indicate population replacement in Early Neolithic Britain. Nat. Ecol. Evol. 3, 765–771 (2019).

214. Sánchez-Quinto, F. et al. Megalithic tombs in western and northern Neolithic Europe were linked to a kindred society. Proc. Natl. Acad. Sci. U. S. A. 116, 9469–9474 (2019).

215. Brace, S. et al. Genomes from a medieval mass burial show Ashkenazi-associated hereditary diseases pre-date the 12th century. Current Biology 32, 4350–4359.e6 (2022).

216. Schiffels, S. et al. Iron Age and Anglo-Saxon genomes from East England reveal British migration history. Nat. Commun. 7, 10408 (2016).

217. Mallick, S. & Reich, D. The Allen Ancient DNA Resource (AADR): A curated compendium of ancient human genomes. Harvard Dataverse, V9 data release 10.7910/DVN/FFIDCW (2024).

218. Mallick, S. et al. The Allen Ancient DNA Resource (AADR) a curated compendium of ancient human genomes. Sci. Data 11, (2024).

219. Li, H. et al. The Sequence Alignment/Map format and SAMtools. Bioinformatics 25, 2078–2079 (2009).

220. Dulias, K. et al. Ancient DNA at the edge of the world: Continental immigration and the persistence of Neolithic male lineages in Bronze Age Orkney. Proc. Natl. Acad. Sci. U. S. A. 119, e2108001119 (2022).

221. Li, H. & Durbin, R. Fast and accurate short read alignment with Burrows-Wheeler transform. Bioinformatics 25, 1754–1760 (2009).

222. Schubert, M. et al. Improving ancient DNA read mapping against modern reference genomes. BMC Genomics 13, 178 (2012).

223. Peltzer, A. et al. EAGER: efficient ancient genome reconstruction. Genome Biol. 17, 60 (2016).

224. Fernandes, R., Millard, A. R., Brabec, M., Nadeau, M. J. & Grootes, P. Food reconstruction using isotopic transferred signals (FRUITS): A bayesian model for diet reconstruction. PLoS One 9, (2014).

225. Tieszen, L. L. & Fagre, T. Effect of diet quality and composition on the isotopic composition of respiratory CO2, bone collagen, bioapatite, and soft tissues. in Prehistoric Human Bone —— Archaeology at the Molecular Level (eds. Lambert, J. B. & Grupe, G.) 121–155 (Springer Berlin, Heidelberg, 1993).

226. Ambrose, S. H. & Norr, L. Experimental evidence for the relationship of the carbon isotope ratios of whole diet and dietary protein to those of bone collagen and carbonate. in Prehistoric Human Bone —— Archaeology at the Molecular Level (eds. Lambert, J. B. & Grupe, G.) 1–38 (Springer Berlin, Heidelberg, 1993).

227. Jim, S., Ambrose, S. H. & Evershed, R. P. Stable carbon isotopic evidence for differences in the dietary origin of bone cholesterol, collagen and apatite: Implications for their use in palaeodietary reconstruction. Geochim. Cosmochim. Acta 68, 61–72 (2004).

228. Howland, M. R. et al. Expression of the dietary isotope signal in the compound-specific δ13C values of pig bone lipids and amino acids. Int. J. Osteoarchaeol. 13, 54–65 (2003).

229. Warinner, C. & Tuross, N. Alkaline cooking and stable isotope tissue-diet spacing in swine: archaeological implications. J. Archaeol. Sci. 36, 1690–1697 (2009).

230. Fernandes, R., Nadeau, M. J. & Grootes, P. M. Macronutrient-based model for dietary carbon routing in bone collagen and bioapatite. Archaeol. Anthropol. Sci. 4, 291–301 (2012).

231. Fernandes, R., Grootes, P., Nadeau, M. J. & Nehlich, O. Quantitative diet reconstruction of a Neolithic population using a Bayesian mixing model (FRUITS): The case study of Ostorf (Germany). Am. J. Phys. Anthropol. 158, 325–340 (2015).

232. Grande, F., Ueda, Y., Masangwi, S. & Holmes, B. Global Nutrient Conversion Table for FAO Supply Utilization Accounts. http://www.fao.org/documents/card/en/c/cc9678en (2024) doi:10.4060/cc9678en.

233. Plummer, M. JAGS: A program for analysis of Bayesian graphical models using Gibbs sampling. in Proceedings of the 3rd International Workshop on Distributed Statistical Computing (DSC 2003) (Vienna, 2003).

234. Felsenstein, J. THEORETICAL EVOLUTIONARY GENETICS. (2019).

235. Taus, T., Futschik, A. & Schlötterer, C. Quantifying Selection with Pool-Seq Time Series Data. Mol. Biol. Evol. 34, 3023–3034 (2017).

236. Wood, S. Generalized Additive Models: An Introduction with R. (chapman and hall/CRC, 2017).

237. Kovács, L. Feature selection algorithms in generalized additive models under concurvity. Comput. Stat. 39, 461–493 (2024).

238. Runge, J. et al. Inferring causation from time series in Earth system sciences. Nat. Commun. 10, 2553 (2019).

239. Shojaie, A. & Fox, E. B. Granger Causality: A Review and Recent Advances. Annu. Rev. Stat. Appl. 9, 289–319 (2022).

240. Ying, X. et al. Continuity Scaling: A Rigorous Framework for Detecting and Quantifying Causality Accurately. Research 2022, 9870149 (2022).

241. Shimizu, S., Hoyer, P. O. & Hyvärinen, A. A Linear Non-Gaussian Acyclic Model for Causal Discovery Antti Kerminen. Journal of Machine Learning Research 7, 2003–2030 (2006).

242. Runge, J., Nowack, P., Kretschmer, M., Flaxman, S. & Sejdinovic, D. Detecting and quantifying causal associations in large nonlinear time series datasets. Sci. Adv 5, eaau4996 (2019).

243. Granger, C. W. J. Investigating Causal Relations by Econometric Models and Cross-spectral Methods. Econometrica 37, 424–438 (1969).

244. Hoyer, P. O., Janzing, D., Mooij, J., Peters, J. & Schölkopf, B. Nonlinear causal discovery with additive noise models. in Advances in Neural Information Processing Systems (Curran Associates, Inc., 2008).

245. Pearl, Judea. Causal Inference in Statistics : A Primer. (John Wiley & Sons Ltd, 2016).

246. Imbens, G. & Rubin, D. B. Causal Inference for Statistics, Social, and Biomedical Sciences: An Introduction. (Cambridge University Press, Cambridge, 2015).

247. Pearl, J. Causality: Models, Reasoning and Inference. (Cambridge University Press, Cambridge, 2009).

248. Rubin, D. B. Practical Implications of Modes of Statistical Inference for Causal Effects and the Critical Role of the Assignment Mechanism. Biometrics 47, 1213–1234 (1991).

249. Rubin, D. B. Comment: Neyman (1923) and Causal Inference in Experiments and and Observational Studies. Statistical Science 5, 472–480 (1990).

250. Rubin, D. B. Bayesian Inference for Causal Effects: The Role of Randomization. The Annals of Statistics 6, 34–58 (1978).

251. Rubin, D. B. ESTIMATING CAUSAL EFFECTS OF TREATMENTS IN RANDOMIZED AND NONRANDOMIZED STUDIES. J. Educ. Psychol. 66, 688–701 (1974).

252. Rosenbaum, P. R. & Rubin, D. B. The central role of the propensity score in observational studies for causal effects. Biometrika 70, 41–55 (1983).

253. Pearl, J. Causal diagrams for empirical research. Biometrika 82, 669–710 (1995).

254. Basu, D. Randomization Analysis of Experimental Data: The Fisher Randomization Test. J. Am. Stat. Assoc. 75, 575–582 (1980).

