## Supplementary material for "Did dietary change drive natural selection? A paleo-empirical evaluation across 6,000 years in Britain": SI

### Section 1 The matching criteria for each SNP and its dietary variables

Among the 14 SNPs included in our analyses, nine loci are considered to be associated with $C_3$ plant consumption: rs12401678, rs2010410, rs272872, rs3891176, rs4073089, rs6265, rs653178, rs7517, and rs7775397. Among these nine SNPs, the phenotypes linked to rs12401678 (circulating glucose levels[1]), rs3891176, rs653178, and rs7775397 (celiac disease susceptibility[2,3]), rs4073089 and rs7517 (insulin secretion; type 2 diabetes[1]), and rs6265 (type 2 diabetes[1]) are all directly related to $C_3$ plant consumption. The phenotypes of these loci are particularly associated with the metabolic and physiological processes involved in the consumption of carbohydrate-rich foods, with no clear mechanistic connection to the other four ancient dietary categories considered in this study (terrestrial herbivores, terrestrial omnivores, marine fish, and freshwater fish).

The remaining two SNPs, rs2010410 (obesity[1]) and rs272872 (ergothioneine levels[3]), do not exhibit direct links to $C_3$ plant consumption. Their inclusion within the $C_3$-associated group reflects, to a large extent, the limitations of our dietary reconstruction model. Obesity is a phenotype influenced by numerous factors[4,5], including physical activity, vegetable intake, and the consumption of fatty meat. Even when considering dietary factors alone, our model cannot explicitly estimate vegetable consumption or distinguish between lean and fat-rich meat intake. As a result, a high-carbohydrate diet represented by $C_3$ plant consumption becomes the most plausible dietary factor related to this phenotype within our modelling framework. A similar rationale applies to ergothioneine levels. Although fungi are the primary dietary source of ergothioneine[6], our model cannot directly incorporate fungal consumption. Within the available dietary categories, $C_3$ plants contain the highest ergothioneine values, making them the closest match for rs272872 under the constraints of our model.

Dairy consumption is considered the putative dietary driver for rs4988235, the locus associated with lactase persistence[3,7]. For SNPs involved in fatty-acid metabolism (rs174546, rs174570, rs174594)[3,8], the potential drivers are taken to be the consumption of marine fish and terrestrial meat (i.e., terrestrial herbivores and omnivores). Marine fish are rich in omega-3 fatty acids[9], whereas terrestrial meat represents a major dietary source of omega-6 fatty acids[10]. Both classes of polyunsaturated fatty acids are key components of fatty-acid metabolic pathways[11], providing a biologically plausible link between these dietary resources and the selection acting on the relevant SNPs. rs7944926 (vitamin D levels[3]) is considered to be associated with the consumption of marine fish, which is the most vitamin-D–rich food category in our dietary model[12,13].

### Section 2 Dietary reconstruction for individuals

In this section, we present the dietary reconstruction results for each human individual. The figures in this section show the posterior mean estimates, while the uncertainty of the parameter estimates is presented in Supplementary Data 3. The results are organized chronologically, covering periods from the Paleolithic to the Post-Medieval. For each period, the data are further categorized by region: England, Scotland, and Wales. To simplify references to individual samples, we use the identifiers in Table S1, as the original Sample IDs may be overly complex. The correspondence between these sample identifiers and their full Sample IDs, along with associated archaeological information, is provided in Supplementary Data 3. We recommend reading the Methods section of the main text, as well as Section 6, before proceeding with this section.

**Table S1| Sample identifiers and their corresponding periods, regions, and materials. Samples from the Beaker 1st stage (2500–1500 BC) and the Beaker 2nd stage (1500–750 BC) derive from the Beaker People Project and the Beakers and Bodies Project**

| Sample identifiers | Period | Region |
|---|---|---|
| Sample_PAE<number> | Paleolithic | England |
| Sample_PAW<number> | Paleolithic | Wales |
| Sample_MLE<number> | Mesolithic | England |
| Sample_MLW<number> | Mesolithic | Wales |
| Sample_NLS<number> | Neolithic | Scotland |
| Sample_NLE<number> | Neolithic | England |
| Sample_NLW<number> | Neolithic | Wales |
| Sample_BAS<number> | Bronze Age | Scotland |
| Sample_BAE<number> | Bronze Age | England |
| Sample_BKCNbone<number> | Beaker 1st stage | Central/Northern England |
| Sample_BKSbone<number> | Beaker 1st stage | Scotland |
| Sample_BASEbone<number> | Beaker 1st stage | Southern England |
| Sample_BKWbone<number> | Beaker 1st stage | Wales |
| Sample_BKCNdentine<number> | Beaker 1st stage | Central/Northern England |
| Sample_BKSdentine<number> | Beaker 1st stage | Scotland |
| Sample_BASEdentine<number> | Beaker 1st stage | Southern England |
| Sample_BKWdentine<number> | Beaker 1st stage | Wales |
| Sample_BABK<number> | Beaker 2nd stage | All |
| Sample_IAS<number> | Iron Age | Scotland |
| Sample_IANE<number> | Iron Age | Northern England |
| Sample_IASE<number> | Iron Age | Southern England |
| Sample_ROS<number> | Roman Iron Age | Scotland |
| Sample_RONE<number> | Roman | Northern England |
| Sample_ROSE<number> | Roman | Southern England |
| Sample_EMS<number> | Early Medieval | Scotland |

| | | |
|---|---|---|
| Sample_EMCN<number> | Early Medieval | Central/Northern England |
| Sample_EMSE<number> | Early Medieval | Southern England |
| Sample_EMW<number> | Early Medieval | Wales |
| Sample_LMS<number> | Later Medieval | Scotland |
| Sample_LMCN<number> | Later Medieval | Central/Northern England |
| Sample_LMSEbone<number> | Later Medieval | Southern England |
| Sample_ LMSEincremental <number> | Later Medieval | Southern England |
| Sample_PMS<number> | Post-Medieval | Scotland |
| Sample_PMNE<number> | Post-Medieval | Northern England |
| Sample_PMSE <number> | Post-Medieval | Southern England |

#### 2A. Paleolithic period

**England**

The Paleolithic individuals[14–18] from England primarily relied on terrestrial resources, with diets mainly based on herbivores, supplemented by small amounts of omnivores and plants (Fig. S1). This dietary pattern reflects the foraging and hunting lifestyle typical of this period. However, two individuals from Doggerland (Sample_PAE1 and 2) show evidence of freshwater resource consumption. These two individuals also exhibit significantly higher $\delta^{15}N$ values compared to the others (see Supplementary Data 3).

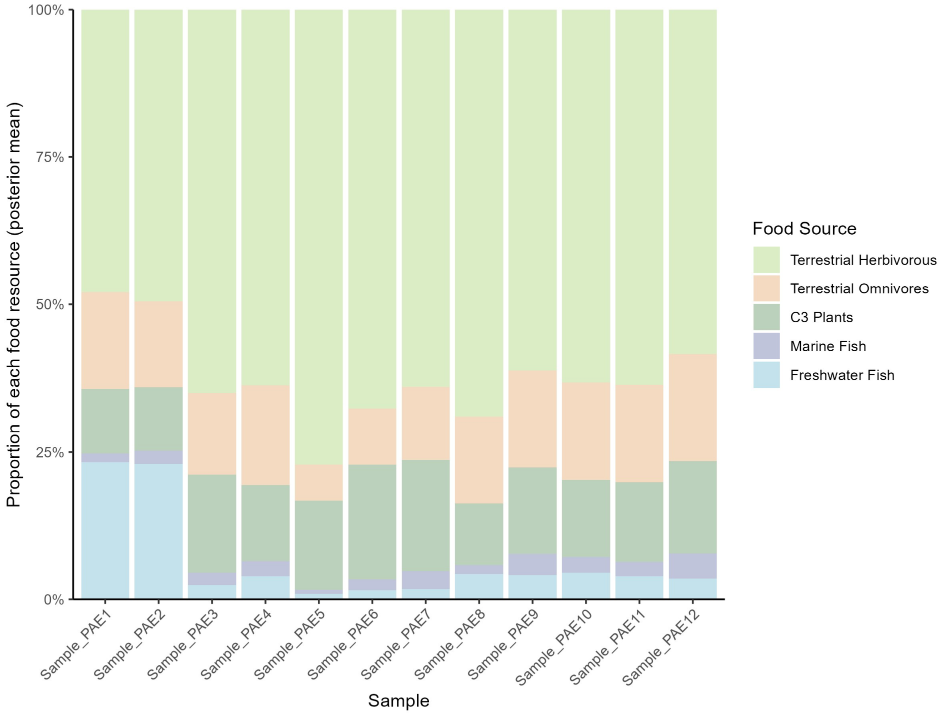

**Fig. S1| Dietary reconstruction for Paleolithic England human samples.**

##### Wales

The subsistence strategy in Paleolithic Wales appears somewhat different (Fig. S2). Samples_PAW1 – 5[17,19] show evidence of marine resource consumption. These individuals, recovered from Kendrick's Cave, come from a location likely situated very close to the sea, making it reasonable that marine resources contributed significantly to their diet. In contrast, Sample_PAW6[20] (from Paviland Cave), which was also likely situated near the coast, shows less reliance on marine resources, although still higher than the levels observed in the samples from England. The $\delta^{15}N$ value of Sample_PAW6 is also lower than those of the other five samples (Supplementary Data 3).

The dietary reconstruction results indicate that the subsistence strategy during the Paleolithic was primarily based on foraging and hunting, with dietary patterns largely determined by local environmental conditions.

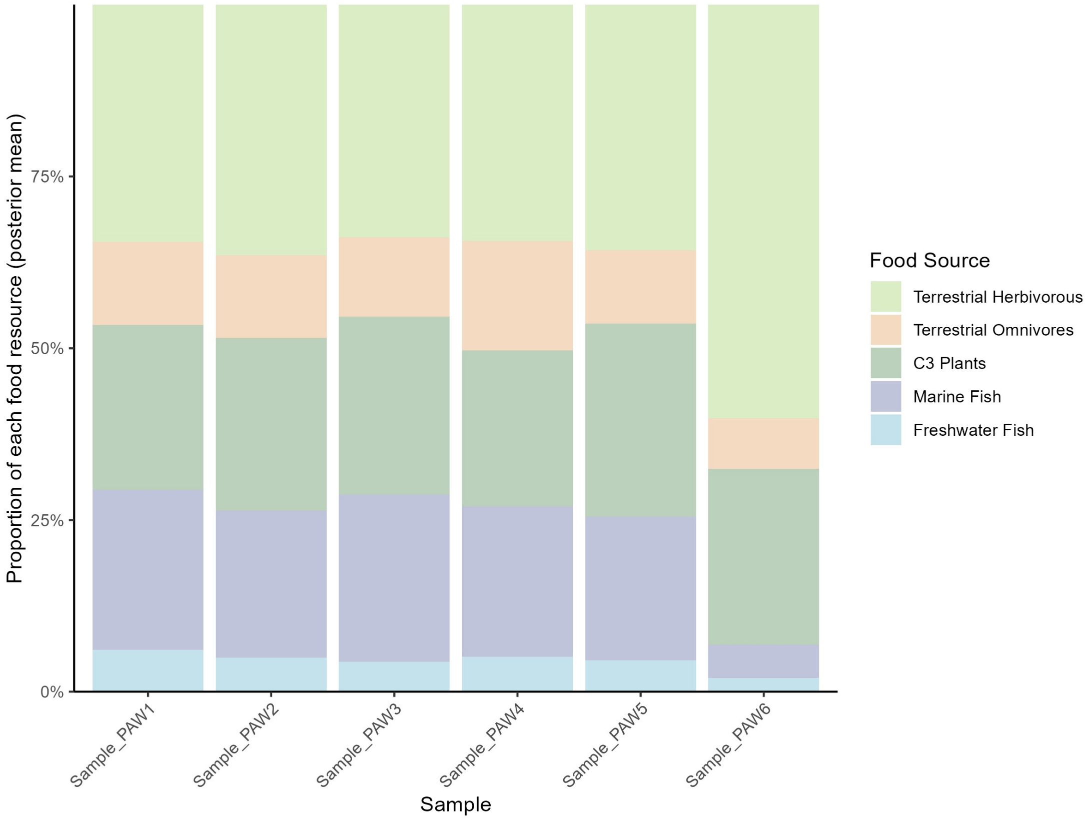

**Fig. S2| Dietary reconstruction for Paleolithic Wales human samples.**

#### 2B. Mesolithic period

##### England

The Mesolithic diet in England can be broadly divided into two categories. The first category[14], represented by individuals from Doggerland (Samples_MLE2–32), relied heavily on freshwater resources (Fig. S3). Among them, several individuals (Sample_MLE11, 15, 17, and 24) also show significant consumption of marine resources. The dietary reconstruction supports the observations from the scatter plot analysis in the original study[14]: humans from the southern North Sea region primarily followed a freshwater/terrestrial dietary regime, with a small number of individuals exhibiting substantial reliance on marine resources.

The other group[21–26] relied heavily on terrestrial resources (Sample_MLE1, and Samples_MLE33–73) (Fig. S3). Within this group, terrestrial herbivores constituted the primary food source, supplemented by smaller amounts of omnivores and plants. The contribution of marine and freshwater fish to their diet was nearly negligible. As noted in previous studies, Aveline's Hole is located far from the contemporary coastline, and therefore, the individuals from this site (Samples_MLE33–53)[21] are not expected to have consumed any marine resources. Similarly, samples from Badger Hole, Cannington Park Quarry, Greylake, Totty Pot, Hazleton, and West Kennet (Samples_MLE54 – 60, Samples_MLE62 – 73)[21,23,25,26], which may also have been located far from the coastline in Mesolithic England, exhibit the same dietary pattern as those from Aveline's Hole. The remaining two samples from Staythorpe Power Station and Tilbury (Sample_MLE1 and 61)[22,23] also exhibit the same subsistence strategy.

**Wales**

Samples from Wales exhibit significantly different dietary patterns (Fig. S4). A large number of individuals from Foxhole Cave, Paviland, and Caldey Island (sites including Daylight Rock, Ogof-yr-Ychen, and Potter's Cave)[23,27,28] show strong signals of marine resource consumption (Samples_MLW2 – 5 and Samples_MLW20 – 28). This is expected, considering that these sites were likely situated close to the coastline. However, Sample_MLW1 (original ID: L2, FX97-41), also from Foxhole Cave, displays a terrestrial-based diet. The reason has yet to be investigated. In contrast, Samples_MLW6–19[23,25], from Mewslade Bay, Worm's Head, and Parc le Breos Cwm, exhibit a terrestrial subsistence strategy. From an isotopic perspective, the $\delta^{13}C$ and $\delta^{15}N$ values of these samples are both lower than those of individuals with marine-based diets (see Supplementary Data 3). Although these individuals were not far from the coast, their diets exhibited predominantly terrestrial patterns, which is somewhat unexpected.

Overall, the Mesolithic dietary patterns reflect a hunter-gatherer subsistence strategy, with populations primarily relying on resources readily available in their local environments. Nevertheless, some individuals from Wales present unexpected dietary signals that warrant further investigation.

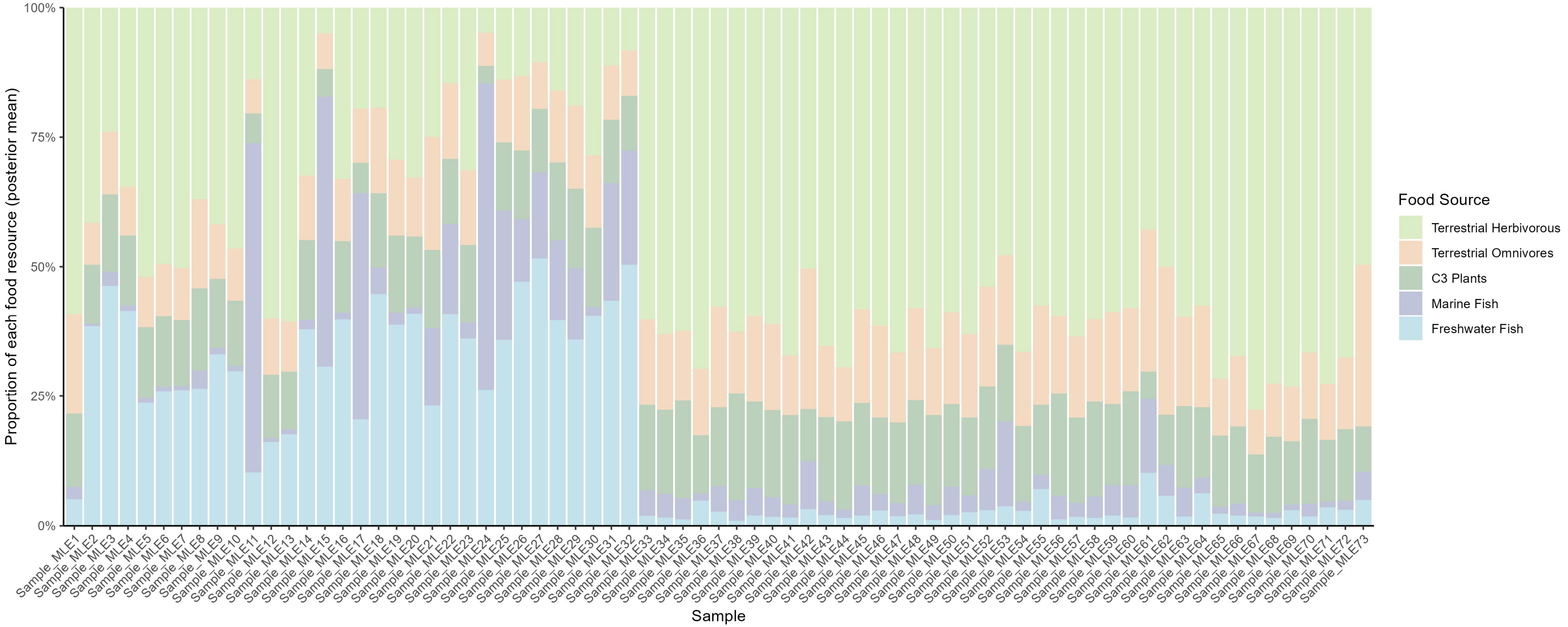

**Fig. S3| Dietary reconstruction for Mesolithic England human samples.**

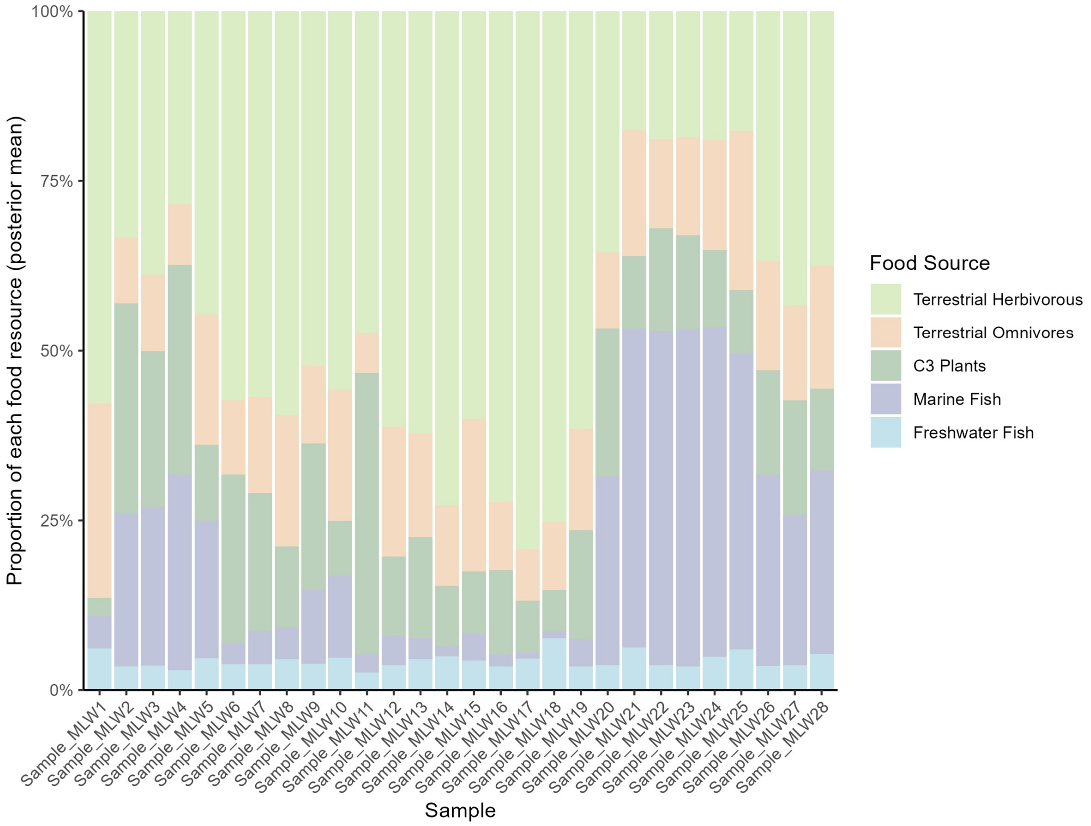

**Fig. S4| Dietary reconstruction for Mesolithic Wales human samples.**

#### 2C. Neolithic period

##### Scotland

With the introduction of agriculture, almost all individuals shifted their diets toward terrestrial resources (Samples_NLS1 – 317)[29–34], primarily based on $C_3$ plants and animal husbandry (Fig. S5). This pattern holds true even for samples from sites located near the coastline. Although the consumption of marine resources varies among individuals, it remains much lower than the proportions observed in the Mesolithic samples from Wales. During the Neolithic transition period, marine resources appear to have been considered a supplementary component of the diet, possibly exploited primarily during periods of famine due to environmental stress[31].

For Samples_NLS318–334 (all from Oronsay)[29,35,36], the reconstruction results indicate a marine-based dietary pattern (Fig. S5). These samples are dated significantly earlier than the others (from 4300 BC to 3600 BC), suggesting that they belong to a transitional period between the Mesolithic and Neolithic. Samples_NLS329 – 334 are even classified as Mesolithic in their original publications. This demonstrates that reliance on marine protein along the west coast of Scotland persisted well beyond the Late Mesolithic period, extending into the 4th millennium BC and the early stages of the Neolithic. This may suggest that Mesolithic subsistence practices persisted on these islands even after the Neolithic had arrived on the mainland.

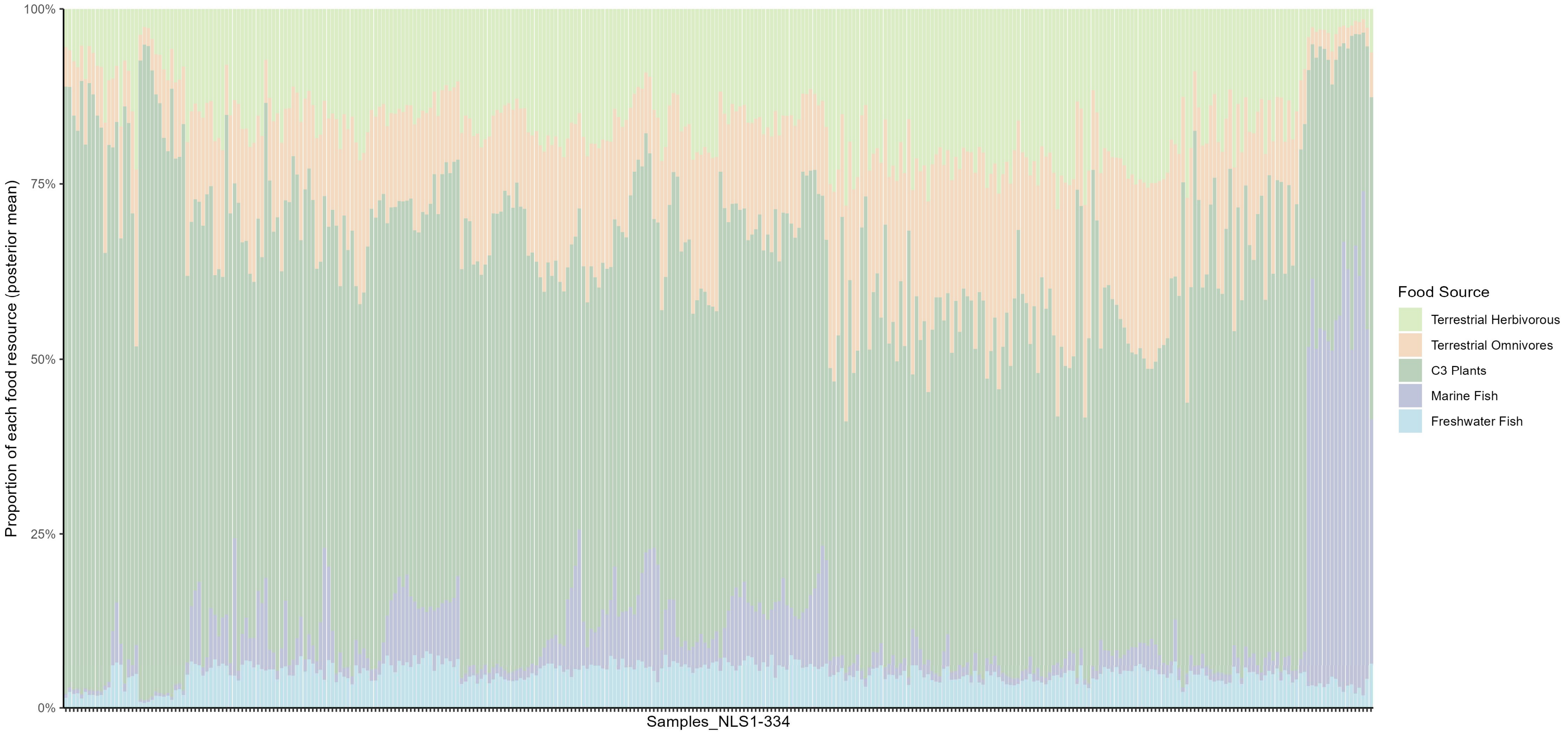

**Fig. S5| Dietary reconstruction for Neolithic Scotland human samples.**

**England and Wales**

Samples from Neolithic England[24,37–41] and Wales[28] do not exhibit any particularly distinctive patterns (Fig. S6, Fig. S7). They reflect a typical agricultural diet, based primarily on $C_3$ ecosystem, with very limited contributions from marine or freshwater sources. Their diet shows no significant difference compared to that of Neolithic Scotland, except for individuals from the Mesolithic–Neolithic transition period, who exhibit high levels of marine protein consumption.

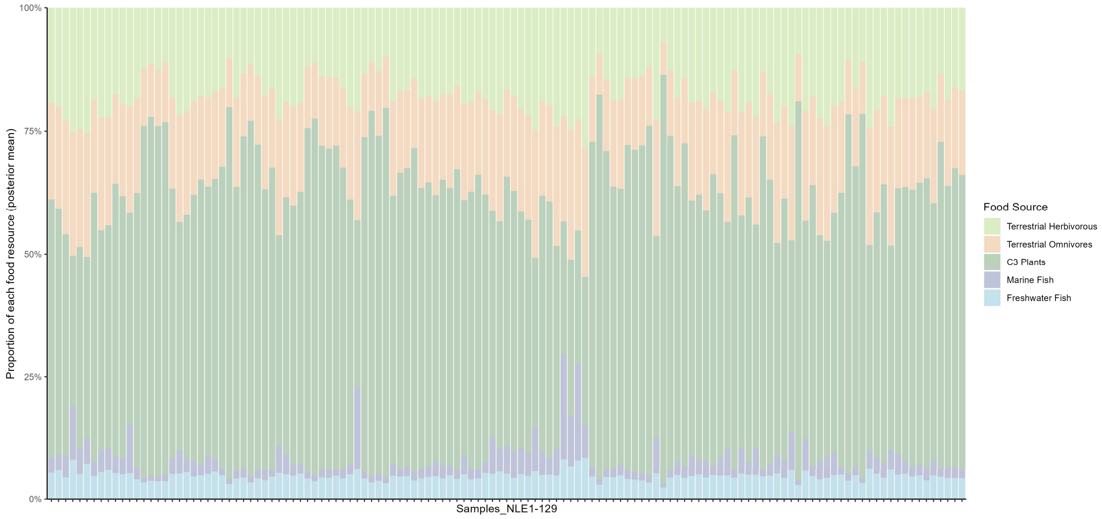

**Fig. S6| Dietary reconstruction for Neolithic England human samples.**

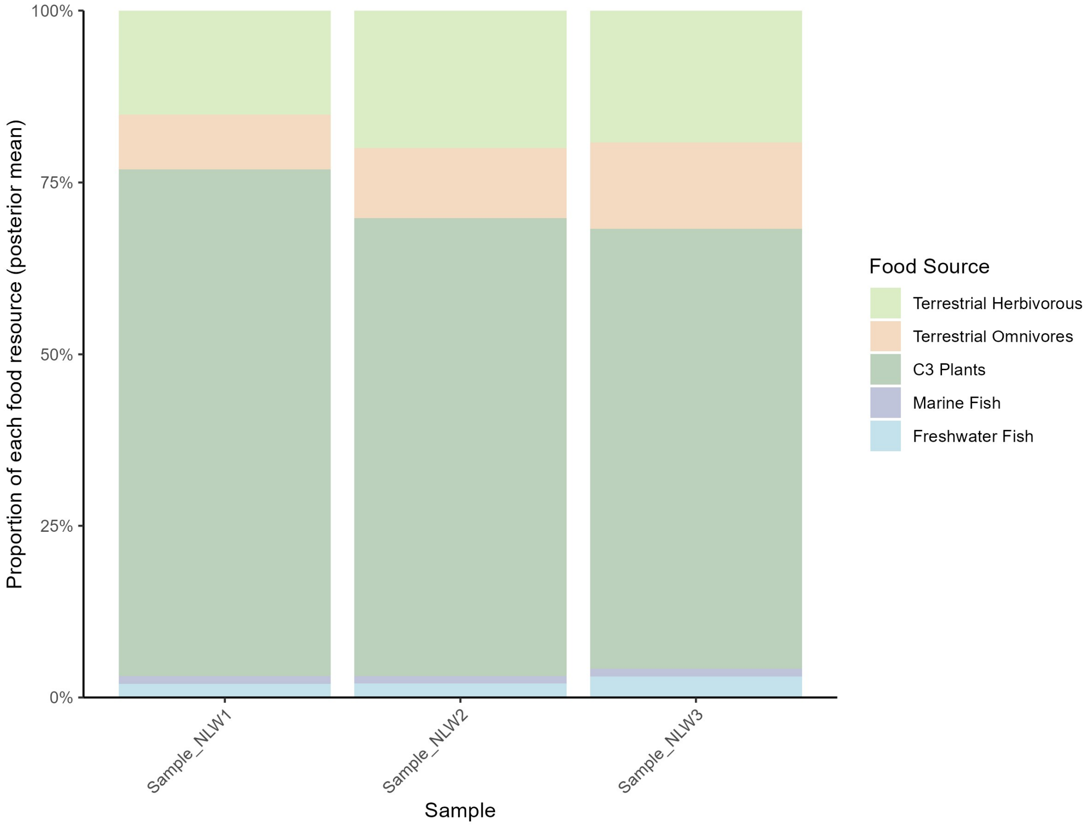

**Fig. S7| Dietary reconstruction for Neolithic Wales human samples.**

Some samples from England were identified by the Bayesian mixing model as having slightly higher marine resource consumption (10% to 20%) (Sample_NLE4, 44, 73, 74, and 75) (Fig. S6). These samples are from Eton College Rowing Course (Sample_NLE4: 2505–2415 BC), Hazleton North (Sample_NLE44: 3780–3640 BC), and Totty Pot (Samples_NLE73–75: 3355–2930 BC). The individuals showing higher levels of marine food consumption do not appear to be geographically or temporally exceptional. Their $\delta^{13}C$ values suggest that they consumed only very limited amounts of marine fish, and in their original studies, their relatively high $\delta^{15}N$ values were interpreted as resulting from increased consumption of terrestrial animal protein[37,38,40]. However, the Bayesian estimation here yields different results. Although we set a very low prior for marine resource consumption, the model still produced relatively high posterior proportions. This discrepancy can be regarded as a conflict between the mathematical model and the descriptive scatter plot interpretation (traditional scatter plot analysis may lack precision because it depends largely on subjective assessment of data distributions without the support of rigorous mathematical modeling). Nevertheless, since only 4% (5/129) of individuals exhibit such signals, this difference is unlikely to affect our overall time-series modeling results.

It is evident that the introduction of agriculture around 6000 years ago led to a relatively homogeneous dietary pattern across Britain, in contrast to the more diverse diets of the Paleolithic and Mesolithic periods. Except for some transitional period samples from the western coast of Scotland, all individuals exhibit a terrestrial-dominated diet, although their levels of animal protein consumption vary (i.e., some individuals consumed more, while others consumed less).

#### 2D. Bronze Age

##### Scotland and England

Similar to Neolithic period samples, Bronze Age individuals (1950 BC to 840 BC) from Scotland[32,42,43] primarily consumed terrestrial resources (Fig. S8). Notably, all of these samples come from coastal areas (Sample_BAS1 and 2: Sculptor's Cave; Sample_BAS3: Isbister; Sample_BAS4: Stenchme; Samples_BAS5–9: Cladh Hallan). Despite their proximity to the sea, only a few individuals (Sample_BAS3, 4, and 9) show slightly elevated marine resource consumption, and even these levels remain significantly lower than those observed in Mesolithic individuals from Wales. The remaining Scottish samples show little to no evidence of marine food consumption.

Samples from Bronze Age England exhibit a similarly terrestrial-based diet (Samples_BAE1–6[37]: 1060–780 BC; Samples_BAE7–9[44]: 2284–1900 BC), with even lower levels of marine resource consumption compared to those from Bronze Age Scotland (Fig. S9). All individuals were recovered from Yarnton, a site located far inland, making their minimal reliance on marine foods unsurprising.

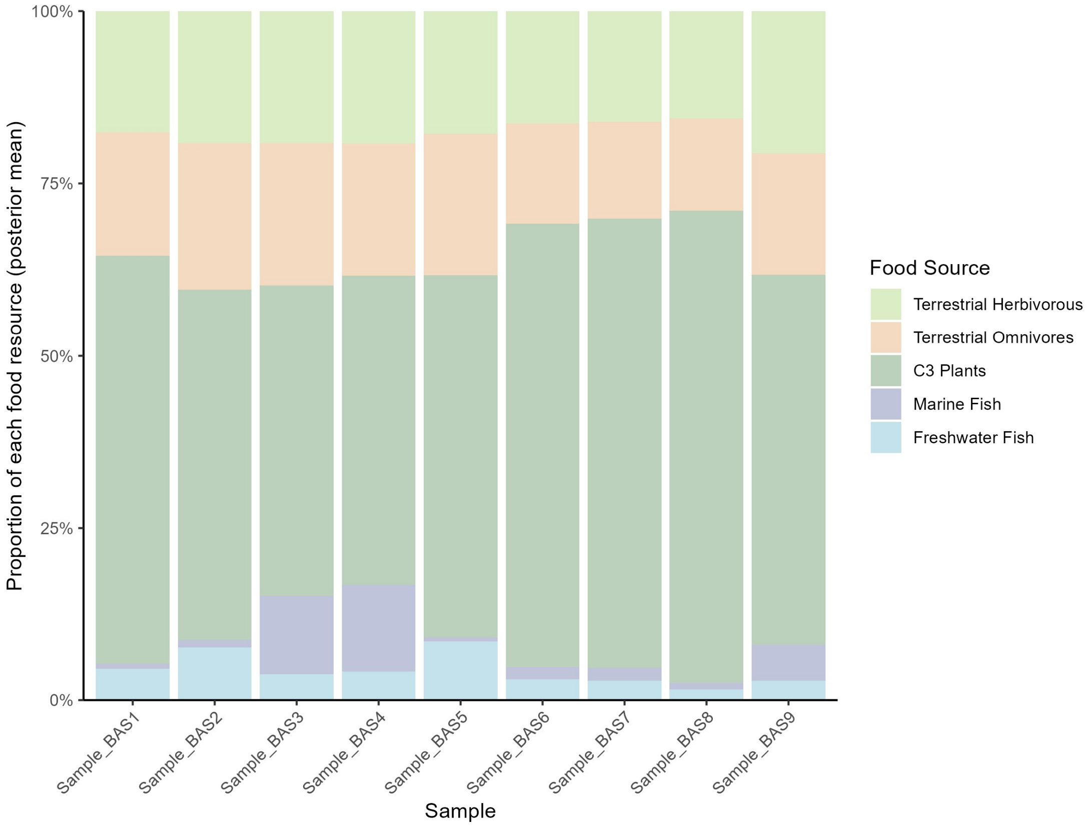

**Fig. S8| Dietary reconstruction for Bronze Age Scotland human samples.**

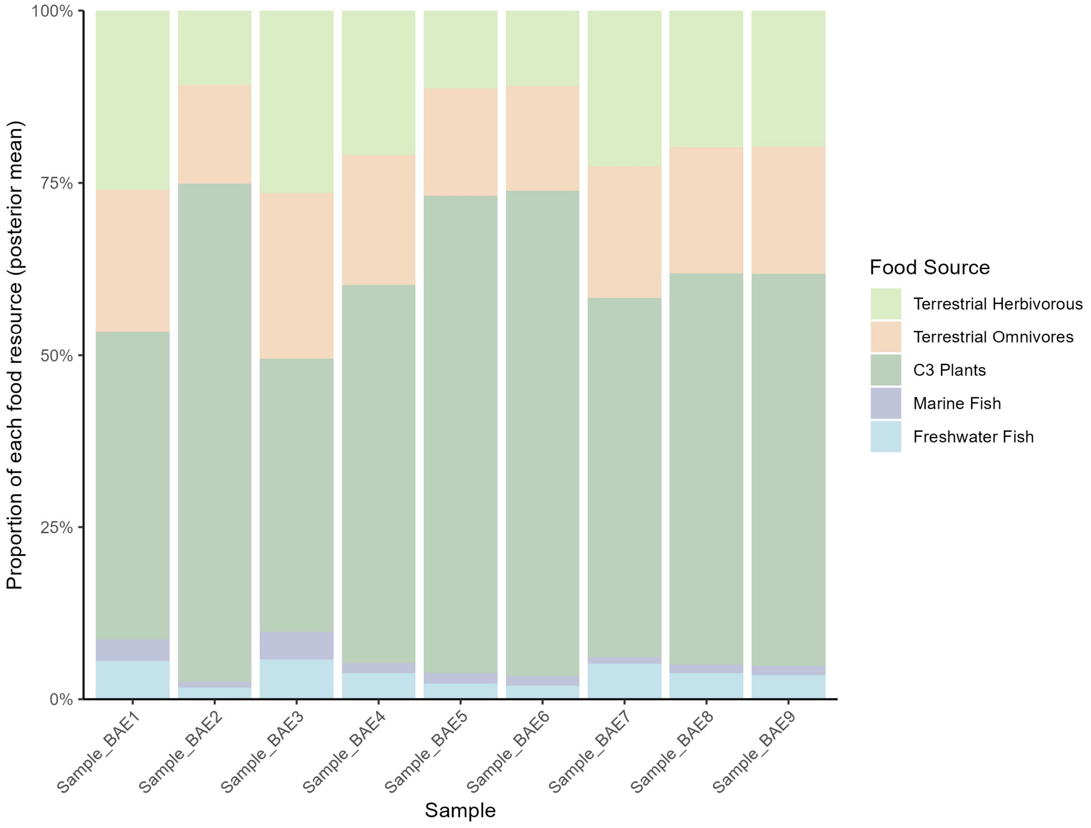

**Fig. S9| Dietary reconstruction for Bronze Age England human samples.**

**The samples from *the Beaker People Project and the Beakers and Bodies Project* (535 samples from 2500 BC to 1500 BC; eight samples from 1500 BC to 750 BC)**

The samples[45,46] dated between 2500 BC and 1500 BC also exhibit a fully terrestrial diet, with virtually no contribution from freshwater or marine resources, regardless of whether they are bone (Fig. S10, Fig. S11, Fig. S12, Fig. S13) or dentine samples (Fig. S14, Fig. S15, Fig. S16, Fig. S17). The only variation between samples lies in the relative intake of terrestrial animal protein: some individuals consumed more animal-derived protein, while others relied more heavily on $C_3$ plants. Diets in Scotland appear to be more homogeneous (Fig. S11, Fig. S15), whereas in England, the consumption of animal protein varies more substantially between individuals (Fig. S10, Fig. S12, Fig. S14, Fig. S16). The remaining eight samples from the Late Bronze Age also exhibit a similar terrestrial diet (Fig. S18).

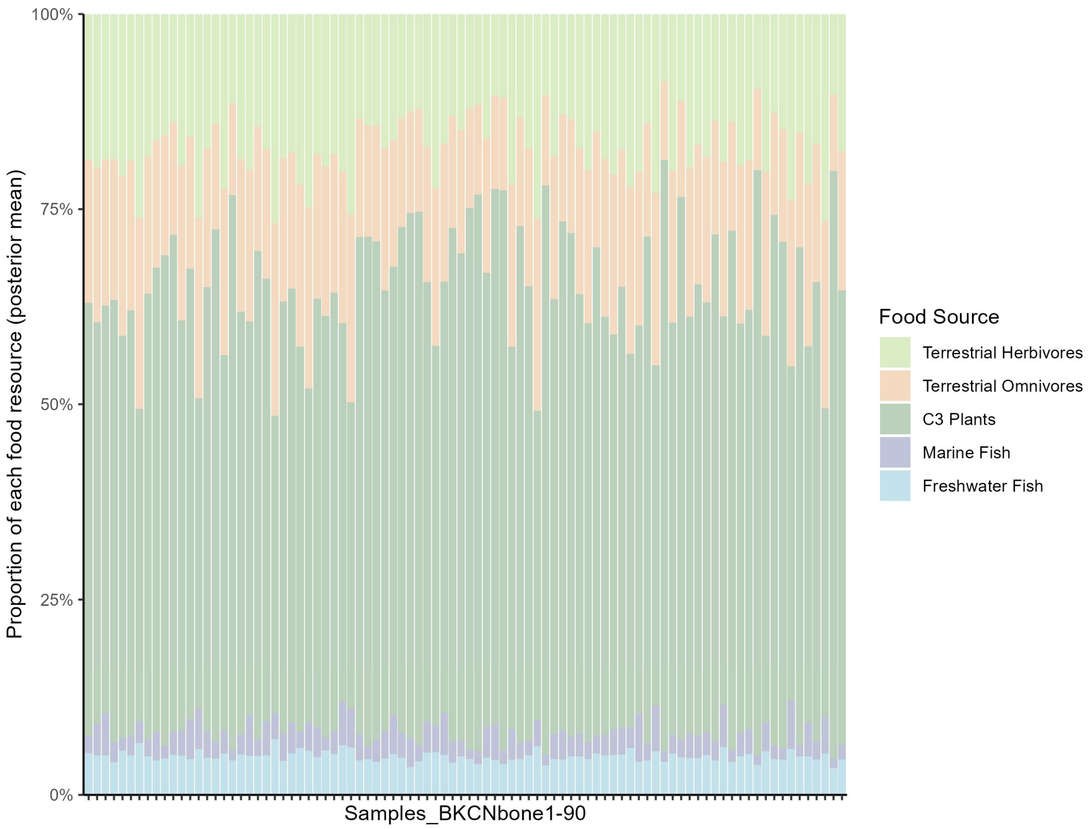

**Fig. S10| Dietary reconstruction for Beaker-period human samples from central and northern England (bone collagen).**

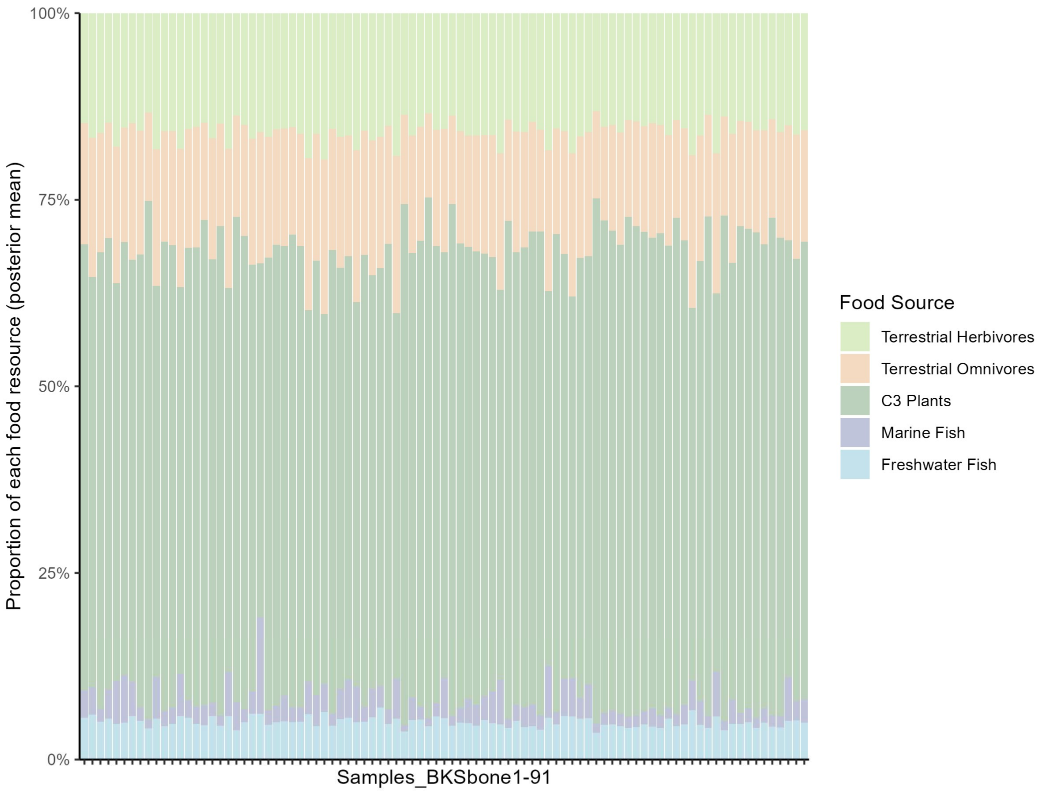

**Fig. S11| Dietary reconstruction for Beaker-period human samples from Scotland (bone collagen).**

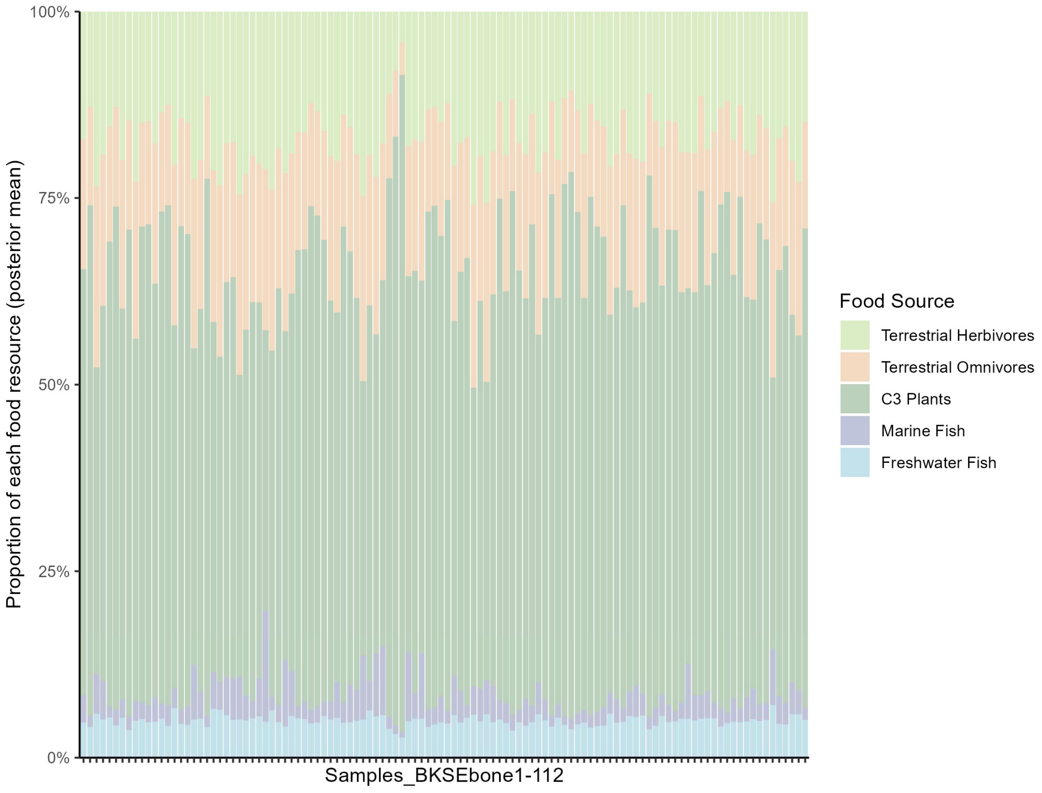

**Fig. S12| Dietary reconstruction for Beaker-period human samples from southern England (bone collagen).**

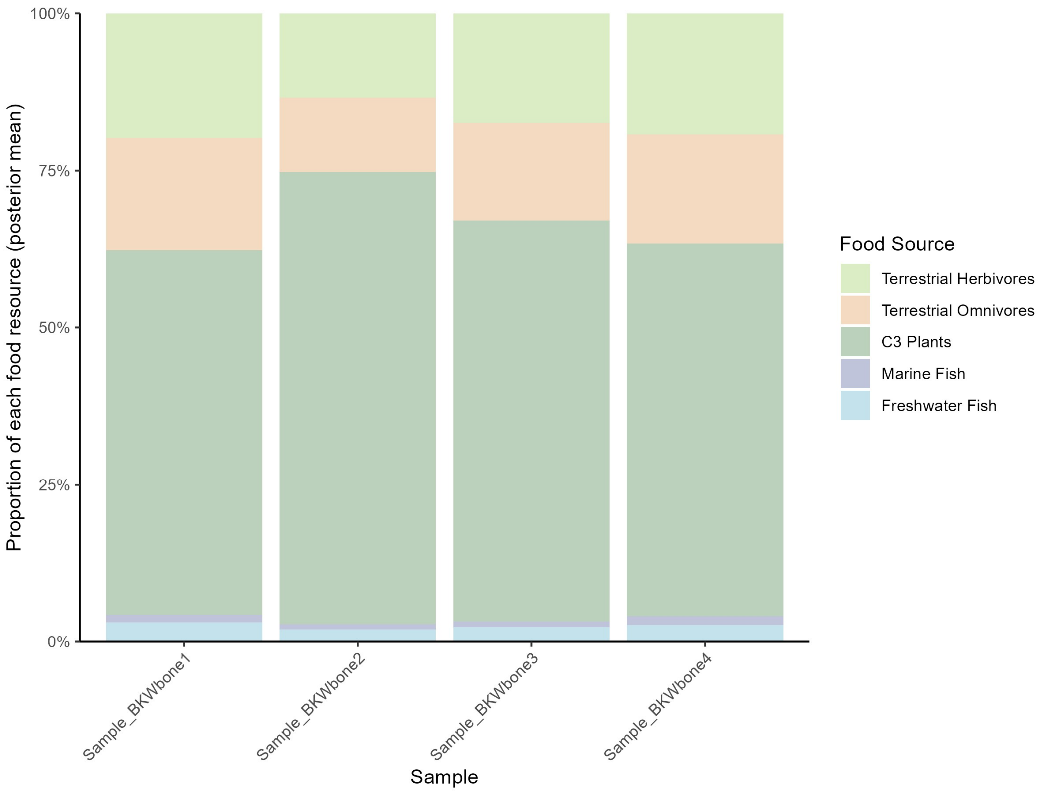

**Fig. S13| Dietary reconstruction for Beaker-period human samples from Wales (bone collagen).**

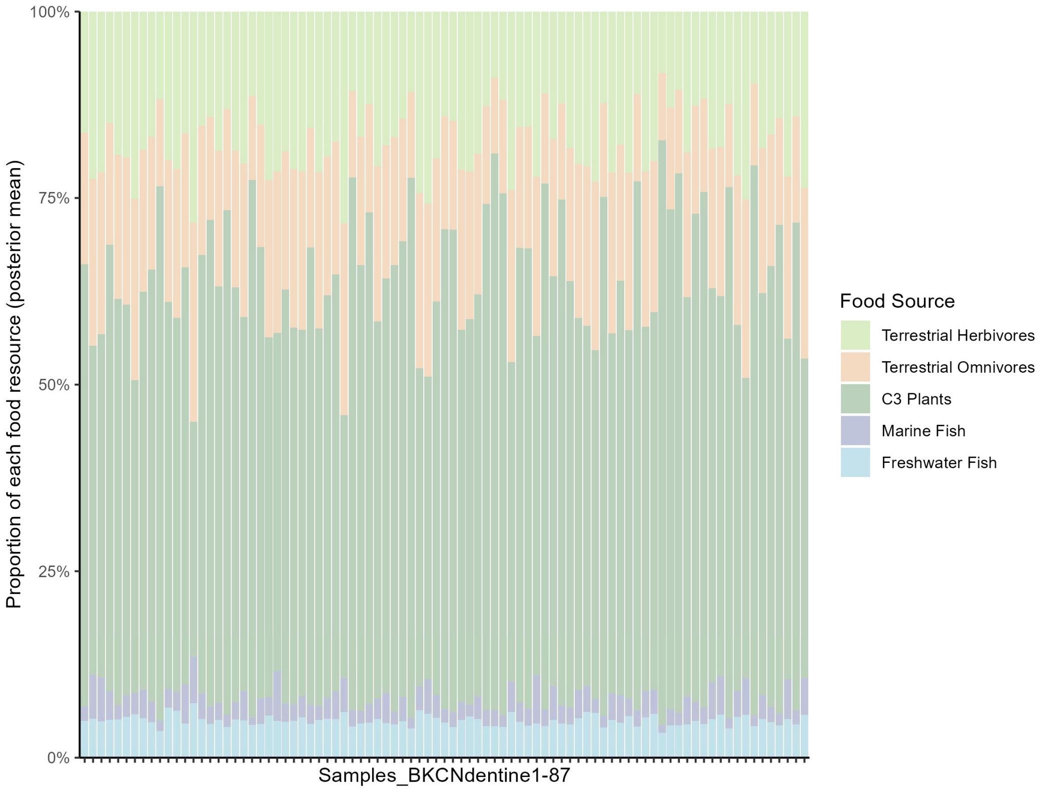

**Fig. S14| Dietary reconstruction for Beaker-period human samples from central and northern England (dentine).**

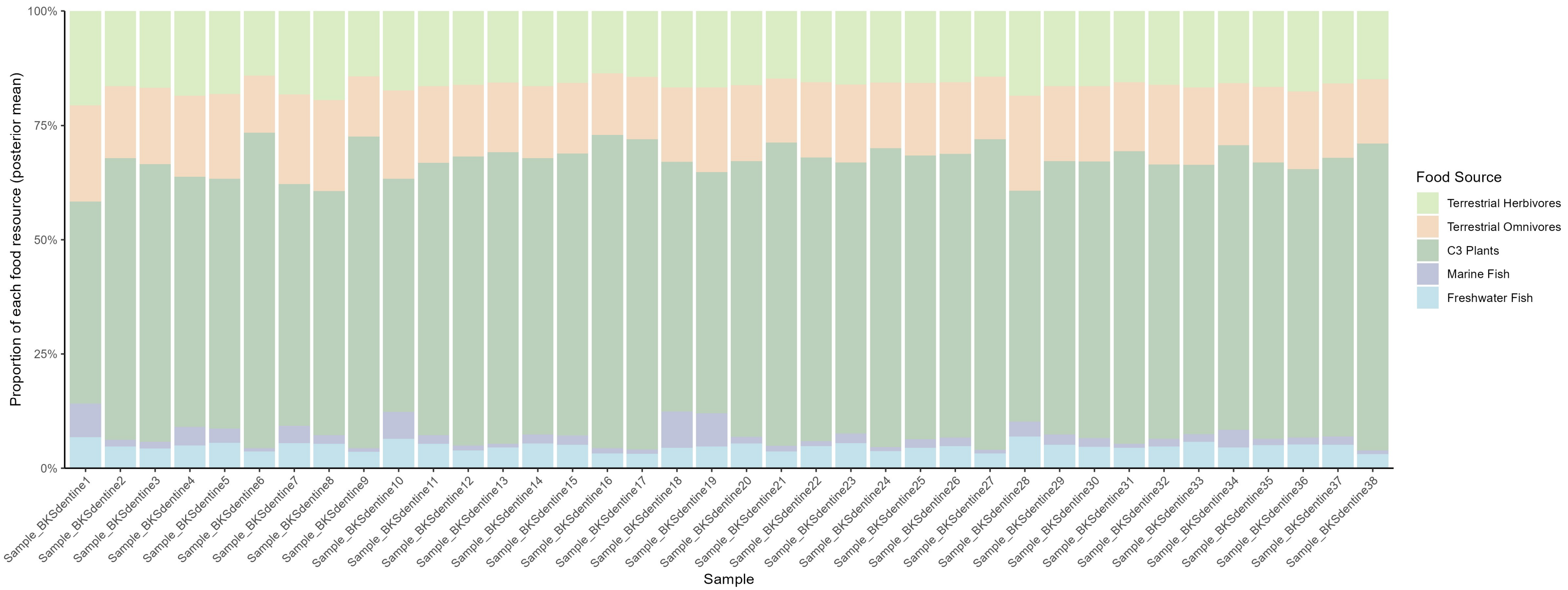

**Fig. S15| Dietary reconstruction for Beaker-period human samples from Scotland (dentine).**

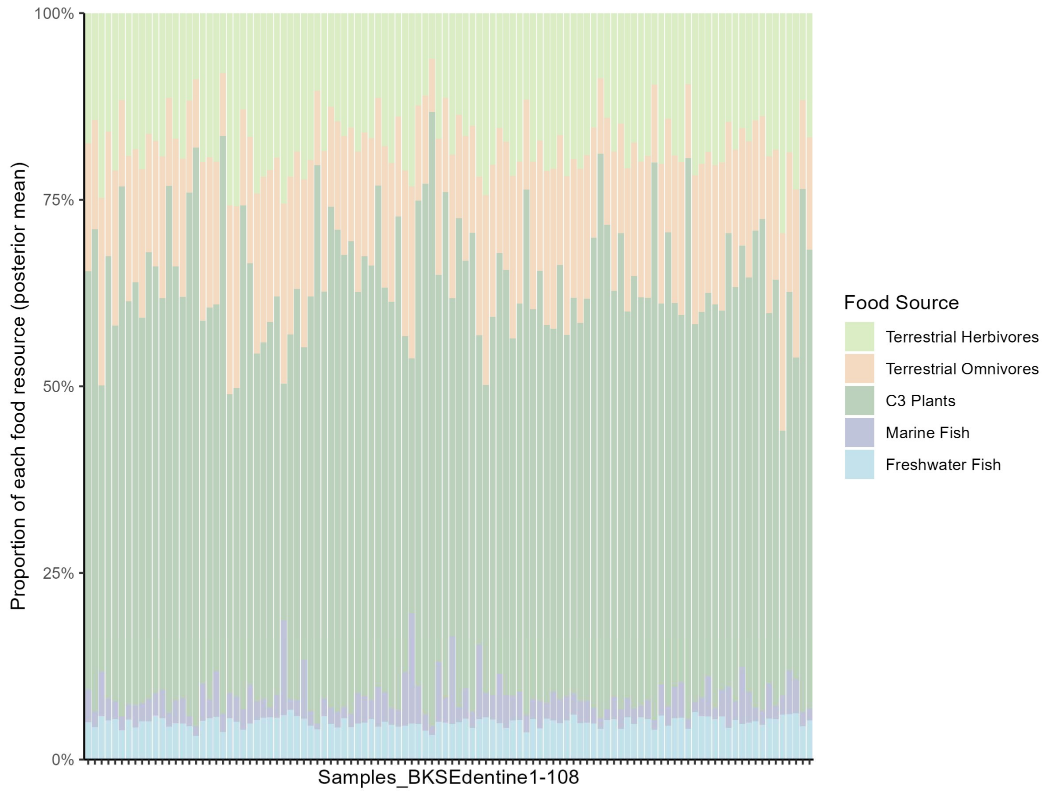

**Fig. S16| Dietary reconstruction for Beaker-period human samples from southern England (dentine).**

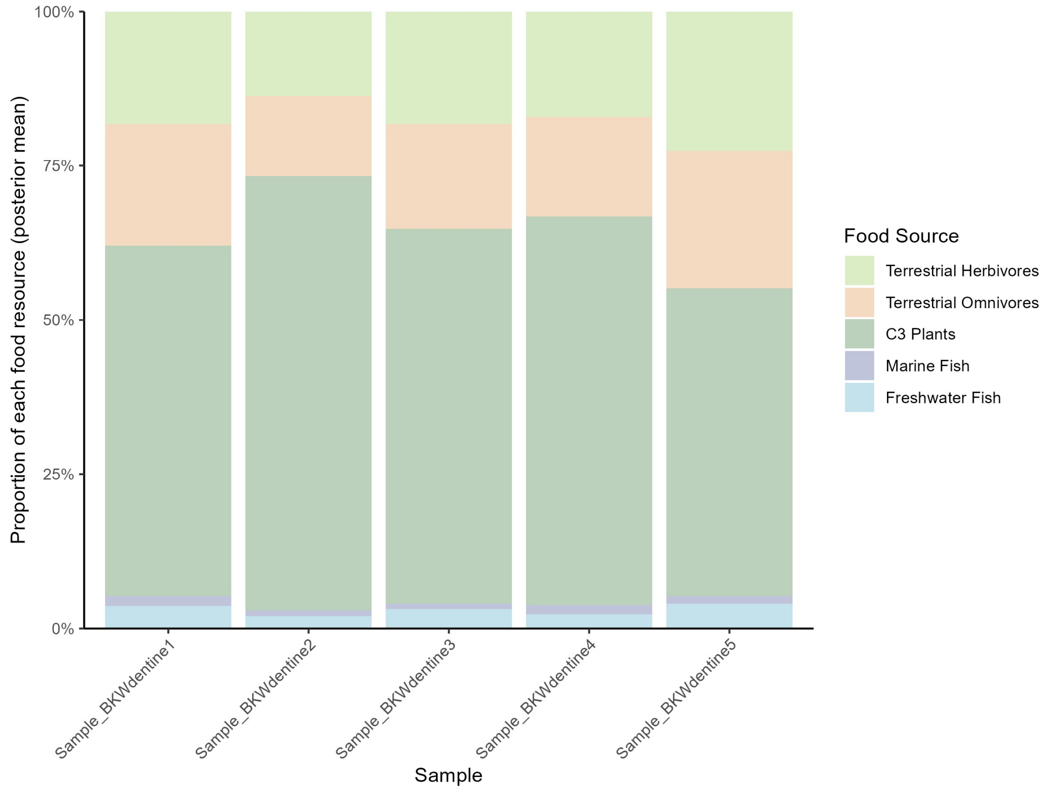

**Fig. S17| Dietary reconstruction for Beaker-period human samples from Wales (dentine).**

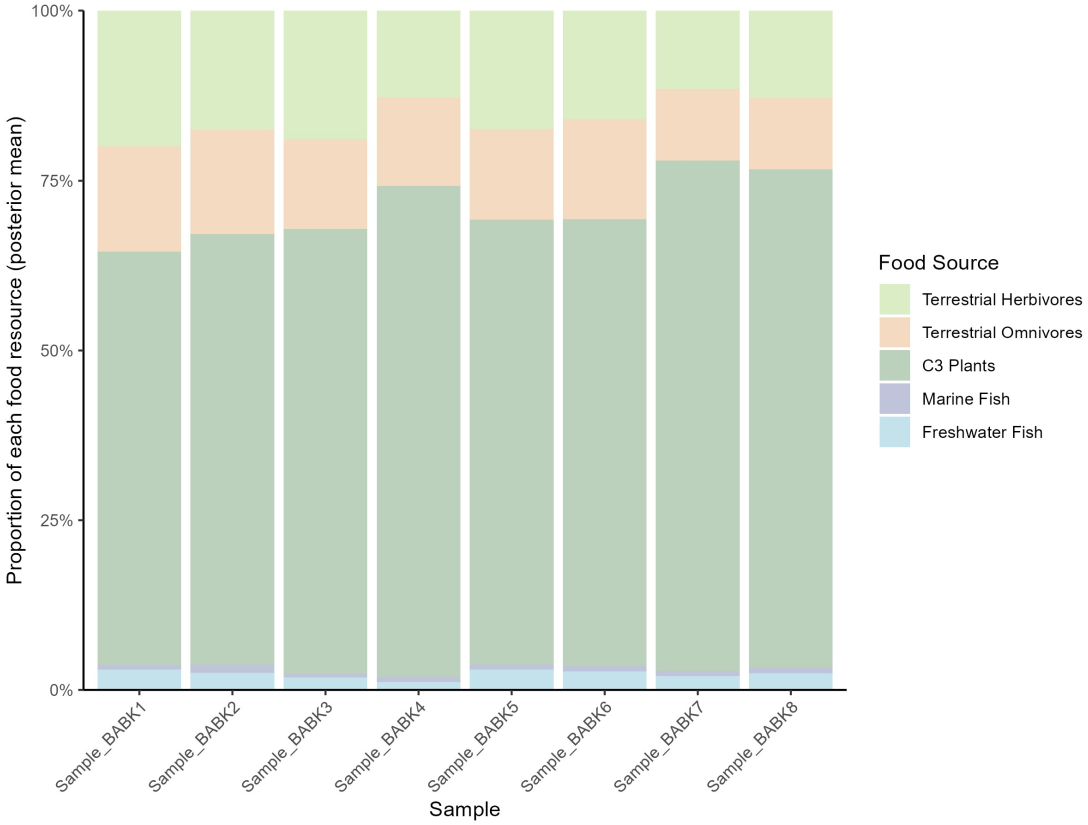

**Fig. S18| Dietary reconstruction for Bronze Age human samples from the Beaker People Project (1500 BC to 750 BC).**

There is evidence suggesting that, starting around 3300 BC, agriculture in Britain began to decline, with most people adopting a more pastoral lifestyle[47–49]. This trend eased somewhat following the Beaker period, but agriculture did not fully recover until around 1500 BC.

Our dietary data partially support this interpretation (Fig. S19). On average, there was an increasing reliance on terrestrial animal protein and a reduction in the consumption of $C_3$ plants from 3300 to 2500 BC, although this dietary shift was far less pronounced than the transition from the Mesolithic to the Neolithic. Subsequently (after 2500 BC), plant consumption increased to levels that were the same as, or even exceeded, those observed in the early Neolithic (before 3300 BC). This contradicts the view that agriculture did not fully recover until around 1500 BC.

Although the proportions of animal protein consumption exhibit patterns indicative of a more pastoral lifestyle, the modest magnitude of these changes appears to support a different hypothesis: that the disruption of agriculture was geographically localized[50]. This spatial heterogeneity likely led to increased animal consumption in certain regions, but at the scale of Britain, the overall dietary transition appears subtle (Fig. S19).

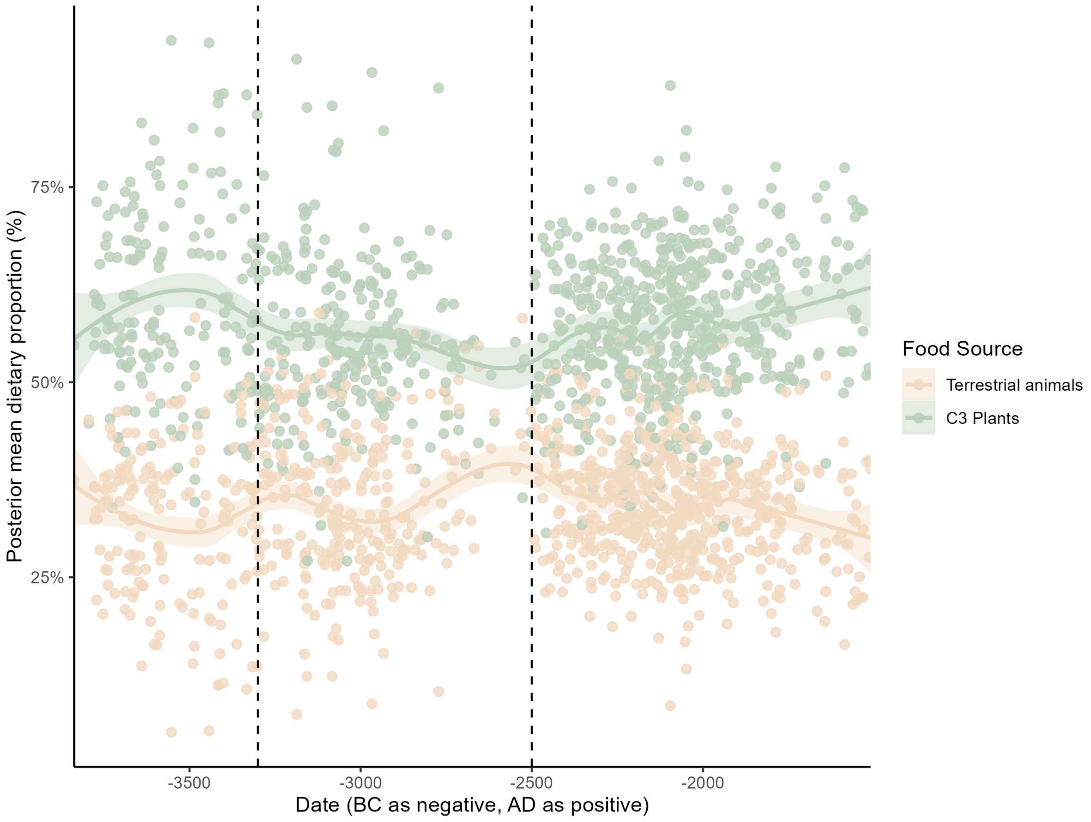

**Fig. S19| Comparative analysis of Neolithic and Bronze Age diets.** Dots represent the posterior mean dietary proportions of individuals. Lines represent smoothed dietary trends over time based on LOESS regression, while shaded areas indicate 95% confidence intervals. 'Terrestrial animals' refers to the combined proportion of herbivores and omnivores.

#### 2E. Iron Age

##### Scotland

Iron Age individuals from Scotland[32,51,52] also exhibit a typical agricultural diet, characterized by strong terrestrial $C_3$ signals and minimal consumption of marine and freshwater resources (Fig. S20). A small number of individuals show evidence of aquatic resource use. Sample_IAS1, which displays slightly elevated $\delta^{13}C$ values (-19.9‰), suggests slightly higher marine input. Two individuals (Sample_IAS32 and 33) exhibit higher freshwater resource consumption. Both were recovered from the Musselburgh site, which is located near the River Esk, making it plausible that the inhabitants had access to freshwater resources.

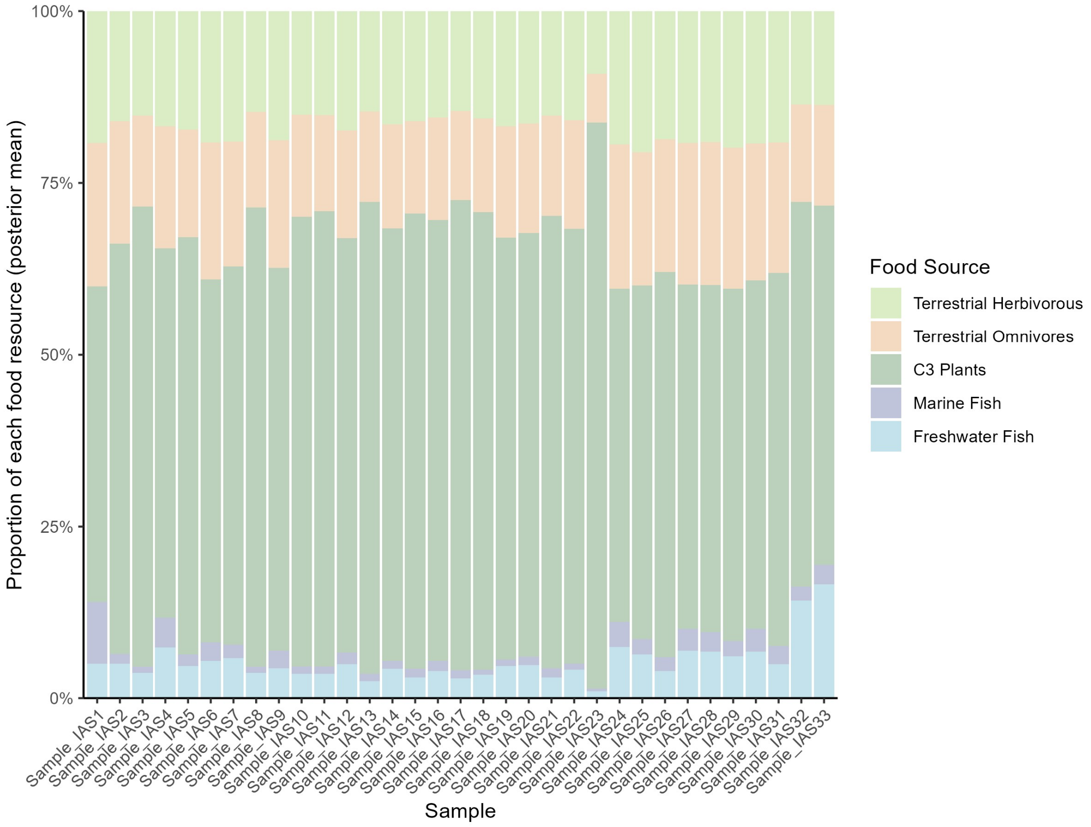

**Fig. S20| Dietary reconstruction for Iron Age Scotland human samples.**

#### England

The northern England individuals[53] are all from Wetwang Slack in East Yorkshire, and their dietary information is generally consistent with the previously described terrestrial pattern (Fig. S21). Two outliers (Sample_IANE4 and 54) appear to have consumed more aquatic resources. These individuals also exhibit isotopic values that deviate from the rest of the group, particularly elevated $\delta^{15}N$ values. The original study suggested that these individuals may have been mobile, originating from a different region where freshwater aquatic resources contributed more substantially to the diet. Given the limited local access to freshwater resources, it is unlikely that they were local inhabitants.

The individuals from southern England[37,44,51,54–59] are also primarily based on terrestrial resources, but they exhibit a more diverse dietary pattern overall (Fig. S22). Individuals from Danebury Ring Hillfort and Suddern Farm (Samples_IASE1–79) display an almost entirely terrestrial diet (Fig. S22).

Samples_IASE80–107 from Yarnton[44] exhibit a more diverse dietary pattern and higher levels of animal protein consumption (Fig. S22). Notably, our modeling results indicate that several individuals from this region show substantial marine resource consumption, which appears to contradict existing conclusions. Previous studies have suggested that marine resources were rarely consumed during the Iron Age. In our priors, we assigned low marine contribution values to these individuals; however, the posterior estimates for six samples (Sample_IASE83, 88, 94, 95, 99, and 101) still indicate significant levels of marine intake, ranging from 10% to 24% (Fig. S22).

The original explanation for the elevated $\delta^{15}N$ values in this area was that animals consumed plants grown on soils fertilized with manure, resulting in higher $\delta^{15}N$ values in terrestrial food sources[44]. However, our comprehensive analysis of faunal isotopic data[(Chen et al in prep)] refutes this hypothesis: even in nearby regions such as Segsbury Camp and Alfred's Castle, animal isotopic values display entirely different patterns. We suggest that there is considerable variance in animal isotopic values within this region, and the original study may have sampled unusually high-$\delta^{15}N$ individuals, leading to a biased interpretation. This also highlights the importance of large-scale data analysis: relying solely on local faunal samples for comparison may lead to biases. Our modeling results suggest that the elevated $\delta^{15}N$ values in this region are primarily due to increased consumption of terrestrial animal protein, with a small number of individuals also showing relatively high levels of marine protein intake.

Our findings further demonstrate that subjective interpretations of isotopic data can sometimes be unreliable. The six samples in question have $\delta^{13}C$ values between -19.8‰ and -20.3‰. While traditional isotopic interpretation would classify them as having a purely terrestrial diet with minimal marine input, as assumed in our model prior settings, our posterior estimates reveal that this is not necessarily the case.

Two samples from Cornwall (Sample_IASE133 and 147)[51] and two from Alington Avenue (Sample_IASE186 and 188)[59] exhibit evidence of marine resource consumption (Fig. S22). Our modeling results support previous studies suggesting that a small number of individuals from Cornwall and Alington Avenue consumed slightly elevated proportions of marine resources.

Four samples from Hampshire (Sample_ IASE157, 161, 162, and 178)[51] and one from Poundbury Camp (Sample_IASE214)[59] exhibit high levels of freshwater resource consumption (Fig. S22). The Hampshire individuals were originally interpreted as non-locals, based on the evidence available at that time suggesting that freshwater resources in the region exhibited relatively high $\delta^{13}C$ values. However, our analysis of faunal isotopic data[(Chen et al in prep)] indicates that freshwater isotopic signatures display a wide range of variation. Even within the same region and period, substantial differences in freshwater $\delta^{13}C$ values can be observed. Therefore, it is plausible that these individuals genuinely consumed greater amounts of freshwater resources. Furthermore, even if they originated from other parts of Britain, the overall isotopic variation in Iron Age terrestrial resources across regions is relatively limited. As such, the elevated isotopic values observed in these individuals are more likely to reflect significant freshwater fish consumption rather than non-local origin. Sample_IASE214 from Poundbury Camp was not discussed in the original publication, but our analysis suggests that it reflects notable freshwater resource consumption.

Our modeling results for the Iron Age indicate that, while terrestrial resources remained dominant, dietary patterns became more diverse compared to the Bronze Age. A small number of individuals began to incorporate more marine and freshwater resources into their diets—a pattern observed during the Neolithic but largely absent in the Bronze Age—although the overall level of aquatic resource consumption remained limited.

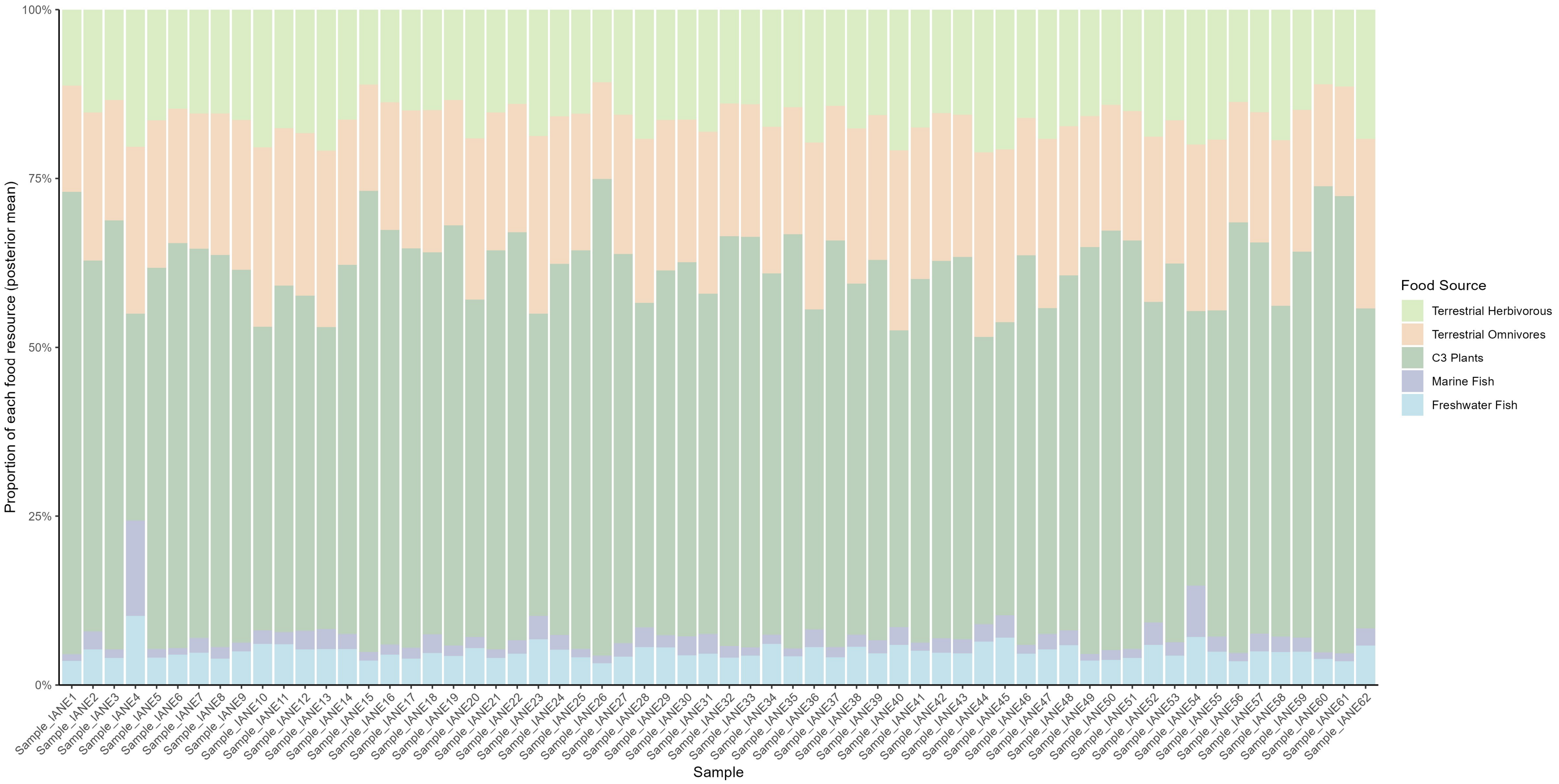

**Fig. S21| Dietary reconstruction for Iron Age northern England human samples.**

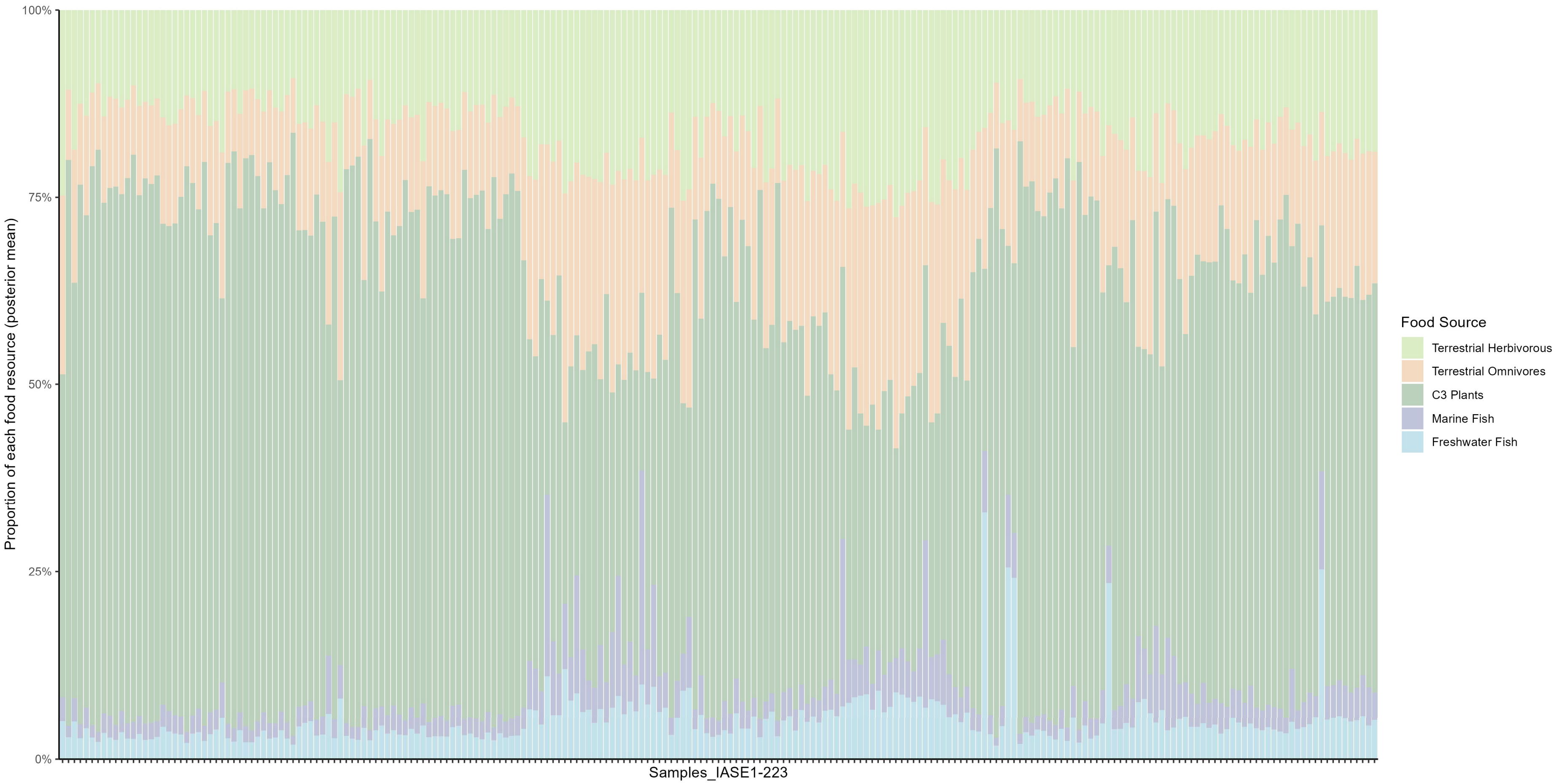

**Fig. S22| Dietary reconstruction for Iron Age southern England human samples.**

#### 2F. Roman period/ Roman Iron Age

##### Scotland

Only 17 individuals[42,52] from the Roman Iron Age in Scotland are available, originating from the Musselburgh site and Sculptor's Cave. Individuals from both sites followed a terrestrial $C_3$-based diet; however, the samples from Musselburgh exhibit a higher proportion of freshwater fish consumption (Samples_ROS1–10) (Fig. S23). Given the site's proximity to the River Esk, greater reliance on freshwater resources is plausible. This interpretation is further supported by archaeological evidence from Inveresk and the surrounding area.

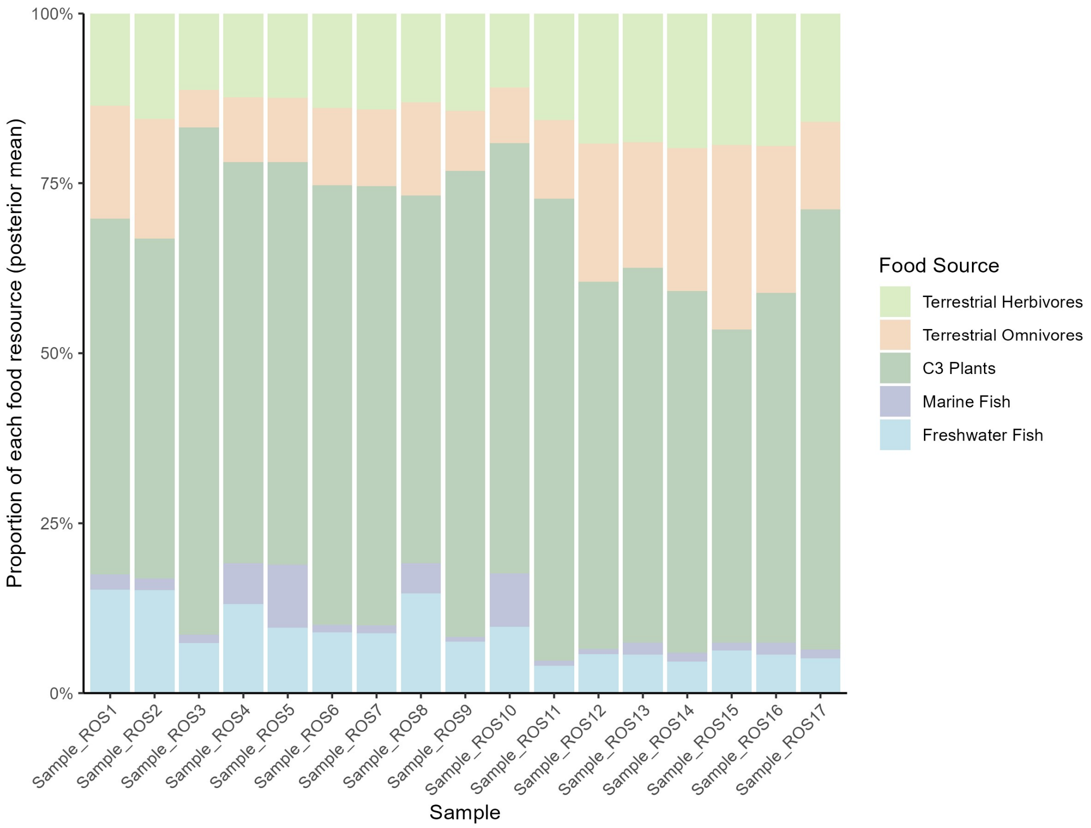

**Fig. S23| Dietary reconstruction for Roman Iron Age Scotland human samples.**

##### England

Individuals from northern England[60–64] are all from North Yorkshire. The most notable characteristic of this region is a more diverse diet, particularly with higher consumption of omnivores as well as marine and freshwater resources, compared to the Iron Age (Fig. S24). Individuals from Blossom Street (Samples_RONE1–17) and Trentholme Drive (Samples_RONE22 – 63)[60] in York are representative of this pattern. Our modeling results support the original interpretation that marine resources were a minor but regular component of the diet for many individuals (Fig. S24). Sample_RONE60 is an exception, exhibiting significant marine resource consumption. Its isotopic values are also notably high ($\delta^{13}C$: –17.1‰; $\delta^{15}N$: 14.1‰). It remains unclear whether this individual was non-local. Nonetheless, this case demonstrates that certain individuals in Roman-period Yorkshire had diets with a substantial marine component, even though such cases were rare overall. This dietary pattern—primarily based on terrestrial $C_3$ resources with a minor contribution from marine and freshwater sources, slightly higher than in the Iron Age—is confirmed by our dietary reconstructions at other sites in York (Samples_RONE122–244)[62,64] (Fig. S24). In addition to the individuals from York, those from a further north at Catterick[61,63] (Samples_RONE18 – 21 and Samples_RONE64–121) also exhibit the same dietary pattern (Fig. S24). However, dietary variation among individuals in Catterick appears to be more limited, with even lower levels of marine and freshwater resource consumption compared to York. This may be attributed to differences in food accessibility in more remote areas.

Individuals from Roman-period southern England[37,44,54,56,57,59,65–74] generally follow the same dietary pattern: a predominantly terrestrial-based diet, with certain populations exhibiting increased consumption of marine or freshwater resources (Fig. S25). Dietary heterogeneity appears to be greater than in earlier periods. This suggests that dietary variation linked to social status was widespread, especially in major cities. During the Roman period, marine resources were generally considered luxury items, more readily accessible to individuals of higher social status[67,72]. Sites showing higher levels of marine resource consumption include Alington Avenue, Poundbury Camp, several locations in Dorset, and London. Other sites—such as Queenford Farm, Gloucester, Hyde Close, Chester Road, Albert Road, and Old Vicarage—also contain a small number of individuals with elevated marine resource intake.

Our model indicates that several individuals from Yarnton[44] (Sample_ROSE47, 48, 50, 51, and 53) exhibited high levels of freshwater resource consumption (Fig. S25). The original study interpreted these individuals as primarily relying on terrestrial resources, but our findings challenge that conclusion. The $\delta^{13}C$ and $\delta^{15}N$ values for these individuals range from –19.5‰ to –19.9‰ and from 11.2‰ to 14.0‰, respectively. The elevated $\delta^{15}N$ values observed in the Yarnton may be attributed to manuring practices. However, based on our isotopic analysis of local food resources[(Chen et al in prep)], the $\delta^{15}N$ values of Roman-period herbivores (6.03‰) and omnivores (8.57‰) from this area are not statistically different from those observed in southern England (herbivores: 7.1‰; omnivores: 8.11‰). This slight enrichment in the $\delta^{15}N$ values of food resources is insufficient to explain the substantial isotopic elevation observed in human remains. Given Yarnton's close proximity to a river, such isotopic signatures are more plausibly explained by increased freshwater resource intake.

Other sites with evidence of high freshwater resource consumption include those from London and Gloucester. Aside from individuals with high levels of marine resource consumption, freshwater resources appear to have been consumed in slightly greater quantities than marine fish among the rest of the population—although neither resource constituted a major component of the diet (Fig. S25).

One notable pattern is observed: most southern England samples from the Roman period exhibit relatively low $\delta^{15}N$ values, while some of them simultaneously show elevated $\delta^{13}C$ values—an unusual combination.

Faunal isotopic baselines for terrestrial animals in this region during this period are similar to those in North Yorkshire[(Chen et al in prep)], yet human $\delta^{15}N$ values are significantly lower. If the high $\delta^{13}C$ values in southern England individuals were interpreted as indicating increased marine resource consumption, their $\delta^{15}N$ values would not support such an interpretation. This isotopic signature has also been observed in certain Neolithic samples; however, during the Neolithic, $\delta^{13}C$ and $\delta^{15}N$ values of terrestrial fauna from southern England were lower[(Chen et al in prep)], making a slight increase in marine input a more plausible explanation.

Some studies have incorporated additional strontium and oxygen isotope analyses, which suggest that individuals with this isotopic pattern may have spent their childhoods in warmer parts of the European continent, or consumed more $C_4$-based diet[70]. For such cases, we excluded the individuals from our modeling as probable immigrants. However, for many individuals exhibiting this pattern, there is no clear evidence of non-local origin. In these cases, we followed the interpretation proposed in the original studies[72,73]—that increased $\delta^{13}C$ reflects greater marine resource consumption. Accordingly, we incorporated this into our priors, and our modeling results indicate that these individuals likely increased their intake of marine resources while simultaneously reducing their consumption of terrestrial animal protein.

Overall, the Roman-period diet appears to have been more diverse than in earlier periods. While it remained largely based on a terrestrial $C_3$ ecosystem, the proportion of marine and freshwater resource intake increased—particularly among individuals of higher social status.

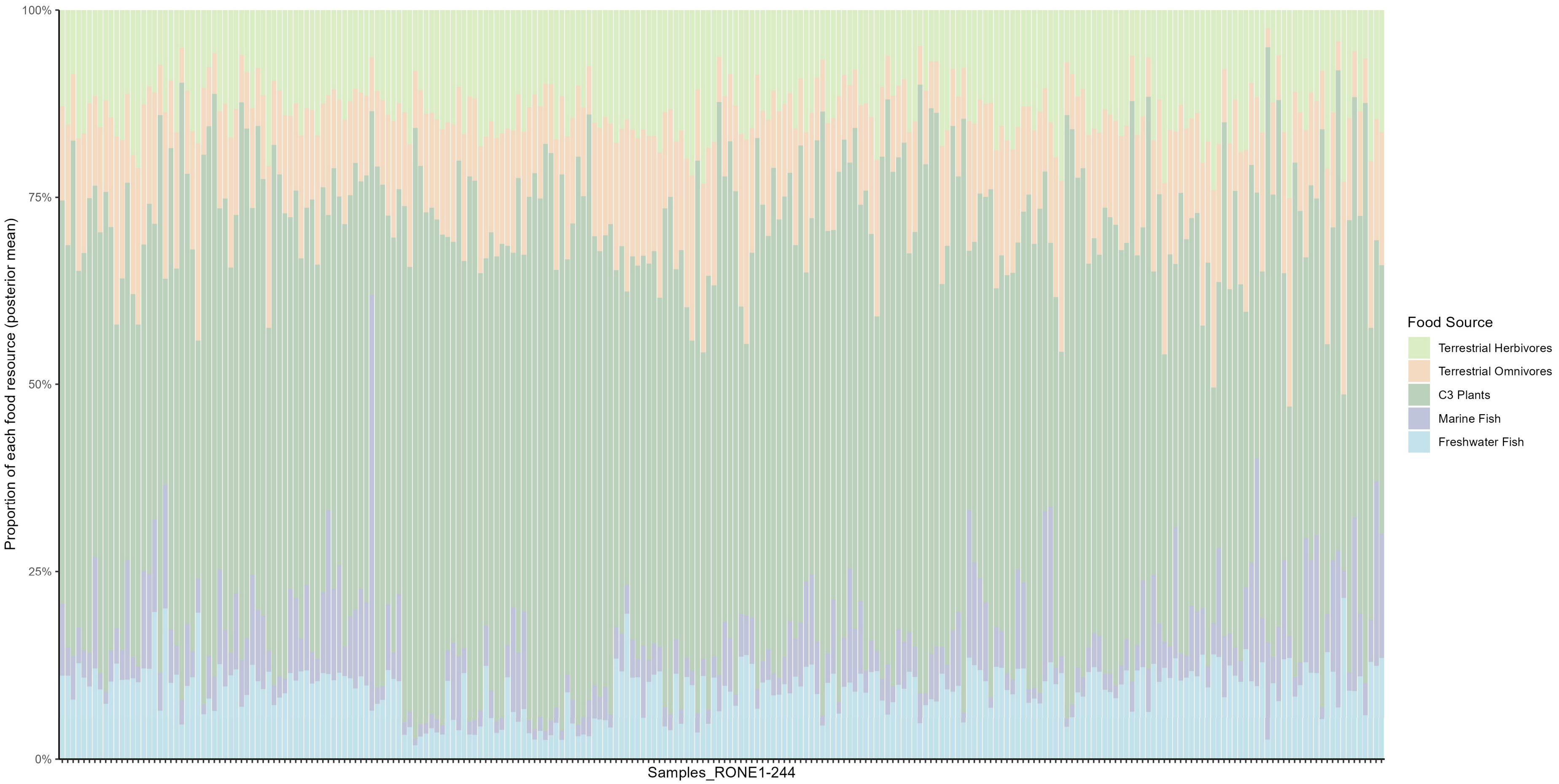

**Fig. S24| Dietary reconstruction for Roman northern England human samples.**

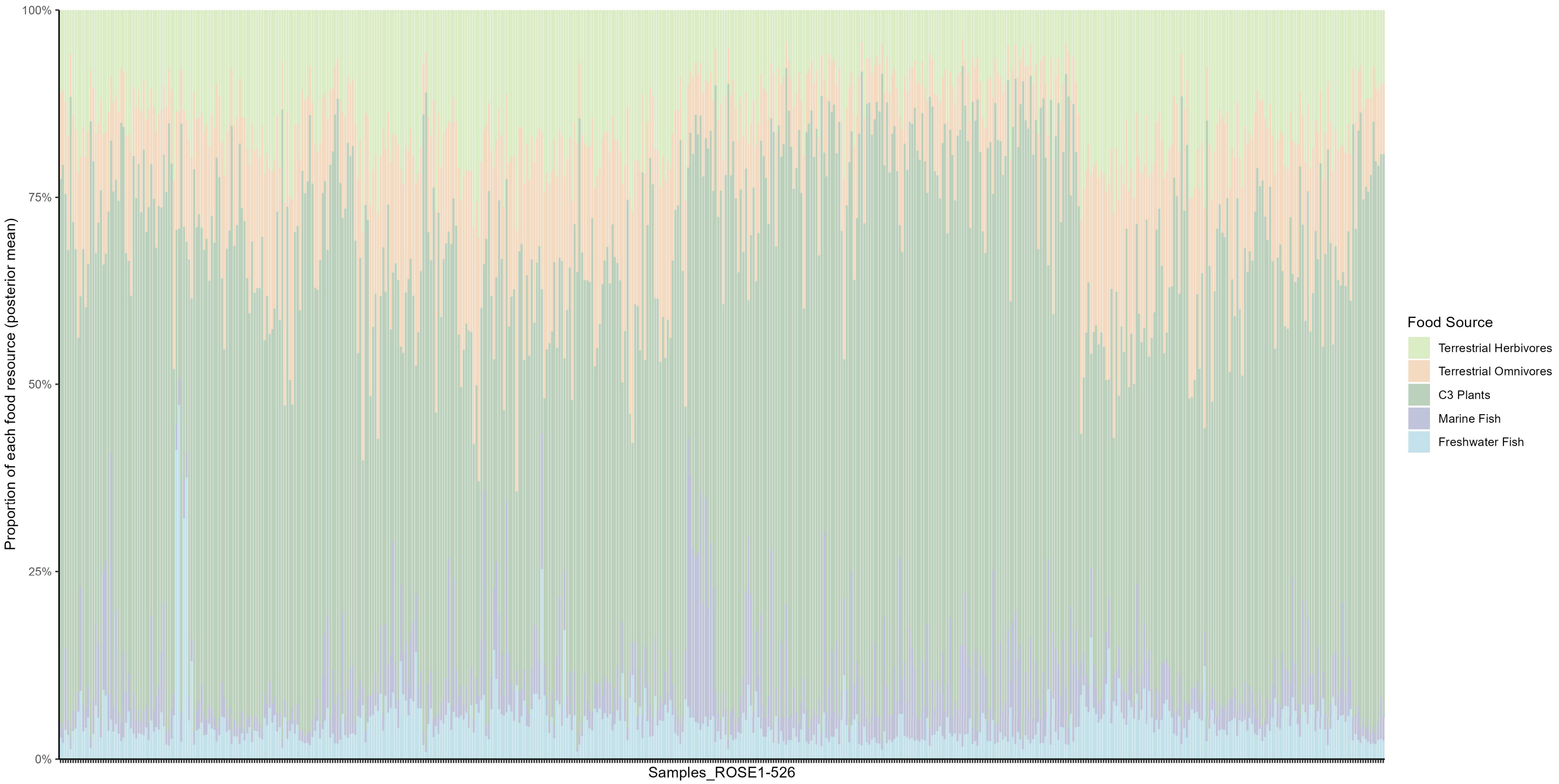

**Fig. S25| Dietary reconstruction for Roman southern England human samples.**

#### 2G. Early Medieval period

##### Scotland

During the Early Medieval period, diets in Scotland remained primarily based on terrestrial $C_3$ resources (Fig. S26)[29,32,75–79]. However, 23 samples from several sites show strong evidence of marine resource consumption. Out of a total of 165 samples from Early Medieval Scotland, these individuals represent approximately 14% of the dataset.

Sites that show evidence of marine resources consumption include Westness (Sample_EMS12 and 13)[75], Cnip (Sample_EMS15 and 16)[75], Portmahomack (Sample_EMS53 and 111)[76,78], Auldhame (Sample_EMS94)[77], Skaill House (Sample_EMS100)[32], and Newark Bay (Sample_EMS127, 131, 134, 135, 137, 139, 140, 143, 144, 145, 154, 158, 159, 160, and 165)[79] (Fig. S26). These samples, with $\delta^{13}C$ values ranging from –16.9‰ to –19.1‰ and $\delta^{15}N$ values from 11.5‰ to 15.92‰, all indicate strong marine resource consumption. Four samples from Westness and Cnip[75] are associated with Viking Age burials, and 15 samples[79] from Newark Bay also date to the Viking period. This suggests that within Viking Age populations, some individuals did indeed consume higher amounts of marine-based foods. However, not all Viking individuals followed a marine-oriented diet[79], and terrestrial resource consumption remained the dominant dietary pattern during this period. The remaining samples that show evidence of marine resource consumption may reflect differences in social status. For example, high-ranking monks may have consumed more fish during special feast days[76].

Another intriguing pattern is observed in the Portmahomack region[76,78], where all samples exhibit strikingly high $\delta^{15}N$ values. However, the original excavation reports note only limited recovery of fish bones, suggesting minimal freshwater fish consumption. Yet $\delta^{15}N$ values of such magnitude cannot be explained by terrestrial resource consumption—for instance, one individual shows $\delta^{13}C$ and $\delta^{15}N$ values of –19.9‰ and 14.6‰, respectively. On average, $\delta^{15}N$ values from this region are even higher than those from Newark Bay, where substantial marine resource consumption was documented.

There is no evidence indicating the influence of manuring or sea spray effects in this area, nor are these factors mentioned in the original study. If manuring or sea spray effects had been present, the isotopic values of local fauna would also be expected to reflect this influence—yet this is not supported by our analysis of faunal isotopic data [Chen et al in prep]. Given this, we suggest that freshwater fish consumption at Portmahomack has been significantly underestimated, as supported by our posterior estimates (Samples_EMS17–85, Samples_EMS102–126) (Fig. S26). Terrestrial resources still dominated the diet in this region; however, 31 out of 94 samples show freshwater resource contributions exceeding 10%, with four individuals exceeding 20%.

**England**

In Early Medieval England, diets in the central and northern regions were predominantly based on terrestrial resources (Fig. S27)[60,80–86]. In this area, marine dietary influence appears to have been even more limited than in Scotland: out of 246 individuals, only 17 show marine resource contributions exceeding 10%. These individuals are mostly concentrated at St Wystan's Church[83] (Samples_EMCN91–107) (Fig. S27). As with high marine consumers identified in Scotland, these individuals are dated to the Viking period and some of them were buried in graves containing Scandinavian-style grave goods.

Individuals showing substantial freshwater resources consumption are primarily from the Trent Valley[81] (Samples_EMCN13 – 38) and Raunds Furnells[82] (Samples_EMCN44–90) (Fig. S27). At these two sites, 27 out of 73 individuals exhibit freshwater resource contributions exceeding 10%; however, only one shows a contribution greater than 15%. Our findings support the interpretation proposed in the original study. Both sites are located close to rivers, suggesting that freshwater resources were readily accessible.

The individuals from the Isle of Man are particularly noteworthy[84]. The original study suggested that elevated $\delta^{15}N$ values might have been influenced by sea spray effects. However, when compared with samples from central and northern England, the $\delta^{15}N$ values from the Isle of Man are not unusually high. Even without applying any manual correction to the $\delta^{15}N$ values, our model did not overestimate marine or freshwater resource consumption, indicating a diet based almost entirely on terrestrial resources. This outcome is consistent with the hypothesis proposed in the original study.

Among the individuals from southern England[44,69,72,85,87–93], 39 out of 395 show marine resource contributions exceeding 10% (Fig. S28). Notably, 31 of these are from St John's College, Oxford[87] (Samples_EMSE69 – 106) (Fig. S28). According to the original study, they were likely more closely related to Viking populations than to Early Medieval. This may explain the relatively high proportion of marine resources in their diets.

Out of a total of 385 individuals, 30 show freshwater consumption proportion between 10% and 15%, while 32 exhibit freshwater consumption exceeding 15% (Fig. S28). Individuals with higher levels of freshwater resource consumption are primarily from Yarnton[44] (Samples_EMSE107 – 115), Ridgeway Hill (Weymouth)[88] (Samples_EMSE116 – 125), Westfield Farm[92] (Samples_EMSE269 – 298) and Beaconsfield[93] (Samples_EMSE299–385) (Fig. S28).

The situation at Early Medieval Yarnton mirrors that of the Roman period. We argue that the original study[44] significantly underestimated the contribution of freshwater resources in this region. Freshwater sources were readily accessible, and as in the Roman period, the significant elevated $\delta^{15}N$ values observed here cannot be explained by manuring only, as the faunal isotopic data do not support this interpretation. Therefore, substantial freshwater fish consumption is the most plausible explanation.

The other archaeological sites showing high freshwater resource consumption are consistent with the conclusions of the original studies. Notably, the samples from Ridgeway Hill (Weymouth) exhibit exceptionally high levels of freshwater intake, in agreement with the original findings. The original analysis noted that the human isotopic values at this site lie within the range of Scandinavian data—particularly resembling those from Björnö (Northern Sweden, 10th–13th century) and Birka (Sweden, Viking Period, 9th–10th century)[88]. These results suggest that the Weymouth individuals likely had a diet similar to that of the Björnö and Birka communities, which has been interpreted as being heavily reliant on freshwater resources. The other two sites are both readily accessible to freshwater resources: Westfield Farm, located in the fenland near the River Ouse, and Beaconsfield, situated close to the River Thames.

**Wales**

Samples from Wales[94] do not exhibit any notable isotopic patterns and instead reflect a typical terrestrial-based diet (Fig. S29), with minimal consumption of marine or freshwater resources. Notably, similar to the Isle of Man, the original study suggested that the elevated $\delta^{15}N$ values along the western coast of Wales might result from sea-spray effects. However, when these data are placed within a broader comparative framework, the $\delta^{15}N$ values from this region are not particularly high relative to other areas. This suggests that the influence of sea-spray effects on isotopic values in this region may have been overestimated.

In conclusion, the diet during the Early Medieval period appears to have remained largely consistent with earlier periods, continuing to be primarily based on terrestrial resources. Marine consumption seems to have declined compared to the Roman period, and marine intake is rarely observed except among individuals identified as potentially associated with Viking populations. This pattern reflects the impact of broader societal transformations on dietary consumption. Historical sources indicate that, by the early Early Medieval period, the infrastructure that had supplied marine foods under the Romans had largely disintegrated[60].

Our modelling results suggest that freshwater resource consumption during the Early Medieval period may have been underestimated—especially at sites such as Portmahomack in Scotland and Yarnton in southern England. The unusually elevated $\delta^{15}N$ values at these locations cannot be adequately explained by manuring practices or sea-spray effects, and greater freshwater intake appears necessary to account for the observed isotopic signatures.

In addition, several riverine sites also show evidence of increased freshwater consumption. These include Trent Valley and Raunds Furnells in the central and northern region of England, as well as Ridgeway Hill (Weymouth), Westfield Farm, and Beaconsfield in southern England. Among them, Weymouth and Westfield Farm likely reflect substantial reliance on freshwater resources. These findings are consistent with previous qualitative isotopic assessments.

Given this, earlier studies suggesting limited freshwater exploitation during the Early Medieval period—even among communities situated near rivers[92]—may need to be reconsidered. Our results indicate that, at least at certain sites, a significant portion of the population did consume freshwater resources. Nevertheless, terrestrial-based diets remained dominant overall.

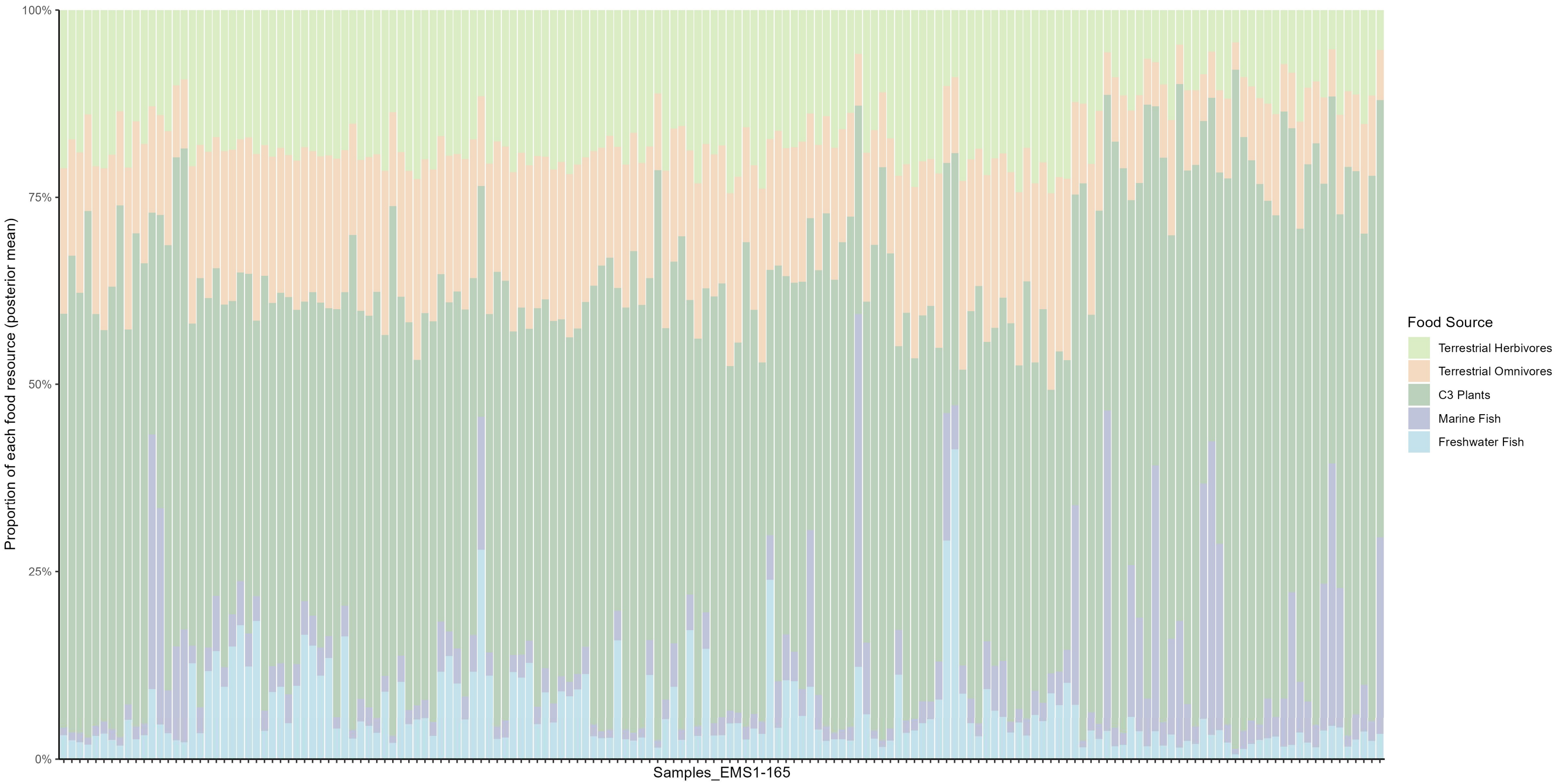

**Fig. S26| Dietary reconstruction for Early Medieval Scotland human samples.**

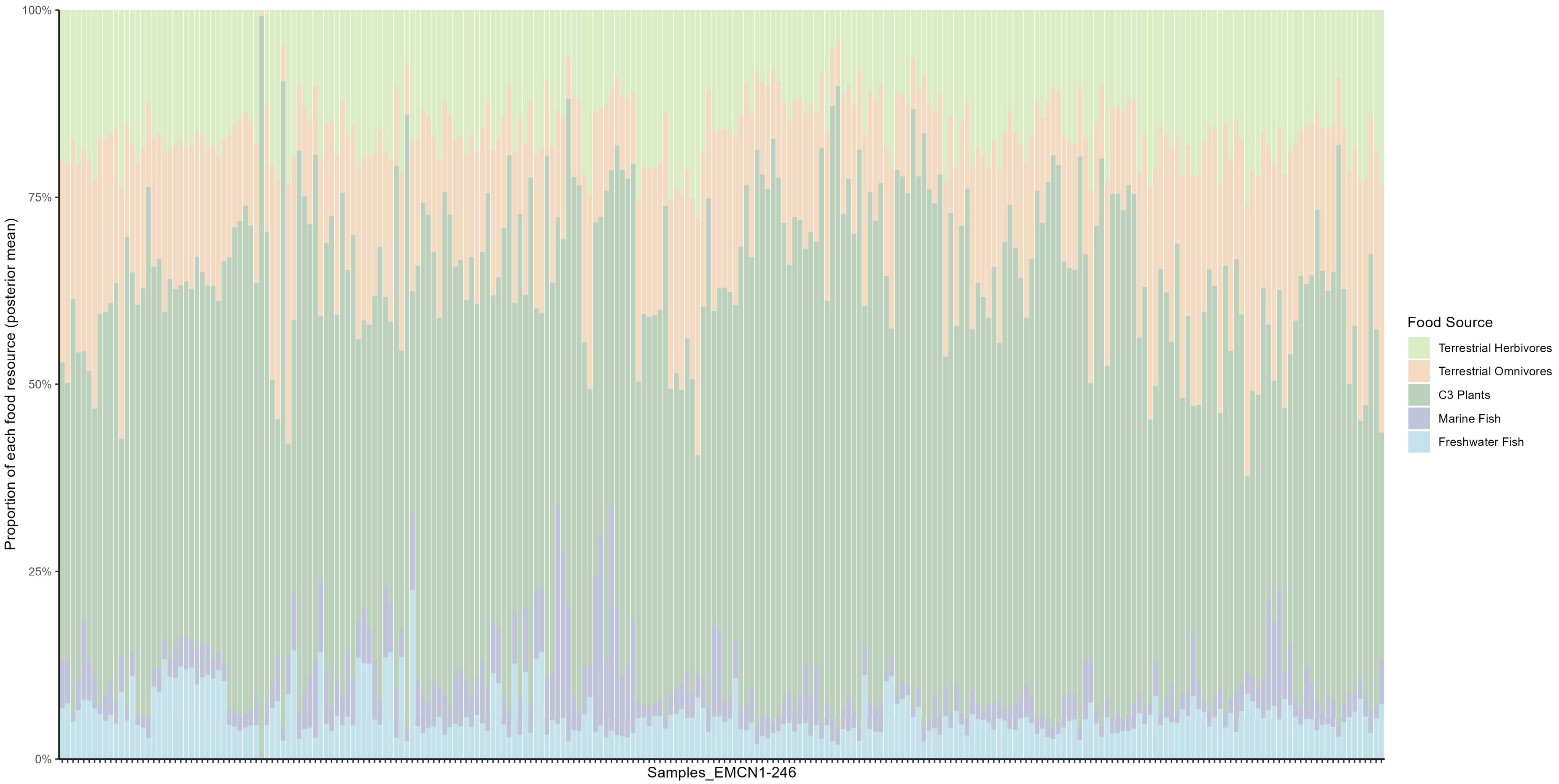

**Fig. S27| Dietary reconstruction for Early Medieval central and northern England human samples.**

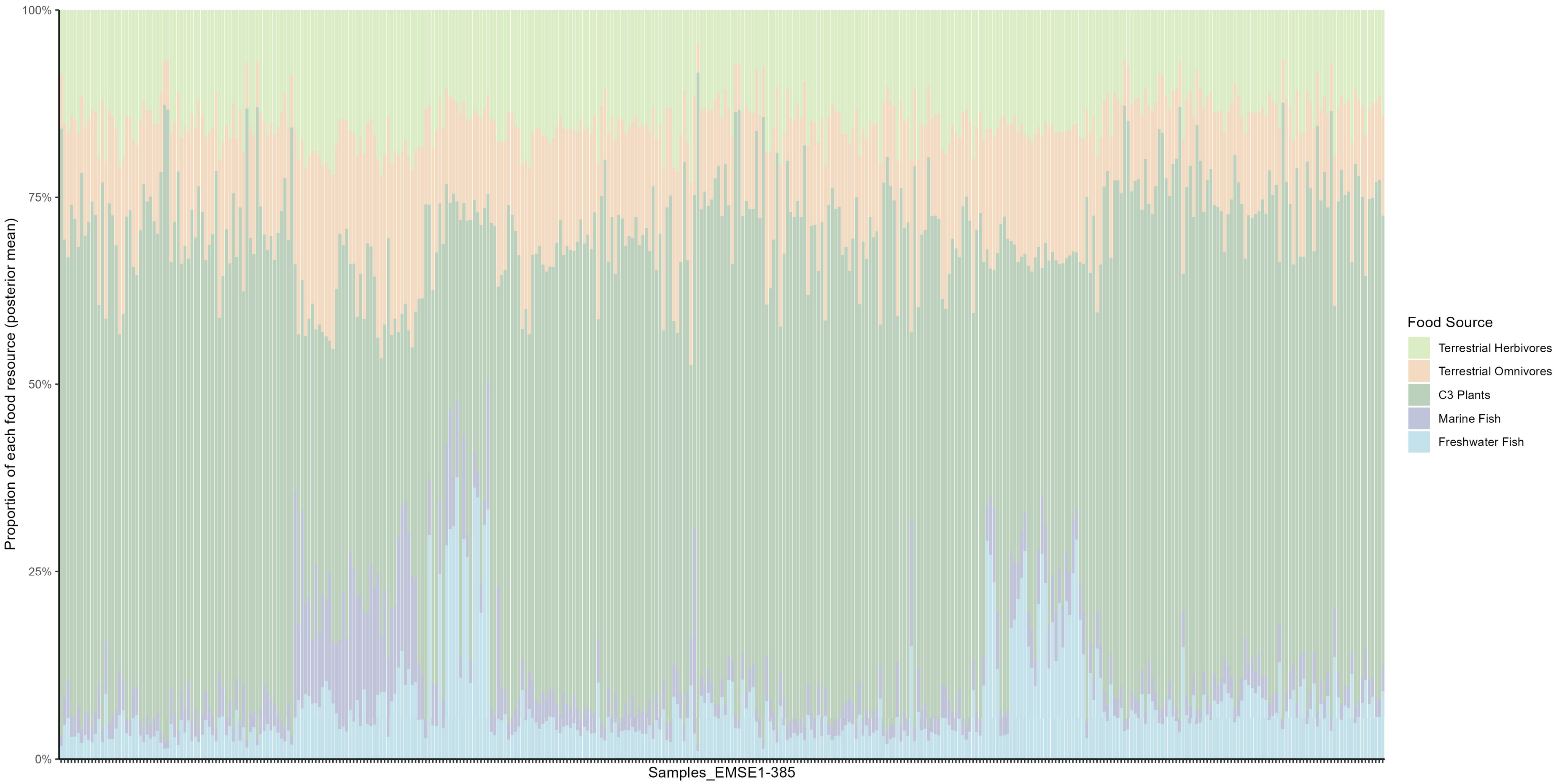

**Fig. S28| Dietary reconstruction for Early Medieval southern England human samples.**

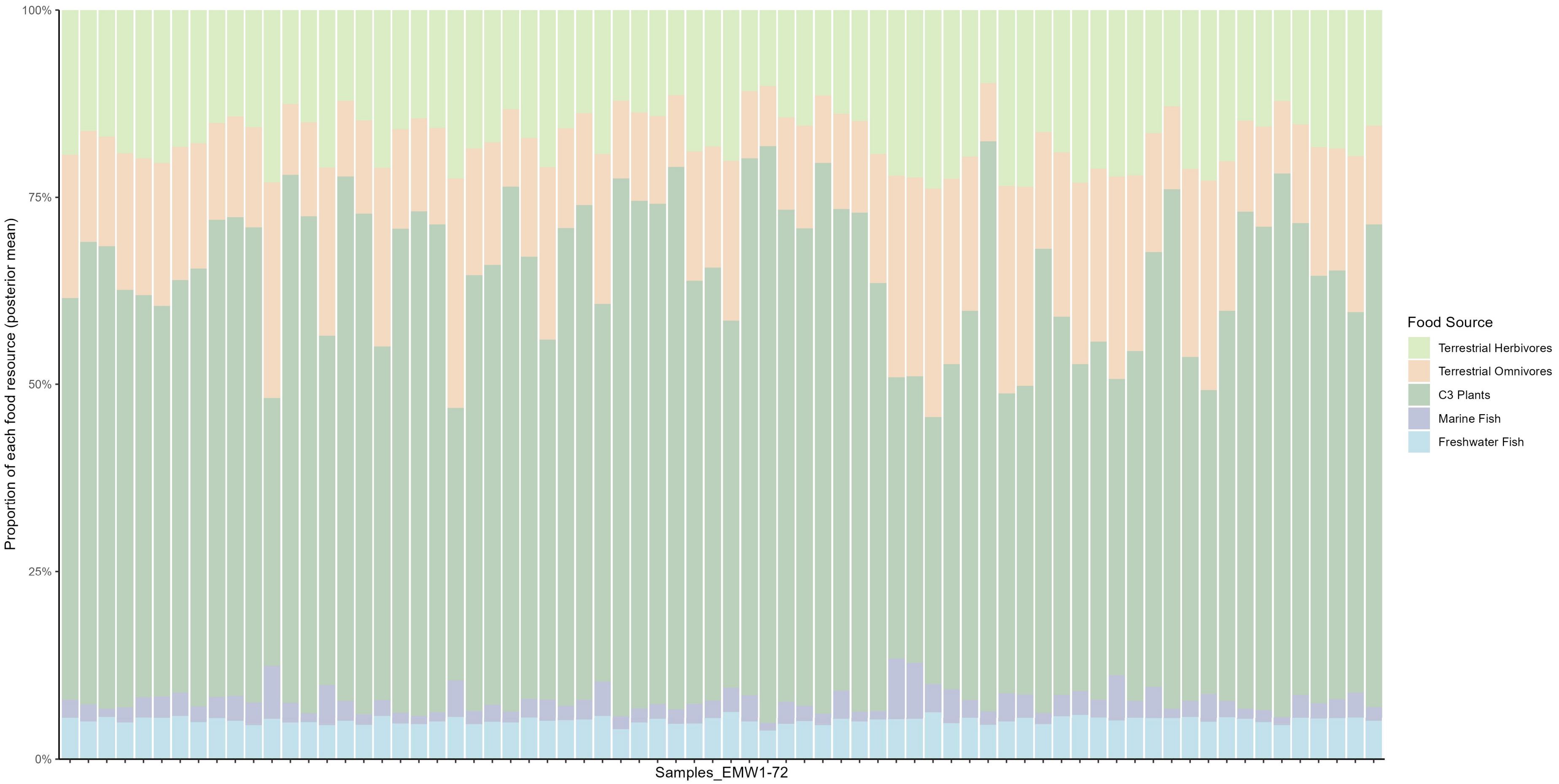

**Fig. S29| Dietary reconstruction for Early Medieval Wales human samples.**

#### 2H. Later Medieval period

##### Scotland

The dietary pattern of the Later Medieval period is particularly distinctive, characterized by a substantial increase in the consumption of marine fish. Evidence from Later Medieval Scotland[32,75–79,95,96] illustrates a marked rise in the number of individuals heavily reliant on marine resources (Fig. S30). While some high-status individuals during the Roman period, as well as certain Viking-associated individuals in the Early Medieval period, also exhibit comparable levels of marine consumption, their proportion within the overall population was considerably smaller than that observed in the Later Medieval period.

In Later Medieval Scotland, the main archaeological sites showing substantial marine resource consumption are located in Orkney[32,75,79] (Samples_LMS35 – 45, Samples_LMS114 – 125, Samples_LMS180 – 201) and Portmahomack[76,78] (Samples_LMS46–111, Samples_LMS156–179) (Fig. S30). These regions contribute a large share of the dataset, with a combined total of 135 individuals out of 201. Among these 135 samples, 79 show clear evidence of significant marine resource intake. Individuals from Orkney reveal some temporal patterns: those dated to the 11th–14th centuries show high levels of marine resource consumption, which declined to Viking-era levels in subsequent periods. However, our analysis suggests that this trend is regionally specific. Several 15th-century individuals from Portmahomack continue to exhibit strong marine consumption, and similar patterns are also observed in the samples from northern England.

At the same time, the consumption of freshwater resources declined significantly. An increase in marine fish consumption accompanied by a decline in freshwater fish consumption is a characteristic feature of the Later Medieval diet in Britain. The increase in marine fish consumption coincided with population growth, the expansion of the Atlantic fish trade, and the increasingly common Christian fasting practices in Britain. This transition has been referred to as the “fish event horizon”[60], beginning around AD 1000, marked by a significant increase in the proportion of marine species in fish bone assemblages.

It is also important to note that the remaining 66 samples from three other sites—Auldhame[77] (Samples_LMS1–34), Whithorn Cathedral Priory[96] (Sample_LMS112 and 113, Samples_LMS126–132, and Samples_LMS152–155), and St. Nicholas Kirk[95] (Samples_LMS133 – 151)—do not show strong evidence of marine resource consumption (Fig. S30). Among these 66 individuals, only five exhibit more than 10% marine intake. Most individuals from these sites display terrestrial $\delta^{13}C$ values. This indicates that even during the Later Medieval period, widespread marine consumption was not uniform across all regions.

**England**

The pattern in northern England is similar to that observed in Scotland, with some sites showing strong evidence of marine resource consumption, while others reflect a predominantly terrestrial diet (Fig. S31). However, compared to earlier periods, the proportion of individuals consuming marine resources increased significantly. Sites that exhibit notable marine intake include Fishergate (York)[60] (Samples_LMCN2 – 49, Samples_LMCN100–254), Priory Close[97] (Samples_LMCN255–333), and Clavering Place[97] (Samples_LMCN334–408) (Fig. S31). These samples represent the majority of the dataset, totaling 357 out of 509 individuals.

Individuals from Fishergate exhibit a clear temporal trend: during the 11th to 12th centuries, the number of individuals with high marine fish consumption was relatively low, whereas in the 13th to 16th centuries, this number increased substantially. The original study interpreted this as evidence that it took several decades—or even more than a century—for this new resource to become fully integrated into the diets of all groups in York's society[60]. This pattern stands in stark contrast to that observed in Orkney, where marine fish consumption was high between the 11th and 14th centuries but subsequently declined. These opposing trends suggest that the temporal dynamics of marine fish consumption in the Later Medieval period were likely region-specific, rather than a uniform pattern shared across Britain.

Individuals from Priory Close and Clavering Place are particularly interesting. These samples, dating from the 13th to 16th centuries, were analyzed using incremental dentine sampling. The results show that they consumed more marine food as they transitioned from adolescence to adulthood. The original study suggests that[97], assuming these individuals were friars, a detectable dietary shift occurred upon their entry into a religious order.

The remaining 152 samples from other sites do not show significant marine resource consumption. These sites include Wharram Percy[98] (Samples_LMCN409–442), St. Giles (Brough)[99,100] (Samples_LMCN61 – 71, Samples_LMCN493 – 509), Towton[100] (Samples_LMCN50 – 60), Warrington[100] (Samples_LMCN443 – 460), Box Lane (Pontefract)[99] (Samples_LMCN72 – 99), and Fishergate House[101] (Samples_LMCN461–492) (Fig. S31).

The inhabitants of Wharram Percy were mostly ordinary peasants, and since lower social classes typically substituted meat with dairy products rather than fish on fast days, the dietary differences observed here may therefore reflect social distinctions[100]. Some sites, such as St. Giles, Warrington, and Box Lane, show evidence of freshwater resource consumption. This is not surprising, as St. Giles and Warrington are located directly on rivers—the Swale and the Mersey, respectively[100]. This suggests that even in the Later Medieval period, when freshwater fish consumption was generally uncommon, some inexpensive freshwater species—especially eels—may still have been part of the daily diet of lower-class individuals.

The situation in northern England further demonstrates that access to marine resources was not universally available during the Later Medieval period. In fact, archaeological sites with low levels of marine consumption are more numerous than those with high marine intake, even though they account for a smaller proportion of the overall population.

There is only one sample from central England—King Richard III (Fig. S31)[102]. His diet displays a remarkably diverse pattern, with substantial intake of animal protein (Sample_LMCN1 in Fig. S31).

The samples from southern England mainly come from Oxford (All Saints, Oxford Castle, Westgate, Christ Church, and Oxford Castle Mound)[103], the Cathedral Close (Hereford)[104], St. Guthlac's Priory (Hereford)[104], and St. Mary Spital Cemetery[105]. The dietary reconstruction results from southern England appear to be inconsistent with the widespread marine resource consumption typically associated with the Later Medieval period (Fig. S32, Fig. S33). Although the proportion of individuals with more than 10% marine resource consumption in southern England is not significantly lower than in Scotland and northern England, the proportion of individuals with high levels of marine consumption (above 25%) is markedly reduced (Fig. S30, Fig. S31, and Fig. S32).

St. Mary Spital Cemetery[105] is the only site that exhibits substantial marine resource consumption. St. Mary Spital Cemetery provides the largest number of samples, with bone isotopic data from 129 individuals and 374 incremental dentine samples from 21 individuals. Among the 129 bone samples (Samples_LMSEbone83 – 211), 70 individuals show evidence of more than 10% marine resource consumption, including 17 individuals with values exceeding 15% (Fig. S32). Of the 374 incremental dentine samples (Samples_LMSEincremental132–505), 127 indicate more than 10% marine intake, with 48 exceeding 15% (Fig. S33). The original study compared isotopic values between individuals classified as “attritional burials” and those associated with famine burials, demonstrating that famine individuals consumed less protein—a finding supported by our modeling results. However, both groups include a proportion of individuals who consumed marine resources, and this proportion does not differ significantly between them. This suggests that access to marine resources was not substantially affected by whether or not individuals had experienced famine.

The marine resource consumption at other sites can be summarized as follows (bone samples: number of individuals with >10% marine resource consumption / total number of individuals): Cathedral Close (2/19) (Samples_LMSEbone41–59)[104], St. Guthlac's

Priory (7/29) (Samples_LMSEbone1 – 29)[104], and Oxford (5/34) (Samples_LMSEbone30–40 and Samples_LMSEbone60–82)[103] (Fig. S32). Among the incremental dentine samples, 29 of 202 individuals from St. Guthlac's Priory (Samples_LMSEincremental506 – 707)[104] and 1 of 131 from Oxford (Samples_LMSEincremental1–131)[103] exhibit this pattern (Fig. S33). It is evident that, apart from the previously discussed St. Mary Spital Cemetery, marine resource consumption at the remaining sites is significantly lower than at the major marine-consuming sites in Scotland and northern England. The Oxford samples are primarily from the early phase of the Later Medieval period (around the 11th century, during the Norman Conquest of England), whereas the samples from the other three sites predominantly date to the late phase of the Later Medieval period. This suggests that the relatively low levels of marine resource consumption observed at these sites are more likely linked to geographic location than to temporal factors.

Taken together with the results from northern England and Scotland, this further supports the observation that substantial marine resource consumption during the Later Medieval period was more common in certain regions and much less so in others. Marine consumption does not appear to follow a consistent chronological pattern: some areas show high levels of marine intake in earlier periods followed by a decline (e.g., Orkney), while others exhibit increased marine consumption in later periods (e.g., Fishergate, York).

Another interesting pattern concerns the consumption of freshwater fish, which is primarily reflected in the incremental dentine samples. We summarized the data as follows (number of individuals with >10% freshwater resource consumption / total number of individuals): Oxford (13/131) (Samples_LMSEincremental1–131)[103], St. Mary Spital Cemetery (46/374) (Samples_LMSEincremental132 – 505)[105], and St. Guthlac's Priory (34/202) (Samples_LMSEincremental506 – 707)[104] (Fig. S33). Although freshwater fish were rarely discussed in the original study, some incremental dentine samples exhibit very high $\delta^{15}N$ values without a corresponding elevation in $\delta^{13}C$ (e.g., −20.6‰, 12.6‰). Such isotopic signatures are difficult to interpret as resulting from increased consumption of omnivores or manuring effects. This is especially relevant in British sites where elevated $\delta^{15}N$ values have previously been attributed to manuring, yet the isotopic values of local animals do not show unusual characteristics. As a result, the role of freshwater fish may have been underestimated—a possibility that has been discussed before at certain Early Medieval sites. Therefore, this pattern observed in southern England during the Later Medieval period may be explained by increased consumption of freshwater fish during adolescence, a key stage of physical growth.

In summary, traditional perspectives characterize Later Medieval diets by a sharp increase in marine resource consumption, and our results support this general trend.

However, our findings also indicate that not all sites show high levels of marine intake. In fact, sites with low levels of marine consumption are more numerous than those with high consumption, although there are fewer individuals in these sites. The temporal pattern of marine consumption is not uniform but varies between sites. Freshwater fish may have served as a significant resource in some lower-status or river-adjacent communities. In addition, samples from southern England suggest a preference for freshwater fish consumption during adolescence. Famine appears to have affected protein intake, but had little impact on access to marine resources.

Although we conducted dietary modeling for each incremental dentine sample (Fig. S33), the incremental samples from southern England were not included in the time-series analysis. This exclusion is due to the exceptionally large number of such samples from the region (707 samples), which could disproportionately inflate the representation of certain age stages. Since dentine increments may reflect dietary patterns specific to adolescence or childhood, their overrepresentation could compromise the overall reliability of the time-series results. In fact, including these incremental samples would result in severely biased estimation of marine/freshwater resource consumption in southern England. Moreover, these samples originate from only three archaeological sites, yet they contribute 707 individual data points—constituting 43% of the entire Later Medieval British dataset (N = 1,637). Our analysis reveals considerable dietary variation across different locations during this period. Therefore, over-reliance on samples from a limited number of sites would inevitably distort the overall reconstruction of dietary patterns across Britain.

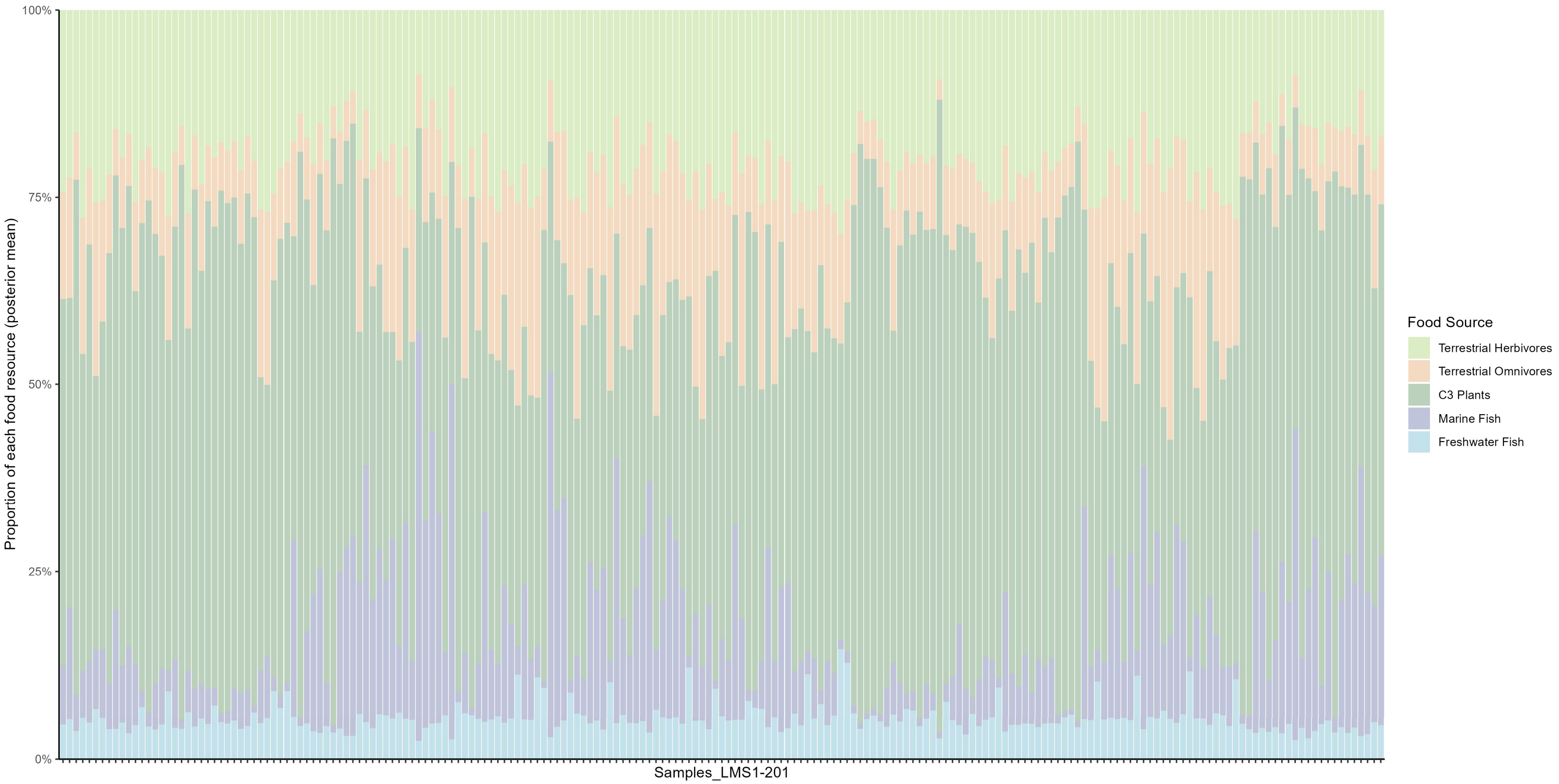

**Fig. S30| Dietary reconstruction for Later Medieval Scotland human samples.**

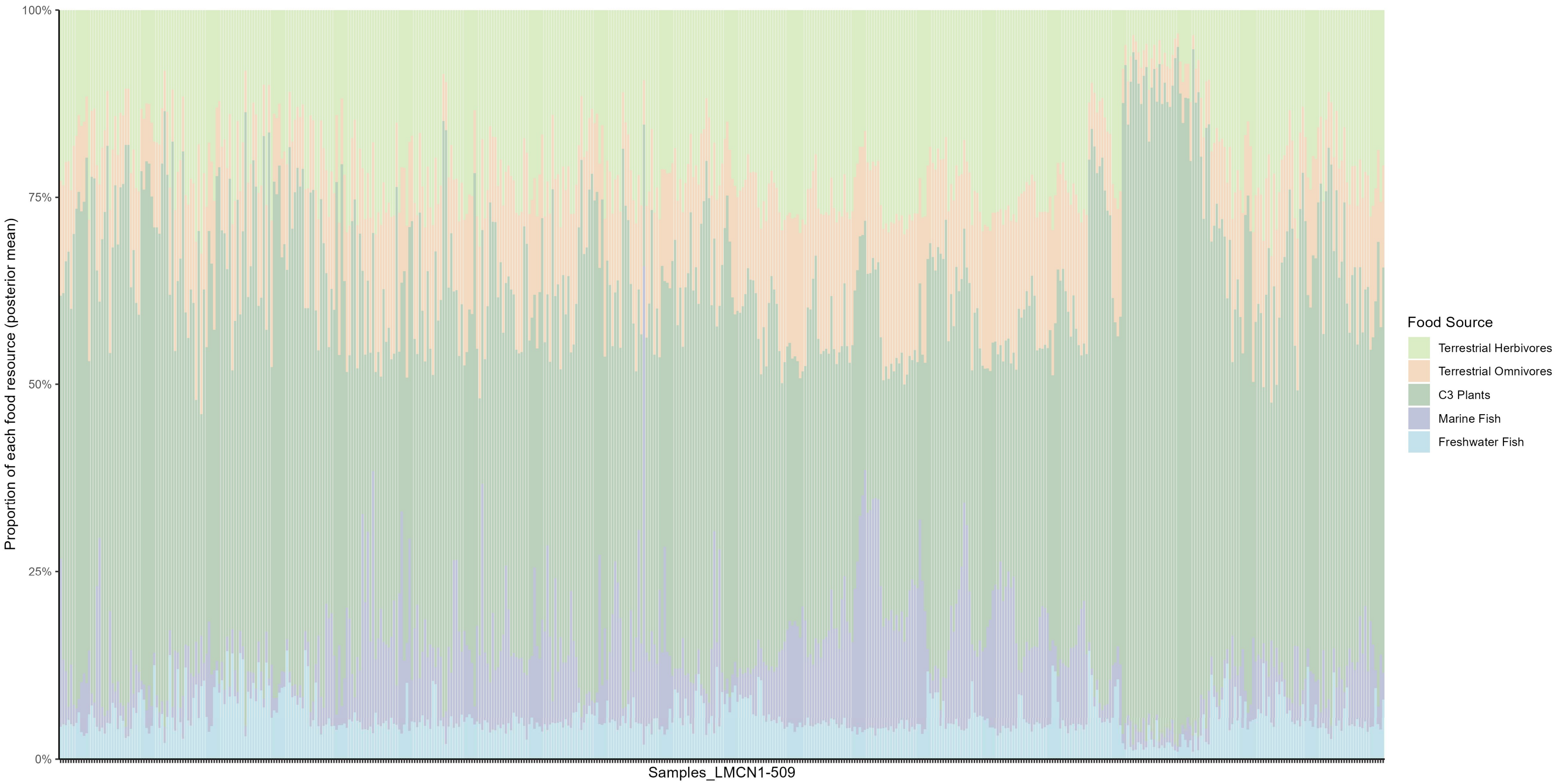

**Fig. S31| Dietary reconstruction for Later Medieval central and northern England human samples.**

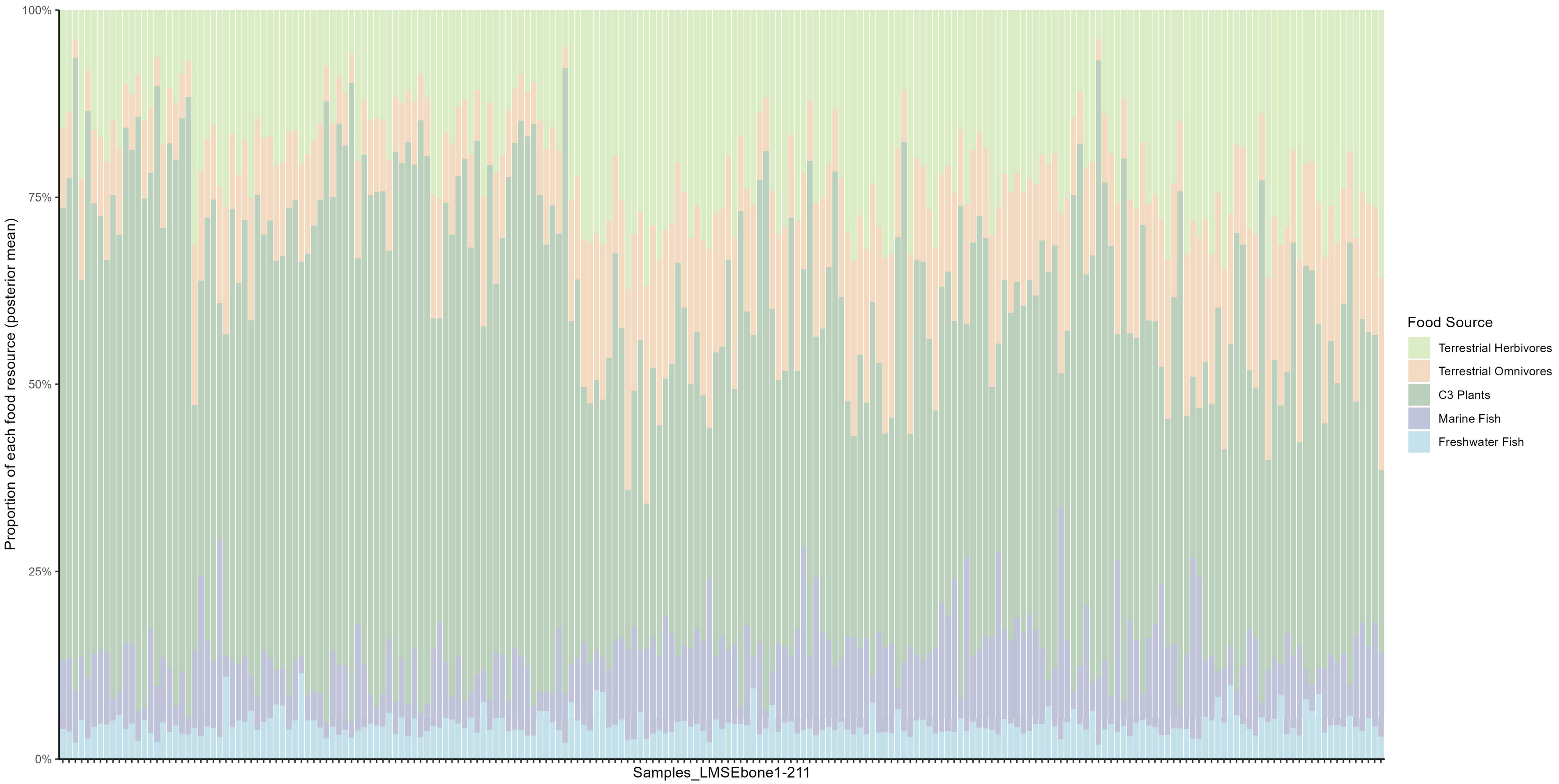

**Fig. S32| Dietary reconstruction for Later Medieval southern England human samples.**

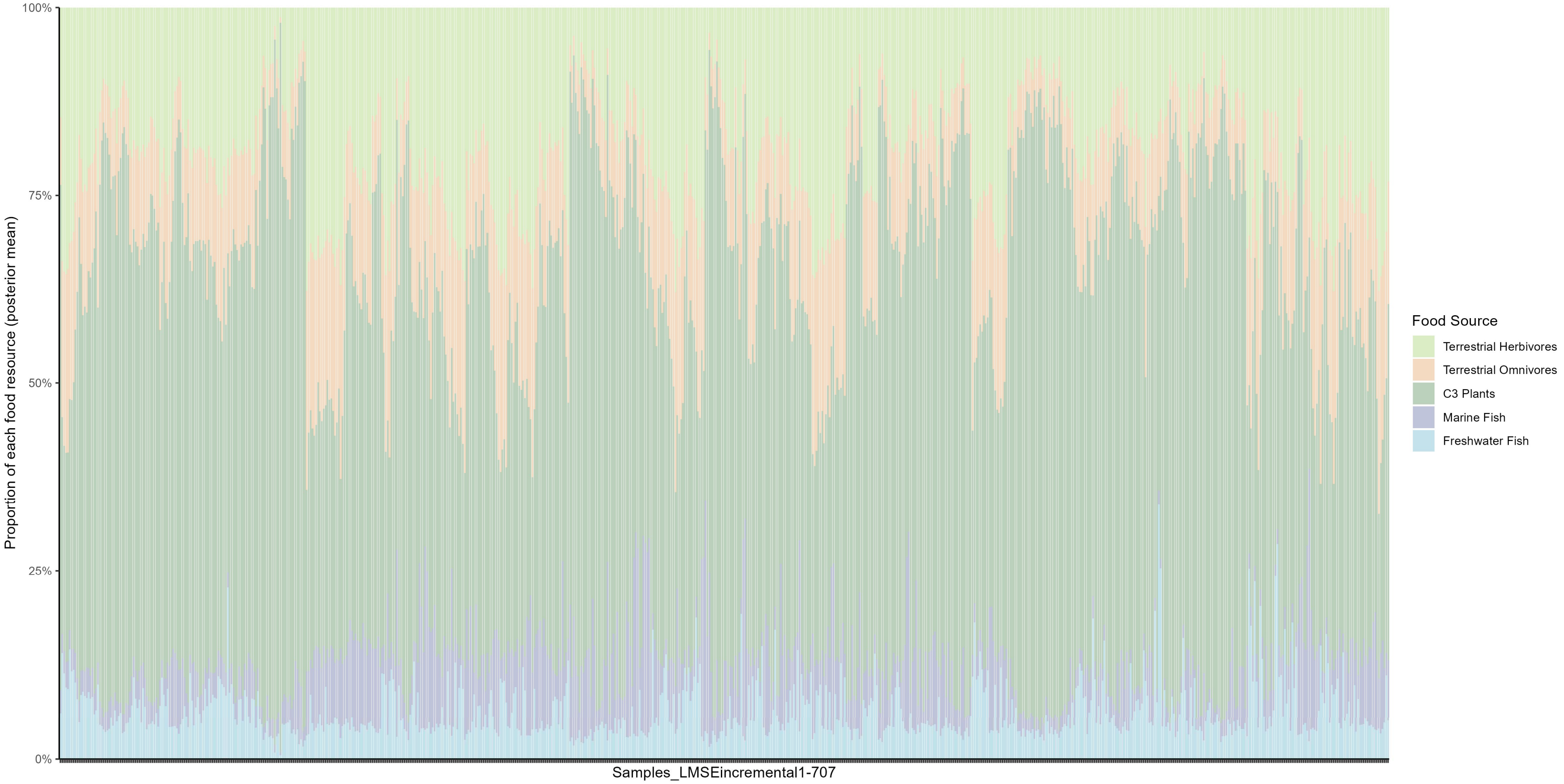

**Fig. S33| Dietary reconstruction for Later Medieval southern England incremental dentine samples.**

#### 2I. Post-Medieval period

##### Scotland

There are relatively few samples from Post-Medieval Scotland (Fig. S34). Those available—Westness (Sample_PMS1)[75], Portmahomack (Sample_PMS2 and 3)[76], Auldhame (Sample_PMS4 and 5)[77], and St. Nicholas Kirk (Samples_PMS6–18)[95]—are drawn from studies primarily concerned with Later Medieval diets. The Post-Medieval individuals were included only incidentally, representing a small subset within broader Later Medieval dietary research. Only two of these 18 individuals show substantial marine resource consumption (Sample_PMS1 and 14). The reliance on marine foods among the Post-Medieval Scottish samples appears to have been limited.

Due to the limited number of samples, it is not possible to determine whether marine resource consumption significantly declined in Post-Medieval Scotland. However, previous studies have noted that although the sample size is small, the Post-Medieval individuals tend to exhibit less enriched in carbon and nitrogen isotopes compared to most Later Medieval samples[76].

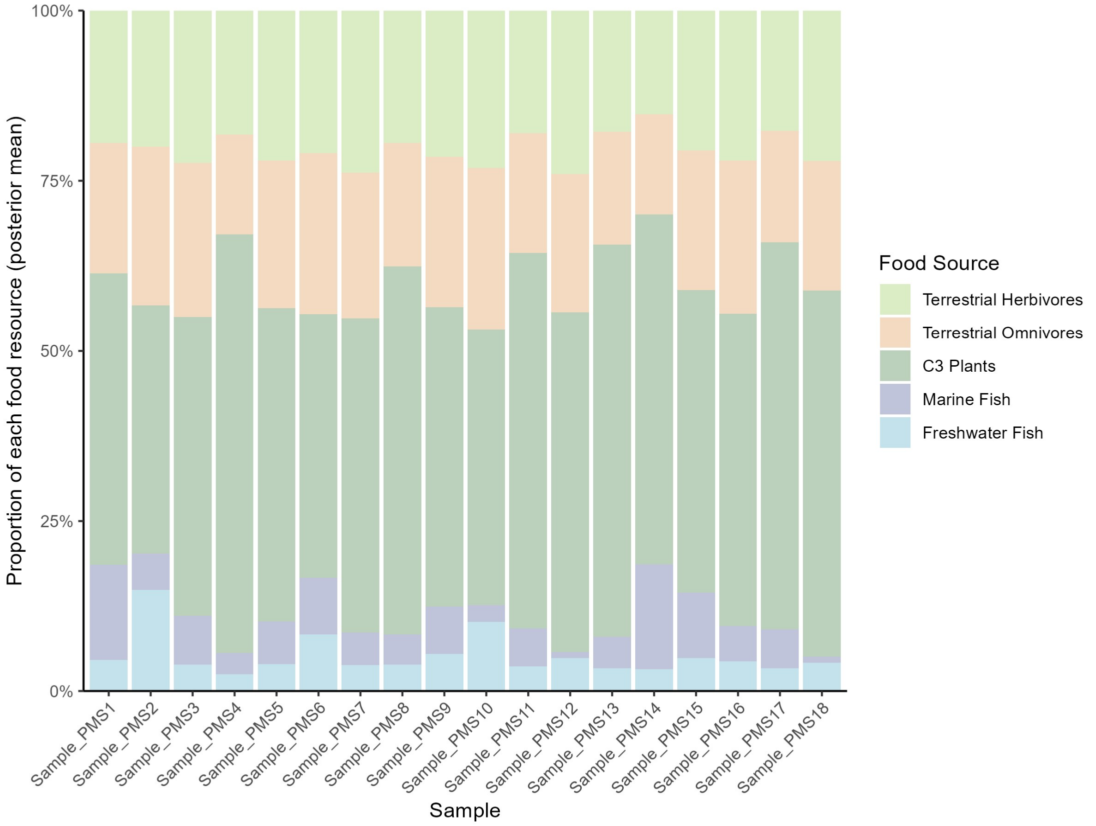

**Fig. S34| Dietary reconstruction for Post-Medieval Scotland human samples.**

##### England

The northern England dataset includes samples from two sites: Palace Green, Durham (Samples_PMNE1–12)[106] and the Church of All Saints, York (Samples_PMNE329–344)[60]. Samples_PMNE13–328[106] represent incremental dentine data derived from the

12 individuals at Palace Green, Durham.

When considering lifetime averages, individuals from Durham predominantly show a terrestrial $C_3$-based diet, with only one individual (Sample_PMNE2) exhibiting a strong marine signal (Fig. S35). However, the incremental dentine data reveal that some individuals may have consumed higher proportions of marine resources during specific periods of their lives (Samples_PMNE13−328) (Fig. S35). This highlights that an individual's diet may vary substantially over the course of their life.

The samples from York display a stronger marine dietary signal (Samples_PMNE329–344) (Fig. S35). According to the original interpretation[60], this may reflect the continued public observance of fasting periods during the Elizabethan era, despite the diminished influence of Catholic fasting regulations following the Reformation. York's relative proximity to the coast may further explain why marine consumption did not decline in the Post-Medieval period.

As with the incremental dentine samples from the Later Medieval period, the Durham incremental samples are not included in the time-series modelling due to their large number. Our analysis suggests considerable regional heterogeneity in diet during this period, and disproportionately expanding the sample size of a single site—such as Durham—could distort the dietary averages for Britain.

The modeling results for southern England[107–113] indicate that 128 individuals—representing 28% of the total—consumed more than 10% marine resources (Fig. S36). This proportion is nearly three times higher than that observed during the Early Medieval period, when only around 10% of individuals exhibited high marine consumption. However, it marks a decline compared to the Later Medieval period, during which the proportion of high marine intake individuals reached 40% in Scotland, 34% in northern England, and 40% among bone samples from southern England.

Sites with elevated marine resource consumption include several from London—namely St Barnabas (Samples_PMSE84–108)[108], Lukin Street (Samples_PMSE229–327)[111], and Spitalfields (Samples_PMSE328–450)[112,113] (Fig. S36). At St Barnabas, most individuals exhibit high marine intake, and historical interpretations suggest that the population may have been relatively affluent. Similarly, the majority of individuals from Lukin Street show strong isotopic signals of marine food consumption. Notably, these results offer the first bioarchaeological support for historical accounts describing the widespread consumption of inexpensive marine foods in Post-Medieval London. A comparable pattern is observed at Spitalfields, where many individuals also show substantial marine resource intake. In the original study, the Spitalfields samples were interpreted as a predominantly terrestrial diet, based on the conclusion that elevated $\delta^{15}N$ values were not accompanied by corresponding increases in $\delta^{13}C$. However, we find that the majority of individuals from this site exhibit isotopic patterns that are fully consistent with substantial marine resource consumption. It remains unclear how the original qualitative assessment concluded a terrestrial diet based on such isotopic values with a mean $\delta^{13}C$ of -18.9‰.

However, not all London sites follow this pattern. Samples from the Queen's Chapel of the Savoy (Samples_PMSE19–83)[108] reveal substantial dietary heterogeneity: while most individuals consumed a predominantly $C_3$-based terrestrial diet, a subset displays strong evidence of marine resource use (Fig. S36). This diversity aligns with historical records indicating that parishioners, seamen, soldiers, patients, and prisoners were buried at the site.

Samples from St. Michael's Litten cemetery in Chichester (Samples_PMSE189–228)[110] predominantly exhibit a terrestrial dietary pattern, although a small number of individuals show evidence of marine resource consumption (Fig. S36).

In contrast, individuals from the Mary Rose (Samples_PMSE1–18 and 451–459)[107,114] display almost no marine intake (Fig. S36). These individuals were associated with the Mary Rose shipwreck, and are therefore all linked to naval dietary practices. The apparent reliance on terrestrial animal protein rather than fish aligns with historical records of naval diet.

Notably, some samples from Plymouth Hospital (Samples_PMSE109–158)[109] exhibit elevated $\delta^{13}C$ values alongside relatively low $\delta^{15}N$ values. The original study interpreted this pattern as evidence of $C_4$ resource consumption. However, based on our isotopic observations, at least some individuals exhibit values consistent with marine resource consumption. Our modeling results also indicate that a considerable number of these individuals are likely to consume marine resources (Fig. S36). These samples are derived from naval hospitals, and according to historical records, it is unlikely that such individuals had regular access to fish-based diets. This discrepancy highlights the difficulty of reliably interpreting dietary sources in the absence of appropriate $C_4$ baseline. The original study also acknowledged that many of these individuals may have spent most of their lives in east coast of North America, and thus their isotopic signatures may not reflect contemporary British dietary patterns.

Samples from another naval institution, the Royal Naval Hospital Haslar (Samples_PMSE159 – 188)[109], Gosport, exhibit a predominantly terrestrial dietary pattern (Fig. S36). Given the complex and often non-local origins of naval personnel[109], and the fact that such individuals represent only a small and unrepresentative subset of the British population, we excluded all three naval-related assemblages from the time-series modeling.

Freshwater resource consumption during this period does not exhibit any distinct patterns and does not appear to have been a major dietary component at any site.

Overall, Post-Medieval diet exhibits several distinct characteristics. In major urban centers such as London and York, marine resource consumption remained substantial. In contrast, individuals from more rural areas appear to have relied far less on marine foods compared to the Later Medieval period—possibly reflecting the declining influence of fasting regulations. Naval diets, meanwhile, seem to have been almost entirely based on terrestrial resources.

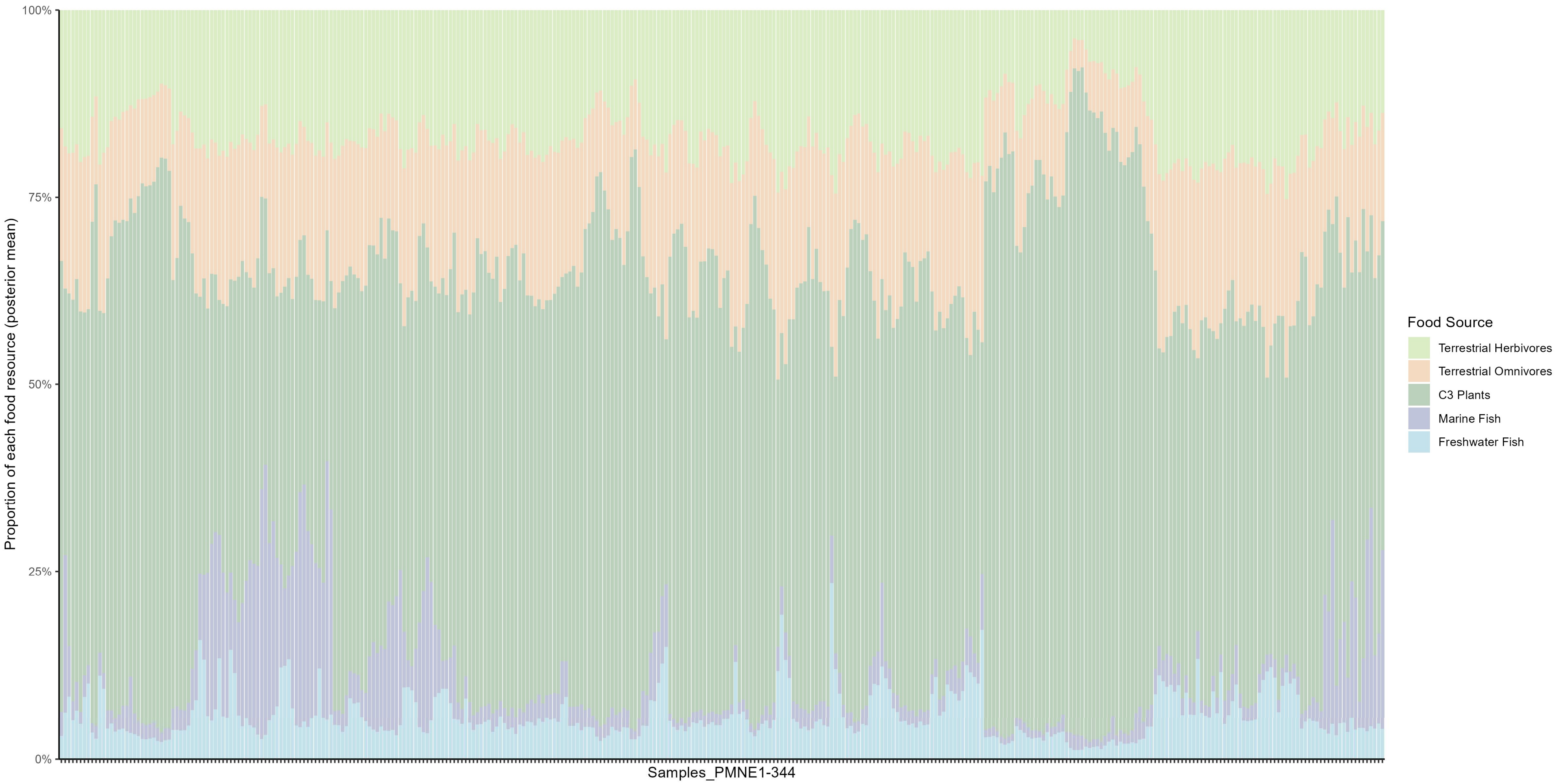

**Fig. S35| Dietary reconstruction for Post-Medieval northern England human samples.**

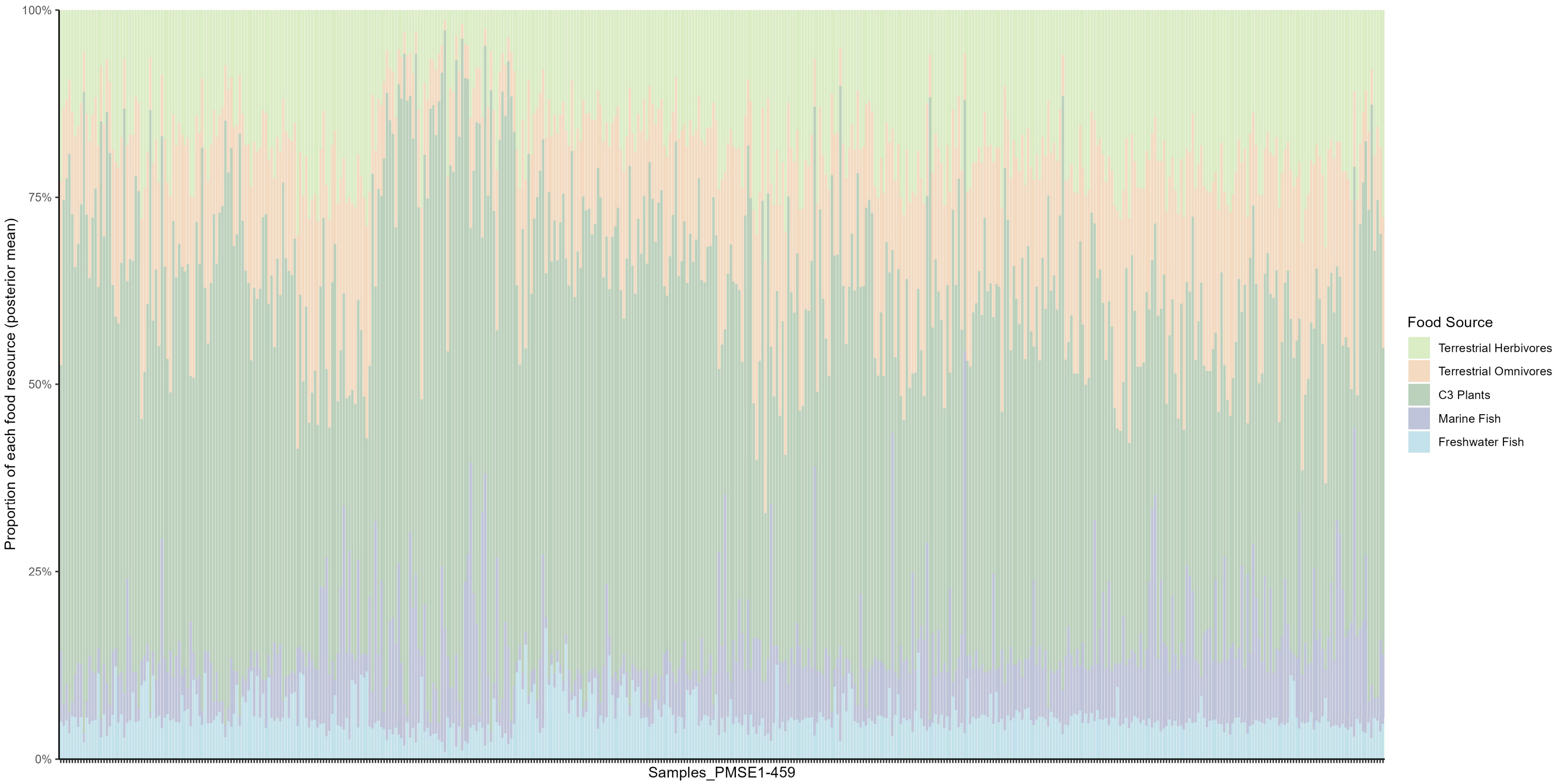

**Fig. S36| Dietary reconstruction for Post-Medieval southern England human samples.**

#### 2J. Detailed results of bootstrap sampling for dietary time-series models

To construct the dietary and dairy time series, we applied a bootstrap resampling. For each time bin, individuals were resampled with replacement 1,000 times to estimate the mean dietary or dairy-use proportion and to characterize the associated uncertainty. Here, we present the probability density distributions of the bootstrap estimates for each food category within each time bin.

Overall, the bootstrap sampling yields robust estimates. Among the time series used for visualization, 273 out of 295 time bins (92.5%) exhibit approximately normal bootstrap distributions (Fig. S37). Notable instability is observed for the time bins centered at 1300 BC and 1100 BC (Fig. S37), which can be attributed to sparse sampling during this period. Estimates for terrestrial herbivores, marine fish, and freshwater fish during the Mesolithic (time bin midpoints from 8500 to 4400 BC) are also less stable (Fig. S37b, c, d), again reflecting limited sample sizes.

By contrast, the time series used for causal discovery are highly stable across nearly all time bins (Fig. S38). This increased stability arises from the use of sliding windows, which incorporate a larger number of samples within each window and thereby reduce sampling variance. For the dairy time series, nine time bins display multimodal distributions (time-bin midpoints at 3100–2800 BC, 1000–700 BC, and 500 BC), whereas the remaining bins show approximately normal distributions (Fig. S39).

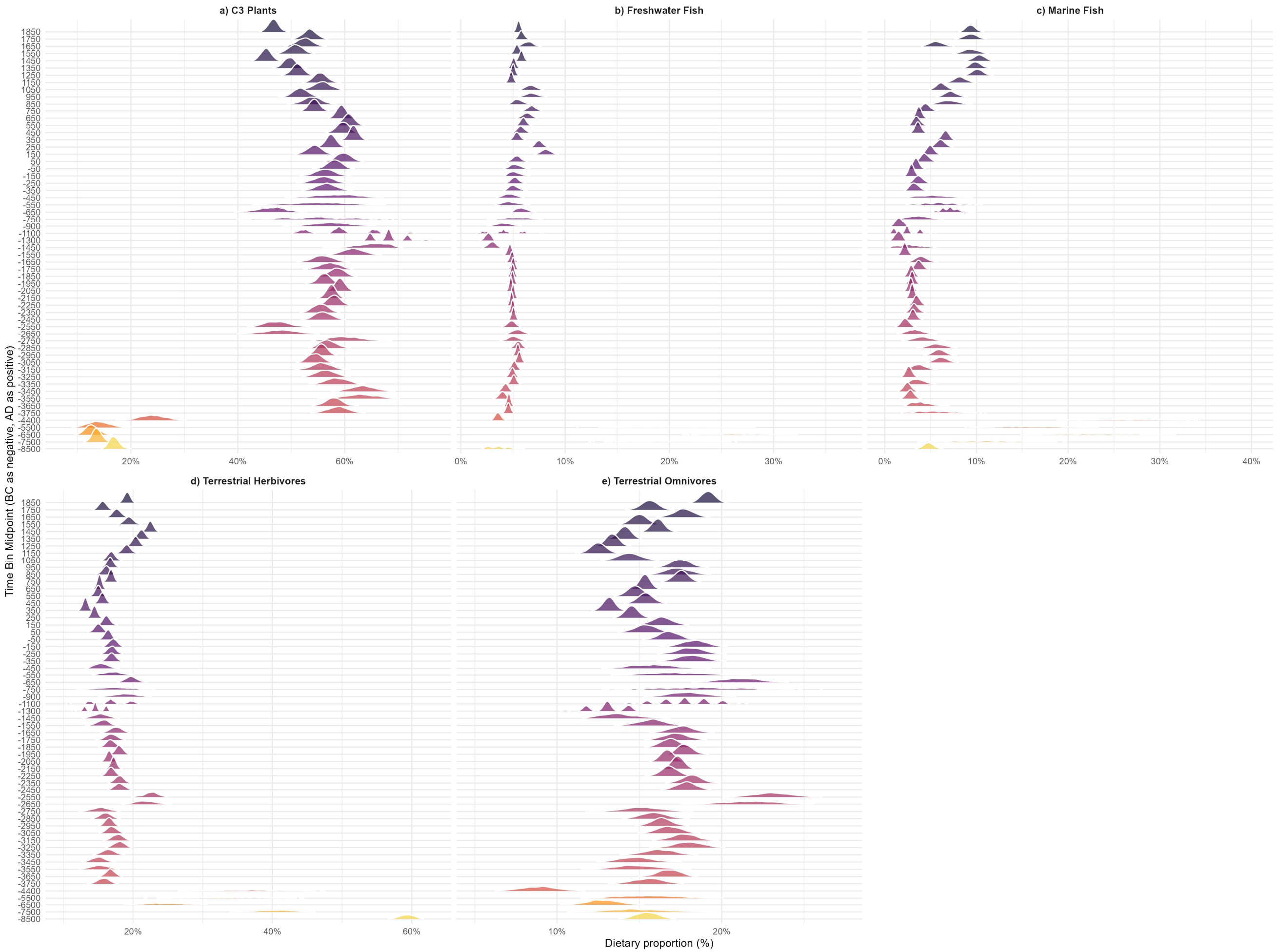

**Fig. S37| Probability density distribution of bootstrap dietary time-series models (visualization) by time bin. a),** $C_3$ plants. **b),** Freshwater fish. **c),** Marine fish. **d),** Terrestrial herbivores. **e),** Terrestrial omnivores.

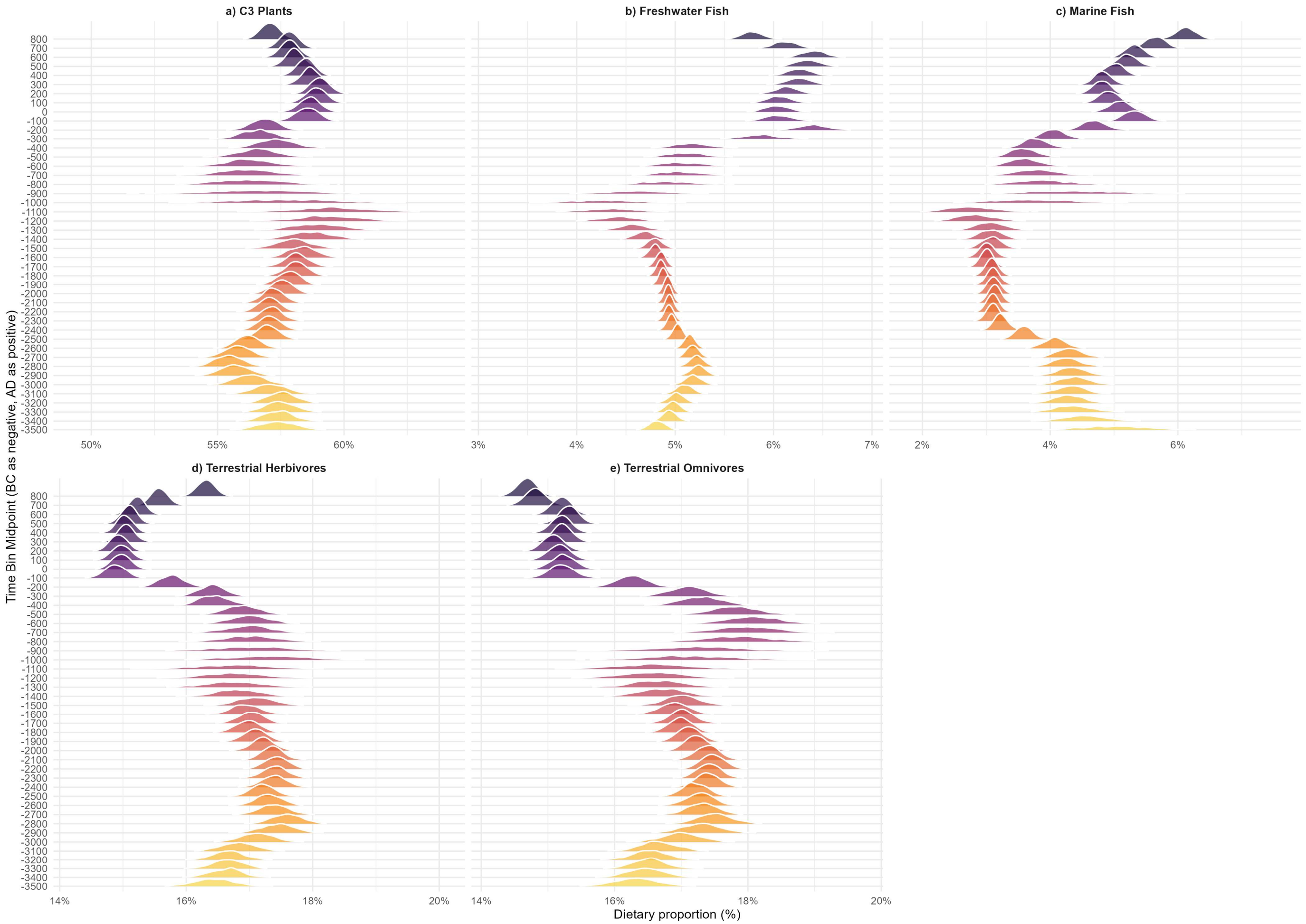

**Fig. S38| Probability density distribution of bootstrap dietary time-series models (causal discovery) by time bin. a),** $C_3$ plants. **b),** Freshwater fish. **c),** Marine fish. **d),** Terrestrial herbivores. **e),** Terrestrial omnivores.

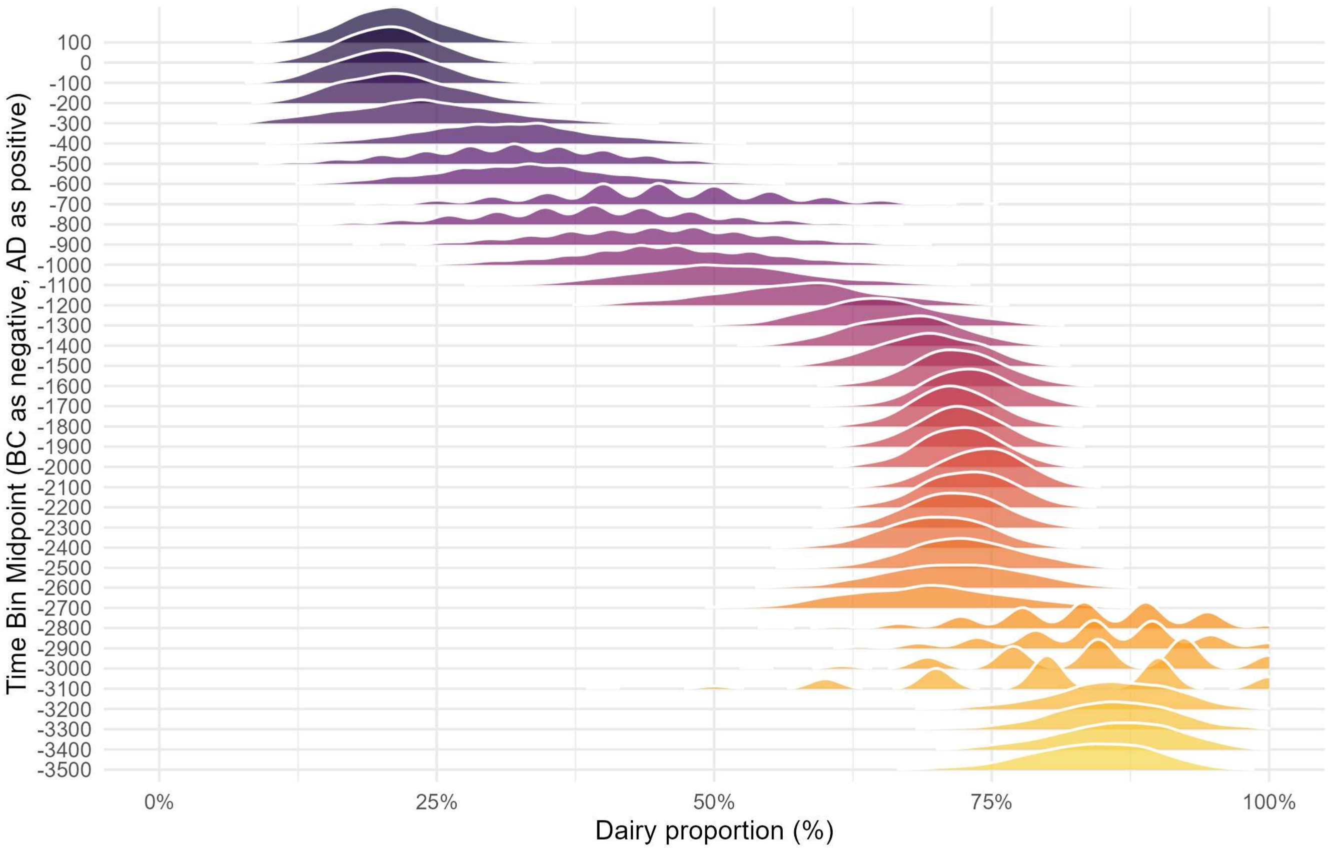

**Fig. S39| Probability density distribution of bootstrap dairy time-series models by time bin.**

#### Section 3 Allele frequency trajectories construction and time-varying selection coefficients estimation (Additional Notes)

This section provides supplementary information on the construction of allele frequency trajectories and the estimation of selection coefficients.

First, for each of the 14 SNPs, we generated 1,000 bootstrap replicates for each time slice and used them to construct the corresponding probability density distributions of derived allele frequencies. To evaluate the robustness of the bootstrap-derived allele frequency estimates, we here display the full probability distributions for all SNPs across all time intervals. The second part presents simulation-based analyses to evaluate the accuracy and reliability of our Generalized Additive Models (GAMs). We regard the parameter estimates as reliable if the allele frequency trajectories simulated under the fitted model can adequately capture the main trends of the observed trajectories. The third part addresses the reasons why certain SNPs yielded fewer estimates of time-varying selection coefficients and discusses which SNPs are unsuitable for further analysis.

##### 3A. Detailed results of bootstrap sampling for allele frequency trajectories

Here, we present detailed results from the bootstrap[115] resampling procedure. After assigning each allele to its corresponding time bin, we performed resampling with replacement within each bin. This process was repeated 1,000 times. This yielded 1,000 bootstrap estimates of the derived allele frequency per bin, from which the empirical frequency distributions were obtained. This procedure was applied to all 14 SNPs. According to the law of large numbers and the central limit theorem[116], when the number of resamples is sufficiently large, the resulting distributions tend to approximate a normal distribution.

The results clearly show that, for all 14 SNPs (Fig. S40), the majority of the bootstrap distributions are approximately normal in shape (585 of 616 bins; 95%), although a minority of bins display visibly multimodal distributions (Fig. S40d, e, k, l, n), which likely reflect the limited number of samples available in those time slices. Overall, these results support the robustness of the bootstrap estimates for 95% of the time bins and provide a transparent view of the uncertainty underlying our inferred allele frequency trajectories.

#### 3B. Simulation results of the fitted GAM model

We simulated allele frequency trajectories using the fitted GAM models and reduced models that retain only the linear terms of the GAM (i.e., trajectories reflecting solely the effect of natural selection), and compared these simulations with the observed allele frequency trajectories (Fig. S41). Except for rs3891176, rs6265, and rs7517 (Fig. S41g, j, l), the smooth terms in the GAMs played a substantial role, leading to pronounced differences between the full GAM models and the selection only (linear term) models. In contrast, for these three SNPs the smooth terms were effectively shrunk toward zero. This occurred because the data for these SNPs did not provide strong support for additional smooth structure, and the penalization framework, together with our allowance for smooth term shrinkage, appropriately suppressed the unnecessary complexity.

Compared with the observed frequency trajectories, the GAM models performed extremely well, with the exception of rs3891176 and rs7775397 (Fig. S41g, m). The GAM models successfully capture the complex nonlinear dynamics present in the data of the remaining 12 SNPs. The overall nonlinear trends can be captured very well. When the observed trajectory of an SNP is particularly smooth (e.g., rs653178), the GAM can even produce a simulated curve that closely matches the empirical data (Fig. S41k). Based on these findings, the selection coefficient estimates for the 12 well-fitted SNPs are deemed reasonable. As a result of trajectory inconsistencies, rs7775397 and rs3891176 are excluded from subsequent causal analyses.

#### 3C. Interpretation of time-varying selection coefficients for rs3891176, rs4988235, and rs7775397

The number of time-varying selection coefficient estimates is smaller for rs3891176, rs4988235, and rs7775397 compared to all the other SNPs (18, 35 and 13 versus 43), particularly for rs3891176 and rs7775397. One contributing factor is that the derived allele appears relatively late in the British dataset for all three SNPs: approximately 3600 BC for rs4988235, 2200 BC for rs3891176, and 1100 BC for rs7775397. In contrast, the derived alleles of all other SNPs had already emerged by 4000 BC.

Another key factor affecting these three SNPs is the prolonged sparsity of derived allele observations following their initial appearance. For example, in the case of rs4988235, the first derived allele appears around 3600 BC, but no additional occurrences are observed for the next 1,000 years. Since our time-varying selection coefficients are estimated using a sliding window approach, a window such as 3600–2600 BC (centered at 3100 BC) would exhibit a strongly negative selection coefficient. This is because the derived allele is only observed at the very beginning of the window and absent thereafter. This pattern does not reflect a true biological signal and thus renders such windows unreliable for estimating selection coefficients. Consequently, time intervals like these are excluded from the analysis. Similar issues are observed for rs3891176 and rs7775397, leading to fewer valid time-specific estimates for these three SNPs compared to the others.

Although rs4988235 has fewer time-varying selection coefficient estimates than the other SNPs (35 vs. 43), the difference is relatively minor, especially when compared to the much lower numbers observed for rs3891176 and rs7775397 (18 and 13 estimates, respectively). Moreover, our model validation indicates that the estimates for rs4988235 are nonetheless reliable (Fig. S41i). Therefore, rs4988235 is retained for causal analysis.

The time-varying selection coefficients of rs3891176 and rs7775397 were excluded from further causal testing. This decision was based on two primary considerations. First, the number of time intervals for which selection coefficients could be reliably estimated was limited for both SNPs. Such short time series result in insufficient library lengths for the convergent cross mapping (CCM) algorithm, potentially preventing convergence and leading to misleading causal results. Second, the fitted GAM simulations did not reproduce the observed trajectories well for these two SNPs (Fig. S41g, m). Therefore, these two SNPs were excluded from further analyses.

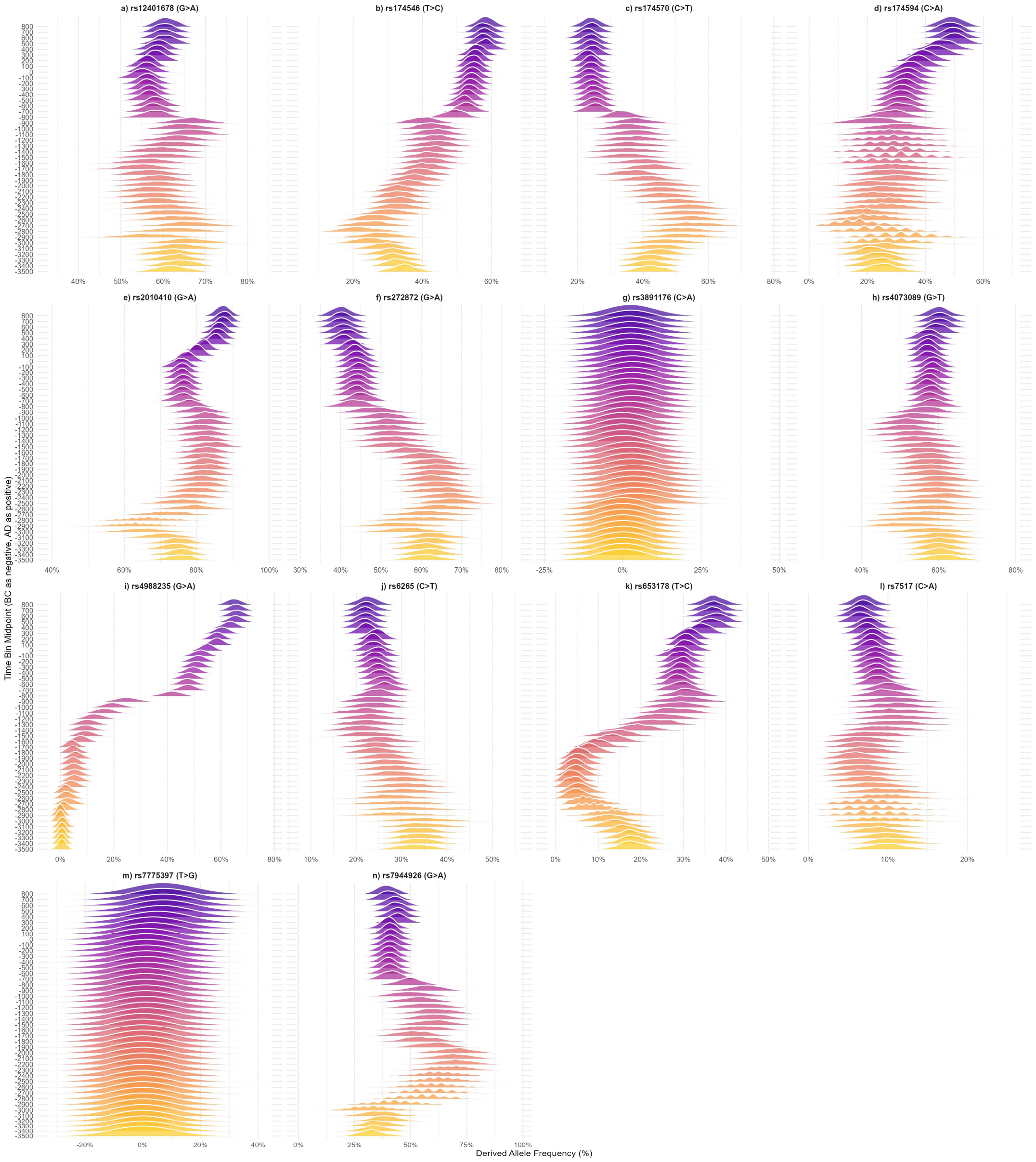

**Fig. S40| Probability density distribution of bootstrap-derived allele frequencies by time bin. a)-n),** Probability density distribution of bootstrap-derived allele frequencies for SNPs rs12401678 to rs7944926 (ordered left to right, top to bottom).

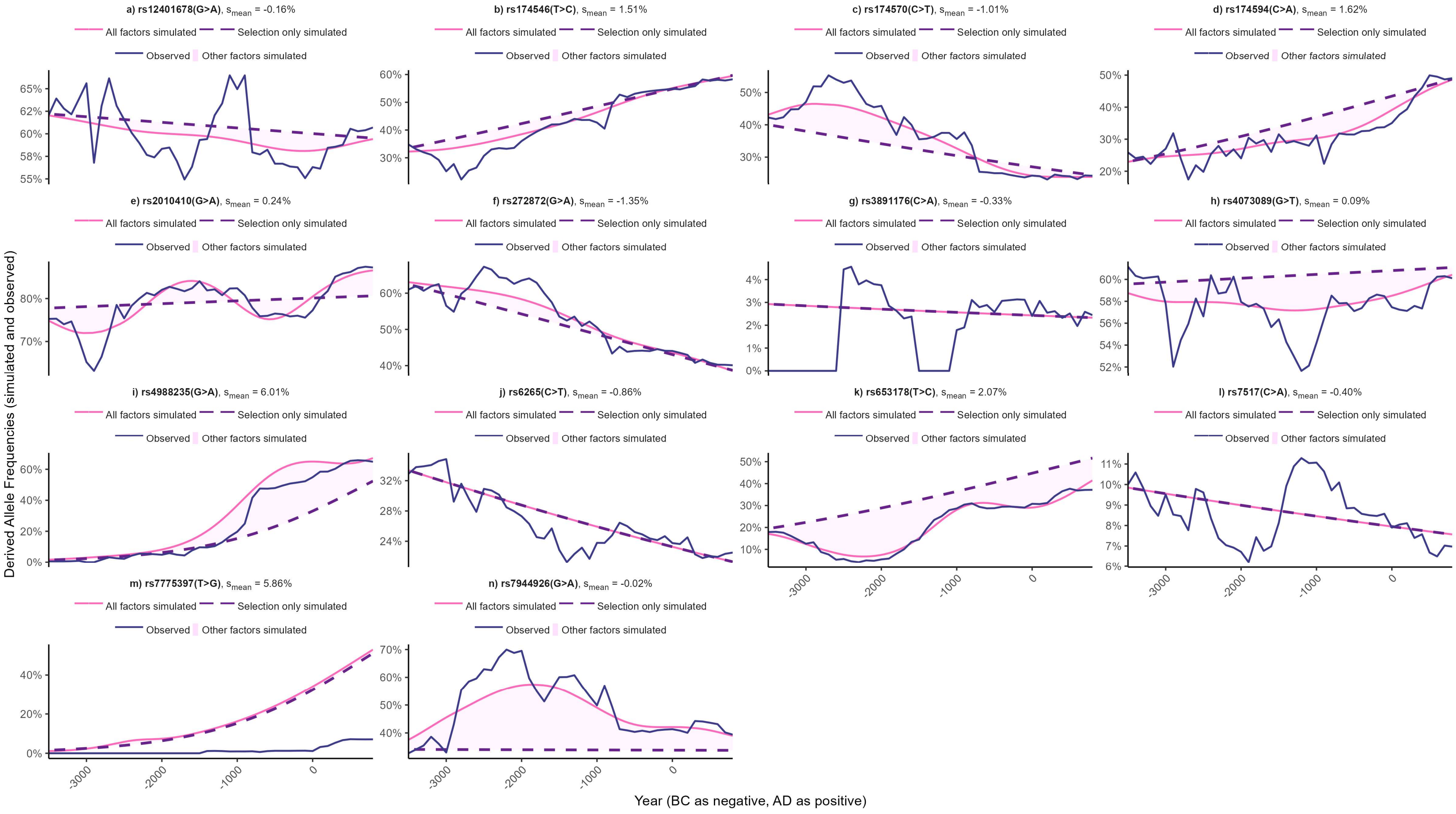
a) rs12401678(G>A), $s_{mean}$ = -0.16%
b) rs174546(T>C), $s_{mean}$ = 1.51%
c) rs174570(C>T), $s_{mean}$ = -1.01%
d) rs174594(C>A), $s_{mean}$ = 1.62%
e) rs2010410(G>A), $s_{mean}$ = 0.24%
f) rs272872(G>A), $s_{mean}$ = -1.35%
g) rs3891176(C>A), $s_{mean}$ = -0.33%
h) rs4073089(G>T), $s_{mean}$ = 0.09%
i) rs4988235(G>A), $s_{mean}$ = 6.01%
j) rs6265(C>T), $s_{mean}$ = -0.86%
k) rs653178(T>C), $s_{mean}$ = 2.07%
l) rs7517(C>A), $s_{mean}$ = -0.40%
m) rs7775397(T>G), $s_{mean}$ = 5.86%
n) rs7944926(G>A), $s_{mean}$ = -0.02%
All factors simulated
Selection only simulated
Observed
Other factors simulated
Derived Allele Frequencies (simulated and observed)
Year (BC as negative, AD as positive)

**Fig. S41| Simulated and observed derived allele frequency trajectories. a)-n),** Simulated and observed derived allele frequency trajectories for SNPs rs12401678 to rs7944926 (ordered left to right, top to bottom). The pink solid line represents simulations from the GAM model incorporating natural selection and other evolutionary forces, while the purple dashed line shows simulations retaining only the linear selection term. The light purple shaded area illustrates the difference between the two simulation models, reflecting the influence of non-selection factors on allele-frequency change. The dark blue solid line depicts the observed allele frequency trajectory.

### Section 4 Robustness tests of CCM, causal analysis review, and details of convergent cross mapping

In this section, we first present the robustness tests of CCM. We then review existing approaches to causal inference and causal discovery, explain our rationale for selecting convergent cross mapping (CCM) as the method of causal analysis for our study. The last part of this section presents the mathematical foundations and algorithmic structure of CCM in detail.

#### 4A. Simulation studies to test the robustness of the CCM results

Among the SNPs that exhibit clear causal signals, rs12401678 and rs653178 display very strong robustness, with 93% and 97% of the simulated CCM curves meeting the causal criterion (Fig. S42a, j). rs4988235 shows weaker robustness but still achieves the criterion in 63% of the simulations (Fig. S42h). For SNPs with possible causal signals, the results are also relatively robust: 95% of the simulated curves for rs174570 agree with its point-estimate CCM conclusion (Fig. S42c), and the corresponding proportion for rs174594 is 69% (Fig. S42d). Among the SNPs without causal signals, most exhibit strong robustness, with the simulated CCM curves broadly supporting the point-estimate results. However, almost half of the simulated CCM curves disagree with the original point-estimate conclusions for rs4073089 (50%) and rs7944926 (53%), indicating that the causal estimates for these two loci are less robust (Fig. S42g, l). Overall, SNPs that show (rs12401678, rs653178, and rs4988235), or are likely to show (rs174570 and rs174594), causal signals demonstrate acceptable robustness, even though not all of them exceed 90% simulation agreement. This indicates that our results are reasonably robust.

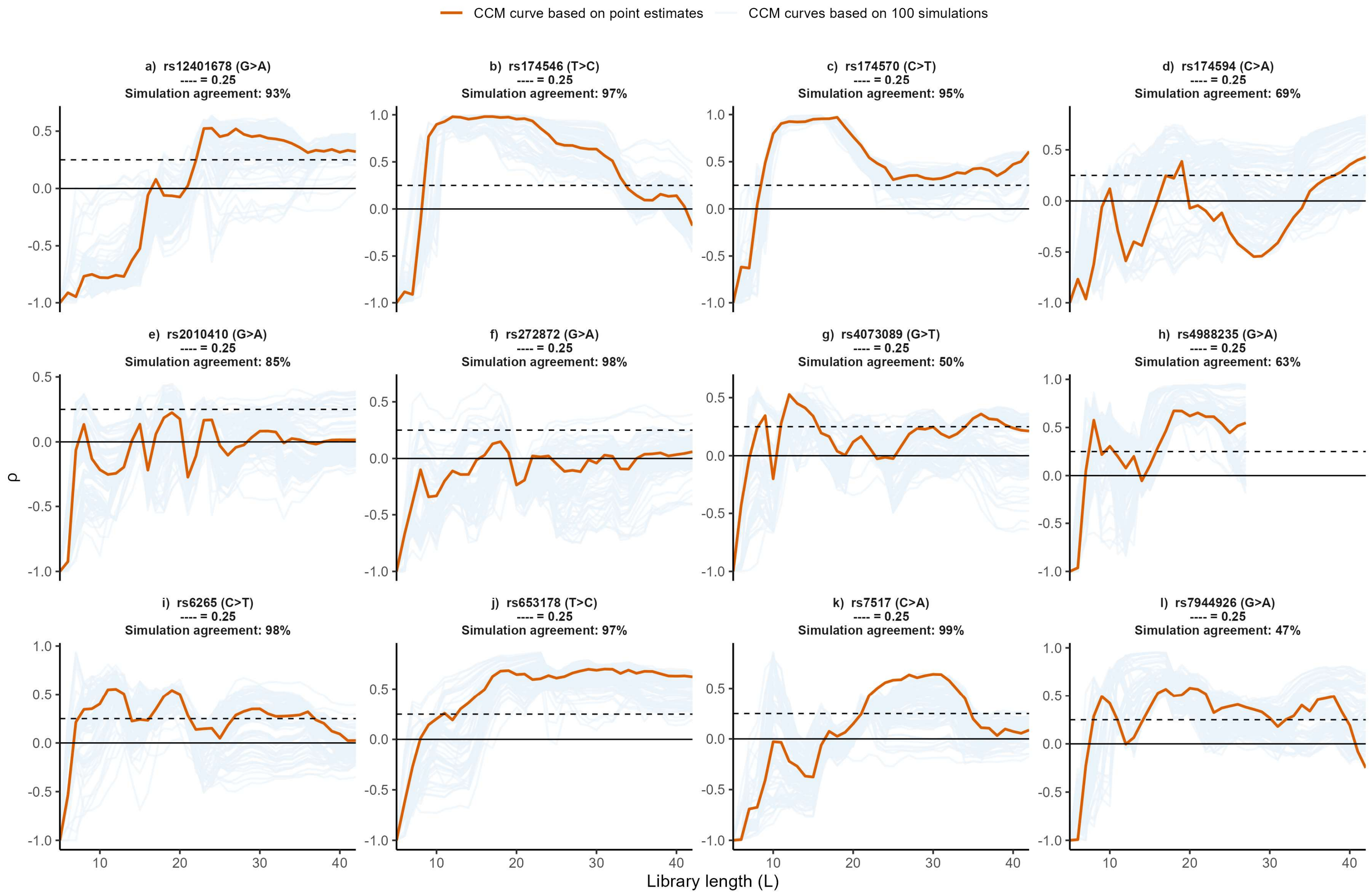

CCM curve based on point estimates
CCM curves based on 100 simulations
a) rs12401678 (G>A)
---- = 0.25
Simulation agreement: 93%
b) rs174546 (T>C)
---- = 0.25
Simulation agreement: 97%
c) rs174570 (C>T)
---- = 0.25
Simulation agreement: 95%
d) rs174594 (C>A)
---- = 0.25
Simulation agreement: 69%
e) rs2010410 (G>A)
---- = 0.25
Simulation agreement: 85%
f) rs272872 (G>A)
---- = 0.25
Simulation agreement: 98%
g) rs4073089 (G>T)
---- = 0.25
Simulation agreement: 50%
h) rs4988235 (G>A)
---- = 0.25
Simulation agreement: 63%
i) rs6265 (C>T)
---- = 0.25
Simulation agreement: 98%
j) rs653178 (T>C)
---- = 0.25
Simulation agreement: 97%
k) rs7517 (C>A)
---- = 0.25
Simulation agreement: 99%
l) rs7944926 (G>A)
---- = 0.25
Simulation agreement: 47%
ρ
Library length (L)

**Fig. S42| CCM robustness analysis across SNPs: CCM curves based on point-estimate time-series (red lines) and 100 CCM curves based on simulated time-series (light blue lines). a)-l),** CCM robustness tests for SNPs rs12401678 to rs7944926 (ordered left to right, top to bottom). Simulation agreement refers to the proportion of simulated CCM analyses that support the causal conclusion inferred from the point-estimate CCM.

#### 4B. A brief review of causal discovery and inference methods

Broadly speaking, causality methods fall into two categories: causal inference (i.e., counterfactual causal effect estimation) and causal discovery. Causal inference methods are predominantly used in econometrics[117–124] and epidemiology[125–129], while various causal discovery algorithms have been widely applied in earth sciences, ecology, and other data-intensive disciplines[130–140]. In this part, we begin by briefly reviewing causal inference and causal discovery approaches other than CCM and discussing why they are not suitable for our data. We then explain why CCM is appropriate for our study.

Causal inference

The fundamental idea of causal inference originates from randomized controlled trials (RCTs) and has developed rapidly following the introduction of the potential outcome framework by Donald Rubin[117,141–146]. At the individual level, a potential outcome represents the value that the outcome variable would assume if the value of a particular treatment variable were altered, assuming all other factors remain unchanged. Clearly, an individual's potential outcome is a counterfactual value and therefore cannot be directly observed or estimated. For instance, when evaluating the effect of taking medication on recovery time from a cold, if an individual takes the medication, we cannot observe what his recovery time would have been had they not taken it. Although the individual may experience another cold in the future and choose not to take the medication, their physiological condition and environmental context at that time would differ from the previous case. Hence, the key assumption of potential outcomes—holding all other factors constant while varying only the treatment—is violated.

Since it is impossible to infer the potential outcome at the individual level, causal inference typically focuses on estimating population-level effects, such as the Average Treatment Effect (ATE). But if we cannot identify an individual's counterfactual outcome, how can we possibly estimate the average of such outcomes across a population? This is where RCTs become essential. By randomly assigning subjects to a treatment group and a control group, and applying the treatment to the former only, we can estimate the ATE as the difference in average outcomes between the two groups. Due to the random assignment, the two groups are assumed to be exchangeable, meaning that they have the same average potential outcomes (i.e., no confounding). As a result, the average outcome in the control group can serve as an unbiased estimate of the potential outcome under no treatment for the treated group.

If the treatment and control groups are not randomly assigned, bias will arise due to confounding factors. For example, when assessing the effect of medication on recovery rates, suppose that the treatment group consists primarily of elderly individuals, while the control group consists of younger individuals. In such a case, one might erroneously conclude that the medication impairs recovery, simply because the treatment group (older individuals) recovers more slowly than the control group (younger ones). Here, age acts as a confounding variable, as it influences both the likelihood of receiving the treatment and the outcome of interest. The lack of random assignment violates the assumption of group comparability, leading to biased estimates of the causal effect. In statistics, this violation is referred to as a failure of the ignorability assumption[145], meaning that the treatment assignment is not independent of potential outcomes. In econometrics, the same issue is termed endogeneity[118,122], indicating that the explanatory variable is correlated with the error term, thereby biasing causal estimates.

However, RCTs are not always feasible in practice. In fields such as econometrics and epidemiology, where causal inference must often rely on observational data, researchers have developed a wide range of methods to adjust for confounding factors and obtain reliable estimates of causal effects from observational data.

In econometrics, commonly used methods for causal inference include instrumental variables (IV)[118,120,122], difference-in-differences (DiD)[119], regression discontinuity design (RDD)[121], and the synthetic control method[123,124]. In epidemiology, researchers have developed a distinct framework known as the G methods[125,126,128,129], including g formula, marginal structural models, and structural nested models. Additionally, Judea Pearl's causal diagram[127,130,133,147] offers a systematic approach to adjusting for confounding, through tools such as the back-door criterion, front-door criterion, and do-calculus. Although causal diagram provides a powerful and general framework in theory, they are less widely adopted among empirical researchers, due to the strong assumptions required for model specification.

The applicability of these causal inference methods is closely tied to the characteristics of the data in different disciplines. Unfortunately, none of these approaches is suitable for our study context. G methods typically require a large number of covariates, which are often readily available in epidemiological studies. However, for our analysis of ancient datasets, such comprehensive covariate information is essentially unattainable. Methods developed in econometrics also prove challenging for our case. Given the long temporal scope of our study, identifying valid IVs is virtually impossible, and RDD is clearly incompatible with our data structure. DiD appeared to be a potentially feasible option, but our attempts to apply it failed due to the violation of the parallel trends assumption. Given the inapplicability of causal inference methods, we are compelled to adopt causal discovery algorithms to test for causal relationships under the constraints of our data structure.

Causal discovery

Common causal discovery algorithms can be broadly categorized into four major classes[132]: Granger causality (GC)[134,140], the nonlinear state-space methods[135,139] (e.g., convergent cross mapping CCM); causal network learning algorithms (e.g., Peter and Clark momentary conditional independence PCMCI[137]), and structural causal models

(SCMs: e,g, LiNGAM[136] method and ANM method[138]). Unlike causal inference methods, all of which generally operate within a counterfactual framework, different causal discovery algorithms adopt distinct definitions of how a causal relationship is identified.

Causal network learning algorithms generally require a large number of high-quality covariates in order to construct robust and interpretable causal graphs. As such, they are not suitable for our context, where covariate data are severely limited due to the nature of ancient datasets. As for GC, we find its definition of causality conceptually problematic. The method posits that if a time series Y, which follows an autoregressive process, shows improved predictive accuracy when another variable X is included in the model, then X is said to "Granger-cause" Y. However, this prediction-based definition of causality diverges significantly from the notion of true causal mechanisms as commonly understood in empirical science. Among the remaining options, SCM-based algorithms appear even less applicable in our context. For instance, the ANM method relies on a particularly strong and often unrealistic assumption: that the effect variable can be expressed as a deterministic function of the cause variable plus a noise term that is statistically independent of the cause. The independence between the cause and the error term, effectively implying the absence of confounding, which is often highly unrealistic in practical settings.

Therefore, after careful consideration, we selected CCM as our primary method for causal analysis. While CCM is not without limitations—for instance, high levels of noise can affect its accuracy[131]—it remains the most practical and suitable approach given the structure of our data. Compared to other mainstream causal inference and discovery methods, CCM offers a more robust framework for uncovering causal signals in complex, nonlinear, and sparse time series data with no requirement for covariates. It also does not rely on smoothly varying or stationary time series, unlike Granger causality tests, making it applicable to a broader range of empirical datasets, such as ours. In addition, CCM is particularly well suited for detecting causality in weakly coupled time series. Excessively strong coupling between time series can cause CCM to lose the ability to distinguish the direction of causality, rendering the method ineffective. To confirm that our data meets this condition, we performed linear regressions between each SNP's selection coefficient trajectory and its corresponding dietary time series (Fig. S43 and Fig. S44). All pairs showed very low correlations, with $R^2$ values ranging from 0.001 to 0.313. This indicates that all SNP–diet combinations fall within the weak-coupling regime, fully satisfying the assumptions under which CCM is most appropriately applied.

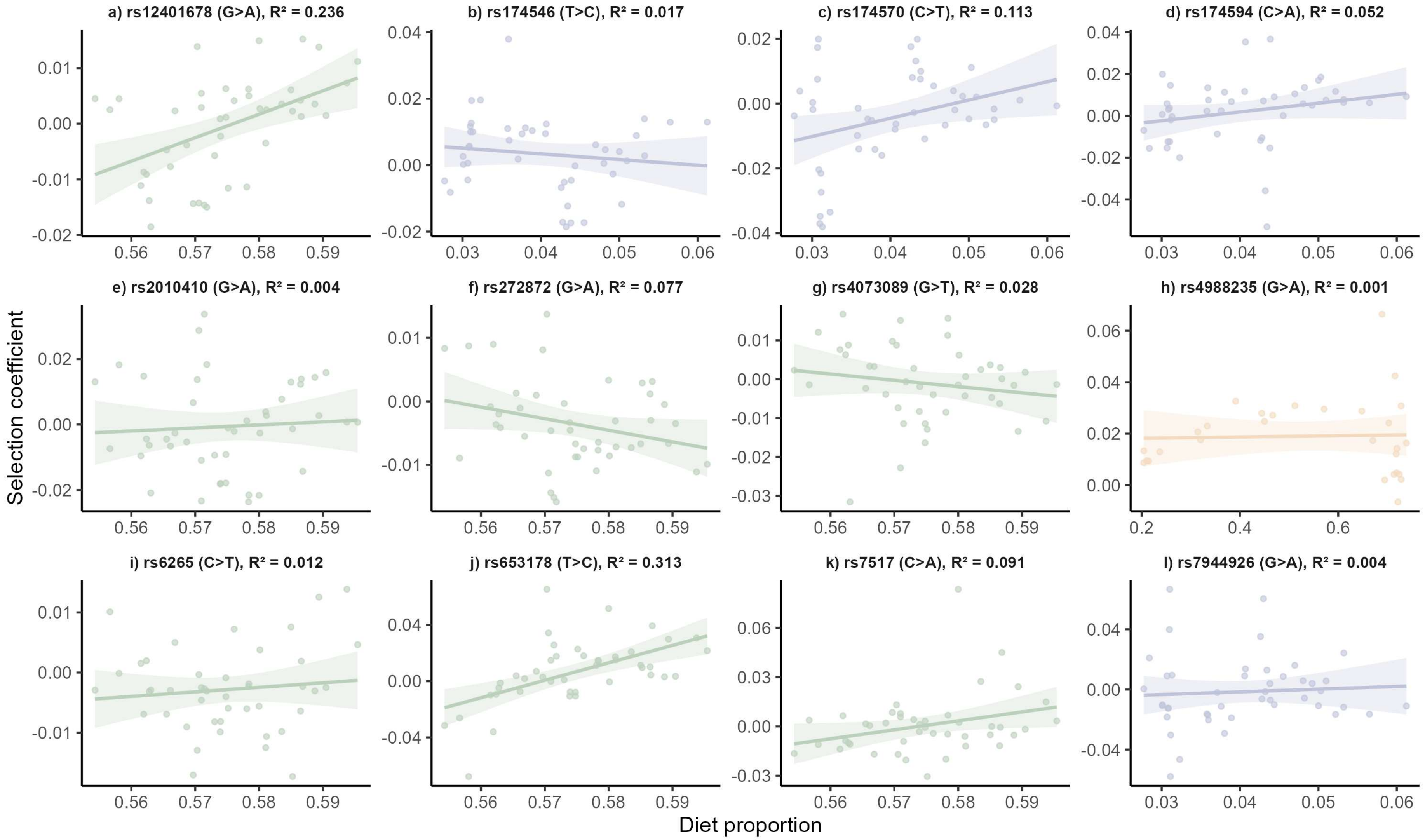
a) rs12401678 (G>A), R² = 0.236
b) rs174546 (T>C), R² = 0.017
c) rs174570 (C>T), R² = 0.113
d) rs174594 (C>A), R² = 0.052
e) rs2010410 (G>A), R² = 0.004
f) rs272872 (G>A), R² = 0.077
g) rs4073089 (G>T), R² = 0.028
h) rs4988235 (G>A), R² = 0.001
i) rs6265 (C>T), R² = 0.012
j) rs653178 (T>C), R² = 0.313
k) rs7517 (C>A), R² = 0.091
l) rs7944926 (G>A), R² = 0.004
Selection coefficient
Diet proportion

**Fig. S43| Linear correlation between diet and selection for each SNP. a)-l),** Linear correlation results for SNPs rs12401678 to rs7944926 (ordered left to right, top to bottom). SNPs associated with marine fish, $C_3$ plants, and dairy are highlighted in blue, green, and yellow, respectively. The linear fits are based on the point estimates of the time-varying selection coefficients and their corresponding point estimates of the dietary proportions at the same time slices.

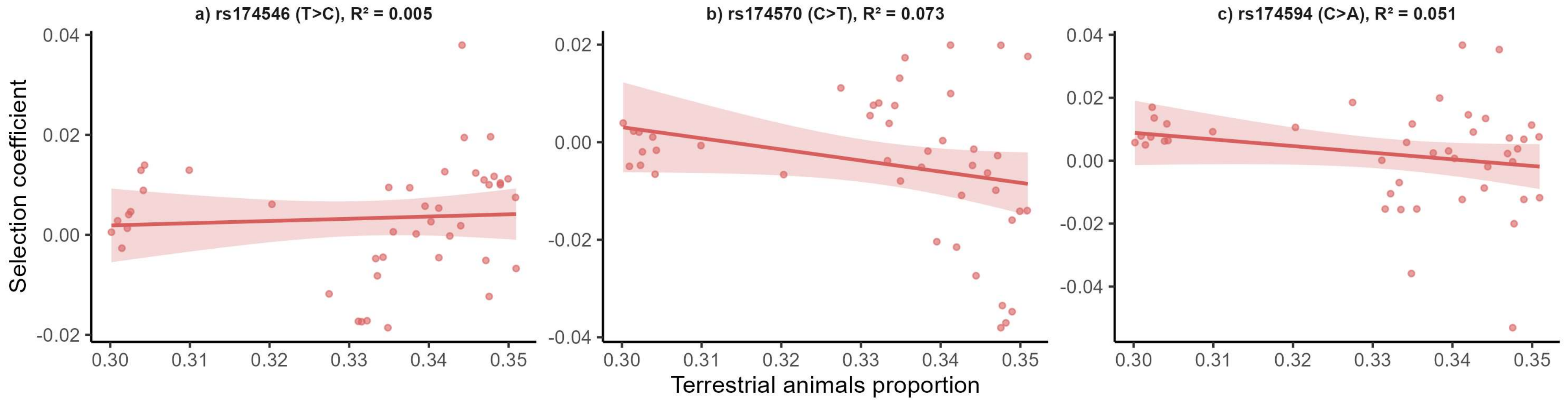

**Fig. S44| Linear correlation between diet and selection for three fatty acid metabolism SNPs (dietary driver: terrestrial animals = herbivores + omnivores). a),** rs174546. **b),** rs174570. **c),** rs174594. The linear fits are based on the point estimates of the time-varying selection coefficients and their corresponding point estimates of the dietary proportions at the same time slices.

#### 4C. Details of convergent cross mapping

The philosophy of CCM is that if the treatment variable X is a cause of the outcome variable Y, then Y must 'know' the behavior of X, and this information can be captured in the state space of Y (denoted as $M_Y$). This idea is based on the principle of state-space reconstruction in dynamical systems[148,149]. Therefore, even if the functional relationship between X and Y is unknown, CCM can still perform causal discovery by measuring the extent to which the state space of Y $(M_Y)$ can reliably estimate the values of X. If $M_Y$ accurately estimates X (i.e., ρ converges to a value significantly above zero), it implies that $M_Y$ contains the causal 'fingerprints' of X, indicating a causal relationship from X to Y. In our study, the outcome variables Y represent the selection coefficients of each SNP, whereas the treatment variables X correspond to the associated dietary variables.

We first performed state-space reconstruction of the outcome variable Y using its time-series data. This reconstruction was based on Takens' time-delay embedding theorem[148,149], which allows the dynamics of a system to be reconstructed from a sequence of scalar observations.

$$M_Y = \{(Y_t, Y_{t-\tau}, Y_{t-2\tau}, \dots, Y_{t-(E-1)\tau}), t = E, E+1, \dots, N\}. \tag{1}$$

$M_Y$ is the so-called shadow manifold of Y, reconstructed from its time-series data. Here, E denotes the embedding dimension, τ is the delay time, and N is the total number of time points in the time series. In our analysis, we set the embedding dimension to E=3 and the delay time to τ=1. The total number of time points is N=29 for rs4988235 and N=44 for all other SNPs.

Then, the reconstructed state space $M_Y$ was used to estimate the value of X. For a specific point $y_t = (Y_t, Y_{t-\tau}, Y_{t-2\tau})$ in $M_Y$, we identified its k nearest neighbors $y_i$. In our analysis, k was set to 4. Euclidean distance was used to calculate the distances between points in the state space.

$$distance(y_t, y_i) = \sqrt{\sum_{j=0}^{E-1} (y_{t-j\tau} - y_{i-j\tau})^2}, i = 1,2,3,4, E = 3, \tau = 1. \tag{2}$$

We then assigned a weight to each neighbor based on its distance to the target point.

$$\omega_i = \frac{exp(-distance(y_t, y_i))}{\sum_{m=1}^{k} exp(-distance(y_t, y_m))}, k = 4. \tag{3}$$

We did not use the normalized form $exp(\frac{-distanc\ (y_t,y_i)}{-distance\ (y_t,y_1)})$ to calculate weights, as is sometimes done to avoid numerical underflow when distances are large. In our case, all data points used to construct $M_Y$ are on a consistent scale, and the pairwise distances are uniformly small. Therefore, using the unnormalized form $exp(-distance\ (y_t, y_i))$ is sufficient and numerically stable.

We then estimated $\widehat{X_t}|M_Y$ and examined the convergence of the prediction.

$$\widehat{X_t}|M_Y = F(M_Y) = \sum_{i=1}^{k} \omega_i \cdot X_i \, , k = 4. \quad (4)$$

In practice, we started with a small library length L=5, using the first five points in $M_Y$ to estimate the corresponding X values: $X_3$ to $X_7$ (excluding the first two due to embedding constraints). We then gradually increased L one point at a time, up to the maximum length N−2. Each time L grows, the corresponding X is estimated. At each step, we calculated the Pearson correlation coefficient ρ between the predicted $\widehat{X_t}|M_Y$ and the true value $X_t$. If X is a true cause of Y, the correlation ρ is expected to increase and stabilize as L grows, ultimately converging to a value significantly greater than zero[139].

We explain the selection of the core CCM parameters as follows. The choice of $\tau$=1 reflects the fact that our time series are discretized into consecutive, equally spaced temporal bins, such that a unit delay represents the smallest meaningful temporal separation in the data. Because our time series are relatively short, larger values of $\tau$ would lead to a sparse reconstructed state space and increased noise in CCM predictions. Accordingly, we adopted $\tau$=1 to ensure that the reconstructed state space retains as many data points as possible. The embedding dimension was set to E=3. Given the limited length of our time series, a low-dimensional embedding should be adopted to ensure stable state-space reconstruction. Higher embedding dimensions would substantially reduce the effective number of embedded points and lead to sparse reconstructions. However, an overly low embedding dimension (e.g., E=2) may be insufficient to adequately reconstruct the underlying state space. Accordingly, we adopted E=3. This embedding dimension has been used in numerous CCM applications involving short time series[150]. Consistent with standard CCM settings[139], the number of nearest neighbors was set to k=E+1=4.

#### Section 5 Limitations

The limitations of this study fall broadly into three domains: dietary reconstruction, selection coefficient estimation, and causal discovery.

To begin with, our reconstruction of ancient diets cannot approach the level of precision available in modern biobank datasets. The dietary categories that can be modelled remain necessarily coarse, limited to five groups: terrestrial herbivores, omnivores, C3 plants, marine fish, and freshwater fish. Many other components of past diets, such as vegetables, mushrooms, or nuts, are invisible to current isotopic approaches and therefore cannot be modelled in any meaningful way. For meat consumption, we can only estimate overall intake and cannot differentiate between diets rich in fat and those based primarily on lean meat. These constraints impose clear limitations when linking reconstructed diets to genetic variants associated with complex traits such as obesity or circulating ergothioneine levels. For such SNPs, the key dietary signals may lie in components that cannot be reconstructed (e.g., vegetables and mushrooms). In addition, they may also depend on nutritional subtleties, including differences between fat and lean tissue consumption, which our model is unable to resolve.

Secondly, the precision of individual-level reconstructions and the uneven distribution of samples may also introduce biases. For example, in our reconstructed dietary time series, marine fish consumption appears extremely high in the Mesolithic period, higher than in the Later Medieval period. However, the number of high–marine-fish consumers in the Later Medieval period is greater than in the Mesolithic. The seemingly elevated Mesolithic values may therefore reflect the sparse sampling for this period, which could inflate the proportion of individuals with high marine fish consumption. This bias does not affect our causal discovery results, because the causal analyses do not include Mesolithic samples. In addition, we use time intervals of 1,000 years with a 100-year sliding step, which helps to smooth out biases arising from individual-level estimation errors and uneven temporal sampling. However, this approach also reduces the temporal variability of the reconstructed dietary time series, which may in turn make causal signals more difficult to detect.

The limitations of our selection coefficient estimates arise primarily from the sample size. We developed a GAM framework with the aim of letting the data speak. Over the past several millennia, allele frequency trajectories have been shaped by a wide range of factors, and it is neither feasible nor realistic to model all of them explicitly. Instead, we introduce a smooth term that is constructed to be uncorrelated with the linear term representing natural selection, allowing the model to capture additional structure in the data while preserving an interpretable selection component. However, although a dataset of more than 1,000 individuals is substantial by ancient DNA standards, it still represents only a tiny fraction of all the people who lived in Britain during this period. If the sampled individuals are not representative of the underlying populations, then the strategy of letting the data speak is inevitably constrained.

CCM is, among the available causal analysis approaches, the method best suited to the structure of our data. Nonetheless, it has several important limitations. It can evaluate causal relationships only between one potential driver and one outcome variable at a time, and its performance is sensitive to noise. As a result, SNPs associated with complex traits, such as those related to type 2 diabetes, do not yield detectable causal signals. Similarly, SNPs influenced by multiple dietary components, such as those involved in lipid metabolism (affected by both marine fish and terrestrial meat consumption), can only be identified as having possible causal signals (i.e., their $\rho$ values at maximum library length exceed 0.25, but the $\rho$ trajectories remain non-convergent).

Overall, by integrating multiple ancient datasets and applying a suite of interdisciplinary methods, we provide an initial attempt to address the long-standing evolutionary question of whether diet has driven natural selection in humans. Our findings highlight the complexity of this topic and indicate that finer-grained work will be needed to further understand the mechanisms involved.

#### Section 6 Supplementary methods

##### 6A. Data collection for isotopic data

Detailed information for each sample—including sample ID, location (region, site name, latitude, and longitude), dating (archaeological period, specific dating results, and dating methods), material used for isotopic analysis (e.g., bone or dentine), isotopic values ($\delta^{13}C$, $\delta^{15}N$), quality control metrics (C/N ratio, C, N, collagen yield), and relevant references—is provided in Supplementary Data 1. For human samples, information on sex and age is also included.

Regarding sample chronology, calibrated radiocarbon dates were used as the primary dating source whenever available. If only uncalibrated radiocarbon dates were reported, these were retained. For samples lacking radiocarbon data, site phase information was used. In cases where the site phase was vague or imprecise, we assigned a date range based on archaeological context (e.g., "late 5th to early 7th century" was defined as 480–620 CE).

For several Paleolithic and Mesolithic individuals from Doggerland[14,15], a now-submerged landmass that once connected southeastern Britain to continental Europe, we randomly selected a coordinate within the Strait of Dover to represent their location. We assigned the same coordinate to all samples to reflect their likely geographic association during this period. The geographic coordinates of the samples from the Beaker People Project and the Beakers and Bodies Project[45,46] were randomly generated within twelve geographic coordinate ranges defined by their respective location groups. These coordinates do not represent the exact locations of each sample, but they do reflect their general geographic distribution.

When constructing the time series, isotopic measurements from multiple skeletal elements of the same individual, including bone, dentine, and incremental dentine samples, were generally treated as separate data points rather than being averaged. This decision was made because our focus is on reconstructing broader dietary patterns across specific time slices in Great Britain. Multiple samples from the same individual are neither entirely independent nor true replicates. Treating them as distinct observations increases the effective sample size and allows for a more robust estimation of mean dietary trends, although it may lead to a slight underestimation of population-level variance. For the purposes of this study, the mean trend is of primary interest, whereas precise estimation of population variance is of secondary importance. For these reasons, we adopted this strategy. Exceptions were made for samples from Later Medieval southern England and Post-Medieval northern England, where incremental dentine samples were excluded from the time-series modeling to avoid a few sites dominating the dataset (see Section 2H and I for details).

In our modeling, we excluded individuals under the age of three as well as those clearly identified as non-local in the original publications. Individuals younger than three years old may rely on a mixture of breastfeeding and solid foods[151], and therefore their isotopic values cannot be used to infer dietary intake in the same way as older individuals. Similarly, the isotopic signatures of confirmed migrants likely reflect food consumption from other regions and therefore cannot be considered representative of diets in Britain. Samples that may have been influenced by breastfeeding—such as first permanent molars, deciduous teeth, or incremental samples representing diets from ages 0 to 3—were also excluded from the analysis.

If a sample is clearly affected by manuring and exhibits anomalous isotopic values, as identified in its original publication, a correction is applied to its $\delta^{15}N$ value by subtracting a fixed offset (1.0‰) before isotopic modelling. Such samples generally show modestly higher $\delta^{15}N$ values than those from nearby contemporaneous sites, without a corresponding increase in $\delta^{13}C$. The magnitude of the enrichment (c. 1.0‰) is substantially smaller than that associated with marine fish consumption.

Individuals with poor data quality (C/N ratios outside the accepted range: 2.9–3.6)[152] were also excluded from the analysis.

After these exclusions, the final set of samples is used for modeling. The priors assigned to each sample (see Methods for the criteria) and the resulting posterior estimates are presented in Supplementary Data 3.

#### 6B. Isotopic values of food resources

Constructing individual-level dietary models to infer proportional dietary contributions ($\theta_i$) using Bayesian estimation[153] requires isotopic data from both the individual ($H_k^{obs}$) and locally accessible food sources ($I_{ijk}$), together with prior information on other model parameters ($W_{jk}, C_{ij}, \theta_i, T_k$). Each accessible food group typically contains multiple samples; the isotopic values within each group are averaged, and standard deviations are calculated. These summary statistics are then used to define the prior distributions for the corresponding model parameters ($I_{ijk}$).

Here we focus on five food resource groups which are isotopically distinguishable: terrestrial herbivores (2359 samples), terrestrial omnivores (860 samples), $C_3$ plants (206 samples), marine resources (280 samples), and freshwater resources (50 samples). Ideally the accessible food source data are contemporary with the individual and from the same region, but such a dataset is rarely available in archaeology. Although we compiled nearly all available isotopic data for ancient British humans and food sources, it was impossible to obtain local isotopic data for all relevant food groups corresponding precisely to each human sample. For most archaeological sites, isotopic data for certain food sources (e.g., terrestrial herbivores and omnivores) are available, while data for others (e.g., marine or freshwater resources) are often missing.

To address this limitation and ensure feasibility, we relaxed the spatial and temporal constraints by aggregating the data into broader region-period groupings. Specifically, we divided Britain into three geographic regions: England (including samples from Doggerland and the Isle of Man), Wales and Scotland. Temporally, the data were divided into eight broad periods:

- Paleolithic (before 9000 BC)
- Mesolithic (9000–4300 BC)
- Neolithic (4300–2200 BC)
- Bronze Age (2200–750 BC)
- Iron Age (750 BC–AD 43)
- Roman/Roman Iron Age (AD 43–410)
- Early Medieval (AD 410–1066)
- Later Medieval/Post-Medieval (AD 1066–1900)

For terrestrial herbivores and terrestrial omnivores, each sample was assigned to a specific region-period combined group based on its latitude, longitude, and chronological information (e.g., England, Iron Age). In cases where a food resource sample spanned multiple periods (e.g., AD 1 to 99, which overlaps with both the Iron Age and Roman period/Roman Iron Age), it was included in both relevant groups. The food resource data from each region-period combined group were used as the accessible food resources for the human individuals associated with that group.

For $C_3$ plants, all plant samples were classified solely by geographic region (England, Wales, or Scotland), irrespective of the archaeological period for the post-agricultural era (i.e., from the Neolithic onward). Plant data from England were used for the dietary reconstruction of human samples from this region across all periods, with the same approach applied to Wales and Scotland. However, we applied correction factors to account for potential manuring effects in later agricultural contexts. Since all our plant samples derive from the agricultural period, their $\delta^{15}N$ values are higher than those of herbivores and omnivores from the Paleolithic and Mesolithic periods. Therefore, for the Paleolithic and Mesolithic, plant $\delta^{15}N$ values were adjusted downward by 4‰ to reflect the absence of manuring practices, ensuring consistency with the expected trophic level differences between plants and herbivores. Additionally, to account for temporal variations in atmospheric $CO_2$ $\delta^{13}C$ values[154], plant $\delta^{13}C$ values for the Paleolithic and Mesolithic periods were adjusted upward by +2‰ and +1‰, respectively. Our data on $\delta^{13}C$ values from terrestrial herbivores and omnivores show minimal variation from the Neolithic through to the Post-Medieval period, with notable differences observed only in the Palaeolithic and Mesolithic (Supplementary Data 2).

There are no published data for ancient $C_3$ plants in Scotland. Therefore, we used modern $C_3$ plant grain samples from Scotland as a substitute. However, it was necessary to correct the $\delta^{13}C$ values to account for changes in atmospheric $CO_2$ over the past century[154–156]. A correction of +4.55‰ was applied to all modern C3 plant samples, based on our data analysis comparing modern and ancient isotopic values of plants between England and Scotland (Table S2). Although some previous studies have suggested a correction of +1.9‰[157], our analyses deemed this value to be too conservative.

For marine resources and freshwater resources, the food source isotopic data were common to all human samples (Table S3 summarizes the correspondence between human individuals and their accessible food resource groups).

The correspondence between human individuals and food resource groups was determined based on detailed analyses of isotopic data from the food groups. However, these results have not yet been published. In this Supplementary Information, any citation of this unpublished manuscript will be indicated using this designation: (Chen et al in prep).

The isotopic values of food resources are typically measured from their bone collagen. However, our Bayesian mixing model requires the isotopic values of the protein and energy fractions. Following the recommendations of previous studies[158,159], we calculated the isotopic values of these fractions using the following equations (see also Supplementary Data 2):

Terrestrial herbivores and omnivores:

$$\delta^{13}C_{Protein} = \delta^{13}C_{Collagen} - 2‰;$$
$$\delta^{15}N_{Protein} = \delta^{15}N_{Collagen} + 2‰;$$
$$\delta^{13}C_{Energy} = \delta^{13}C_{Collagen} - 8‰.$$

C3 plant:

$$\delta^{13}C_{Prote} = \delta^{13}C_{Bulk} - 2‰;$$
$$\delta^{15}N_{Prote} = \delta^{15}N_{Bulk};$$
$$\delta^{13}C_{Energy} = \delta^{13}C_{Bulk} + 0.5‰.$$

Marine fish or freshwater fish:

$$\delta^{13}C_{Protein} = \delta^{13}C_{Collagen} - 1‰;$$
$$\delta^{15}N_{Protein} = \delta^{15}N_{Collagen} + 2‰;$$
$$\delta^{13}C_{Ene} = \delta^{13}C_{Collagen} - 7‰.$$

During the conversion of isotopic values from collagen/dentine to dietary fractions, an additional 0.5 is added to the standard deviations, except in the case of $\delta^{15}N$ for $C_3$ plants, as this value remains untransformed.

**Table S2| The comparison of $\delta^{13}C$ values of ancient and modern plants from England and Scotland**

| $\delta^{13}C$ | Modern Scotland plant biomass (150 samples) | Modern England plant biomass (224 samples) | Modern Scotland grain (20 samples) | Ancient England grain (161 samples) |
|---|---|---|---|---|
| Mean | -29.60‰ | -30.80‰ | -27.20‰ | -23.15‰ |
| SD | 1.59‰ | 2.37‰ | 0.42‰ | 1.17‰ |

We possessed isotopic data for modern plant biomass from both Scotland and England, as well as for modern grain from Scotland. Our aim was to correct the $\delta^{13}C$ values of modern Scottish grain to their ancient equivalents.

An offset of 1.20‰ was observed between modern plant biomass from England and Scotland ($\delta^{13}C$ values of -29.60‰ and -30.80‰, respectively). Based on the $\delta^{13}C$ value of modern Scottish grain (-27.20‰), we assigned a corresponding virtual $\delta^{13}C$ value of -28.40‰ to modern English grain. This virtual value was determined by applying the same biomass offset to the grain data.

Comparing the estimated modern $\delta^{13}C$ value for English grain (-28.40‰) with the $\delta^{13}C$ values of ancient English grains (-23.15‰) reveals an offset of 5.25‰. If we apply this 5.25‰ offset as a correction factor, the mean $\delta^{13}C$ value of ancient Scottish grain would be approximately -21.95‰, slightly higher than that of ancient English grain (-23.15‰). However, after reviewing $\delta^{13}C$ values of herbivores from Neolithic to the Post-Medieval period for Scotland (-21.77‰ to -21.11‰) and England (-22.15‰ to -21.62‰), we ultimately determined to set the $\delta^{13}C$ value for ancient Scottish grain at -22.65‰, corresponding to a correction factor of +4.55‰.

**Table S3| The correspondence between human individuals and their accessible food resource groups used for Bayesian estimation (some groups, such as Bronze Age Wales, are absent because no human samples are available for those groups.)**

| Human | Herbivores | Omnivores | $C_3$ Plants | Marine resources | Freshwater resources |
|---|---|---|---|---|---|
| Paleolithic (England) | Paleolithic (England) | Paleolithic (England) | England (corrected) | All | All |
| Mesolithic (England) | Mesolithic (England) | Mesolithic (England) | England (corrected) | All | All |
| Neolithic (England) | Neolithic (England) | Neolithic (England) | England | All | All |
| Bronze Age (England) | Bronze Age (England) | Bronze Age (England) | England | All | All |
| Iron Age (England) | Iron Age (England) | Iron Age (England) | England | All | All |
| Roman (England) | Roman (England) | Roman (England) | England | All | All |
| Early Medieval (England) | Early Medieval (England) | Early Medieval (England) | England | All | All |
| Later/Post-Medieval (England) | Later/Post-Medieval (England) | Later/Post-Medieval (England) | England | All | All |
| Paleolithic (Wales) | Paleolithic (Wales) | Paleolithic (Wales) | Wales (corrected) | All | All |
| Mesolithic (Wales) | Mesolithic (Wales) | Mesolithic (Wales) | Wales (corrected) | All | All |
| Neolithic (Wales) | Neolithic (Wales) | Neolithic (Wales) | Wales | All | All |
| Early Medieval (Wales) | Early Medieval (Wales) | Early Medieval (Wales) | Wales | All | All |
| Neolithic (Scotland) | Neolithic (Scotland) | Neolithic (Scotland) | Scotland (converted) | All | All |
| Bronze Age (Scotland) | Bronze Age (Scotland) | Bronze Age (Scotland) | Scotland (converted) | All | All |
| Iron Age (Scotland) | Iron Age (Scotland) | Iron Age (Scotland) | Scotland (converted) | All | All |
| Roman Iron Age (Scotland) | Roman Iron Age (Scotland) | Roman Iron Age (Scotland) | Scotland (converted) | All | All |
| Early Medieval (Scotland) | Early Medieval (Scotland) | Early Medieval (Scotland) | Scotland (converted) | All | All |
| Later/Post-Medieval (Scotland) | Later/Post-Medieval (Scotland) | Later/Post-Medieval (Scotland) | Scotland (converted) | All | All |

#### References

1. Le, M. K. *et al.* 1,000 ancient genomes uncover 10,000 years of natural selection in Europe. *bioRxiv* https://doi.org/10.1101/2022.08.24.505188 (2022) doi:10.1101/2022.08.24.505188.
2. Akbari, A. *et al.* Pervasive findings of directional selection realize the promise of ancient DNA to elucidate human adaptation. *bioRxiv* https://doi.org/10.1101/2024.09.14.613021 (2024) doi:10.1101/2024.09.14.613021.
3. Mathieson, I. *et al.* Genome-wide patterns of selection in 230 ancient Eurasians. *Nature* 528, 499–503 (2015).
4. Little, M., Humphries, S., Patel, K. & Dewey, C. Factors associated with BMI, underweight, overweight, and obesity among adults in a population of rural south India: A crosssectional study. *BMC Obes.* 3, 12 (2016).
5. Heald, A. *et al.* A study to investigate genetic factors associated with weight gain in people with diabetes: analysis of polymorphisms in four relevant genes. *Adipocyte* 12, 2236757 (2023).
6. Tian, X., Thorne, J. L. & Moore, J. B. Ergothioneine: An underrecognised dietary micronutrient required for healthy ageing? *British Journal of Nutrition* 129, 104–114 (2023).
7. Evershed, R. P. *et al.* Dairying, diseases and the evolution of lactase persistence in Europe. *Nature* 608, 336–345 (2022).
8. Ye, K., Gao, F., Wang, D., Bar-Yosef, O. & Keinan, A. Dietary adaptation of FADS genes in Europe varied across time and geography. *Nat. Ecol. Evol.* 1, 0167 (2017).
9. Durmuş, M. Fish oil for human health: Omega-3 fatty acid profiles of marine seafood species. *Food Science and Technology (Brazil)* 39, 454–461 (2019).
10. Saini, R. K. & Keum, Y. S. Omega-3 and omega-6 polyunsaturated fatty acids: Dietary sources, metabolism, and significance — A review. *Life Sci.* 203, 255–267 (2018).
11. Fumagalli, M. *et al.* Greenlandic Inuit show genetic signatures of diet and climate adaptation. *Science (1979).* 349, 1343–1347 (2015).
12. Maurya, V. K. & Aggarwal, M. Factors influencing the absorption of vitamin D in GIT: an overview. *J. Food Sci. Technol.* 54, 3753–3765 (2017).
13. Zgaga, L. *et al.* Diet, environmental factors, and lifestyle underlie the high prevalence of vitamin D deficiency in healthy adults in Scotland, and supplementation reduces the proportion that are severely deficient. *Journal of Nutrition* 141, 1535–1542 (2011).
14. van der Plicht, J., Amkreutz, L. W. S. W., Niekus, M. J. L. T., Peeters, J. H. M. & Smit, B. I. Surf'n Turf in Doggerland: Dating, stable isotopes and diet of Mesolithic human remains from the southern North Sea. *J. Archaeol. Sci. Rep.*

10, 110–118 (2016).
15. Amkreutz, L. *et al.* What lies beneath . . . Late Glacial human occupation of the submerged North Sea landscape. *Antiquity* 92, 22–37 (2018).
16. Stevens, R. E., Jacobi, R. M. & Higham, T. F. G. Reassessing the diet of Upper Palaeolithic humans from Gough's Cave and Sun Hole, Cheddar Gorge, Somerset, UK. *J. Archaeol. Sci.* 37, 52–61 (2010).
17. Jacobi, R. M. & Higham, T. F. G. The early Lateglacial re-colonization of Britain: new radiocarbon evidence from Gough's Cave, southwest England. *Quat. Sci. Rev.* 28, 1895–1913 (2009).
18. Richards, M. P., Hedges, R. E. M., Jacobi, R., Current, A. & Stringer, C. FOCUS: Gough's Cave and Sun Hole Cave Human Stable Isotope Values Indicate a High Animal Protein Diet in the British Upper Palaeolithic. *J. Archaeol. Sci.* 27, 1–3 (2000).
19. Richards, M. P., Jacobi, R., Cook, J., Pettitt, P. B. & Stringer, C. B. Isotope evidence for the intensive use of marine foods by Late Upper Palaeolithic humans. *J. Hum. Evol.* 49, 390–394 (2005).
20. Richards, M. P., Pettitt, P. B., Stiner, M. C. & Trinkaus, E. Stable isotope evidence for increasing dietary breadth in the European mid-Upper Paleolithic. *Proc. Natl. Acad. Sci. U. S. A.* 98, 6528–6532 (2001).
21. Schulting, R. J. *et al.* AVELINES'S HOLE: AN UNEXPECTED TWIST IN THE TALE. *Proceedings of the University of Bristol Spelaeological Society* 28, 9–63 (2019).
22. Schulting, R. 'Tilbury man': A mesolithic skeleton from the lower thames. *Proceedings of the Prehistoric Society* 79, 19–37 (2013).
23. Meiklejohn, C., Chamberlain, A. T. & Schulting, R. J. Radiocarbon dating of Mesolithic human remains in Great Britain. *Mesolithic Miscellany* 21, 20–58 (2011).
24. Schulting, R. J., Gardiner, P. J., Hawkes, C. J. & Murray, E. THE MESOLITHIC AND NEOLITHIC HUMAN BONE ASSEMBLAGE FROM TOTTY POT, CHEDDAR, SOMERSET. *Proceedings of the University of Bristol Spelaeological Society* 25, 75–95 (2010).
25. Richards, M. Chapter 12 Human consumption of plant foods in the British Neolithic: Direct evidence from bone stable isotopes. in *Plants in Neolithic Britain and Beyond* (ed. Fairburn, A. S.) vol. 5 123–136 (Oxbow Books, Oxford, 2000).
26. Brunning, R. An Early Mesolithic Cemetery at Greylake, Somerset, UK. *Archaeology in the Severn Estuary* 22, 67–70 (2013).
27. Schulting, R. J. & Richards, M. P. Finding the coastal Mesolithic in southwest Britain: AMS dates and stable isotope results on human remains from Caldey Island, South Wales. *Antiquity* 76, 1011–1025 (2002).
28. Schulting, R. *et al.* Mesolithic and neolithic human remains from Foxhole Cave, Gower, South Wales. *The Antiquaries Journal* 93, 1–23 (2013).

29. Bownes, J. Reassessing the Scottish Mesolithic-Neolithic Transition: Questions of diet and chronology. (University of Glasgow, Glasgow, 2018).
30. Armit, I., Shapland, F., Montgomery, J. & Beaumont, J. Difference in Death? A Lost Neolithic Inhumation Cemetery with Britain's Earliest Case of Rickets, at Balevullin, Western Scotland. *Proceedings of the Prehistoric Society* 81, 199–214 (2015).
31. Montgomery, J. *et al.* Strategic and sporadic marine consumption at the onset of the Neolithic: Increasing temporal resolution in the isotope evidence. *Antiquity* 87, 1060–1072 (2013).
32. Lawrence, D. M. Orkney's first farmers. Reconstructing biographies from osteological analysis to gain insights into life and society in a Neolithic community on the edge of Atlantic Europe. (University of Bradford, 2012).
33. Schulting, R., Sheridan, A., Crozier, R. & Murphy, E. Revisiting Quanterness: new AMS dates and stable isotope data from an Orcadian chamber tomb. *Proceedings of the Society of Antiquaries of Scotland* 140, 1–50 (2010).
34. Gigleux, C., Richards, M. P., Curtis, N., Hutchison, M. & Britton, K. Reconstructing diet at the Neolithic stalled cairn of the Knowe of Rowiegar, Rousay, Orkney, using stable isotope analysis. *J. Archaeol. Sci. Rep.* 13, 272–280 (2017).
35. Charlton, S. *et al.* Finding Britain's last hunter-gatherers: A new biomolecular approach to 'unidentifiable' bone fragments utilising bone collagen. *J. Archaeol. Sci.* 73, 55–61 (2016).
36. Schulting, R. J. & Richards, M. P. The Wet, the Wild and the Domesticated: the Mesolithic-- Neolithic Transition On the West Coast of Scotland. *Eur. J. Archaeol.* 5, 147–189 (2002).
37. Stevens, R. E., Lightfoot, E., Allen, T. & Hedges, R. E. M. Palaeodiet at Eton College Rowing Course, Buckinghamshire: Isotopic changes in human diet in the Neolithic, Bronze Age, Iron Age and Roman periods throughout the British Isles. *Archaeol. Anthropol. Sci.* 4, 167–184 (2012).
38. Hedges, R. E. M., Stevens, R. E. & Pearson, J. A. Carbon and Nitrogen Stable Isotope Compositions of Animal and Human Bone. in *Building Memories: The Neolithic Cotswold Long Barrow at Ascott-under-Wychwood, Oxfordshire* (eds. Benson, D. & Whittle, A.) 239–246 (Oxbow Books, Oxford, 2007).
39. Hedges, R., Saville, A. & O'Connell, T. Characterizing the diet of individuals at the Neolithic chambered tomb of Hazleton North, Gloucestershire, England, using stable isotopic analysis. *Archaeometry* 50, 114–128 (2008).
40. Schulting, R. J., Chapman, M. & Chapman, E. J. AMS 14 C DATING AND STABLE ISOTOPE (CARBON, NITROGEN) ANALYSIS OF AN EARLIER NEOLITHIC HUMAN SKELETAL ASSEMBLAGE FROM HAY WOOD CAVE, MENDIP, SOMERSET. *Proceedings of the University of Bristol Spelaeological Society* 26, 9–26 (2013).
41. Richards, M. 7 Human remains and diet: Stable isotope values. in *HAMBLEDON*

*HILL, DORSET, ENGLAND. Excavation and survey of a Neolithic monument complex and its surrounding landscape* (eds. Mercer, R. & Healy, F.) vol. 2 522–527 (Liverpool University Press, Historic England, Liverpool, 2008).

42. Armit, I., Schulting, R., Knusel, C. J. & Shepherd, I. A. G. Death, decapitation and display? The bronze and iron age human remains from the sculptor's cave, covesea, North-east Scotland. *Proceedings of the Prehistoric Society* 77, 251–278 (2011).
43. Pearson, M. P. *et al.* Evidence for mummification in Bronze Age Britain. *Antiquity* 79, 529–546 (2005).
44. Lightfoot, E. *et al.* An investigation into diet at the site of yarnton, oxfordshire, using stable carbon and nitrogen isotopes. *Oxford Journal of Archaeology* 28, 301–322 (2009).
45. Parker Pearson, M. *et al.* Beaker people in Britain: migration, mobility and diet. *Antiquity* 90, 620–637 (2016).
46. Jay, M. & Richards, M. P. Carbon and nitrogen isotopic analysis. in *The Beaker People: Isotopes, Mobility and Diet in Prehistoric Britain* (eds. Mike Parker Pearson et al.) 303–339 (Oxbow Books, Oxford, 2019).
47. Woodbridge, J. *et al.* The impact of the Neolithic agricultural transition in Britain: A comparison of pollen-based land-cover and archaeological 14C date-inferred population change. *J. Archaeol. Sci.* 51, 216–224 (2014).
48. Stevens, C. J. & Fuller, D. Q. Did neolithic farming fail? The case for a Bronze Age agricultural revolution in the British Isles. *Antiquity* 86, 707–722 (2012).
49. Treasure, E. R., Gröcke, D. R., Caseldine, A. E. & Church, M. J. Neolithic farming and wild plant exploitation in western Britain: Archaeobotanical and crop stable isotope evidence from Wales (c. 4000–2200 cal BC). *Proceedings of the Prehistoric Society* 85, 193–222 (2019).
50. Bishop, R. R. Did Late Neolithic farming fail or flourish? A Scottish perspective on the evidence for Late Neolithic arable cultivation in the British Isles. *World Archaeol.* 47, 834–855 (2015).
51. Jay, M. & Richards, M. P. British iron age diet: Stable isotopes and other evidence. *Proceedings of the Prehistoric Society* 73, 169–190 (2007).
52. Moore, J. *et al.* A multi-isotope (C, N, O, Sr, Pb) study of Iron Age and Roman period skeletons from east Edinburgh, Scotland exploring the relationship between decapitation burials and geographical origins. *J. Archaeol. Sci. Rep.* 29, 102075 (2020).
53. Jay, M. & Richards, M. P. Diet in the Iron Age cemetery population at Wetwang Slack, East Yorkshire, UK: Carbon and nitrogen stable isotope evidence. *J. Archaeol. Sci.* 33, 653–662 (2006).
54. Stevens, R. E., Lightfoot, E., Hamilton, J., Cunliffe, B. & Hedges, R. E. M. Investigating dietary variation with burial ritual in iron age hampshire: An isotopic comparison of suddern farm cemetery and danebury hillfort pit burials. *Oxford Journal of Archaeology* 32, 257–273 (2013).

55. Stevens, R. E., Lightfoot, E., Hamilton, J., Cunliffe, B. & Hedges, R. E. M. Stable Isotope Investigations Of The Danebury Hillfort Pit Burials. *Oxford Journal of Archaeology* 29, 407–428 (2010).
56. Pollard, A. M. *et al.* 'These boots were made for walking': The isotopic analysis of a C4 Roman inhumation from Gravesend, Kent, UK. *Am. J. Phys. Anthropol.* 146, 446–456 (2011).
57. Redfern, R. C., Millard, A. R. & Hamlin, C. A regional investigation of subadult dietary patterns and health in late Iron Age and Roman Dorset, England. *J. Archaeol. Sci.* 39, 1249–1259 (2012).
58. Jay, M. Iron Age Diet at Glastonbury Lake Village: The isotopic evidence for negligible aquatic resource consumption. *Oxford Journal of Archaeology* 27, 201–216 (2008).
59. Redfern, R. C., Hamlin, C. & Athfield, N. B. Temporal changes in diet: A stable isotope analysis of late Iron Age and Roman Dorset, Britain. *J. Archaeol. Sci.* 37, 1149–1160 (2010).
60. Müldner, G. & Richards, M. P. Stable isotope evidence for 1500 years of human diet at the city of York, UK. *Am. J. Phys. Anthropol.* 133, 682–697 (2007).
61. Eckardt, H., Müldner, G. & Speed, G. The Late Roman Field Army in Northern Britain Mobility, Material Culture and Multi-Isotope Analysis at Scorton (N Yorks.). *Britannia* 46, 191–223 (2015).
62. Müldner, G., Chenery, C. & Eckardt, H. The 'Headless Romans': Multi-isotope investigations of an unusual burial ground from Roman Britain. *J. Archaeol. Sci.* 38, 280–290 (2011).
63. Chenery, C., Eckardt, H. & Müldner, G. Cosmopolitan Catterick? Isotopic evidence for population mobility on Rome's Northern frontier. *J. Archaeol. Sci.* 38, 1525–1536 (2011).
64. Müldner, G. Stable isotopes and diet: Their contribution to Romano-British research. *Antiquity* 87, 137–149 (2013).
65. Bonsall, L. A. & Pickard, C. Stable isotope and dental pathology evidence for diet in late Roman Winchester, England. *J. Archaeol. Sci. Rep.* 2, 128–140 (2015).
66. Cheung, C., Schroeder, H. & Hedges, R. E. M. Diet, social differentiation and cultural change in Roman Britain: New isotopic evidence from Gloucestershire. *Archaeol. Anthropol. Sci.* 4, 61–73 (2012).
67. Redfern, R., Gowland, R., Millard, A., Powell, L. & Gröcke, D. 'From the mouths of babes': A subadult dietary stable isotope perspective on Roman London (Londinium). *J. Archaeol. Sci. Rep.* 19, 1030–1040 (2018).
68. Redfern, R. C. *et al.* Going south of the river: A multidisciplinary analysis of ancestry, mobility and diet in a population from Roman Southwark, London. *J. Archaeol. Sci.* 74, 11–22 (2016).
69. Sakai, Y. Transition from the Late Roman Period to the Early Anglo-Saxon Period in the Upper Thames Valley Based on Stable Isotopes. (University of Oxford, Oxford, 2017).

70. Chenery, C., Müldner, G., Evans, J., Eckardt, H. & Lewis, M. Strontium and stable isotope evidence for diet and mobility in Roman Gloucester, UK. *J. Archaeol. Sci.* 37, 150–163 (2010).
71. Nehlich, O. *et al.* Application of sulphur isotope ratios to examine weaning patterns and freshwater fish consumption in Roman Oxfordshire, UK. *Geochim. Cosmochim. Acta* 75, 4963–4977 (2011).
72. Richards, M. P., Hedges, R. E. M., Molleson, T. I. & Vogel, J. C. Stable Isotope Analysis Reveals Variations in Human Diet at the Poundbury Camp Cemetery Site. *J. Archaeol. Sci.* 25, 1247–1252 (1998).
73. Cummings, C. & Hedges, R. Chapter 5: Human remains: Carbon and nitrogen stable isotope analysis. in *The late Roman cemetery at Lankhills, Winchester: Excavations 2000-2005* (eds. Booth, P. et al.) vol. 10 411–421 (Oxford Archaeology Monograph, Oxford, 2010).
74. Fuller, B. T., Molleson, T. I., Harris, D. A., Gilmour, L. T. & Hedges, R. E. M. Isotopic evidence for breastfeeding and possible adult dietary differences from Late/Sub-Roman Britain. *Am. J. Phys. Anthropol.* 129, 45–54 (2006).
75. Barrett, J. H. & Richards, M. P. Identity, gender, religion and economy: New isotope and radiocarbon evidence for marine resource intensification in early historic Orkney, Scotland, UK. *Eur. J. Archaeol.* 7, 249–271 (2004).
76. Curtis-Summers, S., Pearson, J. A. & Lamb, A. L. From Picts to Parish: Stable isotope evidence of dietary change at medieval Portmahomack, Scotland. *J. Archaeol. Sci. Rep.* 31, 102303 (2020).
77. Lamb, A. L., Melikian, M., Ives, R. & Evans, J. Multi-isotope analysis of the population of the lost medieval village of Auldhame, East Lothian, Scotland. *J. Anal. At. Spectrom.* 27, 765–777 (2012).
78. Curtis-Summers, S., Montgomery, J. & Carver, M. Stable isotope evidence for dietary contrast between Pictish and medieval populations at Portmahomack, Scotland. *Mediev. Archaeol.* 58, 21–43 (2014).
79. Richards, M. P., Fuller, B. T. & Molleson, T. I. Stable isotope palaeodietary study of humans and fauna from the multi-period (Iron Age, Viking and Late Medieval) site of Newark Bay, Orkney. *J. Archaeol. Sci.* 33, 122–131 (2006).
80. Macpherson, P. M. Tracing Change: An Isotopic Investigation of Anglo-Saxon Childhood Diet. (University of Sheffield, Sheffield, 2005).
81. Moore, F. E. Diet and subsistence in the Anglo-Saxon Trent Valley: a stable isotope investigation of Broughton Lodge Anglo-Saxon cemetery, Nottinghamshire. (University of Nottingham, Nottingham, 2017).
82. Haydock, H., Clarke, L., Craig-Atkins, E., Howcroft, R. & Buckberry, J. Weaning at Anglo-Saxon raunds: Implications for changing breastfeeding practice in britain over two millennia. *Am. J. Phys. Anthropol.* 151, 604–612 (2013).
83. Jarman, C. L., Biddle, M., Higham, T. & Bronk Ramsey, C. The Viking Great Army in England: new dates from the Repton charnel. *Antiquity* 92, 183–199

(2018).
84. Hemer, K. A., Lamb, A. L., Chenery, C. A. & Evans, J. A. A multi-isotope investigation of diet and subsistence amongst island and mainland populations from early medieval western Britain. *Am. J. Phys. Anthropol.* 162, 423–440 (2017).
85. Mays, S. & Beavan, N. An investigation of diet in early Anglo-Saxon England using carbon and nitrogen stable isotope analysis of human bone collagen. *J. Archaeol. Sci.* 39, 867–874 (2012).
86. Buckberry, J. *et al.* Finding Vikings in the Danelaw. *Oxford Journal of Archaeology* 33, 413–434 (2014).
87. Pollard, A. M. *et al.* 'Sprouting like cockle amongst the wheat': The St Brice's Day Massacre and the isotopic analysis of human bones from St John's College, Oxford. *Oxford Journal of Archaeology* 31, 83–102 (2012).
88. Chenery, C. A., Evans, J. A., Score, D. & Boyle, A. A Boat Load of Vikings? *Journal of the North Atlantic* 7, 43–53 (2014).
89. Hannah, E. L., McLaughlin, T. R., Keaveney, E. M. & Hakenbeck, S. E. Anglo-Saxon diet in the Conversion period: A comparative isotopic study using carbon and nitrogen. *J. Archaeol. Sci. Rep.* 19, 24–34 (2018).
90. Knapp, Z. The Zooarchaeology of the Anglo-Saxon Christian Conversion: Lyminge, a case study. (University of Reading, Reading, 2018).
91. O'Connell, T. C. & Lawler, A. Chapter 5. Economic Resources VI. Stable isotope analysis of human and faunal remains. in *The Anglo-Saxon Settlement and Cemetery at Bloodmoor Hill, Carlton Colville, Suffolk* (eds. Lucy, S., Tipper, J. & Dickens, A.) vol. EAA 131 317–321 (East Anglian Archaeology, Barnsley, 2009).
92. Lucy, S. *et al.* The Burial of A Princess? The Later Seventh-Century Cemetery At Westfield Farm, Ely. *The Antiquaries Journal* 89, 81–141 (2009).
93. Privat, K. L., O'connell, T. C. & Richards, M. P. Stable isotope analysis of human and faunal remains from the Anglo-Saxon Cemetery and Berinsfield, Oxfordshire: Dietary and social implications. *J. Archaeol. Sci.* 29, 779–790 (2002).
94. Hemer, K. A., Lamb, A. L., Chenery, C. A. & Evans, J. A. A multi-isotope investigation of diet and subsistence amongst island and mainland populations from early medieval western Britain. *Am. J. Phys. Anthropol.* 162, 423–440 (2017).
95. Britton, K. *et al.* Isotopes and new norms: Investigating the emergence of early modern U.K. breastfeeding practices at St. Nicholas Kirk, Aberdeen. *Int. J. Osteoarchaeol.* 28, 510–522 (2018).
96. Müldner, G. *et al.* Isotopes and individuals: Diet and mobility among the medieval Bishops of Whithorn. *Antiquity* 83, 1119–1133 (2009).
97. Kancle, L., Montgomery, J., Gröcke, D. R. & Caffell, A. From field to fish: Tracking changes in diet on entry to two medieval friaries in northern England.

*J. Archaeol. Sci. Rep.* 22, 264–284 (2018).
98. Britton, K., Fuller, B. T., Tütken, T., Mays, S. & Richards, M. P. Oxygen isotope analysis of human bone phosphate evidences weaning age in archaeological populations. *Am. J. Phys. Anthropol.* 157, 226–241 (2015).
99. Bownes, J., Clarke, L. & Buckberry, J. The importance of animal baselines: Using isotope analysis to compare diet in a British medieval hospital and lay population. *J. Archaeol. Sci. Rep.* 17, 103–110 (2018).
100. Müldner, G. & Richards, M. P. Fast or feast: Reconstructing diet in later medieval England by stable isotope analysis. *J. Archaeol. Sci.* 32, 39–48 (2005).
101. Burt, N. M. Stable isotope ratio analysis of breastfeeding and weaning practices of children from medieval Fishergate House York, UK. *Am. J. Phys. Anthropol.* 152, 407–416 (2013).
102. Lamb, A. L., Evans, J. E., Buckley, R. & Appleby, J. Multi-isotope analysis demonstrates significant lifestyle changes in King Richard III. *J. Archaeol. Sci.* 50, 559–565 (2014).
103. Craig-Atkins, E. *et al.* The dietary impact of the Norman Conquest: A multiproxy archaeological investigation of Oxford, UK. *PLoS One* 15, e0239640 (2020).
104. Halldórsdóttir, H. H. *et al.* Continuity and individuality in Medieval Hereford, England: A stable isotope approach to bulk bone and incremental dentine. *J. Archaeol. Sci. Rep.* 23, 800–809 (2019).
105. Walter, B. S., DeWitte, S. N., Dupras, T. & Beaumont, J. Assessment of nutritional stress in famine burials using stable isotope analysis. *Am. J. Phys. Anthropol.* 172, 214–226 (2020).
106. Millard, A. R. *et al.* Scottish soldiers from the Battle of Dunbar 1650: A prosopographical approach to a skeletal assemblage. *PLoS One* 15, e0243369 (2020).
107. Bell, L. S., Lee Thorp, J. A. & Elkerton, A. The sinking of the Mary Rose warship: a medieval mystery solved? *J. Archaeol. Sci.* 36, 166–173 (2009).
108. Bleasdale, M. *et al.* Multidisciplinary investigations of the diets of two post-medieval populations from London using stable isotopes and microdebris analysis. *Archaeol. Anthropol. Sci.* 11, 6161–6181 (2019).
109. Roberts, P. *et al.* The men of Nelson's navy: A comparative stable isotope dietary study of late 18th century and early 19th century servicemen from Royal Naval Hospital burial grounds at Plymouth and Gosport, England. *Am. J. Phys. Anthropol.* 148, 1–10 (2012).
110. Dhaliwal, K., Rando, C., Reade, H., Jourdan, A. L. & Stevens, R. E. Socioeconomic differences in diet: An isotopic examination of post-Medieval Chichester, West Sussex. *Am. J. Phys. Anthropol.* 171, 584–597 (2020).
111. Beaumont, J. *et al.* Victims and survivors: Stable isotopes used to identify migrants from the Great Irish Famine to 19th century London. *Am. J. Phys. Anthropol.* 150, 87–98 (2013).
112. Nitsch, E. K., Humphrey, L. T. & Hedges, R. E. M. The effect of parity status on

δ15N: looking for the ‘ pregnancy effect’ in 18th and 19th century London. *J. Archaeol. Sci.* 37, 3191–3199 (2010).

113. Nitsch, E. K., Humphrey, L. T. & Hedges, R. E. M. Using stable isotope analysis to examine the effect of economic change on breastfeeding practices in Spitalfields, London, UK. *Am. J. Phys. Anthropol.* 146, 619–628 (2011).
114. Scorrer, J. *et al.* Diversity aboard a Tudor warship: Investigating the origins of the Mary Rose crew using multi-isotope analysis. *R. Soc. Open Sci.* 8, 202106 (2021).
115. Efron, B. Bootstrap Methods: Another Look at the Jackknife. *The Annals of Statistics* 7, 1–26 (1979).
116. Dekking, F. M., Kraaikamp, C., Lopuhaa, H. P. & Meester, L. E. *Modern Introduction to Probability and Statistics*. (Springer-Verlag, London, 2005).
117. Imbens, G. & Rubin, D. B. *Causal Inference for Statistics, Social, and Biomedical Sciences: An Introduction*. (Cambridge University Press, Cambridge, 2015).
118. Angrist, J. D., Imbens, G. W. & Rubin, D. B. Identification of Causal Effects Using Instrumental Variables. *J. Am. Stat. Assoc.* 91, 444–455 (1996).
119. Card, D. & Krueger, A. B. Minimum Wages and Employment: A Case Study of the Fast-Food Industry in New Jersey and Pennsylvania. *Am. Econ. Rev.* 84, 772–793 (1994).
120. Imbens, G. W. & Angrist, J. D. Identification and Estimation of Local Average Treatment Effects. *Econometrica* 62, 467–475 (1994).
121. Imbens, G. W. & Lemieux, T. Regression discontinuity designs: A guide to practice. *J. Econom.* 142, 615–635 (2006).
122. Angrist, J. D. & Pischke, J.-S. *Mostly Harmless Econometrics: An Empiricist's Companion*. (Princeton University Press, Princeton, NJ, 2008).
123. Abadie, A. & Gardeazabal, J. The Economic Costs of Confict: A Case Study of the Basque Country. *Am. Econ. Rev.* 93, 113–132 (2003).
124. Abadie, A., Diamond, A. & Hainmueller, A. J. Synthetic control methods for comparative case studies: Estimating the effect of California's Tobacco control program. *J. Am. Stat. Assoc.* 105, 493–505 (2010).
125. Robins, J. M., Ngel Hernán, M. A. ´ & Brumback, B. Marginal Structural Models and Causal Inference in Epidemiology. *Epidemiology* 11, 550–560 (2000).
126. Robins, J. A NEW APPROACH TO CAUSAL INFERENCE IN MORTALITY STUDIES WITH A SUSTAINED EXPOSURE PERIOD-APPLICATION TO CONTROL OF THE HEALTHY WORKER SURVIVOR EFFECT. *Mathematical Modelling* 7, 1393–1512 (1986).
127. Greenland, S., Pearl, J. & Robins, J. M. Causal Diagrams for Epidemiologic Research. *Epidemiology* 10, 37–48 (1999).
128. Hernán, M. A., Hernández-Díaz, S. & Robins, J. M. A structural approach to selection bias. *Epidemiology* 15, 615–625 (2004).
129. Hernán, M. A. & Robins, J. M. *Causal Inference: What If.* (Chapman &

Hall/CRC, 2024).
130. Pearl, J. *Causality: Models, Reasoning and Inference*. (Cambridge University Press, Cambridge, 2009).
131. Mønster, D., Fusaroli, R., Tylén, K., Roepstorff, A. & Sherson, J. F. Causal inference from noisy time-series data — Testing the Convergent Cross-Mapping algorithm in the presence of noise and external influence. *Future Generation Computer Systems* 73, 52–62 (2017).
132. Runge, J. *et al.* Inferring causation from time series in Earth system sciences. *Nat. Commun.* 10, 2553 (2019).
133. Pearl, Judea. *Causal Inference in Statistics : A Primer*. (John Wiley & Sons Ltd, 2016).
134. Shojaie, A. & Fox, E. B. Granger Causality: A Review and Recent Advances. *Annu. Rev. Stat. Appl.* 9, 289–319 (2022).
135. Ying, X. *et al.* Continuity Scaling: A Rigorous Framework for Detecting and Quantifying Causality Accurately. *Research* 2022, 9870149 (2022).
136. Shimizu, S., Hoyer, P. O. & Hyvärinen, A. A Linear Non-Gaussian Acyclic Model for Causal Discovery Antti Kerminen. *Journal of Machine Learning Research* 7, 2003–2030 (2006).
137. Runge, J., Nowack, P., Kretschmer, M., Flaxman, S. & Sejdinovic, D. Detecting and quantifying causal associations in large nonlinear time series datasets. *Sci. Adv* 5, eaau4996 (2019).
138. Hoyer, P. O., Janzing, D., Mooij, J., Peters, J. & Schölkopf, B. Nonlinear causal discovery with additive noise models. in *Advances in Neural Information Processing Systems* (Curran Associates, Inc., 2008).
139. Sugihara, G. *et al.* Detecting causality in complex ecosystems. *Science (1979).* 338, 496–500 (2012).
140. Granger, C. W. J. Investigating Causal Relations by Econometric Models and Cross-spectral Methods. *Econometrica* 37, 424–438 (1969).
141. Rubin, D. B. Practical Implications of Modes of Statistical Inference for Causal Effects and the Critical Role of the Assignment Mechanism. *Biometrics* 47, 1213–1234 (1991).
142. Rubin, D. B. Comment: Neyman (1923) and Causal Inference in Experiments and and Observational Studies. *Statistical Science* 5, 472–480 (1990).
143. Rubin, D. B. Bayesian Inference for Causal Effects: The Role of Randomization. *The Annals of Statistics* 6, 34–58 (1978).
144. Rubin, D. B. ESTIMATING CAUSAL EFFECTS OF TREATMENTS IN RANDOMIZED AND NONRANDOMIZED STUDIES. *J. Educ. Psychol.* 66, 688–701 (1974).
145. Rosenbaum, P. R. & Rubin, D. B. The central role of the propensity score in observational studies for causal effects. *Biometrika* 70, 41–55 (1983).
146. Basu, D. Randomization Analysis of Experimental Data: The Fisher Randomization Test. *J. Am. Stat. Assoc.* 75, 575–582 (1980).

147. Pearl, J. Causal diagrams for empirical research. *Biometrika* 82, 669–710 (1995).
148. Takens, F. Detecting strange attractors in turbulence. in *Dynamical Systems and Turbulence, Warwick 1980* (eds. Rand, D. & Young, L.-S.) 366–381 (Springer-Verlag Berlin Heidelberg, 1981).
149. Kostelich, E. J. & Schreiber, T. Noise reduction in chaotic time-series data: A survey of common methods. *PHYSICAL REVIEW E VOLUME* 48, 1752–1763 (1993).
150. Clark, A. T. *et al.* Spatial convergent cross mapping to detect causal relationships from short time series. *Ecology* 96, 1174–1181 (2015).
151. Kendall, E., Millard, A. & Beaumont, J. The "weanling's dilemma" revisited: Evolving bodies of evidence and the problem of infant paleodietary interpretation. *Am. J. Phys. Anthropol.* 175, 57–78 (2021).
152. Deniro, M. J. Postmortem preservation and alteration of in vivo bone collagen isotope ratios in relation to palaeodietary reconstruction. *Nature* 317, 806–809 (1985).
153. Fernandes, R., Millard, A. R., Brabec, M., Nadeau, M. J. & Grootes, P. Food reconstruction using isotopic transferred signals (FRUITS): A bayesian model for diet reconstruction. *PLoS One* 9, (2014).
154. Hare, V. J., Loftus, E., Jeffrey, A. & Ramsey, C. B. Atmospheric CO2 effect on stable carbon isotope composition of terrestrial fossil archives. *Nat. Commun.* 9, 252 (2018).
155. Eggleston, S., Schmitt, J., Bereiter, B., Schneider, R. & Fischer, H. Evolution of the stable carbon isotope composition of atmospheric CO2 over the last glacial cycle. *Paleoceanography* 31, 434–452 (2016).
156. Schmitt, J. *et al.* Carbon isotope constraints on the deglacial CO2 rise from ice cores. *Science (1979).* 336, 711–714 (2012).
157. Bird, M. I., Haig, J., Ulm, S. & Wurster, C. A carbon and nitrogen isotope perspective on ancient human diet in the British Isles. *J. Archaeol. Sci.* 137, 105516 (2022).
158. Fernandes, R., Grootes, P., Nadeau, M. J. & Nehlich, O. Quantitative diet reconstruction of a Neolithic population using a Bayesian mixing model (FRUITS): The case study of Ostorf (Germany). *Am. J. Phys. Anthropol.* 158, 325–340 (2015).
159. Pickard, C. & Bonsall, C. Post-glacial hunter-gatherer subsistence patterns in Britain: dietary reconstruction using FRUITS. *Archaeol. Anthropol. Sci.* 12, 142 (2020).
